# Principles of *in situ* protein sequencing: expansion microscopy-adapted Edman degradation and amino acid recognition

**DOI:** 10.64898/2026.01.29.702630

**Authors:** Camille M. Mitchell-Wang, Sara Z. Tavana, Joanne Z. Peng, Hao Wang, Jiuhan Shi, Chi Zhang, Lilia Evgeniou, Masy Domecillo, Shiwei Mitchell-Wang, Daniel M. Estandian, Alexi G. Choueiri, Evelyn Wong, Sarah Dohadwala, Isken Kenzhebaev, Tay Shin, Nicholas F. Polizzi, Laura L. Kiessling, Edward S. Boyden

## Abstract

The ability to map protein identity, with resolution sufficient to infer interactions, would support analysis of how proteins work together, or malfunction, in biological processes and diseases. Although several emerging technologies aim towards single-molecule protein sequencing, they require proteins to be removed from the nanoscale spatial context of cells and tissues. Expansion microscopy (ExM) has facilitated a diversity of chemical analyses by isotropically separating molecules throughout a specimen after permeation via a charged hydrogel, followed by gel swelling. Here, we adapt key protein sequencing steps - Edman degradation and amino acid recognition - to the ExM gel context. Using testbed peptides in ExM gels, we show that N-terminal amino acids can be recognized over multiple cycles of in-gel Edman degradation. This principle-oriented study demonstrates sequencing chemistry on defined synthetic constructs, rather than endogenous proteins in biological samples. These results establish principles of *in situ* protein sequencing and provide a framework for future *in situ* protein sequencing developments, including the development of higher specificity and affinity amino acid binders.

## Main

Proteins are central to life processes^1^, and their interactions achieve metabolic or signaling outcomes with great efficacy and precision, in a fashion that depends on nanoscale context^2–4^. As just one example, synaptic NMDA receptor signaling can promote neuronal health, whereas extrasynaptic NMDA receptor signaling can result in neuronal death^5^. A strategy to map all protein identities and positions, within intact cellular contexts, ideally at single molecule resolution (e.g., 1 nm or better) so that interactions can be inferred, could enable the generation of novel hypotheses and reveal insights into how proteins interact to mediate essential and disease-related processes. However, existing spatial protein mapping techniques either do not scale to the entire proteome, and/or fall short of single molecule spatial resolution and sensitivity (see **Supplementary Table 1** for a list of spatial protein mapping strategies and their quantitative properties).

To map all protein identities, locations, and putative interactions, ideally one would be able to identify individual proteins, with single molecule resolution (e.g., ∼1 nm or better). To this end, expansion microscopy (ExM) is a tool that involves the chemical anchoring of specific amino acids within a protein to a cell- or tissue-permeating swellable hydrogel; the biological specimen is then chemically softened (e.g., with heat and detergent, or enzymatic, treatment), and then immersed in water, which causes isotropic expansion of the specimen-hydrogel composite^6,7^. The net outcome is that a microscope’s effective resolution is improved by an expansion factor (e.g., ∼4.5-fold, in the original version). ExM has been used in hundreds of experimental studies throughout biology^8^. ExM can be iterated, as a sample can be expanded, then a second hydrogel chemically formed in the space opened up by the first expansion, and finally the specimen expanded again. With novel ExM chemistries that achieve 20-fold expansion in a single step, expansion factors of >100x may be feasible^9–11^. Alternatively, a recent study with a 10-fold expansion protocol followed by super-resolution radial fluctuations (SRRF) imaging was used to separate and resolve a protein’s proteolyzed fragments^12^. Together, these advances place ExM in a nanoscale physical regime motivating its use as a nanotechnological scaffold for *in situ* protein sequencing.

Many efforts are ongoing towards *ex situ* single-molecule protein sequencing. These include using fluorescent N-terminal amino acid binders, or covalent amino acid labels, in conjunction with iterative amino acid removal strategies (e.g. Edman degradation). These technologies operate upon proteins *ex situ*, extracted from the spatial context of intact biological cells and tissues^13–17^. If proteins throughout an expanded cell or tissue were fragmented into pieces, and the pieces separated from each other (**Fig. 1A**), then one could in principle walk down each peptide fragment, amino acid by amino acid using Edman degradation (**Fig. 1B**), applying N-terminal binders (or covalent amino acid labels) to such peptide fragments to identify each N-terminal amino acid in turn (**Fig. 1C**). Finally, computational methods could assemble these sequences into an overall protein identity or even sequence. ExM would support such an effort in at least two ways: it would provide the magnification needed to resolve individual peptide fragments (perhaps assisted by further super-resolution imaging), and would enable peptides to be surrounded by a well-defined chemical environment appropriate for protein sequencing (and to allow them to be more easily followed over the many rounds of imaging required for sequencing, as previously demonstrated for nucleic acids^18–20^).

**Figure 1.**
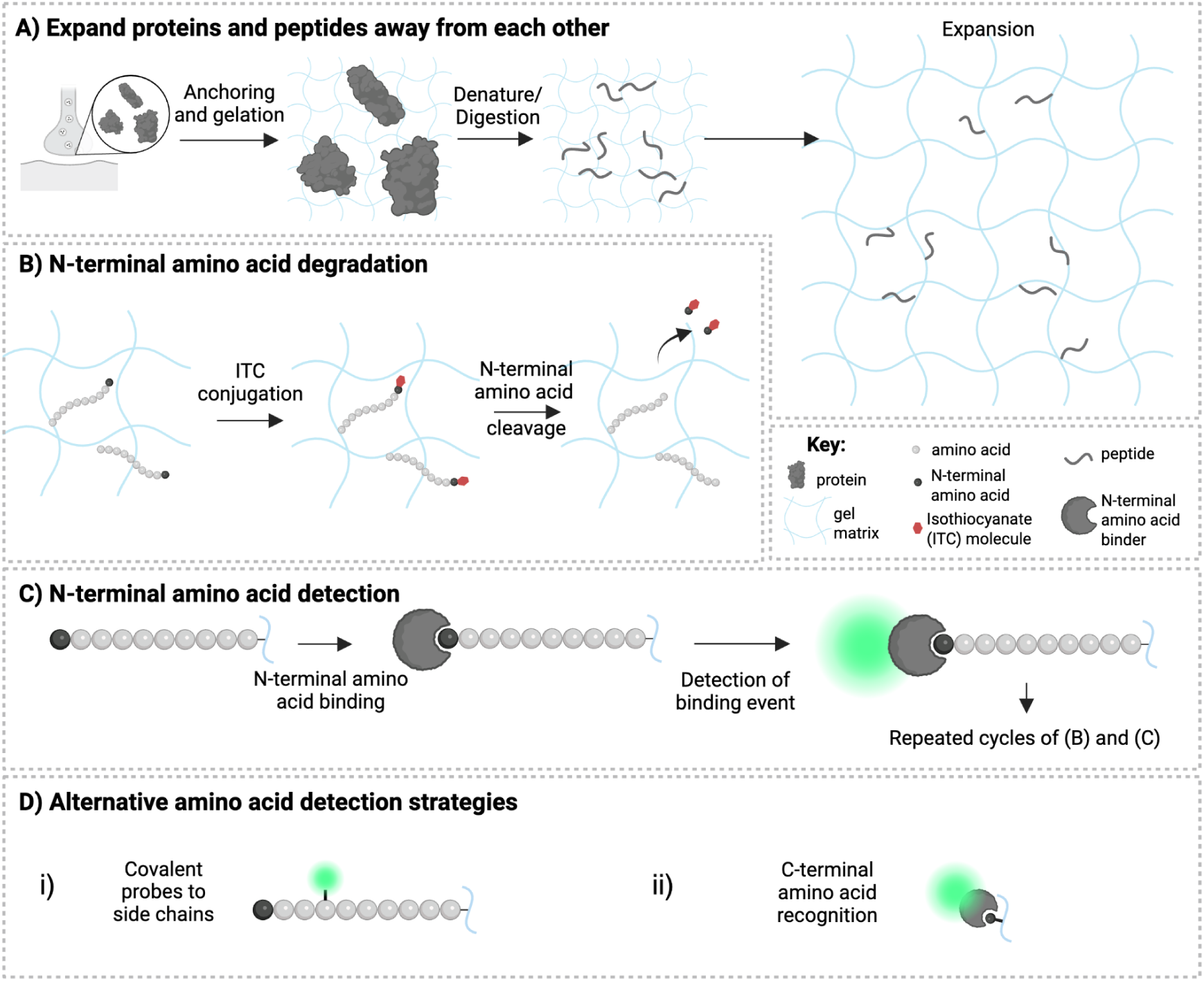
In-gel Edman degradation concept: a chemistry platform to support the framework for *in situ* protein sequencing technology development and application. (A) Expansion of the gel network to separate proteins and protein fragments, prior to peptide sequencing, exploits expansion microscopy (ExM)-style processing of a biological sample (e.g., a cell), including covalent anchoring key amino acids (e.g., lysine side chain modified with 6-((acryloyl)amino)hexanoic acid (AcX), in the original proExM protocol^6,7^), gelation, denaturation and/or digestion of proteins, and expansion, followed by re-embedding of ExM gels in charge-neutral gels to stabilize them in the expanded state (not shown). (B) Edman-in-gel chemistry can be conducted via conjugation of an isothiocyanate (ITC) derivative (e.g., phenylisothiocyanate (PITC)) to N-terminal amino acids of protein fragments or peptides, followed by cleavage of this N-terminal amino acid (e.g., with trifluoroacetic acid (TFA)). (C) The N-terminal amino acid at the end of a protein fragment or peptide can be detected with an N-terminal amino acid binder (e.g., ClpS2 St-V1^25,26^). The binding event is monitored (e.g., with antibodies with or without signal amplification, depicted as a green fluorescent signal next to the binder), before washing away of the binder and a new round of N-terminal amino acid degradation (B). Steps (B) and (C) are performed iteratively to sequentially identify the amino acids of the protein fragment or peptide. (D) Alternative amino acid detection strategies are depicted. (i) One alternative strategy consists in using covalent probes that target sidechains of amino acids^15,32–39^. (ii) Another alternative strategy consists in detecting amino acids conjugated to a modified Edman reagent with their C-termini exposed.

Edman degradation removes amino acids one at a time using a two-step process. First, phenylisothiocyanate (PITC) reacts with the N-terminal amino acid, and then trifluoroacetic acid (TFA) results in cyclization followed by cleavage^21^, at an efficiency of up to ∼99% per round^22^. *Ex situ* single molecule protein sequencing using Edman degradation is an active area of research, e.g., through covalent fluorescent tagging of amino acids^15^, or the application of N-terminal amino acid binders (NAABs)^14,16,23^. Here, we adapt Edman degradation chemistry to ExM to afford an in-gel Edman degradation protocol operating within a nanostructured hydrogel environment. We systematically evaluated solvents, Edman reagents, and reaction conditions to optimize isothiocyanate (ITC)–peptide conjugation in ExM gels. To optimize the chemical reactions that underpin the process, we analyzed testbed peptides with well-defined sequences in gels, analogous to the path taken by early *in situ* nucleic acid sequencing^24^. We demonstrated that we can evaluate N-terminal amino acid binders as they bind to iteratively exposed N-terminal amino acids of testbed peptides. The creation of such binders is an active area of interest for *ex situ* protein sequencing^14,23,25–31^, and further research in this space will be required for a full protein sequencing protocol to be developed. However, the principles revealed here define a nanomaterial-enabled physical regime in which *in situ* protein sequencing becomes feasible.

## Results

### Solvent considerations for in-gel Edman degradation

In-gel Edman degradation is a version of Edman degradation that requires anchoring of peptides via some functional groups (akin to solid phase sequencing), for instance through carboxyl groups as performed in *ex situ* protein sequencing^15,40^ or via the primary amine groups, as performed traditionally in ExM^7^. The sequencing of peptides could be performed on neo N-termini generated by selective peptide cleavage after denaturation/digestion, enabling sequencing of many peptides in parallel (see **Supplementary Note 10** and **Supplementary Figure 13C** for modeled exploration of the effect of anchoring and digestion efficiency on the percent of amino acids that are detected). To adapt Edman degradation to the ExM gel, we had to first discover a solvent that was compatible with gels remaining in the expanded state, while enabling PITC conjugation to peptides. The overall chemical reaction architecture for in-gel Edman chemistry is schematized in **Fig. 2A** (background information on Edman degradation, is in **Supplementary Note 1**, with reagents listed in **Supplementary Table 2**). We cast ExM gels by reacting sodium acrylate, acrylamide, and N,N’-methylenebisacrylamide (BIS, at a slightly high concentration to ensure gel robustness in subsequent steps, resulting in an expansion factor of 3.5x), in a fashion that would either incorporate gel-anchored peptides, or that would omit peptides, as a control. We then cast an acrylamide gel throughout the expanded gel (resulting in a re-embedded ExM gel, or ExMre for short), to stabilize it (at 2.5x expansion, increasing back to 3.5x when exposed to water) and reduce shrinkage under varying solvent and salt compositions (PBS causes ExMre gels to shrink to 2.7x, vs. 1.8x for native ExM gels; **Fig. 2B**; see **Supplementary Note 2** for information on the ionic strength of these solutions), as used previously for *in situ* RNA sequencing^19^.

**Figure 2.**
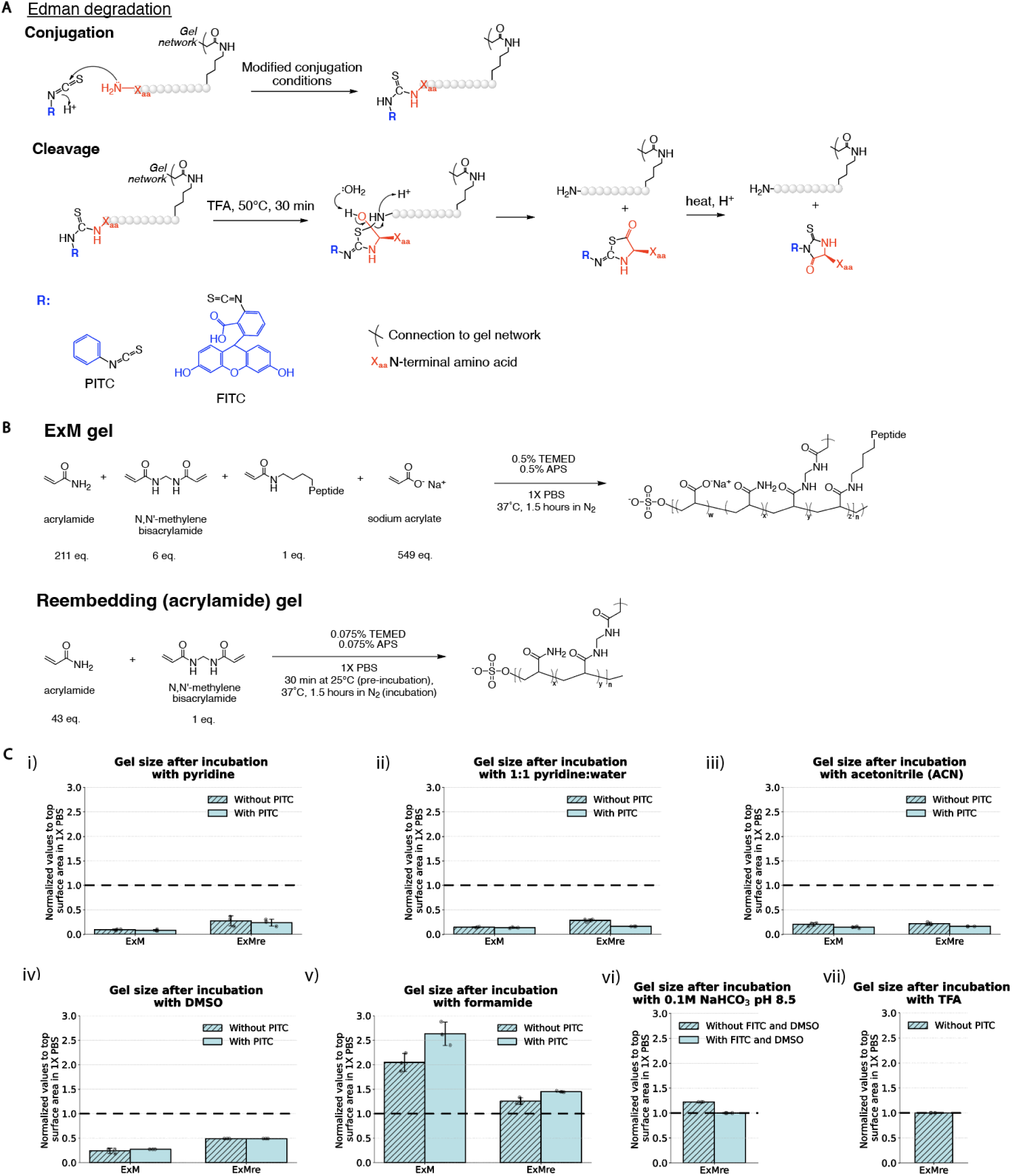
Edman degradation: chemical strategies for in-gel adaptation, and initial characterization of gel compatibility. (A) Proposed reaction scheme of Edman degradation within an ExM gel. To develop conditions, specific peptides were anchored to the gel. The sequencing chemistry includes conjugation of an isothiocyanate (ITC) molecule (i.e., Edman reagent) to the N-terminal amino acid (top). Edman degradation can be performed with a variety of isothiocyanates (denoted “R”, and depicted in blue). The conjugated N-terminal amino acid (denoted “Xaa”, and depicted in red) is then cleaved with TFA, eliminating the thiohydantoin amino acid derivative. (B) Reaction scheme for ExM, and re-embedding of ExM gels. Proteins will be anchored to the gel and then pulled apart from each other. We focus on using synthetic peptides with a C-terminal acryloyl functional group for covalent conjugation to the gel during free-radical polymerization. TEMED, tetramethylethylenediamine; APS, ammonium persulfate. (C) Flat/top side surface size of ExM and ExM re-embedded (ExMre) gels when placed in solvents used in Edman degradation: (i) 100% pyridine, (ii) 1:1 pyridine:water (all ratios throughout are of volumes added, unless otherwise indicated), (iii) acetonitrile (ACN), (iv) dimethylsulfoxide (DMSO), (v) formamide, (vi) 0.1 M sodium bicarbonate (NaHCO_3_). pH 8.5, (vii) trifluoroacetic acid (TFA), with and without Edman reagents (PITC 1:9 ratio PITC:solvent for (i), PITC 1:1000 PITC:solvent for (ii-v); FITC, 5.9 mM, and 23:77 DMSO:0.1 M sodium bicarbonate buffer pH 8.5, for (vi)). (Black dashed line: top/flat surface size of the gel in PBS; error bar: standard deviation; black dots, individual experiments; n=3 separate gelation solutions for (ii-vii); n=3 gels with same starting gelation solution for (i).)

To understand how the reaction conditions might influence the gel, we quantified the flat/top side surface of the gels with a ruler, after placing PBS-washed gels in various PITC conjugation solvents for 30 min at 50 °C, followed by adding PITC (30 min at 50 °C). Pyridine and acetonitrile (ACN) are solvents commonly used for PITC conjugation that resulted in gel shrinkage, and in some cases opacity, which could prevent chemical access to the inside of the gel (**Fig. 2Ci-iii**; see **Supplementary Figure 1** and **Supplementary Figure 2** for images and quantifications of gels cited throughout **Fig. 2**; see **Supplementary Table 4** for full data and statistics for **Fig. 2**), alone or supplemented with PITC (either at 1:9, or 1:1000, which we also found to suffice; **Supplementary Figure 4**). Many other solvents also turned the gel opaque and/or shrank gels (see **Supplementary Figure 1G** for images). We reasoned that solvents not previously used in Edman degradation, such as dimethylsulfoxide (DMSO) and formamide, might solubilize PITC while being compatible with hydrophilic ExM gels. Such solvents did not cause such severe gel shrinkage (**Fig. 2Civ-v**), alone or with PITC. FITC, a nontraditional Edman reagent that is more water soluble, administered in DMSO with sodium bicarbonate led to no change in gel flat/top surface size (**Fig. 2Cvi**), suggesting these conditions might be useful in the first Edman degradation step. Finally, TFA, needed in the cleavage step, did not shrink ExMre gels (**Fig. 2Cvii)**. Thus, we could find solvents compatible with Edman degradation conditions in expanded gel states.

Achieving a full *in situ* protein sequencing platform could benefit from expansion factors on the order of >100x, in order to separate peptides or proteins (or peptides cleaved from the same protein, if desired) so as to reach molecular (<1 nm) resolution on a conventional confocal microscope. The exact expansion factor required, of course, would depend on the imaging modality used: for example, 10x expansion plus SRRF super-resolution could separate peptides that were cleaved from a parent protein (Shaib et al. 2024) (see **Supplementary Table 12** for how various other modalities could be combined to reach this resolution). To this end, while not the focus of the current study, we demonstrated a proof-of-concept achieving high expansion factors >100x. By casting a ∼13-16x gel based on a previously published ExM protocol^11^, re-embedding it using a cleavable N,N’-diallyl-L-tartardiamide (DATD) gel^9,41,42^, casting a second expanding gel, and cleaving the intermediate DATD gel, we were able to then expand the final gel by ∼10-12x to achieve this target expansion factor (∼130-190x; see **Supplementary Figure 22** for the outline of this strategy and the results of this high-expansion gel protocol; see **Supplementary Note 11** for the protocol). Future work will be needed to validate distortion, resolution, and *in situ* yield of such high-expansion gels.

### Testbed peptides anchored throughout ExMre gels for validation of in-gel Edman degradation

Using testbed peptides, we devised a set of mass spectrometry and fluorescent measurements to characterize the efficacy of Edman reagent conjugation and subsequent cleavage. For mass spectrometry, we used a peptide that could be trypsinized from the gel to enable analysis via liquid chromatography coupled with electrospray ionization quadrupole time-of-flight mass spectrometry (or LC/QToF for short) (**Fig. 3A**; see **Supplementary Figure 12** for discussion of the control peptide).

**Figure 3.**
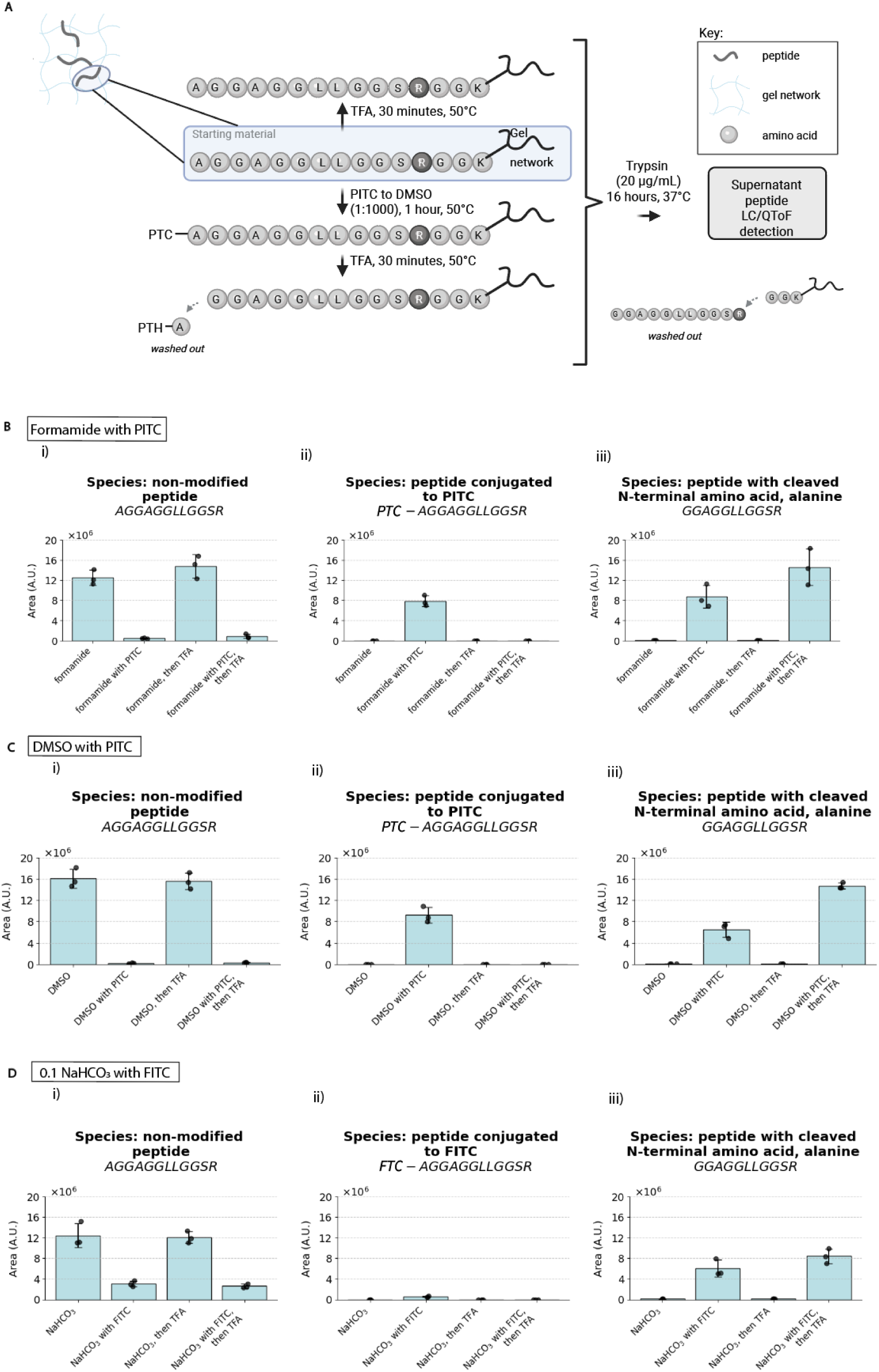
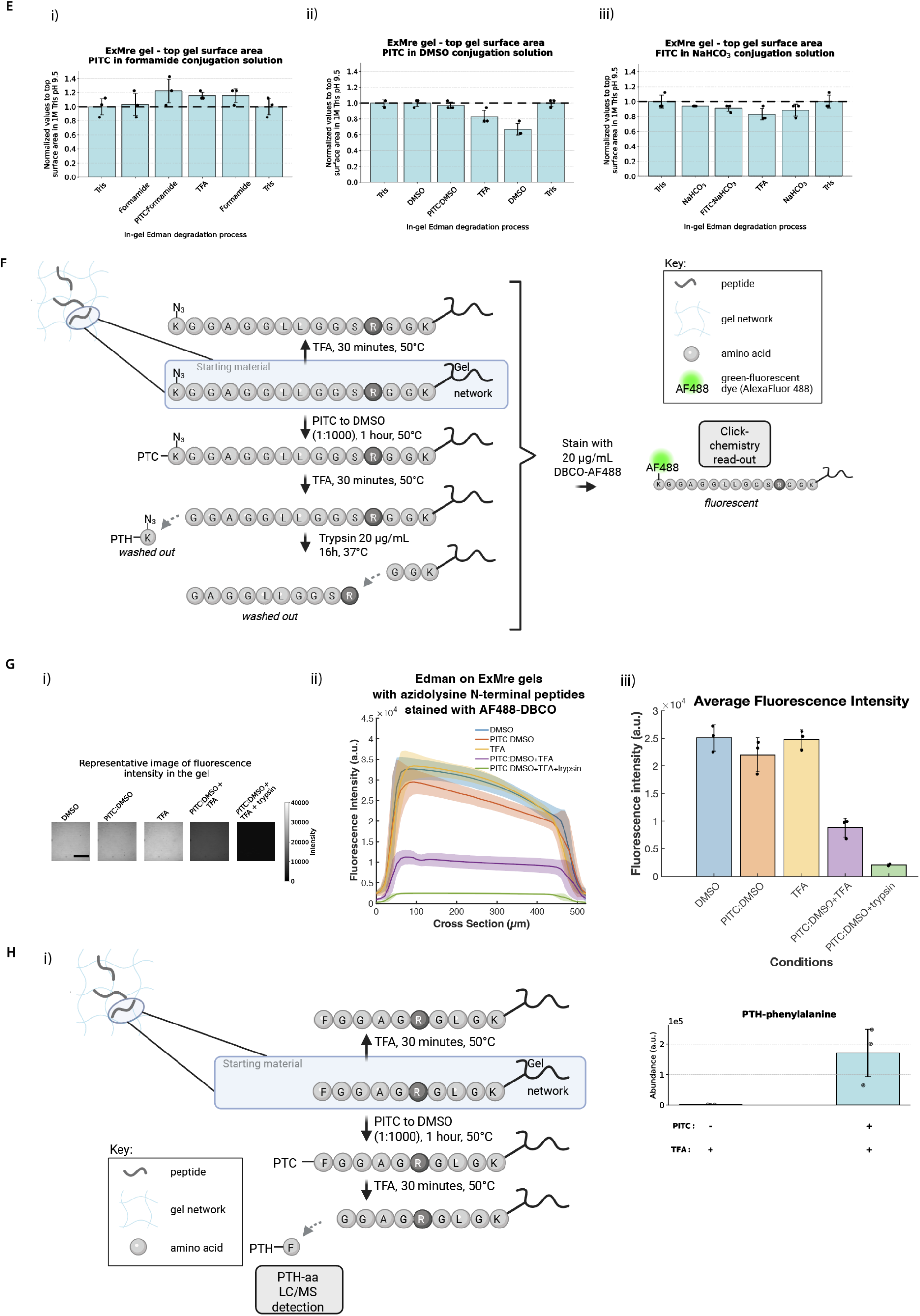
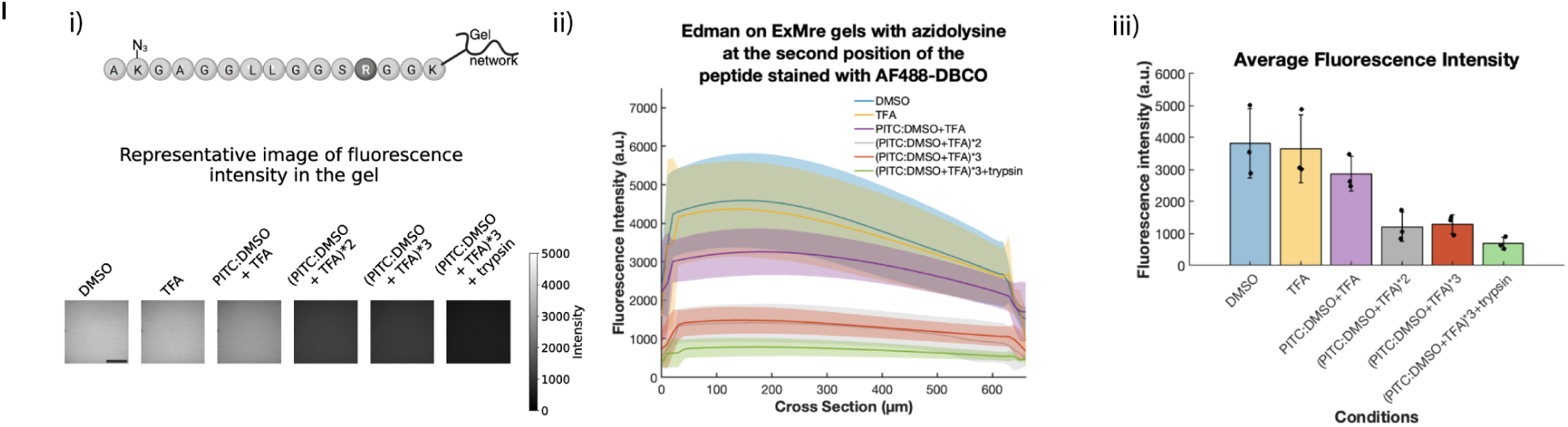
Assessing Edman degradation compatibility on peptides in ExMre gels using LC/QToF trypsinization assay, SPAAC fluorescence assay, and PTH-aa detection. (A) Assay to assess efficiency of Edman reagent conjugation on synthetic peptides in ExMre gels. Synthetic peptide AGGAGGLLGGSRGGK{acr} (abbreviated as A15-peptide; K{acr} denotes an acryloyl functional group on the lysine side chain); amino acids of the peptide chain are depicted as light grey beads, with arginine in the dark grey bead, as site of trypsin cleavage. Peptides are embedded into ExMre gels during free-radical polymerization of the swellable gel, and separate gels are subjected to various in-gel Edman conditions. All ExMre gels are then immersed in trypsin for digestion overnight to acquire the fragment upstream to the trypsin cleavage site at arginine, for further analysis. Supernatant is analyzed with liquid chromatography coupled with electrospray ionization quadrupole time-of-flight mass spectrometry (LC-ESI-QToF MS, abbreviated LC/QToF). (B) In-gel Edman chemistry with formamide as conjugation solvent and PITC as the Edman reagent. Bar graphs representing the relative abundance (arbitrary units, a.u., with all samples processed with spiked-in control compounds; see **Methods** and **Supplementary** Figure 12) of different peptide ion species detected on the LC/QToF. Bar graphs were obtained from measuring the area under the curve (AUC) of the chromatogram of various species (extracted based on the exact mass, see **Methods** for details, and **Source Data** for raw traces). The separate conditions were solvent only (“formamide”), PITC to solvent (1:1000 ratio PITC:solvent) for 1 hour at 50 °C (“formamide with PITC”), TFA for 30 min at 50 °C (“formamide, then TFA”), PITC to solvent (1:1000 ratio PITC:solvent) for 1 hour at 50 °C followed by TFA for 30 min at 50 °C (“formamide with PITC, then TFA”). The relative abundance of the ion species: (i) non-modified peptide (AGGAGGLLGGSR), (ii) peptide conjugated to PITC, phenylthiocarbamyl (PTC)-peptide (*PTC*-AGGAGGLLGGSR), and (iii) peptide with cleaved N-terminal amino acid (GGAGGLLGGSR), were reported throughout the in-gel Edman degradation process in the various conditions (dots, individual experiments; blue bar, mean; error bar, standard deviation, n=3 separate gelation solutions). Conversion rates were obtained by comparing conditions: 1- (solvent with PITC[“A15-peptide”] / solvent [“A15-peptide”]), and details can be found in **Supplementary Table 5**. (C) In-gel Edman chemistry as in B, but with DMSO as solvent. (D) In-gel Edman chemistry as in C, but with 23:77 of DMSO:0.1 M sodium bicarbonate pH 8.5 as conjugation solution with FITC (5.9 mM) as Edman reagent. The peptide conjugated to FITC is fluorescein-thiocarbamyl (FTC)-peptide (*FTC*-AGGAGGLLGGSR). (E) Top/flat surface size of ExMre gels throughout the Edman degradation process. Normalized top/flat surface size at each step for ExMre gels with (i) formamide conjugation with PITC, (ii) DMSO conjugation with PITC, or (iii) 5.9 mM FITC in 23:77 DMSO:0.1 M NaHCO_3_ pH 8.5 (thick dashed black line, top/flat surface size of the gel after washes in 1 M Tris pH 9.5 (1 mL x 3) before Edman degradation; error bar, standard deviation; black dots, individual experiments; n=3 separate gelation solutions). (F) Assay to assess efficiency of Edman degradation on synthetic peptides in ExMre gels. Synthetic peptide: K{N_3_}GGAGGLLGGSRGGK{acr} (abbreviated K{N_3_}15-peptide, where K{N_3_} is 6-azido-lysine), and amino acids of the peptide chain are depicted as light grey beads, with arginine, “R”, with the dark grey bead, as site of trypsin cleavage. The peptides are embedded into the ExMre gels during free-radical polymerization of the first gel, and separate gels are subjected to various Edman conditions. Read-out is performed using strain-promoted alkyne-azide cycloaddition (SPAAC) with dibenzocyclooctyne AlexaFluor 488 (DBCO-AF488) on the embedded peptides followed by analysis of the fluorescence. Note: sloped intensity profiles were likely due to excitation light attenuation in deeper layers of highly fluorescent gels. (G) ExM gels containing K{N_3_}15mer-peptide at 1 mM were cast in a gelation chamber, expanded and re-embedded to reach ∼2.7X expansion factor at a final peptide concentration of ∼50 μM. The separate conditions were DMSO only (“DMSO”), PITC to DMSO (1:1000 ratio PITC:DMSO; “PITC:DMSO”), DMSO followed by TFA, PITC to DMSO (1:1000 ratio PITC:DMSO) followed by TFA, and the same condition but with 20 ug/mL trypsin (“PITC:DMSO+TFA+trypsin”). After SPAAC with 20 μg/mL DBCO-AF488 in PBS, bulk gel fluorescence was imaged using a confocal microscope with a 10 μm Z-step. Analysis was performed on raw images. See Methods for **Yield calculations**. (i) A representative raw image of the fluorescence intensity is depicted for each gel condition, taking the 20th slice of the Z-stack for each (∼200 μm deep into the gel). Scale bar is 500 μm. (ii) The fluorescence intensity of the ExMre gels in different conditions was compared throughout the gel thickness (∼500 μm) (line, mean; shaded area, standard deviation; n=3 separate gelation solutions). (iii) Average fluorescence intensity of ExMre gels in the different conditions across the whole volume imaged (colored bar, mean; black dots, individual experiments; error bar, standard deviation, n=3 separate gelation solutions). (H) (i) Independent assay for in-gel Edman degradation, with phenylthiohydantoin (PTH)-F detection of Edman degraded synthetic peptide: FGGAGRGLGK{acr} (abbreviated “F_1_ peptide”) embedded in ExMre gels, as in (A). Separate gels were subjected to various Edman conditions. The conditions included TFA only for 30 min at 50 °C, and PITC to DMSO (1:1000 ratio PITC:DMSO) for 1 hour at 50 °C followed by TFA for 30 min at 50 °C. Subsequently, TFA was removed from the gels and they were immersed in 50 μL of 1:1 acetonitrile to water and agitated. Read-out was then performed by injecting the supernatant into LC/QToF using **LC Method (see Methods)**. (ii) Results acquired as in (i). Analysis of PTH-F abundance was performed using PTH-F exact mass, 282.0827 ± 0.0056 Da (see **Methods for Edman degradation and PTH detection** for details). (blue bar, mean; error bar, standard deviation; black dots, individual experiments; n=3 separate gelation solutions). (I) ExM gels containing AK{N_3_}15mer-peptide at 1 mM were cast in a gelation chamber, expanded and re-embedded to reach ∼2.7X expansion factor at a final peptide concentration of ∼50 μM. The separate conditions were DMSO only (“DMSO”), TFA only (“TFA”), PITC to DMSO (1:1000 ratio PITC:DMSO) followed by TFA performed over one, two or three cycles (“(PITC:DMSO+TFA*3)+trypsin”), and the same condition but with 20 ug/mL trypsin (“(PITC:DMSO+TFA*3)+trypsin”). After SPAAC with 20 μg/mL DBCO-AF488 in PBS, bulk gel fluorescence was imaged using a confocal microscope with a 10 μm Z-step. Analysis was performed on raw images. See Methods for **Yield calculations**. (i) The AK{N_3_}15mer-peptide with azidolysine at the second position of the peptide is depicted at the top. The representative raw image of the fluorescence intensity is depicted for each gel condition, taking the 10th slice of the Z-stack for each (∼100 μm deep into the gel). Scale bar is 100 μm. (ii) The fluorescence intensity of the ExMre gels in different conditions was compared throughout the gel thickness (∼660 μm) (line, mean; shaded area, standard deviation; n=3 separate gelation solutions). (iii) Average fluorescence intensity of ExMre gels in the different conditions across the whole volume imaged (colored bar, mean; black dots, individual experiments; error bar, standard deviation, n=3 separate gelation solutions).

We note that the phenylthiocarbamyl (PTC)-peptide bond resulting from Edman reagent conjugation is labile under preparation conditions for mass spectrometry (see **Supplementary Note 3** for more detail about this limitation), and thus we focus our analysis on the efficacy of ITC conjugation. As expected, conventional Edman solvents that caused gel shrinkage or opacity did not result in good conjugation (see **Supplementary Note 4**). Formamide and DMSO, in contrast, enabled excellent conversion to the PTC-peptide with a conversion rate of ∼96% for 1:1000 ratio PITC:formamide (**Fig. 3B**; see **Supplementary Table 5** for full data and statistics related to **Fig. 3**) and a conversion rate of >99% for 1:1000 PITC:DMSO (**Fig. 3C**). FITC in 0.1 M sodium bicarbonate resulted in a lower conversion rate of ∼75% (**Fig. 3D**), as expected from the lower FITC conjugation rate reported in the literature^43^. All three protocols (**Fig. 3E**), could be conducted without substantial gel shrinkage.

To assess phenylthiohydantoin release, we devised a strain-promoted alkyne-azide cycloaddition (SPAAC) click chemistry fluorescence experiment (**Fig. 3F)**. We used an N-terminal azidolysine peptide that was clicked to a dibenzocyclooctyne (DBCO) fluorophore after various chemical treatments. When the gel was exposed to our Edman degradation conditions, we observed the expected drop in fluorescence due to cleavage (**Fig. 3Gi-ii**), about ∼70% of the total (**Fig. 3Giii**; trypsin-mediated positive control). We then performed the same experiment on a peptide with an azidolysine on the second amino acid residue, exposing it to three rounds of in-gel Edman degradation. We observed a sizable drop in fluorescence after the second round of in-gel Edman degradation, with an observed ∼83% yield when using trypsin as a positive control, suggesting a per-round degradation yield of ∼90%. This variability in observed yield likely reflects both the limited number of replicates (n=3) and the sensitivity of the degradation chemistry to small differences in reagent purity or protocol execution, and should be further characterized with larger sample sizes in future work, after further optimization to higher % yield, that would justify the focus on attaining large confirmatory sample sizes. We further tested covalently labeling peptides with Atto647N-alkyne dye, known to withstand Edman degradation solvents^15^, prior to in-gel Edman degradation conditions using the azidolysine labeled peptides above. Using similar controls, the results of the N-terminal labeled azidolysine peptide suggested ∼93% yield over one round (**Supplementary Figure 6Fi-iii**). For the peptide with azidolysine at the second position, we observed a yield of ∼74% over two rounds, suggesting a per-round yield of ∼86% (**Supplementary Figure 6Gi-iii**). Thus, we consistently obtained per-round yields in the 70%-90% range, as measured by a range of methods. These per-round yields are comparable to those reported by other recently developed modified sequencing chemistries, including a novel alternative to Edman degradation that reported per-residue yields of 60–93% ^44^. We further explored whether the cleaved phenylthiohydantoin (PTH)-amino acid could be detected directly via LC/QToF (**Fig. 3Hi**), and confirmed PTH-amino acid release under Edman conditions (**Fig. 3Hii**; see **Supplementary Figure 6** and **Supplementary Note 5** for results regarding other amino acids).

### Combining in-gel Edman degradation with bulk N-terminal amino acid detection

We set out to see if an existing N-terminal amino acid binder (NAAB) could detect N-terminal amino acids in the expansion gel. An ideal NAAB would have high affinity and specificity, and a slow rate of dissociation - slow enough to support amplification (e.g., by attaching many fluorophores through hybridization chain reaction (HCR) or enzymatic means^19,45^, which takes minutes to hours) so that ordinary microscopes could be used to identify individual amino acids of individual peptides. Existing NAABs do not yet reach these criteria, with poor affinity, dwell time, and with affinity dependence on the downstream amino acids of the peptide. Making better NAABs is an active area of research^14,23,25–30^, and will likely benefit from efforts in immune and display-based selection, structure based design, and/or AI-driven modeling and design^46^.

We chose as our NAAB *Agrobacterium tumefaciens* ClpS2, engineered for higher specificity towards Phe (ClpS2 St-V1)^25^ (see **Supplementary Note 6** for thoughts on affinity and sequence information). We measured bulk fluorescence throughout the gel upon applying ClpS2 St-V1 with a fused HA tag, followed by addition of a fluorescent anti-HA tag antibody, using ExMre gels equipped with peptides with N-terminal Phe, Trp, Tyr, and Ala (**Fig. 4A**). We discuss some assumptions and calculations to help interpret images amidst fast binder off kinetics, in **Supplementary Note 7**. We observed high fluorescence for the Phe-bearing peptides, and not the other three (**Fig. 4B-D**), for instance with an ∼8 fold difference comparing Phe to Tyr (for all raw data for **Fig. 4** see **Supplementary Table 9**) comparable to our calculated estimate mentioned above (∼7 fold difference comparing Phe to Tyr, see **Supplementary Note 8** for calculations). Having established that N-terminal binding was possible, we next sought to test the incorporation of N-terminal binding into a sequencing pipeline. We created gels with peptides bearing Phe on the N-terminus (denoted F_1_ peptide) or in the second position (denoted G_1_F_2_ peptide), so that the NAAB would bind only at the appropriate stage in sequencing (**Fig. 4Ei**-**ii**). We observed the NAAB to bind F_1_ peptide only in cases when the N-terminal Phe was still remaining (e.g., the DMSO and TFA-only cases), and not when it was occluded (after PITC conjugation) or cleaved (PITC followed by TFA, or the trypsin control) (**Fig. 4F**). In contrast, for G_1_F_2_ peptide, we only saw NAAB binding when the Phe was exposed after Edman degradation (PITC followed by TFA), and not under conditions where it was occluded by the N-terminal amino acid (DMSO, TFA-only) or by that amino acid bearing PITC, or after trypsin elimination of the peptide (**Fig. 4G**). In summary, we were able to demonstrate, in the interest of principles of in situ protein sequencing, bulk-level sequencing of defined peptide substrates using NAABs, over cycles of in-gel Edman degradation. Comparing gel fluorescence for Phe in the 1^st^ vs. 2^nd^ position, we observed a ∼50% drop in fluorescence, suggesting a yield comparable to what was estimated from the click chemistry experiment above.

**Figure 4.**
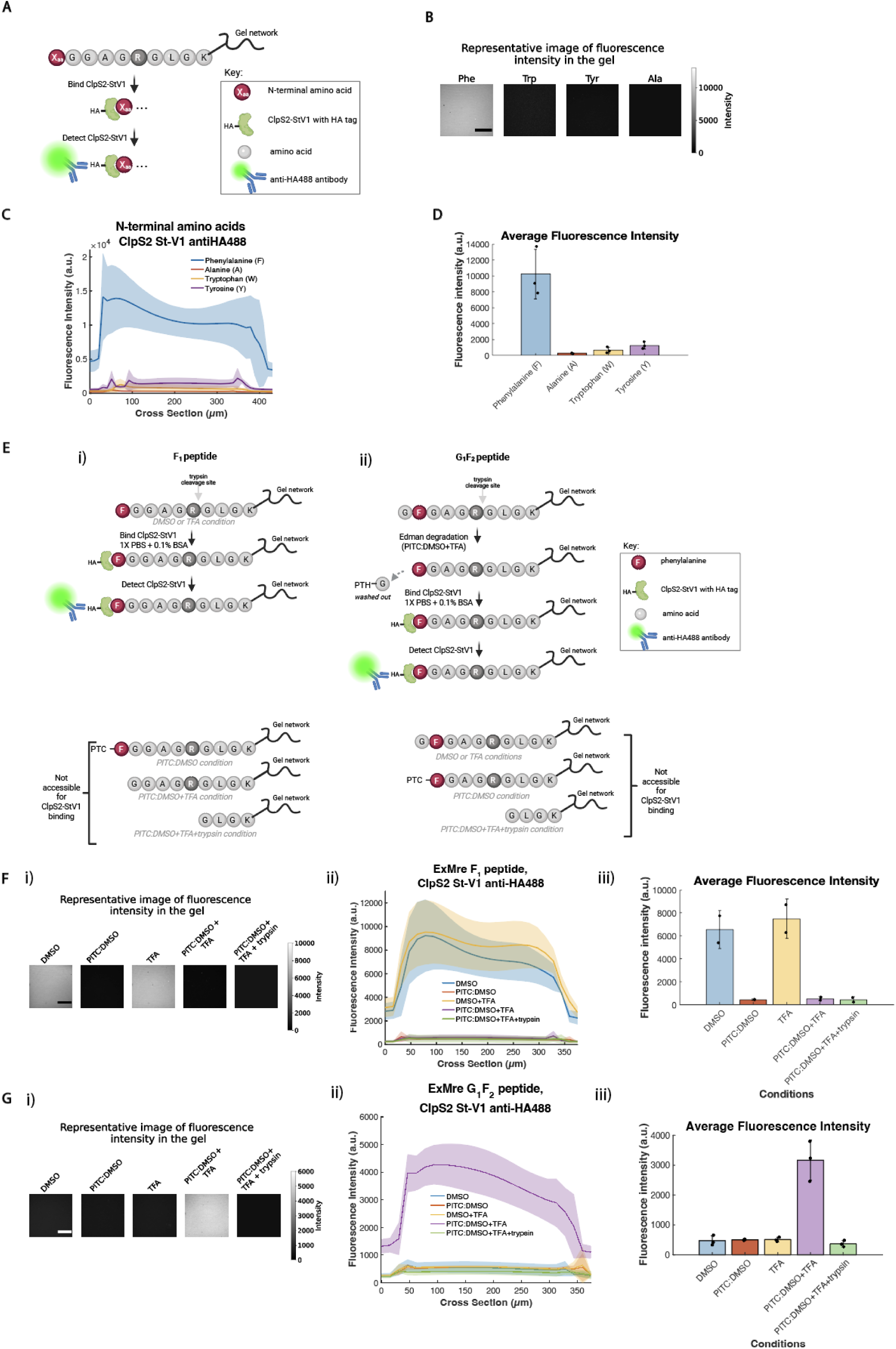
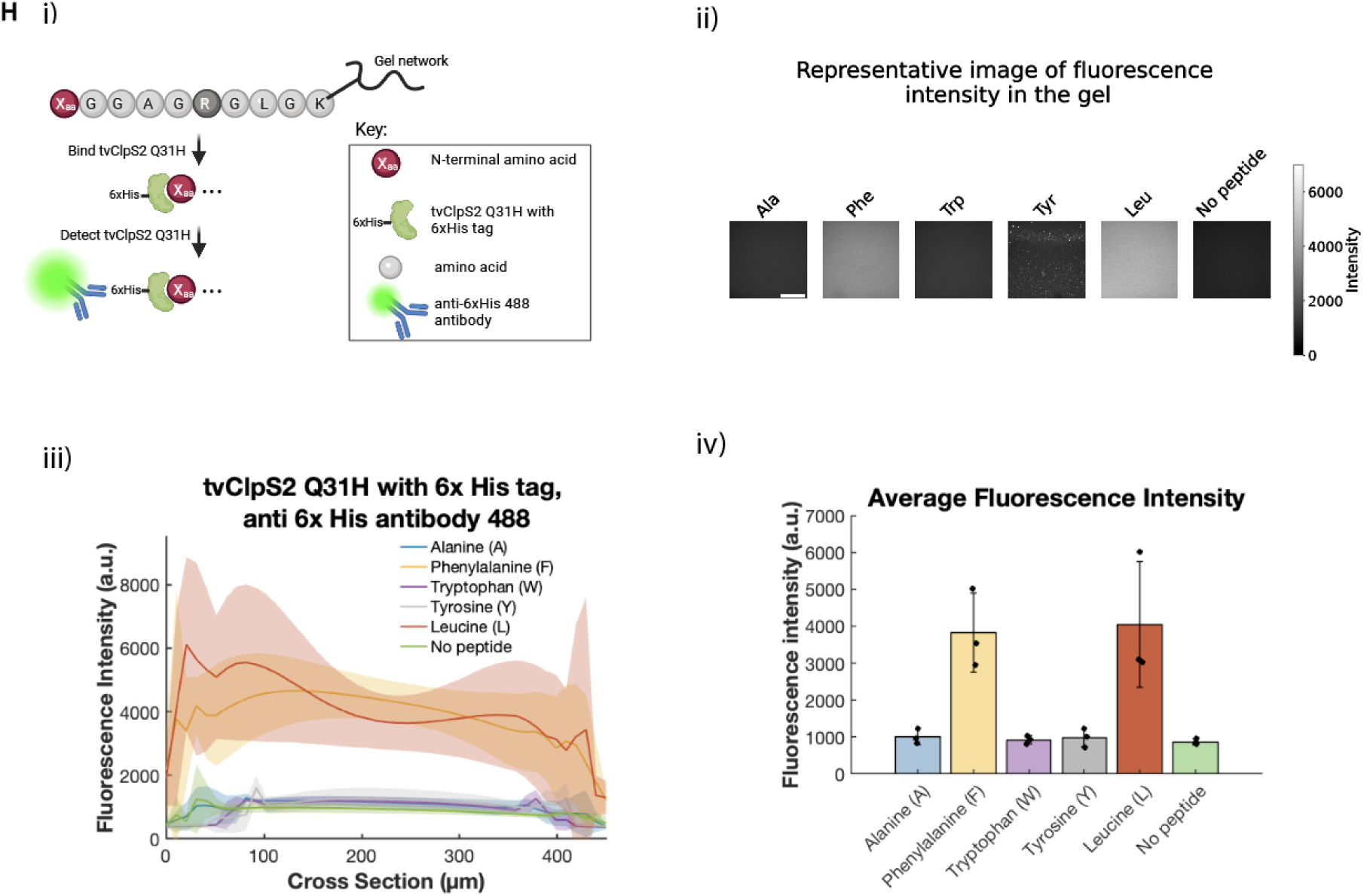
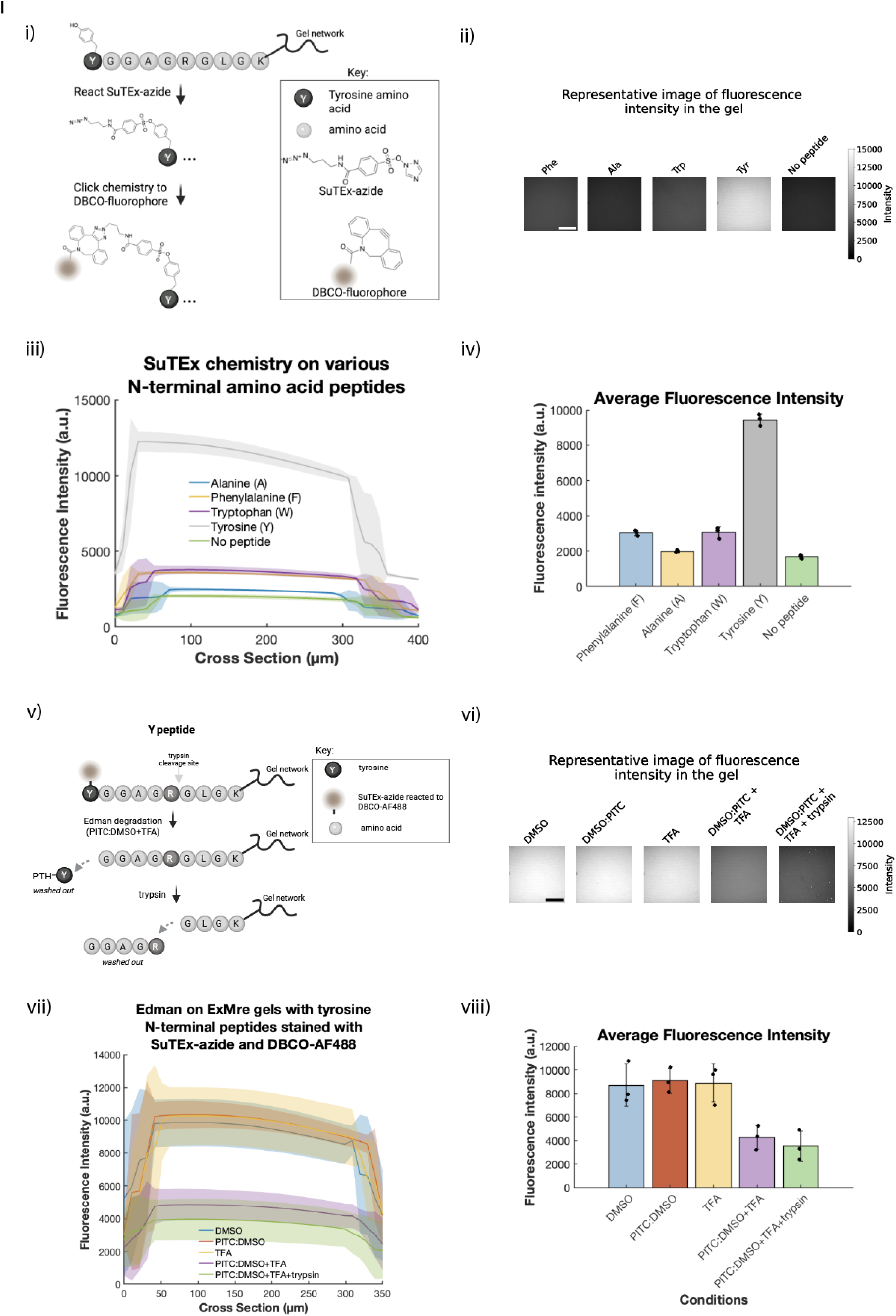
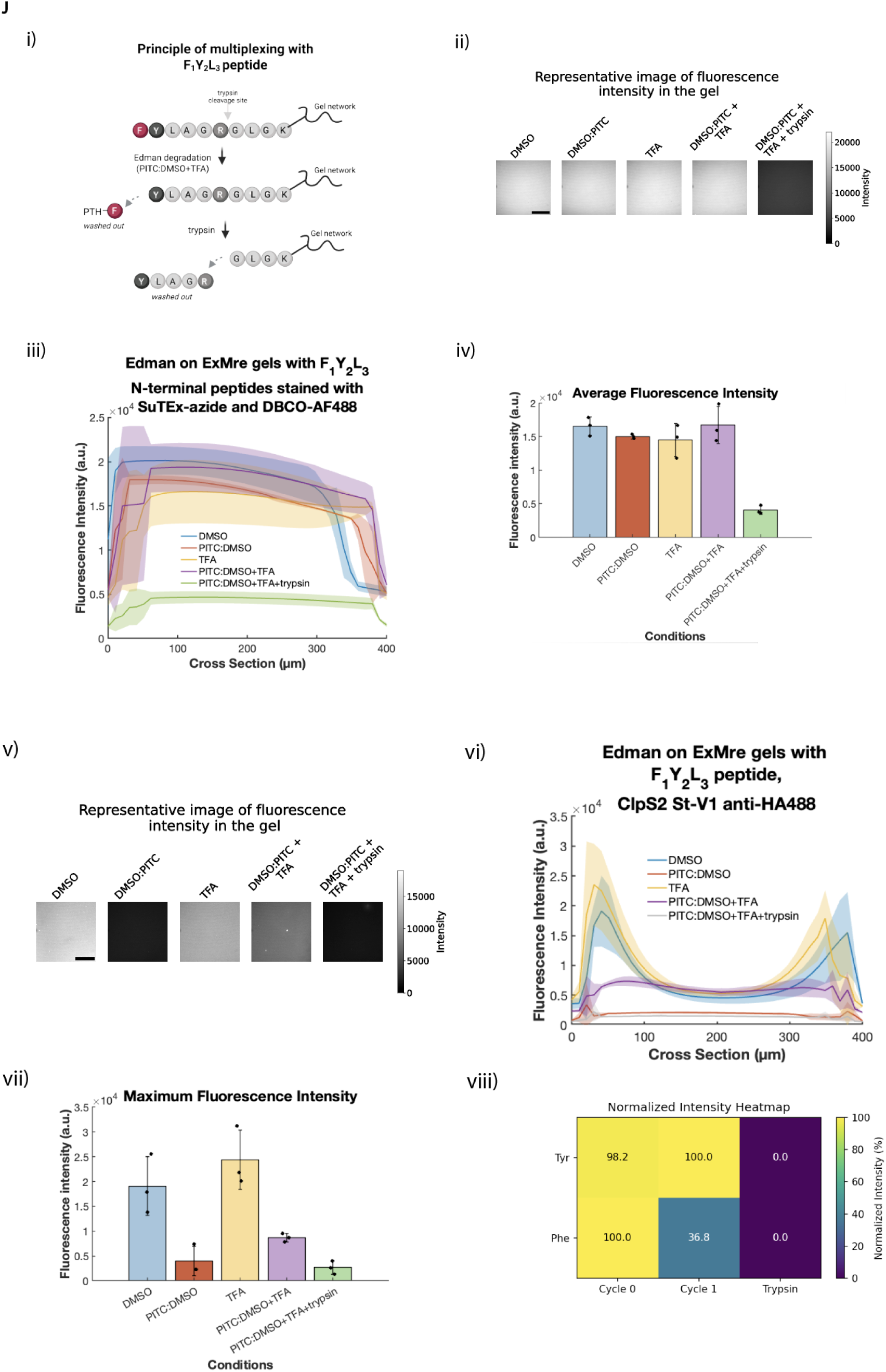
ClpS2 St-V1 serves as read-out for N-terminal phenylalanine on peptides in ExMre gels. (A) Schematic representing different N-terminal peptides with otherwise similar sequence (**Xaa**GGAGRGLGK{acr}, where Xaa can be phenylalanine, alanine, tryptophan, tyrosine). When Xaa is F, abbreviated F_1_ peptide; A, abbreviated A_1_ peptide; W, abbreviated W_1_ peptide; Y, abbreviated Y_1_ peptide. The peptide is depicted as a chain of grey beads, with arginine “R” as the dark grey bead, and the N-terminal amino acid as a red bead. First, 10 μM ClpS2 St-V1 protein with an HA tag is placed in solution to bind peptides in ExMre gels. The protein is briefly washed out, and the gels are stained with anti-HA tag antibody 488. Fluorescence of the gels then reports the relative binding of ClpS2 St-V1 to the various N-terminal peptides via a population read-out using confocal imaging of the ExMre gels. (B) ExM gels containing F_1_ peptide, A_1_ peptide, W_1_ peptide and Y_1_ peptide are separately made at 1 mM concentration cast in a gelation chamber, expanded and re-embedded to reach ∼2.7X expansion factor, reaching a final concentration of ∼50 μM peptide (each gel with a volume of 1.4 μL). ExMre gels were incubated with 10 μM ClpS2 St-V1, washed once, and placed in a solution of 67 nM anti-HA tag antibody 488 in 30 μL solution. The antibody solution is removed right before imaging on a confocal microscope, with a 10 μm step size covering the entirety of the gel. A representative raw image of the fluorescence of the gels is depicted for each ExMre gel containing a different N-terminal peptide, taking the 20th slice of the Z-stack for each (∼200 μm deep into the gel). In black: scale bar, 100 μm. (C) Cross-section of the imaging performed in (B) for the different N-terminal peptides in ExMre gels after staining with ClpS2 St-V1 and anti-HA tag antibody 488. Raw intensity of the gels across the Z-stack were compared (line, mean; shaded area, standard deviation; n=3, with three different peptide aliquots, 3 different gelation mixtures). (D) Average fluorescence intensity of the images obtained in (B) for the different N-terminal peptides in ExMre gels after staining with ClpS2 St-V1 and anti-HA tag antibody 488. Fluorescence intensity of ExMre gels in the different conditions was averaged across the whole volume imaged (black dots, individual experiments; colored bar, mean; error bar, standard deviation, n=3, with three different peptide aliquots, 3 different gelation mixtures). (E) (i) F_1_ peptide, with the F amino acid depicted as a red bead, in ExMre gels. ClpS2 St-V1 staining with anti-HA tag antibody 488 is performed as in (A). (ii) A peptide with sequence GFGAGRGLGK{acr}, abbreviated G_1_F_2_ peptide, where the F amino acid is depicted in red. 1 round of in-gel Edman degradation degrades N-terminal glycine (G) in ExMre gels. Then, ClpS2 St-V1 and anti-HA tag antibody 488 staining is performed as in (A). (F) (i) A representative image of the fluorescence of the gels is depicted for each condition of in-gel Edman degradation for the F_1_ peptide, taking the 14th slice of the Z-stack for each (∼210 μm deep into the gel, with 15 μm z steps). Imaging performed on a confocal microscope (∼375 μm thick Z-stack). Scale bar, 100 μm. (ii) The gels in different conditions are compared in fluorescence intensity throughout their cross-section for the F_1_ peptide (line, mean; shaded area, standard deviation; n=2, different gelation solutions). (iii) Average fluorescence intensity of the ExMre gels in the different conditions for the F_1_ peptide by averaging across the whole volume imaged (black dots, individual experiments; colored bar, mean; error bar, standard deviation, n=2, different gelation solutions). (G) (i) As in Fi, but with G_1_F_2_ peptide, taking the 13th slice of the Z-stack for each (∼195 μm deep into the gel, with 15 μm z steps). (ii) As in Fii, but with G_1_F_2_ peptide (line, mean; shaded area, standard deviation; n=3, different gelation solutions). (iii) As in Fiii, but with G_1_F_2_ peptide (black dots, individual experiments; colored bar, mean; error bar, standard deviation, n=3, different gelation solutions). (H) (i) As in A, schematic representing different N-terminal peptides with otherwise similar sequence (**Xaa**GGAGRGLGK{acr}, where Xaa can be phenylalanine, alanine, tryptophan, tyrosine, leucine). First, 20 μM tvClpS2 Q31H protein with a 6xHis tag is placed in solution to bind peptides in ExMre gels. The protein is briefly washed out, and the gels are stained with 133 nM anti-6xHis tag antibody conjugated to Alexa Fluor 488. Fluorescence of the gels then reports the relative binding of tvClpS2 Q31H to the various N-terminal peptides via a population read-out using confocal imaging of the ExMre gels. (ii) A representative raw image of the fluorescence of the gels is depicted for each ExMre gel containing a different N-terminal peptide, taking the 10th slice of the Z-stack for each (∼100 μm deep into the gel). In black: scale bar, 100 μm. (iii) The gels in different conditions are compared in fluorescence intensity throughout their cross-section (line, mean; shaded area, standard deviation; n=3 different gelation solutions). (iv) Average fluorescence intensity of the ExMre gels in the different conditions (black dots, individual experiments; colored bar, mean; error bar, standard deviation, n=3 different gelation solutions). (I) (i) Schematic representing specific labeling of tyrosine using the SuTEx-azide covalent probe. A stock of 50 mM of the SuTEx-azide probe in ACN is diluted to 3 mM in 1X PBS and 50 μL of this final solution is added to each gel for 2 hours at 37°C. After washing, 20 μg/mL DBCO-AF488 is added to react with the probe for downstream imaging. (ii-iv) As in Hii-iv, but for SuTEx staining. (v) Schematic representing the in-gel Edman degradation on Y_1_ peptide. Peptide is stained after the chemistry is complete using the same reaction as (i). (vi) As in (ii). (vii) As in (iii). (viii) As in (iv). (J) (i) Schematic of the rudimentary multiplexing experiment, involving ClpS2 St-V1 NAAB and SuTEx-azide covalent probe on the F_1_Y_2_L_3_ peptide (sequence: FYLAGRGLGK{acr}). (ii) A representative raw image of the fluorescence of the gels is depicted for each ExMre gel containing a different N-terminal peptide, taking the 20th slice of the Z-stack for each (∼200 μm deep into the gel). In black: scale bar, 100 μm. (iii-iv) As in Hiii-iv, but for SuTEx staining. (v) A representative raw image of the fluorescence of the gels is depicted for each ExMre gel containing a different N-terminal peptide, taking the 8th slice of the Z-stack for each (∼80 μm deep into the gel). In black: scale bar, 100 μm. (vi) As in Hiii-iv, but for ClpS2 St-V1 staining. (viii) Heatmap of the min-max normalized intensities (highest read-out set to 100, minimum read-out set to 0) from the F_1_Y_2_L_3_ peptide before, and after 1 cycle of in-gel Edman degradation, as well as after trypsinization.

We further tested alternative NAABs with different N-terminal specificities, notably a mutated version of *Thermosynechococcus vestitus* ClpS2, abbreviated tvClpS2 Q31H, which exhibits preferential binding to N-terminal leucine, followed by phenylalanine, tyrosine, and tryptophan^29^. Using our platform, we tested the His-tagged tvClpS2 Q31H protein (**Fig. 4Hi)**, where it showed specificity towards N-terminal leucine and phenylalanine, but not tyrosine or tryptophan, or the off-target amino acid alanine and a blank gel (denoted: “no peptide”) (**Fig. 4Hii-iv**). Compared with the previous specificity results, we observed no specific binding to tyrosine or tryptophan, possibly reflecting the effect of the residue at the second position (here a glycine instead of an alanine), which is known to modulate ClpS2 NAAB affinity.

As noted above, single molecule imaging with such binders would not be straightforward on ordinary microscopes, one of our criteria for technological success. For instance, ClpS2 St-V1 exhibits weak affinity (K_d_ of ∼1.1 μM) with a dwell time of ∼10 seconds^14,23,26^. However, single molecule amplification methods used in ExMre gels for read-out with conventional confocal microscopy, such as hybridization chain reaction (HCR) or rolling circle amplification (RCA), require on the order of hours for signal amplification^19,45^. The binder used here would dissociate before any amplification product could be made. Therefore, this NAAB would not be suitable for single molecule detection in an *in situ* setting.

We also set out to test reagents that target non-aliphatic side chains to show the generalizability of our platform beyond non-aliphatic amino acids. However, due to a lack of commercially available NAABs towards non-aliphatic amino acids, we turned towards covalent probes that target reactive functional groups and that exist for up to 11 amino acid side chains^32–39,47^. Therefore, we performed sulfur-triazole exchange (SuTEx) chemistry by testing SuTEx-azide specificity towards tyrosine ^34^ on our platform. We first reacted SuTEx-azide with the embedded peptides, followed by staining with SPAAC click chemistry (**Fig. 4Ii**). Compared to all other N-terminal amino acids, as well as a blank gel (no embedded peptides), our results demonstrated specificity to gels with N-terminal tyrosine (**Fig. 4Iii-iv**).

Using the SuTEx-azide probe, we further tested its ability to monitor degradation of tyrosine from the N-terminus of the peptide after the in-gel Edman degradation chemistry previously developed (**Fig. 4Iv**). The results of this assay demonstrated a consistent fluorescence in the gel in conditions that do not modify the tyrosine side chain, including the PITC conjugation to the N-terminus, but a drop in the fluorescence of the gel when PITC conjugation and TFA cleavage was performed on the peptides (**Fig. 4Ivi-viii**). The results of this assay suggested a ∼ 85% N-terminal amino acid degradation yield (see **Methods: Yield calculations**) of the tyrosine amino acid using trypsin as our positive control, similar to the ∼70-90% per-round yield observed in **Fig. 3G,I** and **Supplementary Figure 6F-G**.

As a rudimentary demonstration of multiplexed amino acid detection, we tested whether we could combine the NAAB read-out for Phe and the covalent probe for Tyr, using a peptide bearing FYL at the N-terminus (denoted F_1_Y_2_L_3_ peptide). After running one round of in-gel Edman degradation with the control conditions, including the trypsinization control, we could assess whether the read-outs correctly reflected the peptide modification and degradation status (**Fig. 4Ji**). Tyr labeling after in-gel Edman degradation showed higher overall fluorescence in all the conditions except the trypsinized peptide, which corroborates with Tyr being at the second position and thus not being degraded away after 1 cycle, but getting cleaved away after trypsinization at the arginine (**Fig. 4Jii-iv**). On the other hand, the fluorescence signal for NAAB binding to Phe was highest in the conditions that do not modify the N-terminus of the peptide, *i.e.* DMSO- or TFA- only conditions. In this case, the signals were higher on the edge of the gels compared to what was observed in **Fig. 4F** on the F_1_ peptide, most likely due to the higher affinity of ClpS2 St-V1 for a peptide with N-terminal Phe and second amino acid Tyr, as reported^26^. Indeed, these peptides would soak up the NAAB more efficiently before they got into the center (see **Supplementary Note 6** that also discusses this). For this reason, we plotted the maximum fluorescence intensity of the gels rather than the average fluorescence intensity, to account for this edge effect phenomenon (**Fig. 4Jvii**). However, the signal was lower when the Phe residue was occluded (after PITC conjugation) or cleaved off (via in-gel Edman degradation or with trypsin) (**Fig. 4Jv-viii**). The signal in the PITC followed by TFA condition, was higher than after PITC conjugation or trypsinization of the peptide. This is consistent with the presence of Tyr, a secondary substrate of ClpS2 St-V1 NAAB^25^.

We summarized the result of this rudimentary multiplexing experiment in **Fig. 4Jviii**, where the min-max normalized intensity values for tyrosine (Y) and phenylalanine (F) read-outs in the various conditions are reported: round 0 (i.e., “DMSO”), round 1 (i.e., “PITC in DMSO 1:1000, then TFA”), or trypsin (i.e., “PITC in DMSO 1:1000, then TFA+trypsin”). Together, these results demonstrate principles of multiplexed in-gel Edman sequencing, reading out distinct fluorescence signatures for two different amino acid identities — one via an N-terminal binder and one via a covalent labeling reagent — across both chemical degradation and enzymatic digestion.

### Oxidation analysis of amino acid side chains in ExM gels

As proteins being attached to polymerized ExM gels experience a free-radical filled environment, we examined the susceptibility of amino acid side chains to oxidation as a result of free-radical polymerization. The results are described in **Supplementary Note 9**. In summary, we observed a low number of oxidation products for N-terminal Tyr, Phe, Trp, His, Pro and Arg after ExM polymerization, but more substantial oxidation products for both Cys and Met residues as a result of the gelation process (other amino acids, not tested, do not have appropriately reactive side chains). There might be different ways to prevent, reverse, or compensate for such modifications amidst our framework for *in situ* protein sequencing; however, one might also be able to guess the identity of a protein even when only some of its amino acids are recognized - and thus a low number of oxidation sites might not hamper the development of a first working *in situ* protein sequencing technology (see **Supplementary Note 10** and **Supplementary Figure 13D** for a simulation of the fraction of correctly identified proteins using a subset of NAABs). Of course, ExM gels that do not involve free radicals are also possible^48,49^.

### In-gel Edman degradation over multiple rounds (3 cycles)

We performed assessment of in-gel Edman degradation over 3 cycles, using the LC/QToF-trypsinization assay analysis of embedded peptides, described above, and recording the various peptide species (peptides with up to 3 cleaved N-terminal amino acids, and PTC-peptide from the conjugation step at each round of Edman). Importantly, performing multiple rounds of in-gel Edman degradation provides a way to confirm cleavage of PTC-peptide in the cleavage step of the in-gel Edman degradation reaction, overcoming the mass spectrometry caveat in quantitating the yield of the cleavage reaction in a single round, as noted in **Supplementary Note 3**. In short, formic acid addition, and other sample processing steps that compromise interpretation of PTC-peptide due to simulating the effects of TFA cleavage, occur only at the final step of the procedure, within the instrument itself. Thus such confounds do not compromise the interpretation of PTC-peptide from second and third round of in-gel Edman degradation, which require a fresh N-terminus from a successful cleavage reaction within the gel itself, followed by a successful conjugation, to detect these species. Gel size was relatively constant throughout (**Fig. 5A**; see **Supplementary Table 6** for all values and statistics for **Fig. 5**), and the products yielded by each step of Edman degradation largely matched what was expected from the 1- and 2-round experiments above (**Fig. 5B**). In summary, conversion rate of peptide upon reaction with PITC was high each round, with 98-99% conversion (**Fig. 5Bi**, **Fig. 5Biii**, **Fig. 5Bv**). Small amounts of some side products were observed (see **Source Data** for **Fig. 5B** for all raw chromatogram traces and mass spectra used in **Fig. 5**), as well as some PTC-peptide from earlier rounds in later rounds of Edman degradation (**Fig. 5Biv**, **Fig. 5Bvi**). These could be the result of reactions involving oxidative desulfuration of the PTC moiety, as reported in the literature^40,50^, or it is possible that some of the novel ingredients used here could result in byproducts (e.g., DMSO and TFA^51,52^), further evidenced by different effects of TFA on gel size after DMSO treatment (**Fig. 3Eii**; **Fig. 5A**) vs. without DMSO (**Fig. 2Cvii**). Perhaps using a different acid (e.g. glacial acetic acid or hydrochloric acid; as previously reported in literature; see **Supplementary Table 2**), temperature, and/or incubation time could further optimize conditions for the ExMre solid-phase conditions. Recent results have also demonstrated the possibility for base-induced Edman degradation^44^.

**Figure 5.**
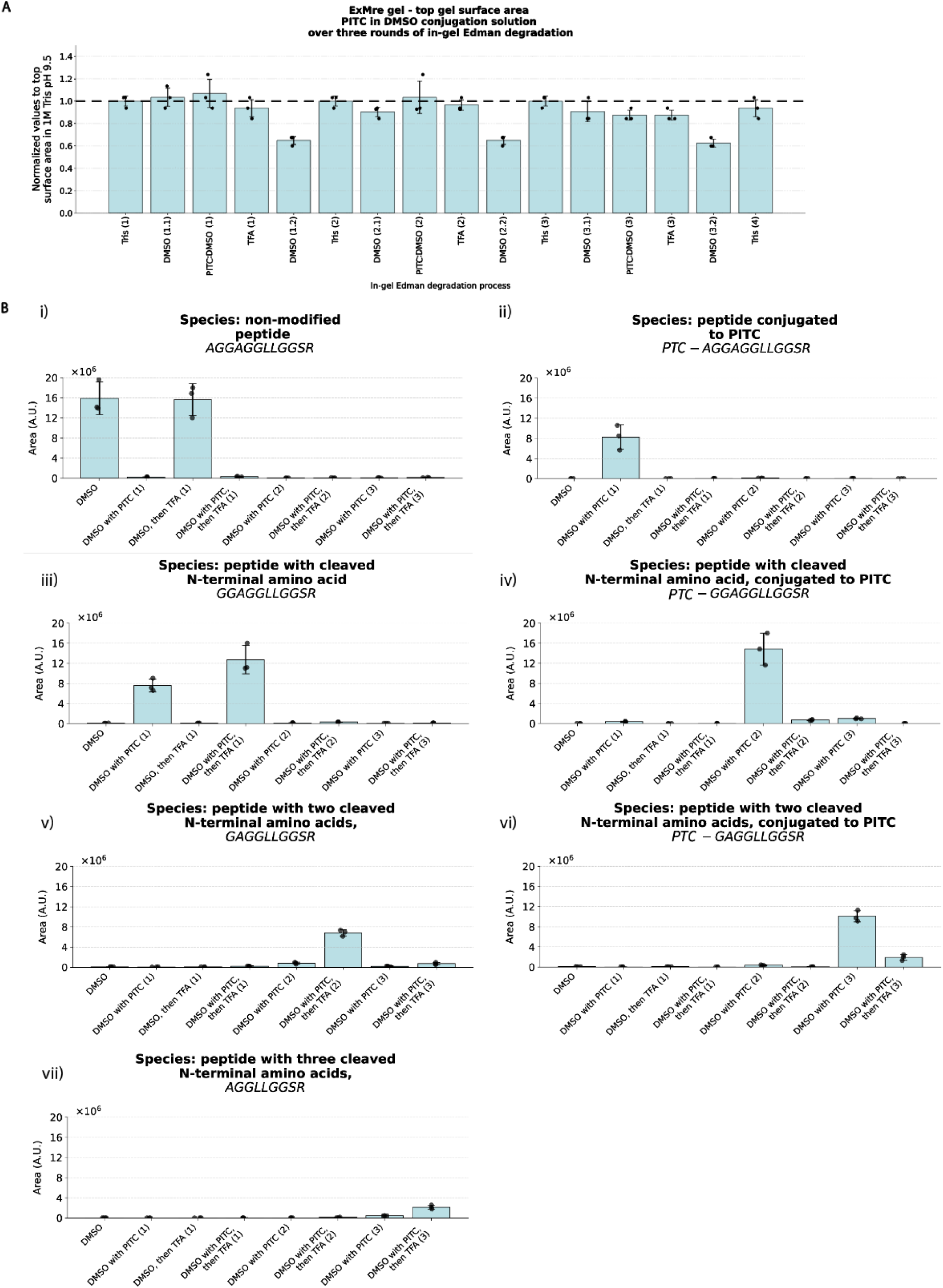
In-gel Edman degradation over multiple rounds (three cycles) using LC/QToF trypsinization assay. (A) Top/flat surface size of ExM and ExMre gels when placed in solvents used in Edman degradation. Normalized top/flat surface size to top/flat surface size in 1X PBS throughout 3 rounds of in-gel Edman degradation for ExMre gels with 1:1000 ratio PITC to DMSO. The round number is specified in parentheses (eg., “1”), and if solution is used at various steps of the same round, the first and second submersion of the gel is specified with an additional number (eg., “1.2”). (Thick dashed black line: top/flat surface size of the gel in 1M Tris pH 9.5 before Edman degradation, error bar: standard deviation, black dots, individual experiments, n=3, three separate gelation solutions.) (B) Bar graphs representing the relative abundance (arbitrary units, a.u.) of different peptide ion species from the LC/QToF, obtained from the area under the curve (AUC) of the chromatogram based on the exact mass of the various species (Agilent MassHunter Qualitative Analysis software). Conversion rates were obtained by comparing conditions: e.g. 1-(solvent with PITC[“A15-peptide”] / solvent [“A15-peptide”]), and details can be found in **Supplementary Table 6i-ii**. The relative abundance of the ion species: (i) non-modified peptide (AGGAGGLLGGSR), (ii) peptide conjugated to PITC (*PTC*-AGGAGGLLGGSR), (iii) peptide with cleaved N-terminal amino acid (GGAGGLLGGSR), (iv) peptide with cleaved N-terminal amino acid conjugated to PITC (*PTC*-GGAGGLLGGSR), (v) peptide with two cleaved N-terminal amino acids (GAGGLLGGSR), (vi) peptide with two cleaved N-terminal amino acids conjugated to PITC (*PTC*-AGGLLGGSR), (vii) peptide with three cleaved N-terminal amino acids (AGGLLGGSR), are reported throughout the Edman degradation process in the gel in the various conditions. Conditions include: DMSO only, DMSO with PITC, TFA only, and sequential DMSO with PITC followed by TFA (black dots, individual experiments; blue bar, mean; error bar, standard deviation, n=3, three separate gelation solutions).

Taken together, the efficiency of conjugation, the completeness of cleavage, and the cumulative signal reduction across iterative cycles collectively define the per-cycle sequencing yield, which we estimate at ∼70-90% from various measurements done after in-gel Edman degradation (**Fig. 3G**, **Fig. 3I**)(**Supplementary Figure 6F-G**). This compounding efficiency across cycles will be an important parameter to optimize for longer sequencing reads, ideally reaching 99% yield like *ex situ* strategies^22^, or alternatively will need to be well-characterized to enable computational correction and sequence reconstruction (as in our theoretical assessment of *in situ* protein sequencing in **Supplementary Note 10**, which assumed a predictable 90% conjugation efficiency and 70% cleavage efficiency throughout each cycle). Note that for binder-based readout, the observed ∼50% fluorescence drop between positions (**Fig. 4G-H**) was lower than predicted from cleavage yield alone, as the short dwell times of current NAABs may non-linearly affect bulk signal intensity, reflecting a convolution of sequencing yield and binding kinetics within the gel environment. However, this convolution would not apply when performing single-molecule readout, where individual binding events are detected directly. In that regime, sequencing accuracy will instead depend on binder affinity and dwell time being sufficient for detection, as well as the labeling efficiency of the NAABs across each cycle.

### An Edman reagent that permits local tethering of an isolated N-terminal amino acid

One concern is that the binding of existing NAABs to an N-terminal amino acid of a peptide is modulated by the second amino acid in the chain (or other downstream amino acids), which could present both chemical and physical hindrance^14,25,26,53^. We thus, over the past several years^53–55^, envisioned a complementary sequencing strategy, with a general workflow as depicted in **Figure 6Ai**, wherein the N-terminal amino acid is first locally tethered to the polymer backbone before being removed from the peptide chain by Edman degradation, for subsequent read-out with an amino acid binder that was designed to bind an isolated amino acid. There would be no second amino acid to interfere with the binder interaction with the first amino acid, and thus perhaps an amino acid binder could have higher affinity and specificity than otherwise possible. As such, ClickP (as we named it), a bifunctional isothiocyanate derivative with an azide group (**Figure 6Aii**), could in principle be used to isolate the N-terminal amino acid of a peptide for later isolated binder interrogation. There could be various ways of implementing this within the gel context, e.g., through initial conjugation of ClickP to the N-terminus of the peptide (i.e., the first step of Edman degradation), followed by local tethering of the N-terminal amino acid to the polymer backbone through click chemistry onto alkyne groups attached to the polymer backbone, followed by cleavage of the amide bond with TFA (see **Supplementary Figure 10B** for our thoughts on this conceptual extension). The amino acid would then remain bound to the polymer backbone, via click chemistry, in local proximity to its parent peptide. The antigen to be detected would simply be an isolated amino acid, attached to the ClickP group (see **Figure 6B**).

**Figure 6.**
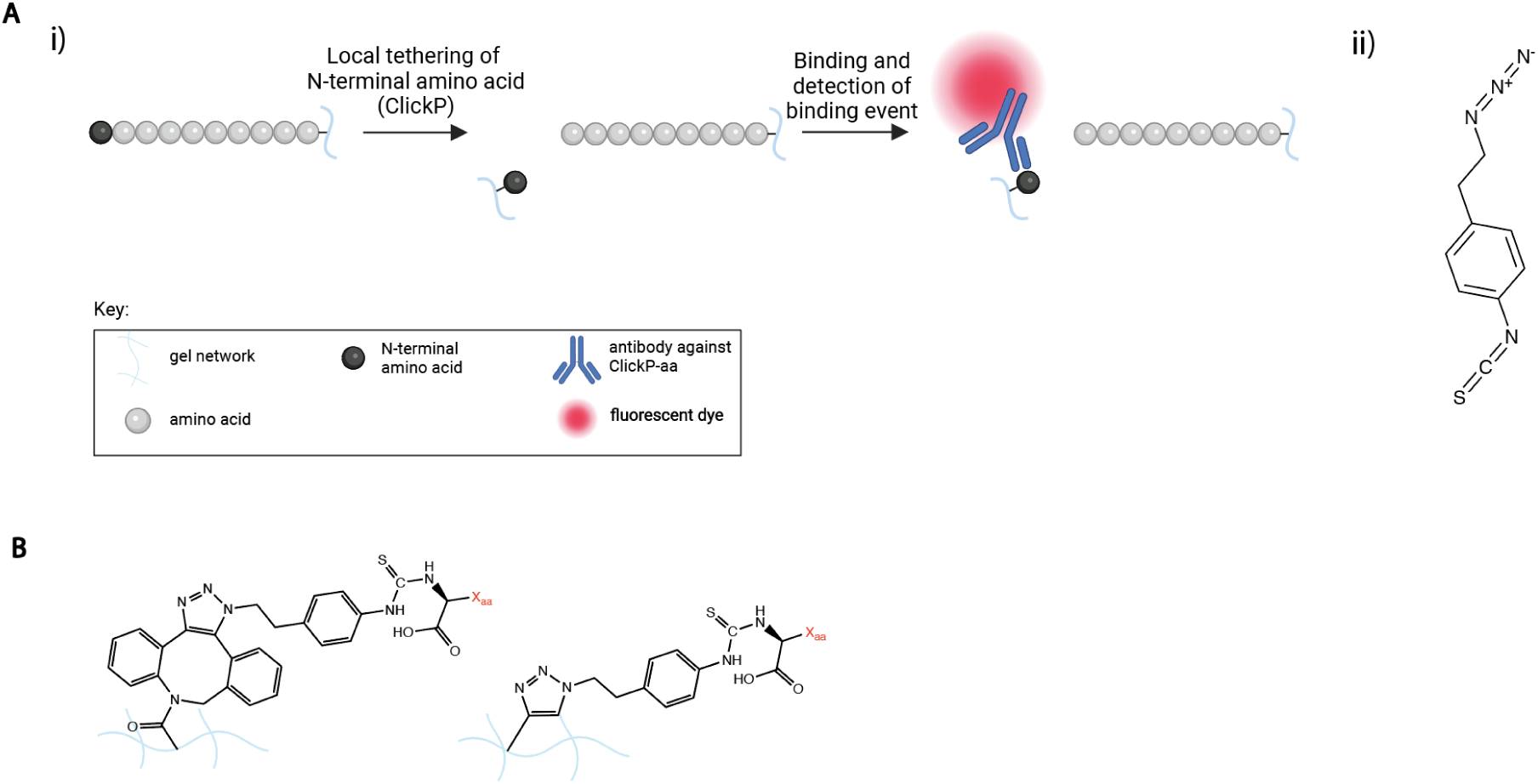
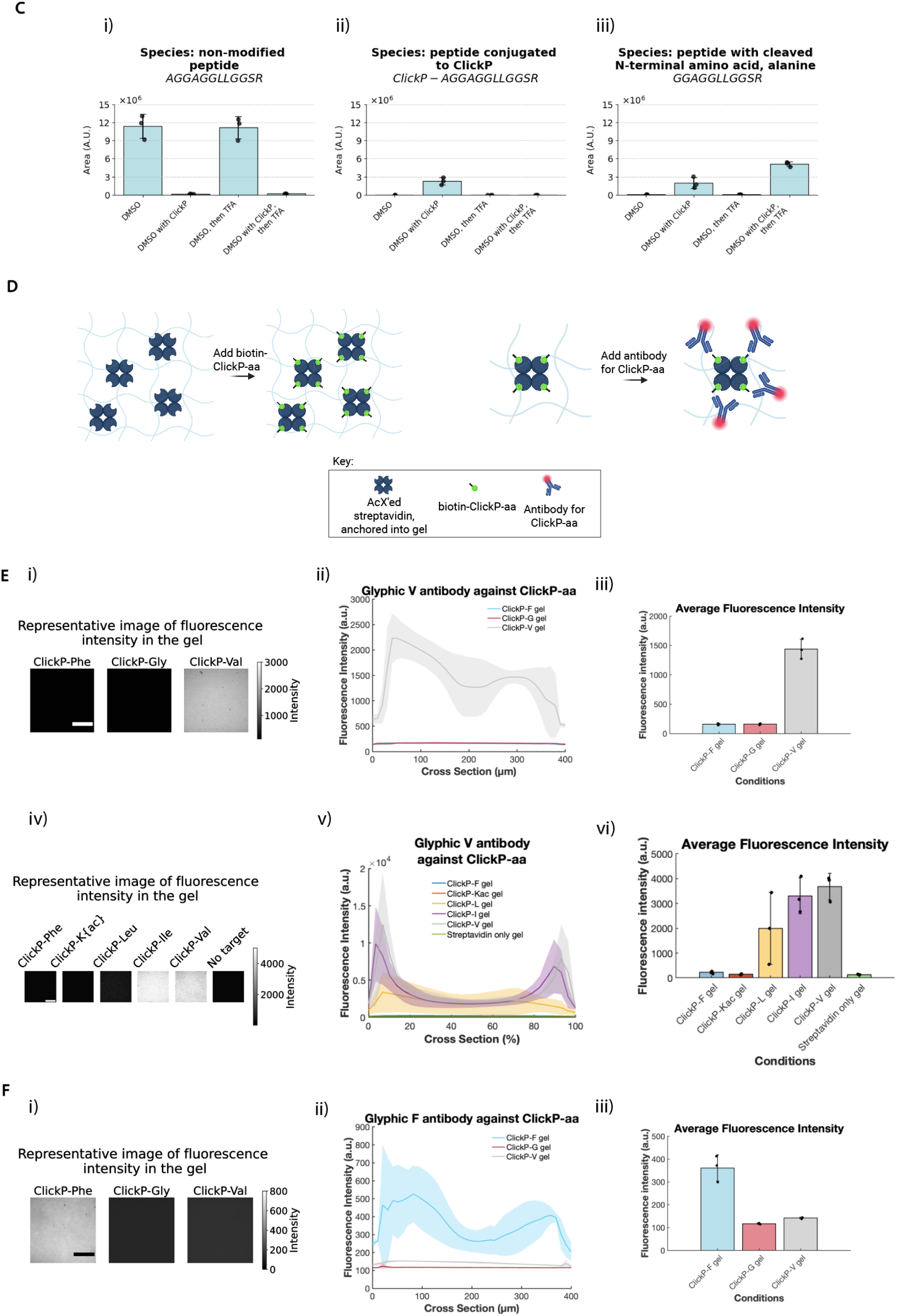
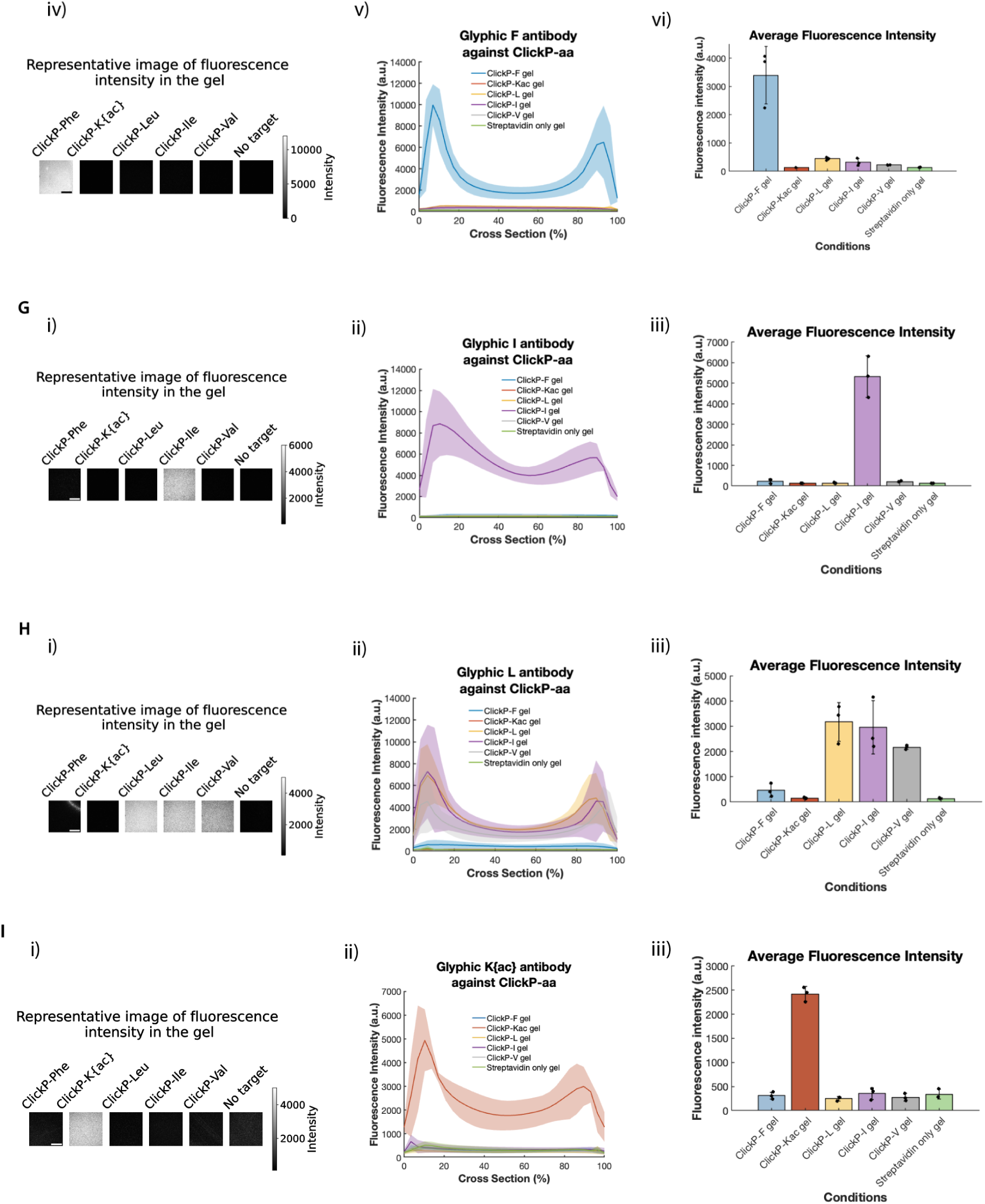
ClickP as Edman reagent, and ClickP-specific binders in ExMre gels. (A) (i) 1-(2-azidoethyl)-4-isothiocyanatobenzene, abbreviated ClickP ITC derivative, is used for local tethering of the ClickP-conjugated N-terminal amino acid to the polymer network, followed by cleavage with TFA. Binders (e.g., commercially available Glyphic antibodies) are used to detect the cleaved N-terminal amino acid bound to the polymer network. N-terminal amino acid local tethering followed by detection are performed iteratively to sequentially identify the amino acids of the chain. (ii) The structure of the ClickP ITC derivative. (B) The final antigen the Glyphic antibodies recognize. Either a reacted DBCO-azide click or a reacted alkyne-azide click group, with a given amino acid and its side chain exposed (amino acid side chain: “Xaa”, in red text), including the free C-terminus of the amino acid.. (C) Bar graphs representing the relative abundance (arbitrary units, a.u.) of different peptide ion species from the LC/QToF, obtained from the area under the curve (AUC) of the chromatogram based on the exact mass of the various species (Agilent MassHunter Qualitative Analysis software). Conversion rates were obtained by comparing conditions: e.g. 1-(solvent with ClickP[“A15-peptide”] / solvent [“A15-peptide”]), and details can be found in **Supplementary Table 7i-ii**. The relative abundance of the ion species: (i) non-modified peptide (AGGAGGLLGGSR), (ii) peptide conjugated to ClickP (4-(2-azidoethyl)phenylthiocarbamoyl-peptide, abbreviated: *ClickP*-AGGAGGLLGGSR), and (iii) peptide with cleaved N-terminal amino acid (GGAGGLLGGSR), is reported throughout the Edman degradation process in the gel in the various conditions (black dots, individual experiments; blue bar, mean; error bar, standard deviation, n=3, three separate gelation solutions). (D) Schematic of the binding assay with biotin-ClickP-aa in ExMre gels with streptavidin. ExMre gels are cast with AcX-reacted streptavidin protein. Separate gels are made for each tested amino acid, by adding biotin-ClickP-amino acid (abbreviated: biotin-ClickP-aa), to bind the streptavidin protein embedded in the gel (left). After the binding of biotin-ClickP-aa to the gels, which exposes the final antigen that Glyphic antibodies recognize, Glyphic antibodies are added to the gels to assess specificity of binding towards the various ClickP-aa (right). (E) Binding assay with biotin-ClickP-aa (aa: G, F, V) in ExMre gels with streptavidin using Glyphic V. First, streptavidin is embedded into ExM gels at a concentration of 0.1 mg/mL and after expansion and re-embedding reach a concentration of ∼5 μg/mL. Biotin-ClickP-aa is added, and after washing, GFP-conjugated Glyphic V is added. (i) A representative raw image of the fluorescence of the gels is depicted for the gels in different conditions, comparing binding of Glyphic V against biotin-ClickP-F, biotin-ClickP-V and biotin-ClickP-G ExMre gels obtained from a confocal microscope, showing a slice 90 μm deep into the gel. Scale bar in white for the ClickP-Phe image (left): 100 μm. (ii) The ExMre gels in different conditions are compared in fluorescence intensity throughout the Z-stack (line, mean; shaded area, standard deviation; n=3 different gelation solutions). (iii) Average fluorescence intensity of ExMre gels in the different conditions by averaging across the whole volume imaged (black dots, individual experiments; colored bar, mean; error bar, standard deviation, n=3 different gelation solutions). (iv) As in (i), but using Glyphic V (without GFP) followed by anti-mouse 488 secondary antibody, comparing binding to more targets, including acetyl-lysine, leucine, isoleucine (abbreviated: K{ac}, Leu, Ile, respectively) and a gel without target. The slice is an image 40 μm deep into the gel. Scale bar in white for the ClickP-Phe image (left): 100 μm. (v) As in (ii), but plotting the complete data of (iv), as gel thickness varied across replicates (300–490 μm), cross-sectional position was normalized to a percentage scale to enable direct comparison (line, mean; shaded area, standard deviation; n=2 different gelation solutions). (vi) As in (iii), but related to the data in (iv) an (v) on the data normalized to a percentage cross-section scale (black dots, individual experiments; colored bar, mean; error bar, standard deviation, n=3 different gelation solutions). (F) (i)-(iii) As in E, but using RFP-conjugated Glyphic F (black dots, individual experiments; colored bar, mean; error bar, standard deviation, n=3 different gelation solutions). (iv)-(vi) As in E, but using Glyphic F followed by anti-mouse 488 secondary antibody. (G) As in E (iv)-(vi), but using Glyphic I followed by anti-rabbit 488 secondary antibody (blank “streptavidin only” gel only has n=2 in this experiment) (H) As in E (iv)-(vi), but using Glyphic L followed by anti-mouse 488 secondary antibody. (I) As in E (iv)-(vi), but using Glyphic K{ac} followed by anti-mouse 488 secondary antibody.

Just as a proof of concept - we wanted to provide the information to benefit the community interested in developing in situ protein sequencing further, even if this was a partial story - we sought to validate whether a binder to such an amino acid could bind to its target in the gel; full realization of a ClickP-based *in situ* sequencing method would require significant further work. We designed binders to target amino acids that have been cleaved off the peptide chain after having been conjugated to ClickP. The ones here were created by Glyphic Biotechnologies (where they are commercially available) - a phenylalanine antibody (denoted F-antibody), a valine antibody (denoted V-antibody), an isoleucine antibody (denoted I-antibody), a leucine antibody (denoted L–antibody) and an acetyl-lysine antibody (denoted K{ac}-antibody) that target ClickP-tethered and decyclized amino acids (where the PTH-aa form is reverted back into PTC form, exposing the C-terminal carboxyl group, as done previously^56^). The K_d_ values for the F-antibody suggest a dissociation constant of ∼75 pM and k_off_ of ∼4.5 * 10^-5^ s^-1^, which corresponds to an average dwell time of ∼6.2 hours (see **Supplementary Figure 9B**); the V-antibody, I-antibody, L-antibody and K{ac} antibodies were similar (see **Supplementary Figure 9A** and **Supplementary Figure 9C**). These dwell times would, unlike the previous NAABs discussed, support amplification strategies in an *in situ* context, which takes minutes to hours. ClickP showed good conjugation, with a conversion rate of ∼99%, and more cleavage after TFA treatment than without, with the caveats acknowledged above regarding interpretation of PTC-amino acid cleavage with mass spec (**Fig. 6ci-iii;** see **Supplementary Table 7** for all values from **Fig. 6C**).

ClickP-antigen for various amino acid side chains (comparing Gly, Val, Phe, Ile, Leu, and acetyl-lysine abbreviated K{ac}) carrying a biotin group were incubated with separate gels containing anchored streptavidin (**Fig. 6D**; chemical structures in **Supplementary Figure 10**). Subsequent antibody (F-antibody and V-antibody, I-antibody, L-antibody, K{ac}-antibody) binding to the antigens was monitored using bulk fluorescence read-out for all antibodies, as in **Fig. 4**. The V-antibody showed binding to the valine target over glycine and phenylalanine (**Fig. 6Ei**-**iii**; see **Supplementary Table 10** for all raw values and statistics, including p-values, for **Fig. 6E-I**). Similarly, the F-antibody showed binding to the phenylalanine target over glycine and valine (**Fig. 6Fi-iii**). Further testing of these antibodies towards other targets, demonstrated cross-reactivity of V-antibody to isoleucine and leucine (**Fig. 6Eiv-vi**). However, F-antibody showed specificity even after testing against all other targets (**Fig. 6Fiv-vi**). Similarly I-antibody and K{ac}-antibody showed specificity against all other targets (**Fig. 6Gi-iii**; **Fig. 6Ii-iii**). Finally, L-antibody showed cross-reactivity to both isoleucine and valine, but not the other targets (**Fig. 6Hi-iii**). Thus, ClickP-amino acid antibodies can bind to their targets in the expansion gel.

### A theoretical assessment of *in situ* protein sequencing

Having established core chemical principles and the framework for *in situ* protein sequencing, while recognizing remaining uncertainties in binder affinity and specificity, we next set out to perform a theoretical assessment of *in situ* protein sequencing, that could guide future developments in the field (details on our strategy and results can be found in **Supplementary Note 10**). In short, the percentage of residues of a protein that remained accessible for read-out through in-gel Edman degradation and N-terminal binding depended on the extent and specificities of fixation, anchoring and digestion chemistries. However, the model suggests that even partial realization of an *in situ* protein sequencing platform could be extremely useful: under the assumptions of our model, with a subset of only 10 amino acid binders, with medium specificity (as defined by our model), 12 rounds of sequencing, with 10% conjugation error, and 30% cleavage error, could correctly identify ∼90% of the proteins in the mycoplasma proteome (see **Supplementary Figure 13D** for the detailed graph).

The human proteome, similarly, can be interrogated quite successfully even with a partially efficacious sequencing scheme with the same conjugation and cleavage errors: one would need 15 amino acid binders, with medium specificity (as defined by our model), 11 rounds of sequencing to correctly identify ∼80% of the human proteome based on assumptions given the available data (see **Supplementary Figure 20**).

## Conclusion

Here, we derive principles of *in situ* protein sequencing. To do so, we focus on a principle-yielding demonstration using synthetic testbed peptides in re-embedded ExM gels; we do not yet demonstrate sequencing of endogenous proteins in biological samples. Building from the idea that expansion microscopy (ExM) can decrowd proteins from each other for sequencing, we demonstrate several principle-oriented chemistries, including forms of Edman degradation compatible with the ExM re-embedded (ExMre) hydrogel, and detections of N-terminal amino acids with NAABs. We thereby establish that iterative, multiple round (3 cycles) sequencing chemistry on peptides can be implemented at the ensemble level within expansion hydrogels operating in a nanoscale physical regime. In addition, the N-terminal amino acid identities embedded within the polymer network can be determined using NAABs before and after N-terminal amino acid degradation. At present, due to the limiting kinetics of existing NAABs to support single molecule *in situ* readout in 3D specimens (with dwell times in the order of seconds), to focus on the principles of *in situ* protein sequencing, we here utilize ensemble-measurements of the binding events during the sequencing. For example, the ClpS2 St-V1 binder exhibits modest affinity and short dwell times that will need to be improved for single-molecule or amplification-based implementations. Nevertheless, these binders were still useful for demonstrating many core principles of in situ protein sequencing, which will support the field going forward. Our current results offer a platform upon which to build further *in situ* protein sequencing technology, including testing of NAAB designs, and validating chemical probes for covalent amino acid sidechain labeling in the gel. We also demonstrate that a bifunctional Edman reagent with a click moiety is compatible with sequencing peptides embedded within ExMre hydrogels. Commercially available binders specific to amino acids tethered via this reagent exhibit high-affinity binding kinetics (dwell times in the order of hours) and retain specificity within the hydrogel-defined physical regime.

Deriving principles of *in situ* protein sequencing required the development and validation of several chemical assays and optimizations. To enable N-terminal amino acid degradation from peptides embedded in ExMre gels, we evaluated solvents conventionally employed in Edman degradation chemistry, finding that traditional Edman conjugation solutions caused substantial gel shrinkage and opacity, whereas solvents such as DMSO did not. This preservation was essential for successful Edman degradation to proceed, as confirmed by the various quantitative assays. Further work will be required to optimize the yield beyond its current ∼70-90% level, which we attribute to interactions between solvents and reagents and the gel network, as well as potential constraints during cyclization that are unique to the nanostructured polymer network of ExM compared to other Edman degradation environments. The free-radical polymerization used in ExM gel formation can also oxidize reactive amino acid side chains, with Cys and Met being the most susceptible. However, our simulations suggest protein identification remains feasible even with partial amino acid recognition. Moreover, binders towards oxidized side-chains or, alternatively, free-radical-avoiding ExM chemistries may provide a route to avoiding this issue altogether.

Existing NAABs have a rate of dissociation that is too fast (∼seconds) for fluorescence-based imaging of single molecules via signal amplification schemes (requiring hours), and the second amino acid of the peptide chain can influence binding to the first^25^. Prior work suggests that single amino acid side chains can be determinants of an epitope^57–61^. For instance, the armadillo modular protein, when using a constant peptide for K_d_ ∼nM baseline affinity, can exhibit single side chain selectivity^62^. Nevertheless, the relatively small epitope size of a single amino acid side chain, compared to the 4-12 amino acids targeted by many antibodies^63^, and the chemical and physical proximity of subsequent amino acid residues in the peptide chain to the N-terminal amino acid, may make NAAB design challenging. One could imagine using PITC or FITC N-terminally conjugated peptides to raise the baseline affinity of the binders (or even an alternative ITC molecule that is both highly compatible with the hydrogel and provides additional groups for binder attachment), while maintaining specificity for the side chain. Binder design may be significantly enhanced by progress in computational protein design, particularly through AI-driven approaches for modeling and designing protein interactions^64–66^.

We proposed an alternative strategy to overcome the limitations of NAABs, via our heterobifunctional ClickP molecule. This molecule features an isothiocyanate moiety that binds to the N-terminal amine of peptides, along with a click group that can locally tether the amino acid conjugated to the ClickP molecule, e.g. to the hydrogel. We showed this molecule to be compatible with in-gel Edman degradation. In addition, commercially available antibodies produced against these specific antigens had strong affinity, with dwell times in the order of hours, and we demonstrated their specificity to their expected target within the ExMre gels. Local tethering of an amino acid away from its parent peptide chain would prevent influence from subsequent amino acids in the peptide chain and, thus, potentially improve specificity. Future work could include development of the entire workflow combining in-gel Edman degradation with local tethering of N-terminal amino acid with ClickP for a complete cycle of degradation, tethering, and detection of the amino acid from a parent peptide within the polymer network. Other modifications of PITC, e.g. one that allows for DNA oligo conjugation^31^, have been put forth as well.

The ExM protocol itself could be improved in terms of expansion factor, for better decrowding of proteins from each other. Indeed, ExM preserves key aspects of protein shape, enabling imaging at 1 nanometer resolution^12^; although we have shown that expansion factors larger than 100x can be achieved, future work will be needed to validate distortion, resolution and *in situ* yield of such high-expansion gels. It would also be beneficial to show that such protocols can be applied to proteins in a densely packed, heterogeneous biological sample, such as in a cell or tissue. Nevertheless, the fact that 10x ExM followed by SRRF super-resolution is capable of separating peptides cleaved from different parts of a protein (Shaib et al. 2024), already offers a potential imaging modality that could in principle support *in situ* protein sequencing.

Finally, our theoretical assessment of *in situ* protein sequencing demonstrates that protein identification remains feasible even with partial sequence coverage and incomplete reaction yields. The results, under the assumptions of our model, also reveal directions for improvement and optimization for single molecule *in situ* protein identification. Our modeling can guide experimental strategies, such as refining selection parameters and reaction efficiencies to improve peptide fragment retention for subsequent protein identification. Importantly, it can be modularly modified to test alternative conditions and constraints, and the underlying assumptions will be updated as they become better constrained by empirical data.

In summary, here we derived principles of *in situ* protein sequencing and demonstrated key chemical and optical assays toward this goal. Future work will focus on improving NAAB affinity (to ∼pM K_d_) and specificity, increasing in-gel Edman cycle yields toward those of traditional Edman degradation (∼99%), and developing expansion protocols that resolve individual peptides within densely packed biological samples (being explored by many groups, and also prototyped here). Ultimately, the approach will be validated on complex biological specimens through a top-down strategy, assessing tissue integrity, reaction homogeneity, and single-amino-acid readout from proteins *in situ*. We anticipate that our framework for *in situ* protein sequencing will open new avenues for spatially resolved proteomics directly within biological tissues.

## Methods

### Software for figure making

**Figure 1A-C**; **Figure 3A,F,Hi; Figure 4A, E; Figure 6A,B,D; Supplementary Figure 13A-B** were made with BioRender. **Figure 2A,B**; **Figure 6A,B; Supplementary Figure 8Ai-Hi; Supplementary Figure 10; Supplementary Figure 11** were made with ChemDraw.

### Synthetic peptide designs

All peptides were ordered from AAPPTEC in aliquots with > 95% purity. 5 mM stock solutions of each peptide were made by diluting lyophilized peptides with UltraPure distilled water, DNase/RNase free (Invitrogen). See **Supplementary Table 3** for exact mass and chemical formulae of all peptides used in the paper, and **Supplementary Figure 11** for structures of all peptides used in the paper.

Edman degradation readouts with liquid chromatography coupled to electrospray ionization quadrupole time-of-flight mass spectrometry (LC-ESI-QToF MS, abbreviated LC/QToF) (results from **Figure 3B-D**, **Figure 5B**, **Figure 6C**) were performed with a peptide that we denoted “A15-peptide” (which stands for AGGAGGLLGGSRGGK{acr}; K{acr} is an abbreviation for lysine modified with an acryloyl functional group). A peptide that we denoted “A9-peptide,” AGGAGK{acr}GLR, was spiked into samples to account for any ionization efficiency fluctuations.

In-gel Edman degradation monitored through bulk fluorescence in **Figure 3G** was performed with a peptide that we denoted “K{N_3_}15-peptide,” K{N_3_}GGAGGLLGGSRGGK{acr}, where K{N_3_} is an abbreviation for lysine modified with an azide functional group. For **Figure 3I**, the bulk fluorescence was performed with “AK{N_3_}15-peptide,” AK{N_3_}GAGGLLGGSRGGK{acr}.

Phenylthiohydantoin-phenylalanine (PTH-F) detection was performed with “F_1_” peptide, FGGAGRGLGK{acr} (**Figure 3H**).

ClpS2 St-V1 bulk fluorescence read-outs comparing different N-terminal amino acids (**Fig. 4A**) in the ExMre gels were performed with “F_1_” peptide, “W_1_” peptide, WGGAGRGLGK{acr}, “Y_1_” peptide, YGGAGRGLGK{acr}, and “A_1_” peptide, AGGAGRGLGK{acr}. ClpS2 St-V1 bulk fluorescence read-outs combined with Edman degradation chemistry (**Fig. 4F-G**) were performed with “F_1_” peptide and “G_1_F_2_ ” peptide, GFGAGRGLGK{acr}. tvClpS2 Q31H bulk fluorescence read-out was performed with “L_1_” peptide in addition to the previous peptides (**Fig. 4H**). Rudimentary multiplexing with ClpS2 St-V1 and sulfur-triazole exchange (SuTEx)-azide with “F_1_Y_2_L_3_” peptide, FYLAGRGLGK{acr}.

Hemagglutinin-tag (HA) peptide with a K{acr} group (or YPYDVPDYAK{acr}) was used in 9% acrylamide gels in **Supplementary Figure 7B**.

Amino acid oxidation analysis and results **A-H**) in ExM gels were performed with XaaGGAGRGLGK{acr} peptides, where Xaa was one of: M, C, W, Y, F, P, H, R. Oxidation analysis, as a result of free-radical polymerization, was performed after trypsin cleavage of the peptides from the gel (or in solution with/without ammonium persulfate (APS; Sigma A3678-25G; we include the catalog number the first time we introduce a chemical in the Methods section) and tetramethylethylenediamine (TEMED; Sigma T7024-25ML)) followed by injection in LC/QToF.

Post-translational oxidation modifications, exact mass and chemical formulae, are detailed in **Supplementary Table 3**.

### Making acrylamide, expansion microscopy (ExM), and ExM re-embedded (ExMre) gels

All washes were performed at room temperature (RT), unless otherwise specified (for all sections).

#### Acrylamide gel: making the empty gel gelation mixture (i.e. no peptide/protein)

The empty gel gelation mixture (used for gel size change measurements in **Supplementary Figure 2**) was prepared to have a final concentration of 9% acrylamide (w/v) (A9099-25G), 10 mM N,N’-methylenebisacrylamide (abbreviated: BIS; Sigma M7279-25G), 1X PBS (from diluting ThermoFisher 70011 10X PBS pH 7.4), in UltraPure water (Invitrogen 10977015; below, whenever water is mentioned, we use this UltraPure water unless otherwise indicated) to a total volume of 100 μL (see **Table 1: Acrylamide gelation solution)**. Subsequently, pre-chilled 10% (v/v) TEMED and 10% (w/v) APS stock solutions were added to the mix (5 μL each) on ice. Casting the gel was performed as described in section **Casting the gel**.

**Table 1:** Acrylamide gelation solution.

| Acrylamide gelation solution | Concentration | Final concentration | Add |
| --- | --- | --- | --- |
| Acrylamide | 50% (w/v) | 9% (w/v) | 20 μL |
| BIS | 130 mM | 9.1 mM | 7.7 μL |
| PBS | 10x | 0.9x | 10 μL |
| Water | | | 62.3 μL |
| APS | 10% (w/v) | 0.45% (w/v) | 5 μL |
| TEMED | 10% (v/v) | 0.45% (v/v) | 5 μL |
| Total volume | | | 110 μL |

#### Acrylamide gel: making the gelation mixture with peptide

The peptide-containing gelation mixture (used for **Supplementary Figure 7A-B**) was prepared to have a final concentration of 9% acrylamide (w/v), 10 mM BIS, 1X PBS, 20 μL of peptide (using a stock of 25 mM for **Supplementary Figure 7A**, and using a stock of 5 mM for **Supplementary Figure 7B**), and water to a total volume of 100 μL (final peptide concentration of 5 mM for **Supplementary Figure 7A**, and final concentration of 1 mM for **Supplementary Figure 7B**).

Subsequently, pre-chilled 10% (v/v) TEMED and 10% (w/v) APS stock solutions were added to the mix (5 μL each) on ice. Casting the gel was performed as described in section **Casting the gel**.

#### ExM gel: making the empty gel gelation mixture

The empty gel gelation mixture (used for gel size change measurements in **Fig. 2c** and in **Supplementary Figure 2**) was prepared with 60 μL StockX monomer solution (see **Table 2: StockX monomer solution**), 10 μL 130 mM BIS, and water to a total volume of 90 μL. Subsequently, pre-chilled 10% TEMED (v/v) and 10% APS (w/v) stock solutions were added to the mix (5 μL each) on ice (see **Table 3: ExM empty gel and peptide-containing gelation solutions**). StockX monomer solution was made as follows as in the previously described protocol^7^. Aliquots of StockX monomer solution were kept at -20 °C. Casting the gel was performed as described in section **Casting the gel**.

**Table 2:** StockX monomer solution.

| StockX monomer solution | Concentration | Amount (mL) |
| --- | --- | --- |
| Sodium acrylate (Combi-Blocks QC-1489) | 38% w/v | 2.25 |
| Acrylamide | 50% w/v | 0.5 |
| BIS | 2% w/v | 0.75 |
| Sodium chloride (J60434.AK) | 5 M | 4 |
| 10X PBS | 10X | 1 |
| Water |  | 0.9 |
| **Total** |  | **9.4** |

**Table 3:** ExM empty gel and peptide-containing gelation solutions.

| Reagent | Concentration | Final concentration | Add |
| --- | --- | --- | --- |
| StockX monomer solution | | | 60 μL |
| Additional BIS (not including concentration of BIS in the StockX solution) | 130 mM (2% (w/v)) | 13 mM | 10 μL |
| Peptide (replace with water for the empty gel gelation solution) | 5 mM | 1 mM | 20 μL |
| APS | 10% (w/v) | 0.5% (w/v) | 5 μL |
| TEMED | 10% (v/v) | 0.5% (v/v) | 5 μL |
| Total volume | | | 100 μL |

#### ExM gel: making the gelation mixture with peptide

The peptide-containing gelation mix (used for **Fig. 3B-E,G,Hii; 4B-D,F-G; 5A-B,F-G; 6C,E-F**) was prepared with 20 μL 5 mM peptide of the given sequence, 60 μL StockX, 10 μL 130 mM BIS, with water to a total volume of 90 μL. Subsequently, pre-chilled 10% TEMED v/v and 10% APS w/v stock solutions were added to the mix (5 μL each) on ice (see **Table 3: ExM empty gel and peptide-containing gelation solutions**). Casting the gel was performed as described in section **Casting the gel**.

#### Casting the gel

After gentle mixing, the mixture (taking ∼70 μL of solution) was pipetted into a gelation chamber, created using a glass slide (VWR, #48300-026), a 22 x 22 mm #1.5 coverslip (VWR, #48366-227), and one or two layers of parafilm (Uline, S-25929) as spacers, as described previously ^67^. **Fig. 2C; 3B-E,G; 5A-B**, **6c; Supplementary Figure 1A-H; Supplementary Figure 2A-M; Supplementary Figure 3A-B; Supplementary Figure 4A-C; Supplementary Figure 7A-B; Supplementary Figure 8A-H**, were made with 2 parafilm-thick (∼260 μm) chambers (“thick” gel), whereas **Figure 3Hii**; **4B-D,F-G; 6E-F; Supplementary Figure 6A-B** were made with 1 parafilm-thick (∼130 μm) chambers (“thin” gel). By spacing the parafilm stacks 10 mm apart on the glass slide, the final gelation chamber was 10 mm long, 22 mm wide, and with height determined by the parafilm thickness. The gelation chamber was placed within a Tupperware box (with two holes drilled to insert a nozzle for nitrogen purging, and tape to close the holes shut after purging) with a damp paper towel, purged with nitrogen for 5 minutes to remove oxygen, and then incubated at 37 °C for 1.5 hr for gelation. Upon completion of the gelation process, the chamber was disassembled, and the gel was cut into 10 mm x 5 mm rectangles before being transferred to 1X PBS. The gels were washed 3 times in 1X PBS, 5 minutes each.

#### Expanding the ExM gels

ExM gels, cut to 10 mm x 5 mm rectangles after gelation, were placed on a glass slide inside a 4 well plate in 1X PBS. 1X PBS was removed and the gels were expanded at RT by washing 3 x 20 min with 5 mL water until they reached ∼3.5x expansion. Expanded gels were then cut down to either 5 mm x 5 mm (solvent testing on ExM gels in **Fig. 2C**, **Supplementary Figure 2A-M**) or used for further re-embedding in **Fig. 2C; 3B-E,G,Hii; 4B-D,F-G; 5A-B, 6C,E-F**.

#### Re-embedded ExM (ExMre) gels

Expanded ExM gels (10 mm x 5 mm before expansion, and reaching ∼35 mm x ∼17 mm in width and length after expansion) were re-embedded following a similar strategy as a previously published protocol ^19^. Any remaining water was removed from the 4 well plate, and the gels were submerged in a 5 mL solution with 3% acrylamide (w/v), 0.15% BIS (w/v) obtained from a 1:13 dilution of acrylamide/BIS 19:1, 40% (w/v) solution (AM9024; Fisher Scientific), 5 mM Tris pH 8, 0.075% APS (w/v), 0.075% TEMED (v/v) for 30 min at RT on a shaker (50 rpm). The re-embedding solution was removed from the 4-well plate. A sandwich of 22 x 22 mm #1.5 coverslips (VWR, #48366-227) glued together were placed on either side of the gel. The sandwich consisted of four coverslips for gels with a height of ∼700 μm (i.e., the same height as expanded ExM gels made with 2 parafilm thick chambers): in **Fig. 2C; 3B-E,G; 5A-B**; **6C; Supplementary Figure 1A-F,H; Supplementary Figure 2A-M; Supplementary Figure 3A-B; Supplementary Figure 4A-C**. The sandwich consisted of two coverslips with a height of ∼350 μm (i.e., the same height as expanded ExM gels made with 1 parafilm thick chambers): in **Fig. 3Hii**; **4B-D,F-G; 6E-F; Supplementary Figure 6A-B**. Then, for all gels, a 50 mm x 24 mm #2 coverslip (VWR, #48382-136) was gently placed on the top of the gel. Gel chambers were placed inside a closed Tupperware box with a damp paper towel (as in section **Casting the gel)**, which was purged with nitrogen for 10 min. Gel chambers were then incubated for 1.5 hr at 37 °C. Expanded re-embedded (ExMre) gels were then cut down to 10 mm x 5 mm (for **Supplementary Figure 3B**) or 5 mm x 5 mm (for **Fig. 2C, 3B-E,G,Hii, 5A-B, 6c; Supplementary Figure 1A-F,H; Supplementary Figure 2A-M; Supplementary Figure 3A; Supplementary Figure 4A-C; Supplementary Figure 6A-B**) or 2 mm x 2 mm size gels (for **Figure 4B-D,F-G, 6E-F; Supplementary Figure 7A-B**) for downstream steps.

### Gel flat/top side surface size tests

#### General solvent tests on gel flat/top side surface size

Three separate batches of ExM gels (in expanded state) and ExMre gels, without embedded peptide, were made, cut into ∼5 mm x 5 mm squares (∼700 μm thick) for solvent testing in **Figure 2C**. Similarly, 9% acrylamide, ExM (in expanded state) and ExMre gels, without embedded peptide, were cut into ∼5 mm x 5 mm squares (∼700 μm thick) for solvent testing in **Supplementary Figure 1** (gel images) and **Supplementary Figure 2** (gel size changes). All gels were washed 3 times 10 min washes in 1X PBS. The exact dimensions of each gel were recorded after the 1X PBS washes by measuring the gel flat/top side surface size on a glass slide using a ruler. In cases where the gel tore, the edges were carefully realigned prior to measurement. When the gel folded into a solid form after exposure to the solvents described below and could not be unfolded, measurements were taken from the accessible edges by rotating the vial to obtain the most representative size.

Glass vials (Chemglass, CG-4912-01) were filled with 300 μL of the following organic solvents: trifluoroacetic acid (TFA) (Sigma, T6508-100ML), 1X PBS, water, phenylisothiocyanate (PITC; Sigma, 317861), acetonitrile (ACN) (Sigma Aldrich, 271004), 1:1 ratio pyridine to water (Millipore Sigma, 270407) (all ratios throughout are of volumes added, unless otherwise indicated), dimethylsulfoxide (DMSO) (Millipore Sigma, 276855), formamide (Fisher Scientific, AM9342), 1M Tris pH 8 (AM9856), 1M Tris pH 9.5 (J62084.K2), 0.1 M sodium bicarbonate pH 8.5 (J60408) (**Fig. 2cii-vii**, and **Supplementary Figure 1Bi-ii, Ci-ii, Di-ii, Ei-ii, F**, **H** and **Supplementary Figure 2A-D**). For other experiments, the glass vials were filled with 270 μL of the following organic solvents: ACN, DMSO, formamide, pyridine, 1:1 pyridine:water, triethylamine (Millipore Sigma, 471283), dimethylformamide (DMF) (Millipore Sigma, 270547), 1:1 ratio DMSO to water (**Supplementary Figure 1A, Biii, Ciii, Diii, Eiii**, and **Supplementary Figure 2E-L**). The glass vials were filled with 231 μL 0.1 M sodium bicarbonate pH 8.5 for **Supplementary Figure 2M**. Gels were submerged in the solvent or buffer on a heat block at 50 °C for 30 min. For **Fig. 2Cii-vii**, and **Supplementary Figure 1Bii, Cii, Dii, Eii**, the solutions were then removed, and the flat/top side surface size of each gel was measured (from the bottom, through the glass vial). Then, PITC or ClickP (Enamine, EN300-37440925) at 1:1000 ratio PITC:solvent or ClickP:solvent, respectively, or fluorescein isothiocyanate (FITC) ‘Isomer 1’ (ThermoFisher, F1906) at a final concentration of 5.9 mM FITC (23:77 of DMSO:0.1 M sodium bicarbonate pH 8.5), was added to fresh solvent or buffer. This solution was added into each glass vial to submerge the gels at 50 °C for 30 min, followed by gel flat/top side surface size measurement again (from the bottom, through the glass vial). For **Supplementary Figure 2E-L**, PITC was added at 1:9 ratio PITC:solvent in the same original solvent (30 μL added to the same 270 μL solvent), and for **Supplementary Figure 2M** FITC in the same original buffer at a final concentration of 5.9 mM FITC (23:77 ratio DMSO:0.1 M sodium bicarbonate pH 8.5; 69 μL of 10 mg/mL FITC in DMSO added to 231 μL buffer) and incubated for 30 min at 50 °C. The gel size was measured again after the solution was removed (from the bottom, through the glass vial).

#### Surface flat/top side size during Edman degradation chemistry

ExMre gels, with embedded peptide, were cut into ∼5 mm x 5 mm squares (∼700 μm thick) after 3x 10 min 1 mL washes in 1M Tris pH 9.5 at RT in 1.5 mL Eppendorf tubes. The exact dimensions of each gel flat/top side surface size throughout the in-gel Edman degradation were recorded for **Fig. 3ei-iii** and **Fig. 5A**. After the 1M Tris pH 9.5 washes, gels were placed on a glass slide and the gel surface size was measured on the glass slide using a ruler. In all the next steps, the gels were submerged in 300 μL of solution and a heat block at 50 °C in glass vials. Gels were transferred to glass vials, washed 2 times with solvent (eg. DMSO or formamide) 5 min each, and gel flat/top side surface size was measured again after removal of the solution from the second wash (measured from the bottom of the glass vial). Then, PITC to solvent (1:1000 ratio, e.g., DMSO or formamide) was added to the gel for 1 hour before removal, followed by the next measurement. The gel was then washed 3 times with solvent (eg. DMSO or formamide) 5 min each before TFA was added for 30 min at 50 °C. After discarding the TFA, the gels were measured again (from the bottom of the glass vial). The gels were washed 2 times with solvent (e.g., DMSO or formamide) 5 min each, and the gel flat/top side surface size was measured again after removal of the last wash. Then, the gels were washed 3 times with Tris pH 8 (**Fig. 3Ei-iii**), 10 min each at RT, measured, and transferred to 1.5 mL plastic Eppendorf tubes. For **Fig. 5A**, the gels were washed 3 times with 1M Tris buffer pH 9.5, 10 min each at RT, measured, before the next cycle of in-gel Edman degradation. During the two subsequent rounds of in-gel Edman degradation for **Fig. 5A** the flat/top side surface size was similarly measured.

#### In-gel Edman degradation

In-gel Edman degradation was performed as outlined in the schematic in **Fig. 3A**. First, a heating block in a chemical hood was preheated to 50 °C. Gels were washed in 1.5 mL Eppendorf tubes, 3 times 5 min with 1 mL of 1M Tris pH 9.5 at RT. The gel samples were carefully transferred into glass vials with solvent on the heat block. In all the next steps, the gels were submerged in 300 μL of solution in glass vials that were placed on a 50 °C heat block. Samples were washed with solvent or buffer (e.g., DMSO for **Fig. 3C**, formamide for **Fig. 3B**, 0.1 M sodium bicarbonate pH 8.5 for **Fig. 3D**) two times, 5 minutes each. Samples were then incubated with PITC to solvent (1:1000 ratio) for a duration of 1 hour (**Fig. 4F-G**, **Supplementary Figure 3B** performed with 1:9 of PITC:solvent; **Supplementary Figure 4a** with 1:100 ratio of PITC:solvent; **Supplementary Figure 4c** with 1:10,000 ratio of PITC:solvent; **Fig. 6c** performed with 1:1000 ratio ClickP:DMSO). For FITC conjugation (**Fig. 3D**), 5.9 mM FITC in 23:77 of DMSO:0.1 M sodium bicarbonate pH 8.5 was added to the samples for a duration of 2 hours, in the dark. Following this, samples were washed 3 times 5 minutes in solvent or buffer (except **Supplementary Figure 3B** which was performed with 1 solvent wash; **Fig. 4F-G** was performed with 2 solvent washes). After removal of the solvent, gels were submerged in TFA for 30 min. TFA was removed and samples were washed with solvent twice, 5 minutes each (this step was omitted for PTH detection in **Fig. 3H**, see **Edman degradation and PTH detection** methods section). Glass vials were discarded, and gels were placed back into Eppendorf tubes at RT. If only 1 in-gel Edman degradation round was performed, as in **Fig. 3B-D, G**; **Fig. 4F-G**; **Fig. 6C**, the tubes were washed 3 times 10 min each, with 1 mL of 1M Tris buffer pH 8 at RT before trypsinization. If another round of in-gel Edman degradation was performed, as in **Fig. 5A-B**, the tubes were filled with 1ml of 1M Tris buffer (pH 9.5), and washed 3 times 10 min each at RT, and the pH was checked by immersing pH strips in the buffer solution after the last wash. (Other gels that served as controls with 1 or 2 rounds of in-gel Edman degradation during the multiround experiment were washed 3 times 10 min with 1 mL of 1M Tris buffer pH 8 and stored at 4 °C.) After the last round was performed, the tubes were washed 3 times 10 min with 1 mL of 1M Tris buffer pH 8 at RT before trypsinization.

### LC/QToF analysis of synthetic peptides in the gel

#### Gel preparation

Edman degradation read-outs with LC/QToF were performed with A15-peptide in ExMre gels. Each gel size was 5 mm x 5 mm x ∼700 μm after re-embedding (see section: **Methods: Re-embedded ExM (ExMre) gels**) .

#### Trypsinization of peptides from the gels

Peptides were trypsinized from the gels after in-gel Edman degradation (see **Fig. 3A**; right hand side). After the 3 times 10 min 1 mL 1M Tris buffer (pH 8) washes, they were submerged in 50 μL of 20 μg/mL trypsin (New England Biolabs, #P8101S) in 50 mM Tris-HCl pH 8 in the bottom of the 1.5 mL Eppendorf tube (ensuring they were completely submerged in trypsin solution). The gels were then incubated overnight (∼16 hours) at 37 °C. The supernatant (∼50 μL) was placed in a mass-spectrometry vial (ThermoFisher, 6PSV9-1P), with the glass insert (ThermoFisher, 6PME03C1SP) and screwed shut (ThermoFisher, 6PSC9ST101X).

The A9-peptide was spiked into all the samples, during trypsinization, at a 5 μM final concentration (the A9-peptide was added along with the 20 μg/mL trypsin for the overnight, ∼16 hours, incubation at 37 °C) to monitor any fluctuations in ionization efficiency of the samples (the abundance of A9-peptide from experiments in **Fig. 3B-D**; **Fig. 5B**; **Fig. 6C** were plotted in **Supplementary Figure 12**).

#### Mass spectrometry (LC/QToF)

Samples were brought to the Department of Chemistry Instrumentation Facility (DCIF) at MIT for injection into the LC/QToF. This instrument was a high-resolution Agilent 6545 mass spectrometer coupled to an Agilent Infinity 1260 LC system running a Jet Stream ESI source (mass accuracy of 1-3 ppm using real-time calibration, with a mass resolving power of 45,000 (FWHM) at m/z of 2722; measurable m/z range from 50 to 10,000). The column used was a reversed-phase ZORBAX Eclipse AAA, 3.0 x 150 mm, 3.5 µm, C18 (Agilent part number: 961400-302). For **Supplementary Figure 3b**, a reverse-phase ZORBAX StableBond 300 C3, 2.1 x 150 mm, 5 µm column was used (Agilent part number: 883750-909). Before each run, the instrument was positively tuned, and 10 μL blank water washes were used to wash the column twice. 4 μL of the trypsinized supernatant was injected into the mass spectrometer for ExMre gels with 5 x 5 mm top/flat surface size in **Fig. 3B-D**; **Fig. 5B**; **Fig. 6C**; **Supplementary Figure 3A**; **Supplementary Figure 4A-C;** and for the control curve in **Supplementary Figure 5** (with an estimated concentration of ∼50 μM peptide in ExMre gels, ∼50 pmol (∼50 ng) of trypsinized peptides were injected). 2 μL of the trypsinized supernatant was injected into the mass spectrometer for ExMre gels with 10 x 5 mm top/flat surface size in **Supplementary Figure 3B**. 1 μL of the trypsinized supernatant was injected into the mass spectrometer for non-expanded ExM gels with 10 x 5 mm top/flat surface size for oxidation analysis in **Supplementary Figure 8**. 10 μL of the supernatant was injected into the mass spectrometer for PTH-F detection in **Fig. 3Hii**. The samples were injected using **LC Method**, which details the mobile phase during the ∼23 min estimated run time.

#### LC Method: chromatographic separation method (LC/QToF)

**LC Method: chromatographic separation method (LC/QToF):**
| Time (min) | Solvent A (%) | Solvent B (%) | Flow (mL/min) | Max Pressure Limit (bar) |
| --- | --- | --- | --- | --- |
| 0.00 | 99.5 | 0.5 | 0.300 | 400.00 |
| 1.50 | 99.5 | 0.5 | 0.300 | 400.00 |
| 15.00 | 40.0 | 60.0 | 0.300 | 400.00 |
| 18.00 | 5.0 | 95.0 | 0.300 | 400.00 |
| 19.00 | 5.0 | 95.0 | 0.300 | 400.00 |
| 20.00 | 99.0 | 1.0 | 0.300 | 400.00 |

From 0.00 to 1.50 min, the eluate was sent to waste and not to the Q-ToF. The Q-ToF was turned on at 1.50 min. Solvent A: 0.1% formic acid in water. Solvent B: 0.1% formic acid in ACN.

#### Analysis of LC/QToF data

Bar graphs representing the relative abundance (arbitrary units, a.u.) of different peptide ion species from the LC/QToF were obtained from the area under the curve (AUC) of the chromatogram based on the exact mass of the various species (Agilent MassHunter Qualitative Analysis software). The relative abundance of the ion species was reported throughout the Edman degradation process in the gel in the various conditions.

Specifically, to obtain the AUC in an automated fashion (**Fig. 3B-D**, **Fig. 5B**, **Fig. 6C, Supplementary Figure 3A-B, Supplementary Figure 4A-C, Supplementary Figure 5** and **Supplementary Figure 12**), Agilent MassHunter Qualitative Analysis software was opened with the files to analyze. Using the “Compound Analysis” tab, with “Find by Formula”, the given formula and exact mass (see **Supplementary Table 3**) was used to extract the information about each species of interest from the total ion chromatograms (TICs). They were saved as an excel spreadsheet for downstream analysis and plotting. Plots were then made in python (see **Code**).

The results from this search were manually inspected to ensure correct peak identification. The raw TIC, and the corresponding mass spectra of the species of interest were exported as metafiles (‘.emf’) (available in **Source Data**, with an arrow pointing towards retention time and peak of interest).

For **Figure 3Hii** and **Supplementary Figure 8A-H**, the AUC for the species of interest were obtained by manual integration by right clicking the chromatogram and selecting “Extract chromatogram”. In the pop-up window, the mass of the [M+H]^+^ species was inputted, and the detected peak(s) from this extraction were integrated into a single value (the integration considers an error of ± 20 ppm (Δm) : Δm = (ppm error × exact mass) / 1,000,000 for detection).

#### In-gel Edman degradation with PTH detection

F_1_ peptide (3 separate vials received from AAPPTEC), were cast in ExM gels with a peptide concentration of 1mM in the gelation solution, and ∼130 μm (i.e., 1 parafilm) thick gelation chamber (see section **Methods: ExM gel: making the gelation mixture with peptide** and **Re-embedded ExM (ExMre) gels**). After they were re-embedded into ExMre gels, they were washed 3x with 1X PBS 10 min, and cut into 5 x 5 mm squares on a glass slide. In-gel Edman degradation was performed with PITC to DMSO (1:1000 ratio PITC:DMSO), as described in section **In-gel Edman degradation**, with some modifications detailed here. Positive control included both PITC and TFA treatments, whereas negative control omitted the PITC. After the 30 min TFA incubation at 50 °C, the 300 μL of TFA was removed, and the gel was directly resuspended in 50 μL of 1:1 ACN to water. After shaking the glass vial 10 times, the 50 μL supernatant was pipetted up and down 20 times. The gel shrank and became slightly opaque in this process. The supernatant, containing the PTH-aa, was placed in a mass spec vial for downstream analysis, where 10 μL of the sample was injected into the LC/QToF (an estimated 2 pmol was injected, if we assume a ∼70% Edman efficiency). Analysis of PTH-F abundance was performed using PTH-F exact mass: 282.0827 ± 0.0056 Da (using an error of ± 20 ppm (Δm) : Δm = (ppm error × exact mass) / 1,000,000). Positive control of PTH-phenylalanine, in **Supplementary Figure 6B**, was purchased from TCI, P0367. A stock concentration of 12.5 mM in 1:1 ACN to water (volume ratio, used throughout unless otherwise indicated) with 0.1% TFA was made and stored at -20 °C in 100 μL aliquots. The stock solution was diluted to 100 μM in 1:1 ACN to water and 10 μL of this sample was injected into the LC/QToF (∼1 nmol).

#### Control curve for calculating yield on LC/QToF

Peptide sequences were the fragments from A15 peptide expected after its trypsinization from the gel (fragments recovered in **Fig. 3B-D**): AGGAGGLLGGSR (non-modified peptide), and GGAGGLLGGSR (peptide with cleaved N-terminal amino acid). These sequences were ordered in 3 different 0.8 mg peptide aliquots of each peptide from AAPPTEC (> 95% purity). Peptides were prepared together in solution at various concentrations from 5 μM to 60 μM for the 3 separate replicates for each concentration (“C_product” in **Supplementary Figure 5**: 5, 10, 20, 30, 40, 50, 60 μM, and “I_product” was the abundance of the peptide). In addition, in each solution, a control peptide, namely the A9-peptide, was spiked at 5 μM final concentration in all triplicate samples for each concentration and was used as the standard (“C_standard” was 5 μM, and “I_standard” was the abundance of the A9-peptide). Peptide abundances were analyzed on Agilent MassHunter Qualitative Analysis software. The method for **Analysis of LC/QToF data** was performed for those species to obtain the AUC in an automated fashion.

#### Strain-promoted alkyne-azide cycloaddition (SPAAC) read-out of in-gel Edman degradation

Edman degradation followed by SPAAC for bulk fluorescence read-outs (**Fig. 3F-G, I**) were performed with K{N_3_}15-peptide and the AK{N_3_}15-peptide. The ExM gelation solution containing peptides at 1 mM concentration was cast in a gelation chamber with thickness ∼260 μm (i.e., 2 parafilm thickness), expanded and re-embedded to reach ∼2.7X expansion factor in 1X PBS. Thus, the final peptide concentration in the gels reached ∼50 μM. ExMre gels were cut to 5 x 5 mm and reached a thickness of ∼700 μm with expansion, with each gel containing ∼900 pmol peptide.

The steps were the same as the protocol written in the section **In-gel Edman degradation**, but with K{N_3_}15-peptide and the AK{N_3_}15-peptide, and with modifications detailed here. As in the in-gel Edman degradation, PITC to DMSO (1:1000 ratio PITC:DMSO) was used for the conjugation step and after removing TFA, the gels were washed 3 times, 10 min each, with 1 mL of 1M Tris buffer pH 8 in 1.5 mL Eppendorf tubes (**Fig. 3G**), or 1 mL of 1M Tris buffer pH 9.5 (**Fig. 3i**). For **Fig. 3I**, the gels were 2 x 2 mm top/flat surface size (as compared to 5 x 5 mm for **Fig. 3G)** and the in-gel Edman degradation was then performed for a total of 3 times. The gels with condition “PITC:DMSO+TFA+trypsin” (**Fig. 3G**) or “(PITC:DMSO+TFA)*3+trypsin” (**Fig. 3I**) were then submerged in 50 μL of 20 μg/mL trypsin in 50 mM Tris-HCl pH 8 in the bottom of the 1.5 mL Eppendorf tube (ensuring they were completely submerged in trypsin solution). The gels for other conditions were submerged in 50 μL of 50 mM Tris-HCl pH 8 (without trypsin). The gels were then incubated overnight (∼16 hours) at 37 °C. Then, they were washed in 1X PBS 3 times, 10 minutes each, at RT. 20 μg/mL of DBCO-AF488 (Lumiprobe, #218F0) was prepared, from a stock of 10 mg/mL fluorophore in DMF stored at -20 °C, in 1X PBS (500 μL per tube). Gels were submerged in 500 μL of the solution and checked to be well suspended within the Eppendorf tube (not stuck to the edge throughout the staining process). The click reaction was left to react overnight (∼16 hours) at 4 °C, and washed 4 times the next day with 1X PBS, 10 min per wash prior to imaging.

#### Yield calculations

For 1-round in-gel Edman degradation yield, *i.e.* **Fig. 3G**, **Fig. 4Iv-viii, Supplementary Figure 6Fi-iii**, the yield calculation based on the SPAAC assay for the K{N_3_}15-peptide (**Fig. 3G** and **Supplementary Figure 6Fi-iii**) and the SuTEx-azide assay for the Y_1_ peptide (**Fig. 4Iv-viii**) was defined as:

% yield = (“1”-“4”)/(“1”-“5”)

where the numbers represent the separate conditions as indicated in **Fig. 3G**: where (1) DMSO only, (2) PITC to DMSO (1:1000 ratio PITC:DMSO), (3) TFA only, (4) PITC to DMSO (1:1000 ratio PITC:DMSO) followed by TFA, and (5) same as (4) but with the additional trypsinization step.

For 2-round in-gel Edman degradation yield, *i.e.* **Fig. 3Ii-iii,** and **Supplementary Fig. 6Gi-iii**, the yield calculation based on the SPAAC assay for the AK{N_3_}15-peptide was defined as:

% yield = (“1”-“4”)/(“1”-“5”)

where the numbers represent the separate conditions as indicated in **Fig. 3I**: where (1) DMSO only, (2) TFA only, (3) PITC to DMSO (1:1000 ratio PITC:DMSO) followed by TFA, (4) 2 rounds of PITC to DMSO (1:1000 ratio PITC:DMSO) followed by TFA, and (5) 3 rounds of PITC to DMSO (1:1000 ratio PITC:DMSO) followed by TFA, and (6) but with the additional trypsinization step.

### Copper(I)-catalyzed Azide-Alkyne Cycloaddition (CuAAC) read-out of in-gel Edman degradation

For **Supplementary Figure 6F-G**, separate gels with K{N_3_}15-peptide and the AK{N_3_}15-peptide, as in **Fig. 3G** and Fig. 3**I** were used after trimming them to 2 x 2 mm flat/top surface area. In this case, gels of the same peptide and gelation solution were all placed in a microcentrifuge tube for CuAAC with alkyne-Atto647N fluorophore prior to in-gel Edman degradation. For this, a master mix with 603 μL of Copper(II)-THPTA catalytic buffer, 1.5x (Lumiprobe, K5150) was added to 1.8 μL of alkyne-Atto647N (Leica Microsystems, AD-647N-145), 277.2 μL of water. Then 96 μL of this master mix was added to each of the gels in the microcentrifuge tubes. Each tube was purged with nitrogen for 10 seconds, followed by the addition of 2 μL of freshly prepared 50 mM sodium ascorbate (SB0830, Bio Basic) in water, mixing and another purging with nitrogen for 10 seconds. These gels were then incubated overnight at room temperature to react. The next day, they were washed with 1X PBS 3x 10 min at RT prior to running in-gel Edman degradation, as in the section **In-gel Edman degradation**. The gels were washed with 1X PBS 3x 10 min at RT prior to imaging.

### Amino acid read-outs: ClpS2 St-V1 N-terminal binding and Glyphic antibody

#### Binding ClpS2 St-V1 and anti hemagglutinin-tag (HA) antibody to different N-terminal peptides in gel

For **Fig. 4A-D**,**H-I** ExM gels were cast at a concentration of 1mM peptide in the gelation solution, for all peptides: F_1_ peptide, W_1_ peptide, Y_1_ peptide, A_1_ peptide, L_1_ peptide. This was done with 3 different aliquots of 0.8 mg lyophilized peptide (from AAPPTEC, > 95% purity) reconstituted to a concentration of 5 mM in water. The gels were made using a ∼130 μm thick gelation chamber (i.e., 1 parafilm thickness). The gels were then expanded and re-embedded, as detailed in **Methods: ExM gel: making the gelation mixture with peptide** and **Re-embedded ExM (ExMre) gels** using three separate gelation solutions. The gels were cut to 2 x 2 mm top/flat surface size and stored at 4°C in 1X PBS, reaching a thickness of ∼350 μm after expansion, with each gel containing ∼70 pmol peptide for **Fig. 4A-D, F-I**. For **Supplementary Figure 7A**, the same peptides were reconstituted to a concentration of 25 mM in water to make 9% acrylamide gels using ∼260 μm thick gelation chamber (i.e., 2 parafilm thickness) to reach a thickness of ∼310 μm, cut to 2 x 2 mm with each gel containing ∼4 nmol peptide. For **Fig. 4A-D, F-G,** after 3 times 1X PBS washes, 10 min each (after the in-gel Edman degradation for **Fig. 4F-G**), these gels were then placed in plastic microcentrifuge tubes (VWR, 87003-292) with 500 μL water and washed 3 times, 5 minutes each, with 500 μL water.

A stock of ClpS2 St-V1 in a buffer of 50 mM sodium phosphate, 150 mM NaCl, pH 7.4 was made. The stock vial was spun down to remove any protein aggregates, and a concentration of ∼116 μM was measured in the supernatant using NanoDrop (molecular weight of ClpS2 St-V1 is 13034 g/mol, and an extinction coefficient of 8940 M^-1^cm^-1^ at 280 nm). The protein sequence ClpS2 St-V1 (and for ClpS2, and ClpS2 V1) is in **Supplementary Note 6.2**. The protein was aliquoted and stored at -20 °C. Before use, the protein was diluted down to 10 μM in a buffer solution of 0.1% BSA in water, with ∼4.3 mM sodium phosphate, 13 mM NaCl (for **Fig. 4A, F-G**), or 20 μM in a buffer solution of 0.1% BSA in water, with 8.6 mM sodium phosphate, 26 mM NaCl (for **Supplementary Figure 7A**). 30 μL of the solution was incubated with each gel for 6h (for **Fig. 4A, F-G**) or 1h (for **Supplementary Figure 7A**). For **Fig. 4A,F-G**, gels were washed with 0.1% BSA (Fisher Scientific, #NC1303417) in 1X PBS for ∼1-3 min, and replaced with a 30 μL solution of 10 μg/mL (67 nM) (for **Fig. 4A**) or 20 μg/mL (133 nM) (for **Fig. 4F-G**) of anti-HA tag antibody 488 (ThermoFisher, Catalog # 26183-A488) incubated at 4 °C for 7 days (for **Fig. 4A**) or 3 days (for **Fig. 4F-G**). For **Fig. 4A**, after 5 days of antibody incubation, the gels were mixed with a pipette tip to make sure they were not stuck to one side of the plastic tube (which could disrupt equal diffusion of protein from either side of the gel). For **Fig. 4F-G**, after 1 day of antibody incubation, the gels were mixed with a pipette tip to make sure they were not stuck to one side of the plastic tube (which could disrupt equal diffusion of protein from either side of the gel). For **Supplementary Figure 7A**, gels were washed with 0.1% BSA in 1X PBS for ∼1-3 min, and replaced with 100 μL solution of 200 μg/mL anti-HA tag antibody 647 (1.3 μM) (ThermoFisher, Catalog # 26183A647) incubated at 4 °C overnight (O/N).

#### Binding to HA-tagged peptides in acrylamide gels

For **Supplementary Figure 7B**, the HA-tagged peptides were reconstituted to a concentration of 5 mM in water to make 9% acrylamide gels using ∼260 μm thick gel chambers (i.e., 2 parafilm thickness; as detailed in **Methods: Acrylamide gel: making the gelation mixture with peptide**). The gels were cut 2 x 2 mm top/flat surface size and stored at 4 °C in 1X PBS (reaching a thickness of ∼310 μm in 1X PBS, with each gel containing ∼750 pmol peptide). After 3 times 1X PBS washes, 10 min each, these gels were then placed in plastic microcentrifuge tubes with 500 μL water and washed 3 times, 5 minutes each, with 500 μL water. 30 μL of the 20 μM ClpS2St-V1 was incubated with each gel for 1h. The gels were washed with 0.1% BSA (Fisher Scientific, #NC1303417) in 1X PBS, and replaced with 30 μL solution of 20 μg/mL anti-HA tag antibody 647 (ThermoFisher, Catalog # 26183A647) incubated at 4 °C O/N.

#### In-gel Edman degradation for binding experiments with F_1_ peptide and G_1_F_2_ peptide in ExMre gels

ExM gels were cast at a concentration of 1mM peptide, for both peptides: F_1_ peptide, and G_1_F_2_ peptide using 3 different aliquots of 0.8 mg lyophilized peptide (from AAPPTEC, > 95% purity) reconstituted to a concentration of 5 mM in water. The gels were made using 1 parafilm thickness gel chamber. The gels were expanded and re-embedded (as detailed in **Methods: ExM gel: making the gelation mixture with peptide** and **Re-embedded ExM (ExMre) gels**). ExMre gels were cut 5 x 5 mm squares and stored at 4 °C in 1X PBS. In-gel Edman degradation was performed on the gels as previously described (see section **In-gel Edman degradation**), except for the two following modifications: 1:9 of PITC:DMSO by adding 30 μL PITC to 270 μL DMSO, and with only 2 washes with DMSO, 5 min each, at 50 °C, between conjugation (PITC) and cleavage (TFA) steps. After TFA was removed, the gels were washed 3 times with 1M Tris pH 8, 10 min each, at RT. The gels for “PITC:DMSO+TFA+trypsin” condition were then submerged in 50 μL of 20 μg/mL trypsin in 50 mM Tris-HCl pH 8 in the bottom of the 1.5 mL Eppendorf tube (ensuring they were completely submerged in trypsin solution). The gels for other conditions were submerged in 50 μL of 50 mM Tris-HCl pH 8. All gels were then incubated for 2 hours at 37 °C. Each ExMre gel was further cut into 2 x 2 mm squares on a glass slide.

#### Binding tvClpS2 Q31H and anti 6x His tag antibody to different N-terminal peptides in gel

For **Fig. 4Hi-iv**, 2 x 2 mm top/flat surface size previously made for F_1_ peptide, W_1_ peptide, Y_1_ peptide, A_1_ peptide, L_1_ peptide, and no peptide gel were placed in plastic microcentrifuge tubes (VWR, 87003-292) after 2 times 1X PBS washes, 5 min each at RT. Then 30 μL of 20 μM of tvClpS2 Q31H (from a stock of 216.8 μM in 1X PBS with 2 mM DTT and 0.5 mM EDTA) in 1X PBS 0.1% BSA was added to the gels and incubated for 4 hours at RT. Then, gels were washed with 0.1% BSA in 1X PBS for ∼1-3 min, and replaced with a 30 μL solution of 20 μg/mL anti 6x His tag antibody 488 at 4 °C overnight (**Fig. 4Hi-iv**).

#### Sulfur-triazole exchange (SuTEx) covalent probe to tyrosine

For **Fig. 4Ii-iv**, 2 x 2 mm top/flat surface size previously made for F_1_ peptide, W_1_ peptide, Y_1_ peptide, A_1_ peptide, L_1_ peptide, and no peptide gels, from three separate gelation solutions, were placed in plastic microcentrifuge tubes (VWR, 87003-292) after 2 times 1X PBS washes, 5 min each at RT. The purified SuTEx-azide probe was resuspended in ACN at a final concentration of 50 mM. The probe was further diluted to 3 mM in 1X PBS and 50 μL of this final solution was added to each gel. The gels were incubated at 37°C for 2 hours. The gels were then washed 3x 1X PBS with 400 μL for 10 min at RT. 100 μL of 20 μg/mL DBCO-AF488 in 1X PBS was added to each gel for overnight (∼16 hours) incubation at 4°C. The gels were washed 3x 1X PBS 400 μL for 10 min at RT prior to imaging.

#### In-gel Edman degradation with SuTEx-azide covalent probe experiments with Y_1_ peptide in ExMre gels

For **Fig. 4Iv-viii**, 2 x 2 mm top/flat surface size gels previously made for Y_1_ peptide, from three separate gelation solutions, were placed in plastic microcentrifuge tubes (VWR, 87003-292). In-gel Edman degradation was performed on these gels as previously described (see section **In-gel Edman degradation**). The gels for “PITC:DMSO+TFA+trypsin” condition were then submerged in 50 μL of 20 μg/mL trypsin in 50 mM Tris-HCl pH 8 in the bottom of the 1.5 mL Eppendorf tube (ensuring they were completely submerged in trypsin solution). The gels for other conditions were submerged in 50 μL of 50 mM Tris-HCl pH 8. All gels were then incubated for overnight (∼16 hours) at 37 °C, then washed 3x 10 min at RT with 400 μL 1X PBS. The gels were incubated with 50 μL of 3 Mm SuTEx-azide probe (from a 100 mM stock in ACN) at 37°C for 2 hours. The gels were washed 3x 1X PBS with 400 μL for 10 min at RT prior to staining with 250 μL of 20 μg/mL DBCO-AF488 in 1X PBS for overnight (∼16 hours) incubation at 4°C. The gels were washed 3x 1X PBS 400 μL for 10 min at RT prior to imaging.

#### Multiplexed in-gel Edman degradation with SuTEx-azide covalent probe with F_1_Y_2_L_3_ peptide in ExMre gels

For **Fig. 4Ji-iv**, the F_1_Y_2_L_3_ was cast at a concentration of 1mM peptide in the gelation solution. This was done with an aliquot of 4 mg lyophilized peptide (from Genscript, FYLAGRGLGK{acr} > 95% purity) reconstituted to a concentration of 5 mM in water. The gels were prepared similarly to **Fig. 4Iv-viii**, including by first cutting gels to 2 x 2 mm top/flat surface size, then performing in-gel Edman degradation followed by trypsinization of “PITC:DMSO+TFA+trypsin” condition. The gels were then cut into two 2 x 1 mm top/flat surface size and placed in separate microcentrifuge tubes for washing 3x 10 min at RT with 400 μL 1X PBS: one set of the gels was incubated with SuTEx-azide, as in **Fig. 4Iv-viii**, and the other set was incubated with ClpS2 St-V1, as in **Fig. 4F-G**, but the antibody was incubated 2 days at 4°C before imaging (rather than 3 days in **Fig. 4F-G**).

#### Glyphic binder specificity in gel and Biolayer Interferometry (BLI) curves

As depicted in **Fig. 6D**, antigens from Glyphic Biotechnologies were tested by embedding streptavidin in ExMre gels and subsequently adding biotinylated ClickP-amino acid (abbreviated: biotin-ClickP-aa). For AcX modification of streptavidin, 1.5 μL of 10 mg/mL AcX, (ThermoFisher, A20770) stored at -20 °C in DMSO, was added to 25 μL of 10 mg/mL streptavidin (ThermoFisher, 21122) resuspended in water and stored at 4 °C, to a total volume of 50 μL in 0.1 M NaHCO_3_ pH 8.5. This solution was left to incubate overnight at room temperature. Using the ∼5 mg/mL AcX-modified streptavidin stock, ExM gels were cast, using ∼130 μm thick gelation chamber (i.e., 1 parafilm thick), to a final concentration of ∼0.1 mg/mL AcX-modified streptavidin. The gels were expanded and re-embedded (as in **Methods: ExM gel: making the gelation mixture with peptide** and **Re-embedded ExM (ExMre) gels**), to an approximate concentration of ∼5 μg/mL (0.1 mg/mL / 2.7^3^) and stored in 1X PBS at 4 °C. The ExMre gels were cut into 2 x 2 mm squares, placed in separate plastic microcentrifuge tubes, and washed 3 times with water. The gels were incubated with biotin-ClickP-aa (where aa: G, V, F, L, I, K{ac}) at a concentration of ∼16.7 μg/mL (for **Fig. 6Ei-iii** and **Fig. 6Fi-iii**) or 5 μg/mL (for **Fig. 6Eiv-vi**, **Fig. 6Fiv-vi, Fig. 6Gi-iii, Fig. 6Hi-iii** and **Fig. 6Ii-iii**), freshly made from a stock of 0.5 mg/mL for G, V, F, 0.56 mg/mL for L, 0.71 mg/mL for I and 1.14 mg/mL for K{ac} (from Glyphic) in a total volume of 100 μL in 1X PBS with 0.1% BSA for 24 hours at room temperature. These gels were washed 3 times, 20 min each, with 400 μL 0.1% BSA in 1X PBS. For **Fig. 6ei-iii** and **Fig. 6fi-iii** they were then incubated for 2 days with 30 μL of 10 μg/mL RFP-conjugated Glyphic antibody against ClickP-F (abbreviated: Glyphic F) or 5 μg/mL GFP-conjugated Glyphic antibody against ClickP-V (abbreviated: Glyphic V) in 0.1% BSA in 1X PBS at 4 °C. For **Fig. 6Eiv-vi**, **Fig. 6Fiv-vi, Fig. 6Gi-iii, Fig. 6Hi-iii** and **Fig. 6Ii-iii,** they were incubated for 4 days with 30 μL of 5 μg/mL primary Glyphic antibodies (without fluorescent proteins) in 0.1% BSA in 1X PBS at 4 °C, followed by 4 times 30 min 400 μL 0.1% BSA in 1X PBS washes.

These gels were then incubated with secondary antibodies at 1:200 dilution in 0.1% BSA in 1X PBS at 4 °C for 3 days, with anti-mouse 488 antibody (ThermoFisher: Catalog # A32766TR) on the anti V, F, L, K{ac} Glyphic antibodies, and anti-rabbit 488 (ThermoFisher: Catalog # A32790TR) on the anti I Glyphic antibody. Prior to imaging, the gels were washed with 400 μL one time 40 min, followed by one time 20 min in 0.1% BSA in 1X PBS at RT. BLI results of the Glyphic antibodies in **Supplementary Figure 9** were provided by the company.

### Imaging

All imaging was performed using a spinning disk confocal microscope (Andor Dragonfly Spinning Disk confocal).

For **Fig. 3G**,**I** **Fig. 4I**, **Supplementary Figure 6F-G**, the gels were placed on a 24-well glass plate with 1 mL 1X PBS per well. Before imaging, the 1X PBS solution was removed. For **Fig. 3G**, and **Supplementary Figure 6F-G,** the gels were imaged using a 10X magnification lens acquiring a Z-stack covering the whole gel thickness, with a 10 μm Z-step size (520 μm total Z-stack, except condition “DMSO”, replicate 3, with a 510 μm total Z-stack for **Fig. 3G**), 40% laser power, 100 ms exposure time (**Supplementary Figure 6G**), 100% laser power, 500ms (**Supplementary Figure 6E**) with 488 laser line and 525/50 emission filter. For **Fig. 3I**, **Fig. 4I**, the gels were imaged using a 40X magnification lens, acquiring a Z-stack covering the whole gel thickness, with a 10 μm Z-step size (390 μm total Z-stack and 100% laser power for **Fig. 4Iii-iv**, and 350 μm total Z-stack and 100% laser power for **Fig. 4Iv-viii;** 660 μm total Z-stack and 40% laser power for **Fig. 3I**, except condition “TFA”, replicate 2, with a 650 μm total Z stack) 100 ms exposure time, with 488 laser line and 525/50 emission filter.

For **Fig. 4B-D**, and **Fig. 4F-H**,**J,** the supernatant antibody solution was first removed from the gels via a transfer pipette before imaging. They were placed on a droplet of 50 μL of 1X PBS with 0.1% BSA on a 24-well plate for the smooth transfer of the gel, and the droplet was then removed after the transfer. Imaging was performed with 488 laser and an emission filter of 525/50, with 100% laser power and 500 ms exposure time, using a 40X objective (water immersion) and 10 μm step size (**Fig. 4B-D, H**) or 15 μm step size (**Fig. 4F-G**) covering the entirety of the gel cross-section (for a final ∼430 μm Z-stack for **Fig. 4B-D**; **∼**375 μm Z-stack for **Fig. 4F-G**; ∼450 μm Z-stack for **Fig. 4H**; ∼400 μm Z-stack for **Fig. 4J**).

For **Fig. 6E-I**, the gels were placed on a 24-well plate on a droplet of 50 μL 1X PBS 0.1% BSA in the center of the well to transfer the gel before imaging. Imaging was performed with a 40X objective (water immersion), and a Z-stack obtained from 10 μm step size across the depth of the gels (a final 400 μm Z-stack for **Fig. 6Ei-iii, Fig. 6Fi-iii;** a final 300-490 μm Z-stack for **Fig. 6Eiv-vi, Fig. 6Fiv-vi, Fig. 6Gi-iii, Fig. 6Hi-iii and Fig. 6Ii-iii** which were then normalized to % cross-section) using 500 ms exposure time, 100% laser power with excitation wavelength in 488 channel and an emission filter of 525/50 (for the GFP-conjugated Glyphic V antibody; and secondary antibodies 488) or 561 and an emission filter of 582/15 (RFP-conjugated Glyphic F antibody).

For **Supplementary Figure 7A**, the imaging was performed after removing the antibody (anti-HA tag antibody 647) solution using a confocal microscope with 10X objective, 100% laser power in 637 laser with 676/37 emission filter, 400 ms exposure time, 10 μm Z-steps covering the entire gel cross-section.

For **Supplementary Figure 7B**, the imaging was performed after removing the antibody (anti-HA tag antibody 647) solution using a confocal microscope with 10X objective, 100% laser power in 637 laser with 676/37 emission filter, 100 ms exposure time, 10 μm Z-steps covering the entire gel cross-section.

### Image analysis

For **Fig. 3G**, **Fig. 4B-D**, **Fig. 4F-I**, **Figure 6Ei-iii,Fi-iii, Supplementary Figure 6F-G** images (‘.ims’ files) were opened in FIJI and gel Z-stacks were inspected by plotting the Z-axis profile. The final Z-stack thickness for all conditions was set to capture the entire thickness of the gel and was made identical for each replicate (see **Imaging** section). For **Figure 6Eiv-vi,Fiv-vi,G,H,I,** the images were similarly opened in FIJI and gel Z-stacks were inspected, but the entire thickness of the gel was not identical for each replicate, and ranged from 300 μm to 490 μm. As the gel thickness varied across replicated, the cross-sectional position of each Z-slice was normalized to a common percentage scale, where 0% corresponds to the top surface of the gel and 100% corresponds to the bottom surface. This normalization was performed by dividing each slice position (in μm) by the total thickness of that replicate’s gel, allowing fluorescence intensity profiles to be directly compared across replicates regardless of absolute gel thickness (see **Imaging** section). All images were then saved as ‘.tif’ files.

The plots of gel fluorescence throughout the Z-stack (**Fig. 3Gii**, **Fig. 4C, Fii, Gii, Hiii, Iiii, Ivii, Jiii, Jvi** and **Fig. 6Eii,v, Fii,v, Gii, Hii, Iii**) and bar plots of the average fluorescence throughout the gel volume (**Fig. 3Giii**, **Fig. 4D, Fiii, Giii, Hiv, Iiv,viii, Jiv,vii,** and **Fig. 6Eiii,vi, Fiii,vi, Giii, Hiii, Iiii**) were made in MATLAB (see GitHub **Code**). For the Z-stack plot showing fluorescence throughout the gel, the mean and standard deviation of fluorescence for each Z-slice (over the replicates of the same condition for the same Z-slice) was calculated and plotted in different colors for each condition. For the bar plots, the average fluorescence of the gels across the whole volume was computed (i.e., for each Z-slice of a given sample, the mean fluorescence intensity of all pixel values was computed.

Then, for each sample, the average intensity across all its Z-slices was calculated, and plotted as a separate data point. Finally, the average intensity for each condition was obtained by computing the mean of these sample-wise average intensities).

The representative images of each gel (**Fig. 3Gi, 4B, 4Fi, 4Gi, 4Hii, 4Iii,vi, 4Jii,v 6Ei,iv, 6Fi,iv, 6Gi, 6Hi, 6Ii**) was obtained from one gel replicate for a given condition. The same arbitrarily selected depth, reported in the figure captions, was taken for each condition by making a substack of a single Z-slice (e.g. to obtain the image for Z-slice # 20: Image → Stacks → Tools → Make Substack, input “20-20”). A scale bar was also added to every image representing the first condition of each experiment. These average intensity profiles for each condition were then plotted in Python using the *mpimg* Matplotlib image module, with an intensity scale for comparison (see GitHub **Code**).

### Oxidation analysis in ExM gels

Amino acid oxidation analysis in ExM gels were performed with XaaGGAGRGLGK{acr} peptides, where Xaa was one of 8 amino acids: M, C, W, Y, F, P, H, R (as described in **Synthetic peptide designs** section). Post-translational oxidation modifications for each amino acid can be found in **Supplementary Figures 8A-H**.

Three conditions were prepared for each of the N-terminal acid peptides. Two in solution, where condition 1 was: 20 μL of 5 mM peptide with 90 μL water, and condition 2 was: 20 μL of 5 mM peptide with 80 μL water, 5 μL of 10% TEMED (v/v), and 5 μL 10% APS (w/v), incubated at 37 °C for 1h30m. The condition 3 was ExM gelation: 20 μL of 5 mM peptide with 60 μL of StockX, 10 μL of 130 mM BIS, 5 μL of 10% TEMED (v/v), and 5uL 10% APS (w/v) gelled into 20 x 10 mm surface size chamber with ∼260 μm chamber thickness (i.e., 2 parafilm thickness). Gelation was performed as described in section **ExM gel: making the gelation mixture with peptide**.

Condition 1 and condition 2 were further diluted by taking 15 μL of the sample and adding 50 μL 20 μg/mL trypsin in 1M Tris pH 8 for a total volume of 65 μL.Condition 3 gels were cut into 5 mm x 10 mm ExM gel using a razor blade (right after gelation; without expanding) and washed twice with 1X PBS, 5 min each, then once with 1M Tris pH 8 for 5 min. The wash solution was then replaced with 50 μL of 20 μg/mL trypsin in 1M Tris pH 8 on top of the gel. Condition 1, condition 2 and condition 3 were all incubated for 4 hours at 37 °C with the trypsin solution. After trypsinization, the samples were placed in the fridge until analysis on LC/QToF. 50 μL of the solutions, for each condition, were placed in separate mass spec vials for analysis. Condition 1, 2, and 3 samples were injected into the LC/QToF (see section on **Mass spectrometry**).

Analysis of the N-terminal amino acid oxidation was performed as detailed in **Analysis of LC/QToF data** using the information about post-translational oxidation modification of peptides documented in **Supplementary Table 3**. For each N-terminal amino acid, the abundance (a.u.) of the original fragment and the fragment with various post-translational oxidation modifications was extracted for the three different sample conditions and documented in a table (see **Supplementary Figures 8Aii-Hii**). Then, using these values, the relative abundance of each of the fragments were compared using a bar graph. For this, the abundance of each species was normalized with max absolute scaling for plotting (see **Supplementary Figures 8Aiii-Hiii**). The raw total ion chromatograms and mass spectra were exported as metafiles (‘.emf’), and available in **Source Data**.

### Theoretical assessment of *in situ* protein sequencing

All analyses were performed either on mycoplasma genitalium proteome (483 Swiss-Prot reviewed proteins; UniProt ID 243273: by searching “taxonomy_id:243273” in UniProt), or the human proteome (20,421 Swiss-Prot reviewed proteins, UniProt ID 9606: by searching “taxonomy_id:9606” in UniProt). All references throughout for mycoplasma proteome and the human proteome are for these two proteomes, respectively. All code can be found on GitHub, at: https://github.com/camimarie/insituprotein/. Amino acid residues are abbreviated using their single letter amino acid code. N-terminal is abbreviated “N-term”.

### Part 1: results of exploring the chemistries and their effects on preserving protein sequence coverage in the gel

#### Percent amino acid count

For **Supplementary Figure 13C** (mycoplasma proteome) heatmap, the protein database went through a three step chemical process that modified the primary sequence of amino acids. A gridsearch was performed with various combinations of P(fix), P(anchor), P(digest), where these represent the probability of these three chemical steps to modify a subset of amino acid side chains. In addition, the anchoring could be either AcX or epoxide, and digestion could be either endoprotease Lys-C (abbreviated Lys-C), Proteinase K (abbreviated ProK) or trypsin. The gridsearch values were: P(fixed): 0.00, 0.01, 0.05, 0.10, 0.50, 1.00; P(anchored): 0.0, 0.1, 0.2, 0.3, 0.4, 0.5, 0.6, 0.7, 0.8, 0.9, 1.0; P(digestion): 0.0, 0.1, 0.2, 0.3, 0.4, 0.5, 0.6, 0.7, 0.8, 0.9, 1.0. First, the proteome (mycoplasma or human) was selected from UniProt and loaded into the computational environment. Then, N-term, K, C, R, and Y residues were modified with probability P(fixed), where they were replaced with a “J” (arbitrarily, since “J” is not overlapping with any other amino acid single letter code) after fixation.

Similarly, for anchoring, N-term and K residues for AcX, or N-term, K, C, H, Y, D and E for epoxide, were modified with probability of P(anchored), and replaced with a “Z” (since “Z” was not overlapping with any other letter). Lysine residues that were already converted to a J could not be modified to a Z in this anchoring step). For digestion, residue K was digested at its C-terminus for Lys-C, residues R and K were digested at their C-terminus for trypsin, whereas for ProK, Y, W, E, L, V, A, I, F, T residues were digested at their C-terminus. For all digestion types (Lys-C, trypsin, ProK), digestion did not occur if they had already been converted to a J (fixed) or Z (anchored). A fragment was only retained if it had at least one anchored residue to the gel matrix from the P(anchor), otherwise it was considered lost. Each grid in the grid search represents a unique combination of those three parameters, where each unique combination was run 10 times. The “percent amino acid count” was calculated as the percentage of residues of a protein that are not modified (i.e., remains a canonical amino acid), and remain accessible for read-out over 15 rounds of in-gel Edman degradation after the chemical steps. This was computed by summing over all the amino acids of a fragment that are unmodified by any chemical process, stopping at the last anchor of a fragment (subsequent amino acids would be lost and not be readable by in-gel Edman degradation) and, if the final anchor is not within the first 15 amino acids of the fragment, stopping at the 15th amino acid.

The N-term modifications in either the fixation or anchoring steps prevented read-out of the given fragment, for the mycoplasma proteome. The mean percent amino acid count was calculated by averaging over the 10 replicated computational experiments and also over the proteome and plotted on the heatmap in **Supplementary Figure 13C.**

For **Supplementary Figure 18** (human proteome), it was the same as described for **Supplementary Figure 13c** (mycoplasma proteome), but the N-terminal fragment was always considered inaccessible due to N-terminal acetylation. In addition the gridsearch values for P(fixed), P(anchored) and P(digestion) were the same values as above.

For **Supplementary Figure 14A** and **Supplementary Figure 14C**, the distribution of the percent amino acid count (defined above) was plotted as a distribution for all the proteins in the proteomes averaged over the 10 rounds of computational experiments for P(fixation)=0.05, P(anchoring)=0.80, P(digestion)=0.80, for both mycoplasma and human proteomes, where the error bar represented the standard deviation over the multiple computational repetitions. For **Supplementary Figure 14B, Supplementary Figure 14D**, the distribution of the percent amino acid count was plotted as a distribution for all the proteins in the proteomes over the 10 rounds of computational experiments for the highest percent amino acid count for each condition (best performing parameters given the condition) as calculated and plotted in **Supplementary Figure 13C** and **Supplementary Figure 18** (for the mycoplasma proteome and human proteome, respectively). The bin-size is 40.

#### Fragment lengths

For **Supplementary Figure 15**, the fragment length was defined as the number of amino acids for a given fragment retained in the gel, including amino acids that are fixed or anchored (unlike the percent amino acid count), but not considering amino acids that are located downstream, C-terminal, of the last anchoring amino acid of the fragment. We also do not cap at 15 amino acids, unlike the percent amino acid count. The results for these fragment lengths were plotted as distributions of protein count versus fragment length, where the mean and median values were recorded by plotting a vertical line along with their values. The y-axis was log-scale, and standard deviations were the result of 10 computational repetitions, with bin-size of 40. Results of the fragment lengths using a parametrization of P(fixation)=0.05, P(anchoring)=0.8, P(digestion)=0.8, were plotted for the mycoplasma proteome, human proteome (**Supplementary Figure 15A-B**, respectively), which were the same parameterizations used in **Part 2**.

### Part 2: Simulating *in situ* protein sequencing with NAABs and in gel Edman degradation Reference fragment dataset (*T_s_*)

The reference fragment dataset represents the database of ground-truth sequences present in the gel, after fixation, anchoring, gelation, digestion and expansion, but prior to the *in situ* protein sequencing read-out. To generate the dataset, first, similarly to **Part 1** methods, the mycoplasma proteome was selected from UniProt and loaded into the computational environment. Then, the primary sequence of amino acids was modified according to parameters for P(fixed), P(anchored), P(digestion), where these represent the probability of these three chemical steps to modify a subset of amino acid side chains. However, in **Part 2**, these values were fixed with P(fixed)=0.05, P(anchored)=0.8, P(digestion)=0.8 using paraformaldehyde, AcX and trypsin for fixation, anchoring and digestion in the model, respectively. The amino acid side chain specificities of the paraformaldehyde fixation, anchoring with AcX, and digestion with trypsin, remained the same as in **Part 1**. Only fragments with at least one anchored residue to the gel matrix from the P(anchoring) were retained. Subsequently, an idealistic in-gel Edman degradation read-out was conducted on the remaining fragments by considering all the sequencing chemistries, including the conjugation and cleavage of in-gel Edman degradation to be 100%. In addition, the idealistic read-out could detect all amino acids with 100% accuracy, except ones that have been fixed or anchored at previous steps (fixation and anchoring). In this way, the fragments generated were perfectly corresponding to the ground-truth sequence that would be expected when reading them out in idealistic conditions. In addition, if Edman leads to the loss of the only anchor, amino acids following are gone and will be substituted with an unknown read (“X”) since there will be no further amino acids to readout. After, amino acid sidechains modified to J or Z are unrecognized by binders, and thus will return X (given that modifications like J and Z cannot be distinguished during read-out, and will also result in failure to bind) and stored as a fragment. This was performed 1,000 times, and the fragments from these computational experiments were then aggregated into the reference fragment dataset, also named ***T_s_*** (peptides anchored in the expansion gel independent of read errors). For **Supplementary Figure 16**, the number of unique fragments generated were compared for 1, 10, 100, 1,000, and 10,000 simulations and plotted for mycoplasma.

#### Error-prone fragment dataset and assignment to proteins

The proteome (mycoplasma) was selected from UniProt and loaded into the computational environment. The first steps of the error-prone fragment dataset were the same as the reference fragment dataset generation up until the modification of the primary protein sequences into fragments through digestion (with values for P(fixed)=0.05, P(anchored)=0.8, P(digestion)=0.8 using paraformaldehyde, AcX and trypsin for fixation, anchoring and digestion in the model, respectively). Only fragments with at least one anchored residue to the gel matrix from the P(anchoring) were retained. After, amino acid sidechains modified to J or Z are unrecognized by binders, and thus will return X (given that modifications like J and Z cannot be distinguished during read-out). At this stage, instead of considering an idealistic in-gel Edman degradation read-out, the read-out was simulated with errors, including the sequencing chemistry and the binder kinetics.

#### Sequencing chemistry errors

Specifically, for the sequencing chemistry errors, the conjugation of PITC was considered to have a probability of success of 0.9, and the cleavage with TFA with a 0.7 success rate. Amino acids with a modified sidechain (by anchoring or fixation) were considered to be cleavable via in-gel Edman degradation, but when reaching the last anchor of the peptide, the rest of the fragment was lost and subsequent amino acids would be read as unknown (i.e. “X”).

#### Binder read-out errors and error-prone fragment dataset (*F_s_*)

For simulation of the binding, after the PITC conjugation step, binders were bound probabilistically to PITC-conjugated N-terminal amino acids, and this was performed sequentially for every binder. Results were performed with several subsets of binders that recognized various PITC-conjugated N-terminal amino acids. The 20 amino acid binder subset with recognizers towards all canonical amino acid side-chains (but not the X sidechains, of course). The 15 amino acid binder subset contained the following recognizers: {A, N, D, E, Q, G, I, L, K, F, S, T, P, Y, V}, the 10 amino acid binder subset contained the following recognizers: {A, N, D, E, Q, I, L, K, F, V}, and the 5 amino acid binder subset contained the following recognizers: {N, E, L, F, V}. The binding was simulated in a condition with excess binder, and where binding reaches equilibrium (using the Langmuir equation for binding), such that after the washing, the probability of being bound to on-target and off-target was defined as: 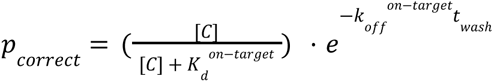and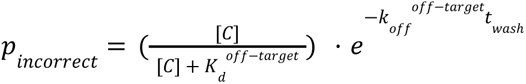, where [*C*] is the concentration of binder, 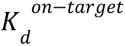 the equilibrium dissociation constant for the on-target, 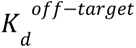 the equilibrium dissociation constant for the off-target, 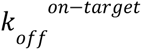 the dissociation rate for the on-target, and 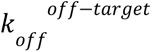 the dissociation rate for the off-target, and *t _was_*_ℎ_ is the time elapsed during the dissociation phase (during the washing). **Supplementary Figure 17** was plotted with these equations (plotting *p_correct_* - *p_incorrect_*) by modifying the values of binder washing, *t_wash_* from 0 to 200 min, and the concentration of binder, [*C*], from 1 pM to 100 mM. These were plotted for a binder with different levels of specificity according to table **Supplementary Table 12**: very high, high, medium, low, and very low specificities, where the equilibrium dissociation constant and dissociation rates for on-target and off-target were modified in the equation for each of these specificity levels. Very high, high, and medium resulted in the same on-target and off-target binding profiles, and thus only one (medium) specificity was included in **Supplementary Figure 13D**. For the results in **Supplementary Figure 13D** (mycoplasma proteome), the *t_wash_* was considered as 30 min, and the [*C*] as 1 μM, and results were generated for different levels of binder specificities according to table **Supplementary Table 12**. For each in-gel Edman degradation round, independent reads were obtained for each amino acid (20, 15, 10, and 5 times) since the binding was performed sequentially for each binder. Thus, several read-outs were associated with the same position in a peptide fragment (this was taken into consideration for the downstream Hidden Markov Model (HMM)). After performing the binding, the N-terminal amino acid was cleaved with sequencing chemistry errors mentioned above, and the process was repeated from 5 to 15 rounds of in-gel Edman degradation and sequential binder read-out.

This resulted in error-prone fragment sequences for each protein, which collectively became the error-prone fragment dataset (*F_s_*).

### Hidden Markov Model (HMM) based matching and fraction of proteome correctly identified

#### Pre-filtering uninformative fragments

Following the generation of the error-prone dataset, which included for every protein in the mycoplasma proteome at each condition (number of rounds × number of binder cases × number of specificity cases × 10 repetitions = 1,760 conditions), low quality fragments were filtered. First, uninformative “query fragments”, originating from the error-prone dataset *F_s_*, were defined as fragments that consisted of more than half of their total Edman degradation rounds contained only unknown reads (“X”), e.g. rounds where no binder sequentially bound at all. These fragments were removed. Second, uninformative “reference fragments”, originating from the reference dataset *T_s_*, were defined as fragments that consisted of more than half of the total reads as unknown reads, denoted by an “X”. These fragments were also removed.

#### Algorithmic details for two-stage HMM-based matching

Next, for each remaining fragment, a two-stage HMM-based matching was performed. The HMM models the joint probability *P*(*F*|*T*, θ) that a true fragment *T* generates an observed fragment *F* given the experimental error parameters θ, which include the following:

● Conjugation failures (c): probability that no conjugation occurs in an Edman round.
● Cleavage failures (d): probability that the N-terminal amino acid is not cleaved.
^●^ Binder errors: per-binder probabilities describing on and off-target binding, where unbound is denoted as “X”: *p_bound_*_, *on*−*target*_, *p_unbound_*_, *on*−*target*_, *p_bound_*_, *off*−*target*_, *p_unbound_*_, *off*−*target*_ .

At each Edman round, each amino acid from the true fragment *T* emits *m* observations, one per binder in the binder sets described above. The model sums over all possible emission paths and state transitions, including insertion (failed cleavage) and stay (failed conjugation) transitions.

To map each observed fragment, subject to Edman conjugation failures, Edman cleavage failures, and binder errors, back to its source, we framed the problem as follows: for every “true” fragment *T* in the error-free trie, *T_s_* (reference fragment dataset), what was the probability that *T* under set parameters θ would generate the observed, error-prone fragment *F* in the error-prone dataset (*F_s_*). Thus, for each error-prone fragment *T*, the Viterbi log-likelihood, 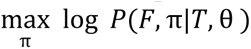 was computed, which represents the probability of the most likely path π∗ generating the observed fragment *F*, given each candidate sequence *T*. After computing these Viterbi scores for all candidates, we apply a pruning step to discard less likely candidates: 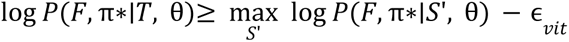, where *S*′ is the highest scoring fragment across all candidates. ɛ*_vit_* was set as 5.0, which was chosen to be deliberately wide as there is increased variance with certain conditions: in log-likelihood space, any candidate that scores 1/e^5^ ≈1/150 of the top score is retained. On the pruned candidate set, a forward dynamic programming algorithm is employed that computes the total likelihood 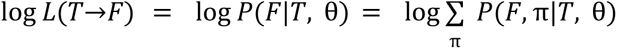, summing over almost all possible paths with a milder beam ɛ *_for_* = 10. 0. Note that no candidates are ever removed here, as the forward epsilon only prunes unlikely paths within the possible paths.

#### Fraction of the proteome correctly identified

For each of the top 10 candidate fragments by forward log-likelihood *T* ∈ *T_S_* (if such fragments exist and have not been filtered out), we log-summed fragment likelihoods weighted by counts using the reference fragment dataset’s protein occurrence counts for each fragment *T*. We then normalize and rank proteins by their total weight. The highest scoring protein is returned as the fragment’s originating protein only if the weight is at least 1.5x the second-highest scoring protein. Otherwise, the fragment is labeled “uncertain”. For each protein, we accumulate all the fragments’ labels. If no valid predictions are produced, we return “no predictions”. Otherwise, we count fragment-level votes per predicted protein: the protein is identified only if the top vote count is at least double the runner up, otherwise we return “uncertain”. We report the counts of “correct”, “false positive”, “uncertain”, and “no predictions”. From these decisions, the fraction correctly identified over the whole proteome was a metric for how many of the proteins in the mycoplasma proteome were correctly identified from the simulation of the error-prone fragments (not including uncertain or false positive matches). For **Supplementary Figure 13d**, the whole simulation was performed 10 times and the average and standard deviation of the fraction of correctly identified proteins was reported on the graph of results. In **Supplementary Figure 19**, we plot the average and standard deviation of the fraction of false positively identified proteins.

## Supporting information

Source Data For Specific Figures

## Acknowledgements

We thank E. Anslyn, J. Choi, A. Sinha, J. Chang, J. Mou, R. Raines, M. Kumar, S. Babakhanova for thoughtful discussions. We are grateful to J. Chang, J. Yao, K. Slaw for delivering the protein materials. We thank Jagannath Swaminathan from Erisyon Inc for providing us with the SuTEx-azide material, the synthesis of which was guided by Prof. Ken Hsu’s (University of Texas -Austin) lab. We also thank William Siegall and Prof. Zvi Kelman for providing us with the tvClpS2 Q31H protein material. We thank Z. Niu for contributing to the CuAAC chemistry in ExM gels and providing the sodium ascorbate. We would like to thank D. Sarkar and J. Kang for discussion around high-expansion gels to reach molecular resolution. For funding, E.S.B. acknowledges Ashar Aziz, Lisa Yang, John Doerr, HHMI, Jed McCaleb, James Fickel, the ERC Synergy Grant under the European Union’s Horizon 2020 research and innovation programme (grant agreement No 835102), NIH R01AG087374, R01EB024261, 1R01AG070831, Tom Stocky and Avni Shah, Kathleen Octavio, Lore McGovern, Open Philanthropy and Good Ventures, the Keck Foundation, and the Richard King Mellon Foundation. N.F.P acknowledges funding from NIH (R00GM135519). C.M.M. acknowledges MIT Health and Life Sciences Collaborative Graduate Fellowship (MIT HEALS) 2025/2026, Friends of the McGovern Institute Student Fellowship 2024/2025, MIT Center of Neurobiological Engineering Training Program 2023/2024, Brain and Cognitive Sciences (BCS) Hilibrand Graduate Student Fellowship 2021-2024. S.Z.T. acknowledges support from the Brain and Cognitive Sciences (BCS) Singleton Fellowship (2024/2025), the BCS Hubert J. P. Schoemaker (1976) Fellowship (2024–2025), and the MIT Praecis Presidential Fellowship (2023–2024). S.W. was supported by MIT Presidential Graduate Fellowship and Weedon Alzheimer’s Graduate Fellowship.

## Author information

### Contributions

C.M.M. and E.S.B. led the study, receiving early valuable insights from L.L.K, as well as early experimental input from C.Z., H.W., D.M.E., S.W.. C.Z. designed the initial experimental framework for in-gel Edman degradation. C.M.M., H.W., S.Z.T. designed the final experimental framework and the final protocol for in-gel Edman degradation. C.Z., C.M.M., H.W. contributed to the early conceptualization of the study, and performed early feasibility studies. Gel size changes were performed by C.M.M. and S.Z.T., and mass spectrometry was performed by C.M.M, S.Z.T., J.S. with experimental insights from C.Z., L.L.K.. Data analysis for gel size change and mass spectrometry was performed by C.M.M and S.Z.T.. Developing the in-gel Edman degradation protocol was done by C.M.M with help from H.W., S.Z.T., J.S. and J.P., and early input from C.Z., S.W., L.L.K.. ClpS binder protein and binder insight was provided by N.F.P and M.D.. Testing of ClpS binders was done by C.M.M., with early input from H.W. Amino acid oxidation experiments and data analysis was performed by C.M.M and L.E.. The creation and initial work on ClickP was performed by D.M.E. and A.G.C.. The development and testing of ClickP-AA binders was performed by D.M.E., A.G.C., E.W., and S.D.. Testing of Glyphic binders was done by C.M.M., with input from Glyphic. Imaging and imaging data analysis was performed by C.M.M.. Computational modeling was designed and developed by J.P. and C.M.M, performed by J.P., and written by C.M.M., J.P., E.S.B.. C.M.M and E.S.B wrote and edited other sections of the manuscript. T.S., S.W., C.M.M. and H.W. designed and performed the experiments to develop the high-expansion gels. C.M.M., S.Z.T., I.K. contributed to revisions on the experimental work and C.M.M., J.P. contributed to the revision on the computational work. E.S.B. supervised the project.

### Corresponding authors

Correspondence to Ed Boyden.

### Ethics declarations

ESB is a co-inventor on multiple patents related to expansion microscopy, and co-founder of a company seeking commercial interests in the space. CMM, CZ and ESB are co-inventors on a pending patent related to *in situ* single-molecule peptide sequencing. DME contributed to this manuscript prior to the founding of Glyphic Biotechnologies.

## References

1. Alberts, B., et al. Protein Function. Molecular Biology of the Cell. 4th edition (2002).

2. Haucke, V., Neher, E. & Sigrist, S. J. Protein scaffolds in the coupling of synaptic exocytosis and endocytosis. Nat. Rev. Neurosci. 12, 127–138 (2011).

3. Good, M. C., Zalatan, J. G. & Lim, W. A. Scaffold proteins: hubs for controlling the flow of cellular information. Science 332, 680–686 (2011).

4. DiRusso, C. J., Dashtiahangar, M. & Gilmore, T. D. Scaffold proteins as dynamic integrators of biological processes. J. Biol. Chem. 298, 102628 (2022).

5. Hardingham, G. E. & Bading, H. Synaptic versus extrasynaptic NMDA receptor signalling: implications for neurodegenerative disorders. Nat. Rev. Neurosci. 11, 682–696 (2010).

6. Chen, F., Tillberg, P. W. & Boyden, E. S. Expansion microscopy. Science 347, 543–548 (2015).

7. Tillberg, P. W. et al. Protein-retention expansion microscopy of cells and tissues labeled using standard fluorescent proteins and antibodies. Nat. Biotechnol. 34, 987–992 (2016).

8. Dolgin, E. ‘Expansion microscopy’ turns ten: how a tissue-swelling method brought super-resolution imaging to the masses. Nature 637, 752–754 (2025).

9. Chang, J.-B. et al. Iterative expansion microscopy. Nat. Methods 14, 593–599 (2017).

10. Sarkar, D. et al. Revealing nanostructures in brain tissue via protein decrowding by iterative expansion microscopy. *Nat*. Biomed. Eng. 6, 1057–1073 (2022).

11. Wang, S. et al. Single-shot 20-fold expansion microscopy. Nat. Methods 1–7 (2024).

12. Shaib, A. H. et al. One-step nanoscale expansion microscopy reveals individual protein shapes. Nat. Biotechnol. 1–9 (2024).

13. Alfaro, J. A. et al. The emerging landscape of single-molecule protein sequencing technologies. Nat. Methods 18, 604–617 (2021-6).

14. Reed, B. D. et al. Real-time dynamic single-molecule protein sequencing on an integrated semiconductor device. Science (2022) doi:10.1126/science.abo7651.

15. Swaminathan, J. et al. Highly parallel single-molecule identification of proteins in zeptomole-scale mixtures. Nat. Biotechnol. 10.1038/nbt.4278 (2018).

16. Rodriques, S. G., Marblestone, A. H. & Boyden, E. S. A theoretical analysis of single molecule protein sequencing via weak binding spectra. PLoS One 14, e0212868 (2019).

17. Ritmejeris, J., Chen, X. & Dekker, C. Single-molecule protein sequencing with nanopores. Nat Rev Bioeng 3, 303–316 (2024).

18. Wang, G., Moffitt, J. R. & Zhuang, X. Multiplexed imaging of high-density libraries of RNAs with MERFISH and expansion microscopy. Sci. Rep. 8, 4847 (2018).

19. Alon, S. et al. Expansion sequencing: Spatially precise in situ transcriptomics in intact biological systems. Science 371, eaax2656 (2021).

20. Labade, A. S. et al. Expansion in situ genome sequencing links nuclear abnormalities to hotspots of aberrant euchromatin repression. bioRxivorg 2024.09.24.614614 (2024) doi:10.1101/2024.09.24.614614.

21. Edman, P., Högfeldt, E., Sillén, L. G. & Kinell, P. O. Method for Determination of the Amino Acid Sequence in Peptides. Acta Chemica Scandinavica 4, 283–293 (1950).

22. Edman, P. & Begg, G. A Protein Sequenator. Eur. J. Biochem. 1, 80–91 (1967).

23. Tullman, J., Marino, J. P. & Kelman, Z. Leveraging nature’s biomolecular designs in next-generation protein sequencing reagent development. Appl. Microbiol. Biotechnol. 104, 7261–7271 (2020).

24. Mitra, R. D., Shendure, J., Olejnik, J., Edyta-Krzymanska-Olejnik & Church, G. M. Fluorescent in situ sequencing on polymerase colonies. Anal. Biochem. 320, 55–65 (2003).

25. Tullman, J., Callahan, N., Ellington, B., Kelman, Z. & Marino, J. P. Engineering ClpS for selective and enhanced N-terminal amino acid binding. Appl. Microbiol. Biotechnol. 103, 2621–2633 (2019).

26. Tullman, J., Christensen, M., Kelman, Z. & Marino, J. P. A ClpS-based N-terminal amino acid binding reagent with improved thermostability and selectivity. Biochem. Eng. J. 154, 107438 (2020).

27. Callahan, N., Tullman, J., Kelman, Z. & Marino, J. Strategies for Development of a Next-Generation Protein Sequencing Platform. Trends Biochem. Sci. 45, 76–89 (01/2020).

28. Havranek, J. J. & Borgo, B. Molecules and methods for iterative polypeptide analysis and processing. US Patent (2014).

29. Callahan, N., Siegall, W. B., Bergonzo, C., Marino, J. P. & Kelman, Z. Contributions from ClpS surface residues in modulating N-terminal peptide binding and their implications for NAAB development. Protein Eng. Des. Sel. 36, gzad007 (2023).

30. Ikonomova, S. P. et al. Engineering GID4 for use as an N-terminal proline binder via directed evolution. Biotechnol. Bioeng. 122, 179–188 (2025).

31. Zheng, L., Sun, Y., Eisenstein, M. & Soh, H. T. Peptide sequencing via reverse translation of peptides into DNA. bioRxiv (2024) doi:10.1101/2024.05.31.596913.

32. Hacker, S. M. et al. Global profiling of lysine reactivity and ligandability in the human proteome. Nat. Chem. 9, 1181–1190 (2017).

33. Weerapana, E. et al. Quantitative reactivity profiling predicts functional cysteines in proteomes. Nature 468, 790–795 (2010).

34. Hahm, H. S. et al. Global targeting of functional tyrosines using sulfur-triazole exchange chemistry. Nat. Chem. Biol. 16, 150–159 (2020).

35. Jia, S., He, D. & Chang, C. J. Bioinspired thiophosphorodichloridate reagents for chemoselective histidine bioconjugation. J. Am. Chem. Soc. 141, 7294–7301 (2019).

36. Ma, N. et al. 2H-azirine-based reagents for chemoselective bioconjugation at carboxyl residues inside live cells. J. Am. Chem. Soc. 142, 6051–6059 (2020).

37. Zanon, P. R. A. et al. Profiling the proteome-wide selectivity of diverse electrophiles. Nat. Chem. 17, 1712–1721 (2025).

38. Xie, X. & Lin, S. Targeting and manipulating tryptophan interactions on proteins. ACS Chem. Biol. 19, 1211–1213 (2024).

39. Taylor, M. T., Nelson, J. E., Suero, M. G. & Gaunt, M. J. A protein functionalization platform based on selective reactions at methionine residues. Nature 562, 563–568 (2018).

40. Laursen, R. A. Solid-phase Edman degradation. An automatic peptide sequencer. Eur. J. Biochem. 20, 89–102 (1971).

41. Yu, C.-C. J. et al. Expansion microscopy of C. elegans. Elife 9, (2020).

42. Shin, T. W. et al. Dense, continuous membrane labeling and expansion microscopy visualization of ultrastructure in tissues. Nat. Commun. 16, 1579 (2025).

43. Muramoto, K., Nokihara, K., Ueda, A. & Kamiya, H. Gas-phase microsequencing of peptides and proteins with a fluorescent Edman-type reagent, fluorescein isothiocyanate. Biosci. Biotechnol. Biochem. 58, 300–304 (1994).

44. Deol, H. et al. After 75 years, an alternative to Edman degradation: A mechanistic and efficiency study of a base-induced method for N-terminal peptide sequencing. J. Am. Chem. Soc. 147, 13973–13982 (2025).

45. Chen, F. et al. Nanoscale Imaging of RNA with Expansion Microscopy. Nat. Methods 13, 679–684 (2016-8).

46. Fry, B., Slaw, K. & Polizzi, N. F. Zero-shot design of drug-binding proteins via neural selection-expansion. bioRxiv 2025.04.22.649862 (2025) doi:10.1101/2025.04.22.649862.

47. Vantourout, J. C. et al. Serine-selective bioconjugation. J. Am. Chem. Soc. 142, 17236–17242 (2020).

48. Gao, R. et al. A highly homogeneous polymer composed of tetrahedron-like monomers for high-isotropy expansion microscopy. Nat. Nanotechnol. 16, 698–707 (2021).

49. Lee, H., Yu, C.-C., Boyden, E. S., Zhuang, X. & Kosuri, P. Tetra-gel enables superior accuracy in combined super-resolution imaging and expansion microscopy. Sci. Rep. 11, 16944 (2021).

50. Niall, H. D. Automated Edman degradation: the protein sequenator. Methods Enzymol. 27, 942–1010 (1973).

51. Otaka, A., Koide, T., Shide, A. & Fujii, N. Application of dimethylsulphoxide(DMSO) / trifluoroacetic acid(TFA) oxidation to the synthesis of cystine-containing peptide. Tetrahedron Lett. 32, 1223–1226 (1991).

52. Spetzler, J. C. & Meldal, M. Evaluation of strategies for ?one-pot? deprotection, cleavage and disulfide bond formation in the preparation of cystine-containing peptides. Lett. Pept. Sci. 3, 327–332 (1997).

53. Estandian, D. M. Enabling tools for de-novo single molecule protein sequencing. (Massachusetts Institute of Technology, 2021).

54. Estandian, D. M., Choueiri, A. G., Boyden, E. S. & Wassie, A. Single-molecule protein and peptide sequencing. Preprint at https://patents.google.com/patent/US20230305017A1/en (2023).

55. Choueiri, A. G. Single-Molecule Protein Sequencing (I) and Genetically Dominant mRNA Therapies to Combat Viral Evolution (II). (Massachusetts Institute of Technology, 2022).

56. Farnsworth, V. & Steinberg, K. The Generation of Phenylthiocarbamyl or Anilinothiazolinone Amino Acids from the Postcleavage Products of the Edman Degradation. Anal. Biochem. 215, 200–210 (12/1993).

57. Braun, M. B. et al. Peptides in headlock – a novel high-affinity and versatile peptide-binding nanobody for proteomics and microscopy. Sci. Rep. 6, 19211 (2016).

58. Hodges, R. S., Heaton, R. J., Parker, J. M., Molday, L. & Molday, R. S. Antigen-antibody interaction. Synthetic peptides define linear antigenic determinants recognized by monoclonal antibodies directed to the cytoplasmic carboxyl terminus of rhodopsin. J. Biol. Chem. 263, 11768–11775 (1988).

59. Pinilla, C., Appel, J. R. & Houghten, R. A. Functional importance of amino acid residues making up peptide antigenic determinants. Mol. Immunol. 30, 577–585 (1993).

60. Rodgers, J. R. & Rich, R. R. 6 - Antigens and antigen presentation. in Clinical Immunology (Fourth Edition) (eds. Rich, R. R. et al.) 77–89 (Elsevier, London, 2013).

61. Stark, S. E. & Caton, A. J. Antibodies that are specific for a single amino acid interchange in a protein epitope use structurally distinct variable regions. J. Exp. Med. 174, 613–624 (1991).

62. Stark, Y. et al. Modular binder technology by NGS-aided, high-resolution selection in yeast of designed armadillo modules. Proc. Natl. Acad. Sci. U. S. A. 121, e2318198121 (2024).

63. Buus, S. et al. High-resolution Mapping of Linear Antibody Epitopes Using Ultrahigh-density Peptide Microarrays. Mol. Cell. Proteomics 11, 1790–1800 (2012).

64. Khakzad, H. et al. A new age in protein design empowered by deep learning. Cell Syst. 14, 925–939 (2023).

65. Cao, L. et al. Design of protein-binding proteins from the target structure alone. Nature 605, 551–560 (2022).

66. Jumper, J. et al. Highly accurate protein structure prediction with AlphaFold. Nature 596, 583–589 (2021).

67. Asano, S. M. et al. Expansion Microscopy: Protocols for Imaging Proteins and RNA in Cells and Tissues. Curr. Protoc. Cell Biol. 80, e56 (2018-9).

