## Supplementary material for "Principles of *in situ* protein sequencing: expansion microscopy-adapted Edman degradation and amino acid recognition": Source Data For Specific Figures

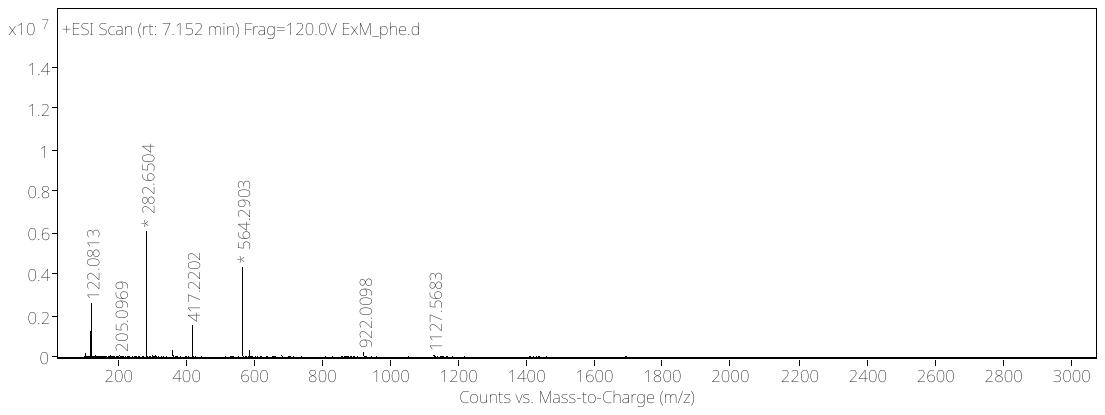

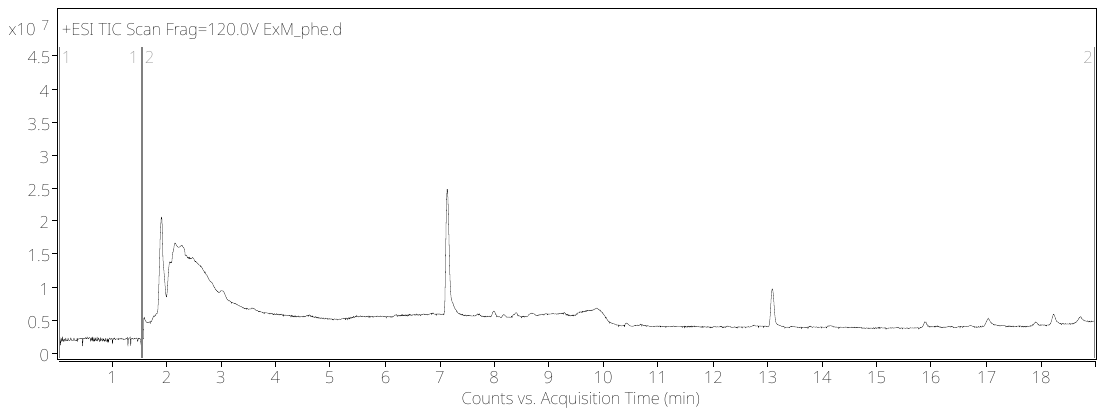

**Condition**

**Oxidation – phenylalanine results**

Mass spectrum of peak

Mass spectrum of peak

LC/Q-ToF Total ion chromatogram – C18 column, method 1

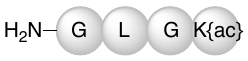

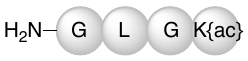

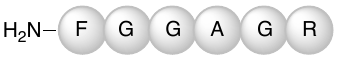

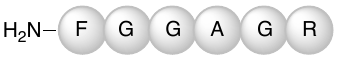

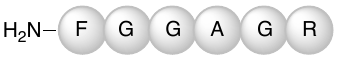

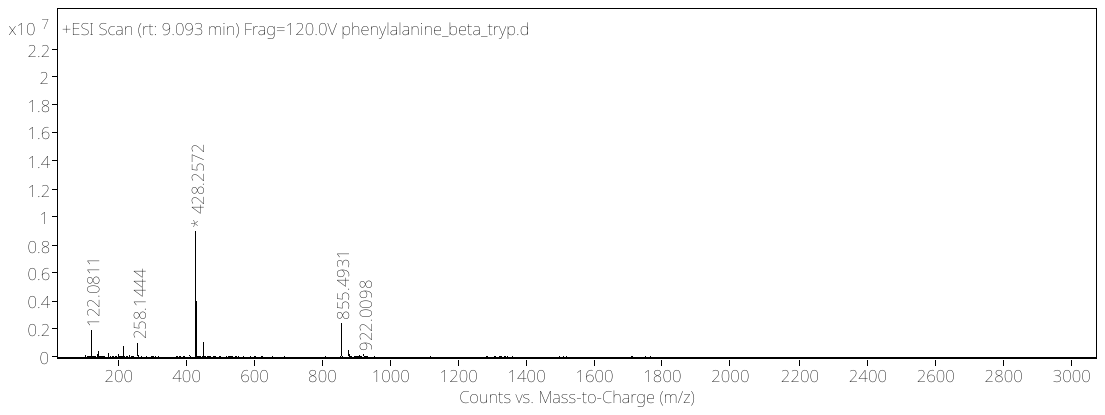

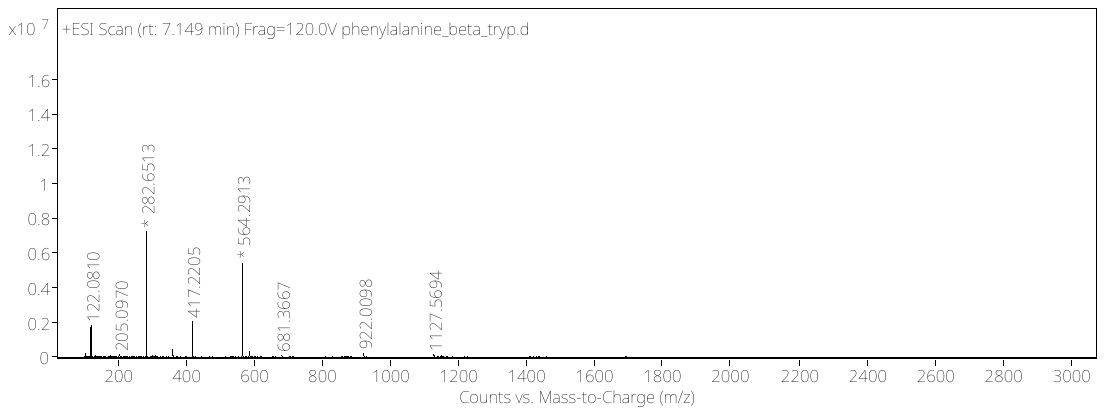

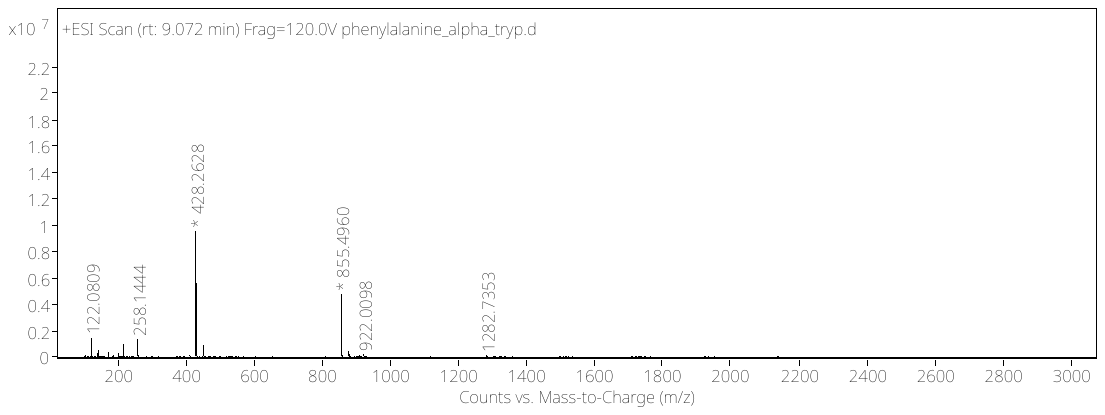

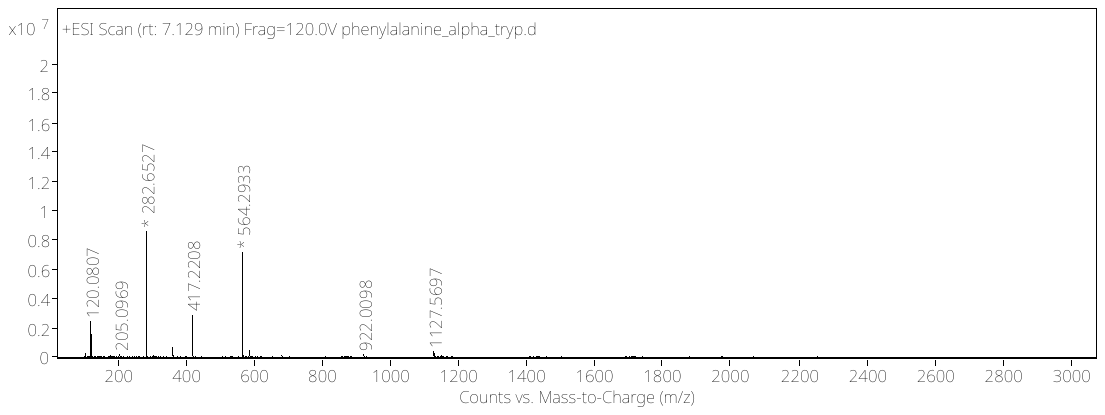

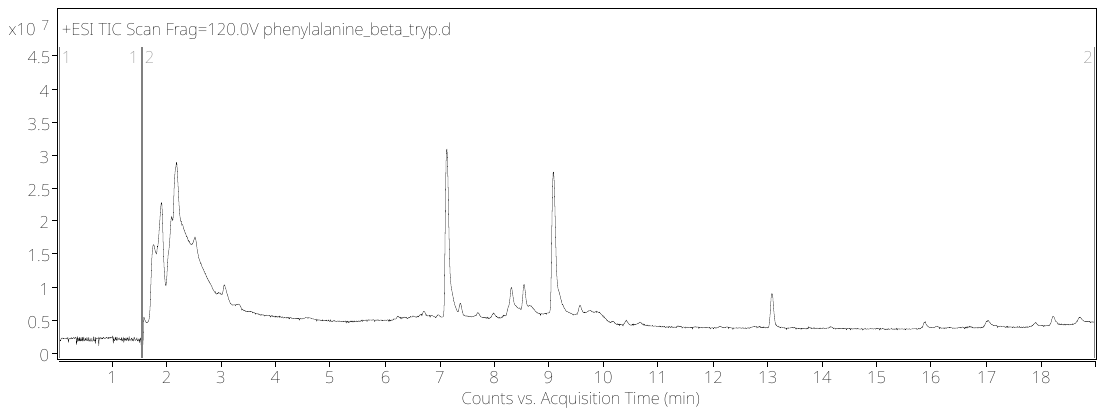

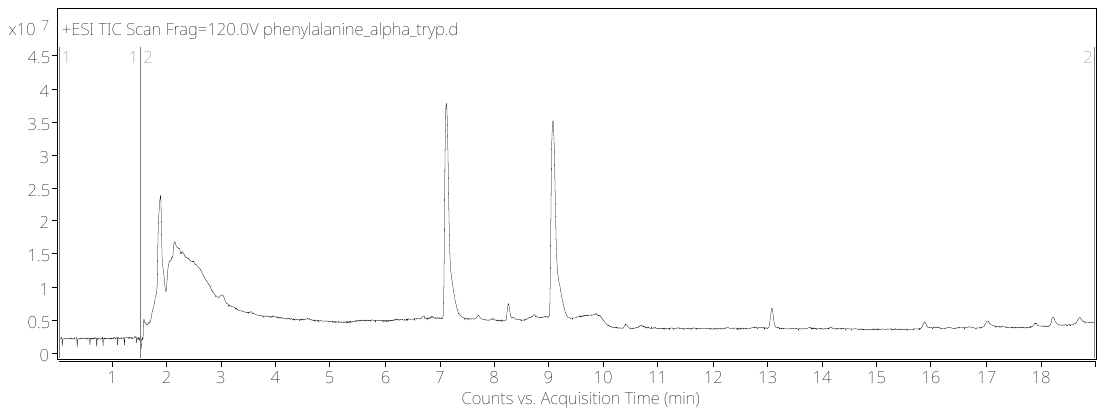

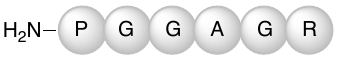

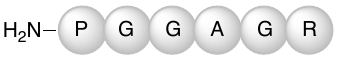

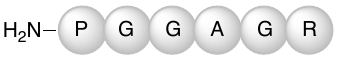

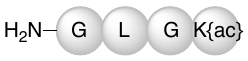

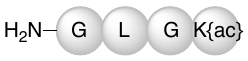

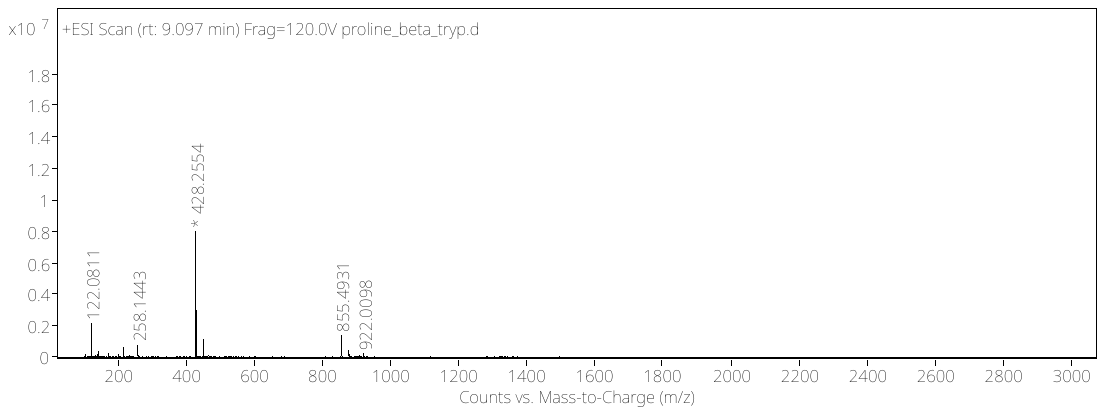

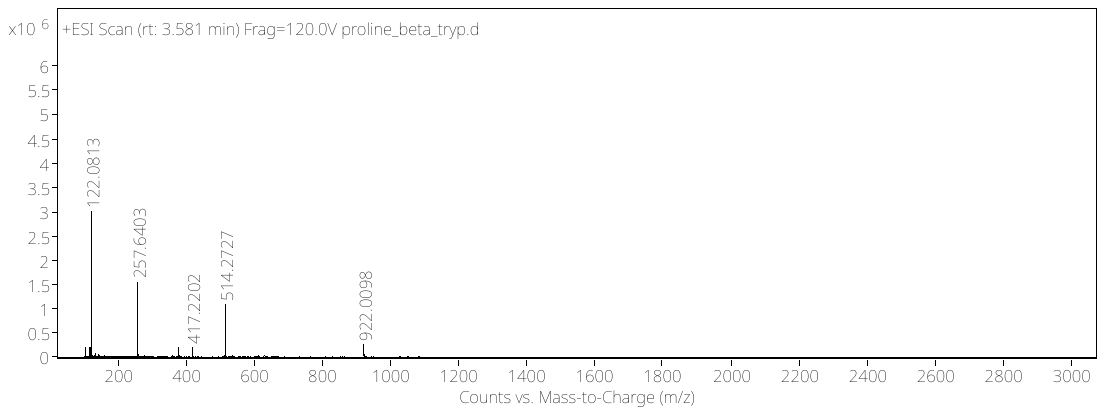

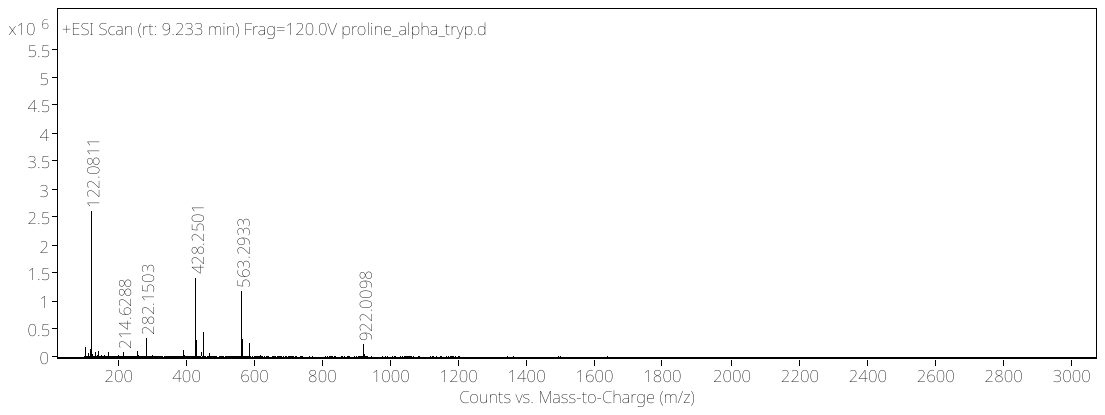

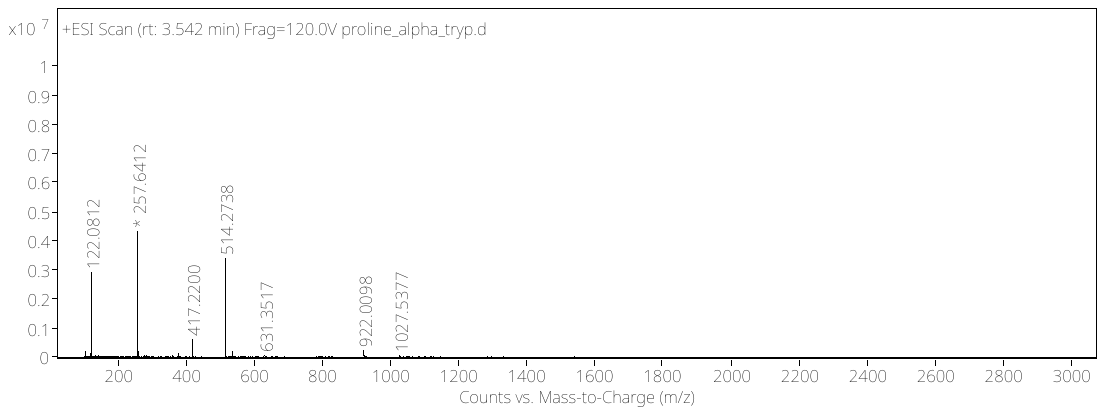

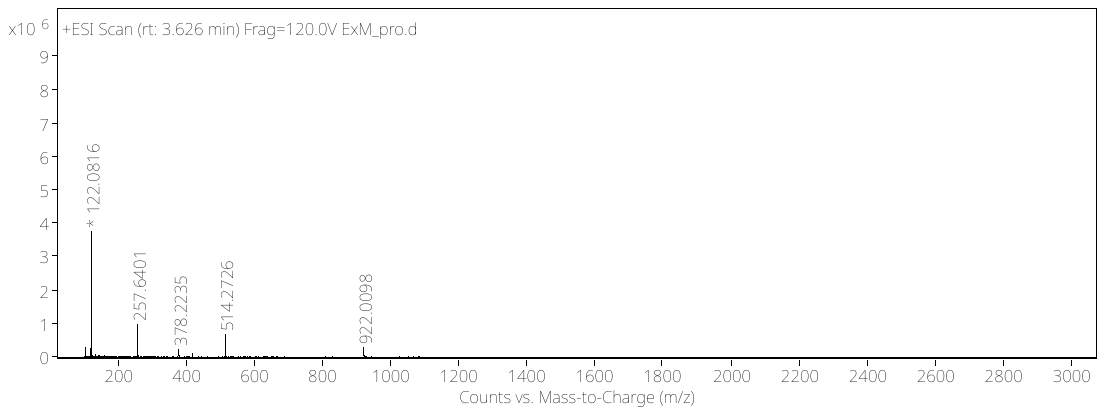

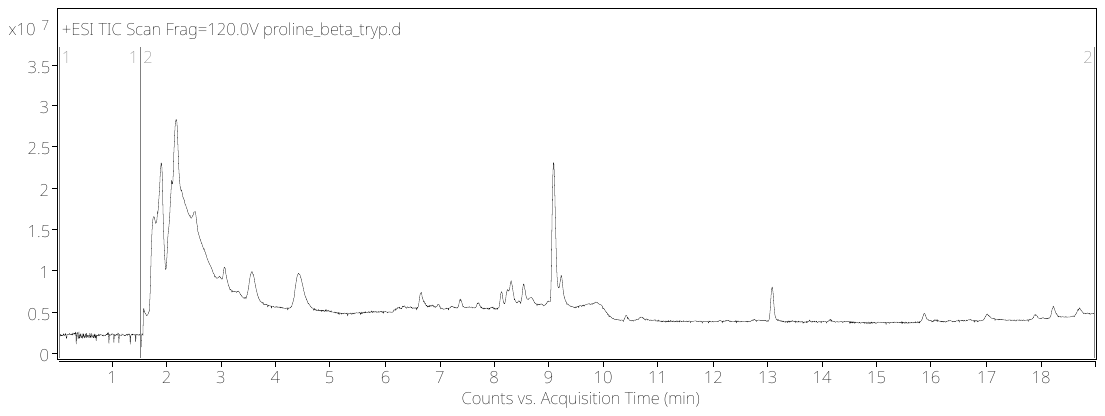

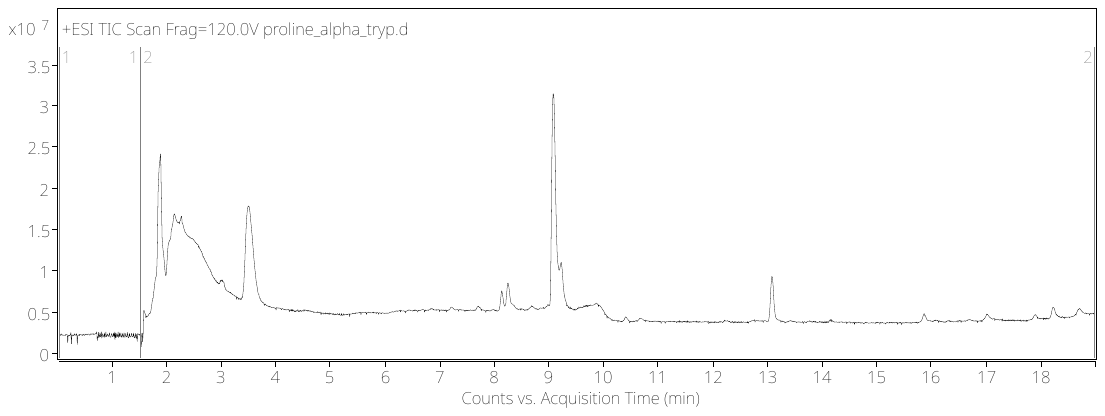

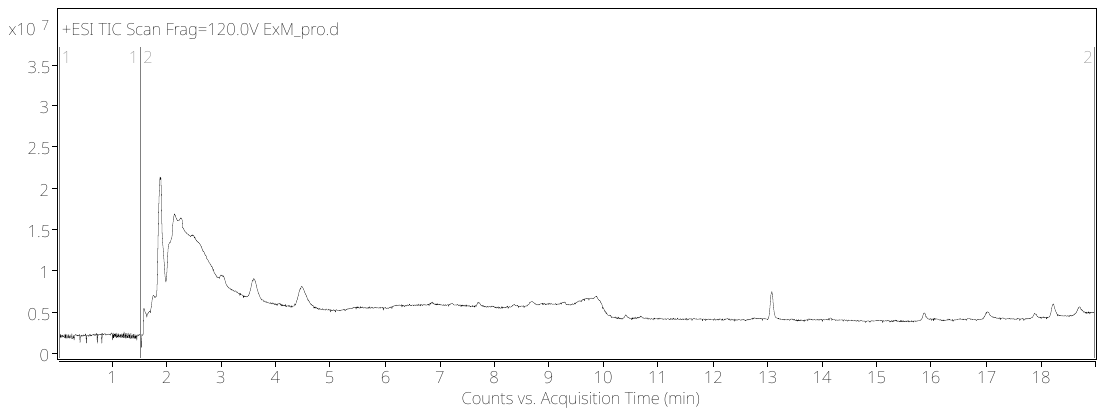

**Source Data - Supplementary Fig. 8d:**

The three conditions tested for oxidation with F_1_ peptide (FGGAGRGLGK{acrylic acid}): ExM (top row), in solution with trypsin (middle row), in solution with trypsin and APS, TEMED (bottom row). LC/Q-ToF total ion chromatogram for each condition (center) and arrows pointing towards trypsinized N-terminal fragment: phenylalanine peptide FGGAGR (exact mass: 563.2816), and trypsinized C-terminal fragment C-terminal fragment: GLGK{acylic acid} (exact mass: 427.2431). The circled mass on the mass spectrum matches the peptide overlayed on the graph.

In solution with trypsin

In solution with trypsin,

APS, TEMED

ExM

**Source Data - Supplementary Fig. 8g:**

The three conditions tested for oxidation with P_1_ peptide (PGGAGRGLGK{acrylic acid}): ExM (top row), in solution with trypsin (middle row), in solution with trypsin and APS, TEMED (bottom row). LC/Q-ToF total ion chromatogram for each condition (center) and arrows pointing towards mass spectrum of trypsinized N-terminal fragment: proline peptide PGGAGR (exact mass: 513.2659), and trypsinized C-terminal fragment: GLGK{acylic acid} (exact mass: 427.2431). The circled mass on the mass spectrum matches the peptide overlayed on the graph.

**Condition**

LC/Q-ToF Total ion chromatogram – C18 column, method 1

Mass spectrum of peak

Mass spectrum of peak

ExM

In solution with trypsin

In solution with trypsin,

APS, TEMED

**Oxidation – proline results**

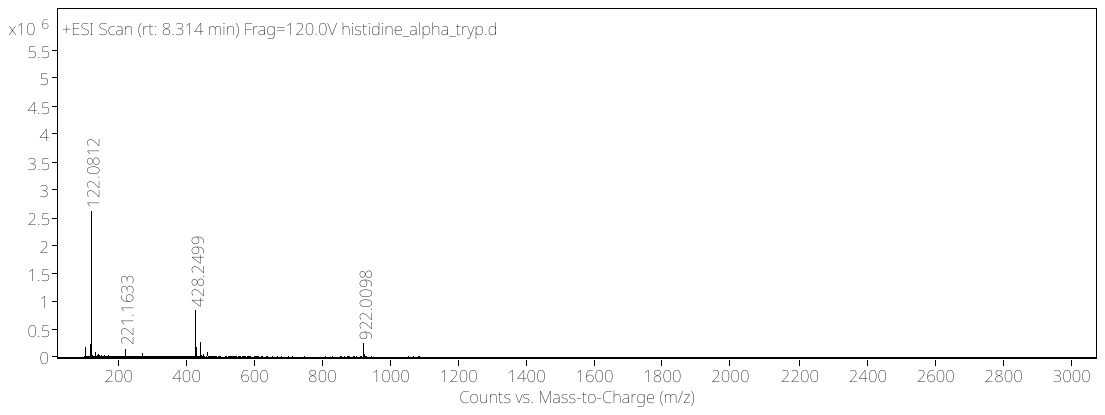

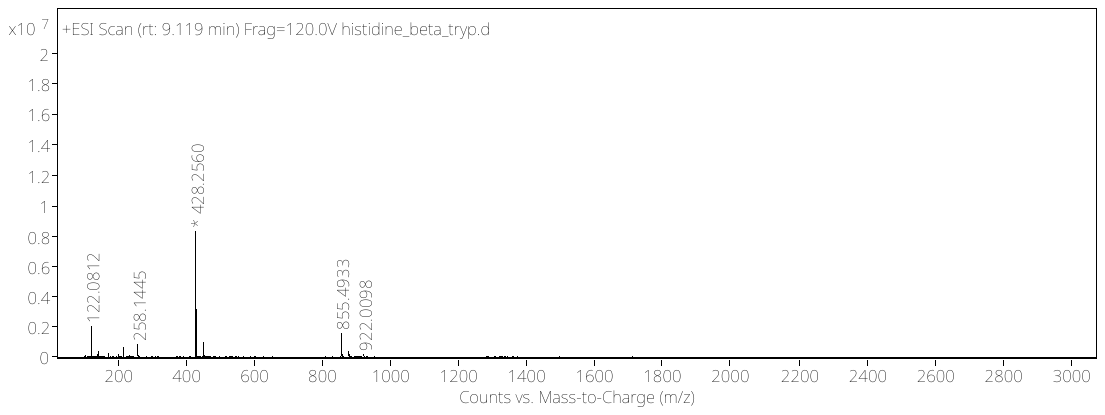

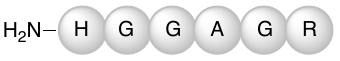

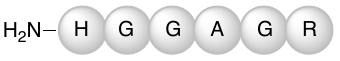

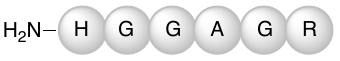

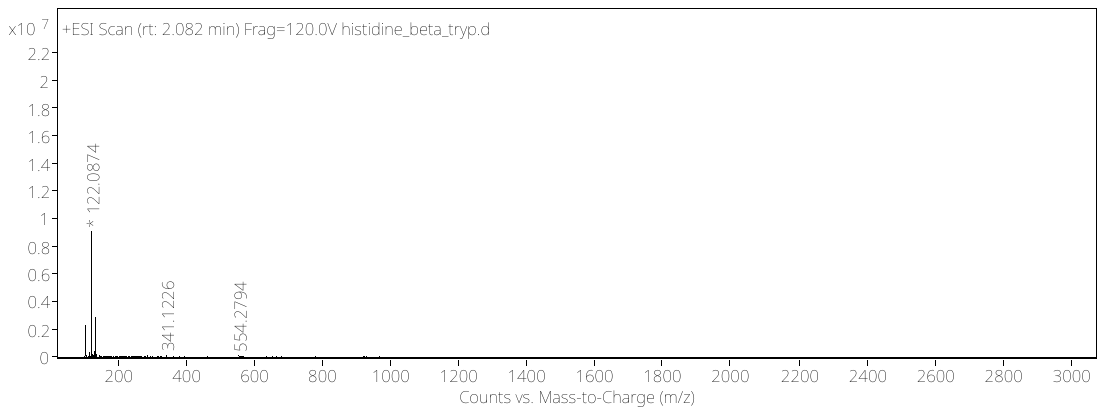

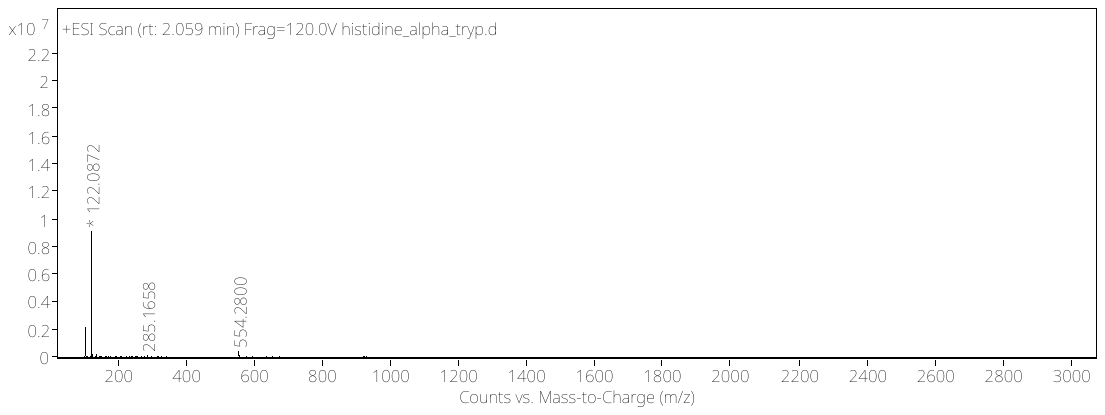

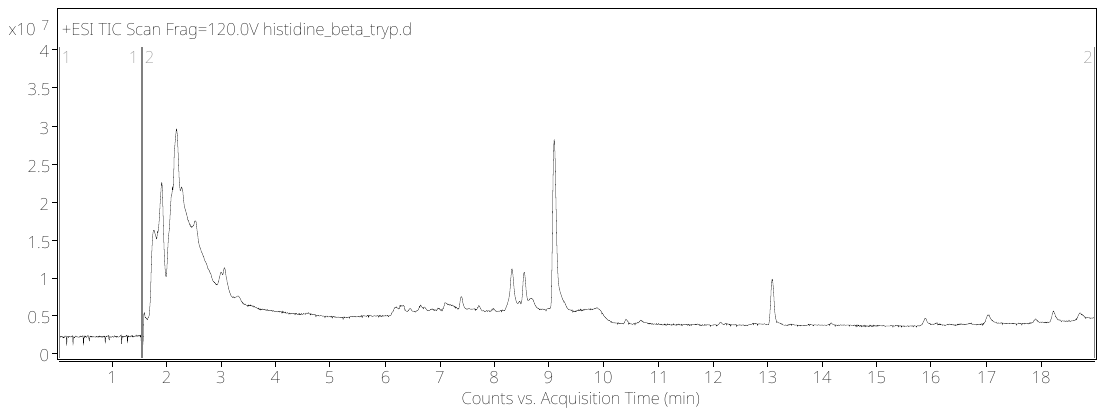

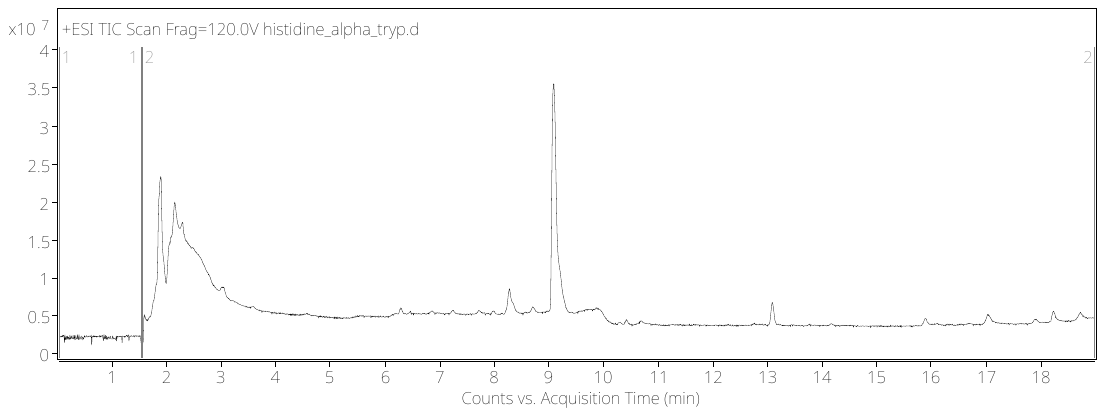

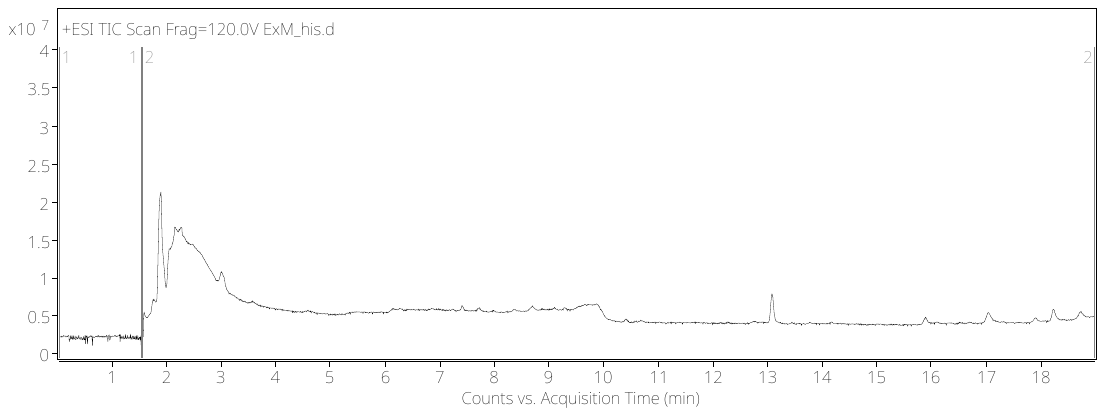

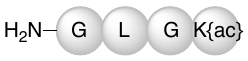

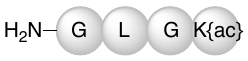

**Source Data - Supplementary Fig. 8f:**

The three conditions tested for oxidation with H_1_ peptide (HGGAGRGLGK{acrylic acid}): ExM (top row), in solution with trypsin (middle row), in solution with trypsin and APS, TEMED (bottom row). LC/Q-ToF total ion chromatogram for each condition (center) and arrows pointing towards mass spectrum of trypsinized N-terminal fragment: histidine peptide HGGAGR (exact mass: 553.2721), and trypsinized C-terminal fragment: GLGK{acylic acid} (exact mass: 427.2431). The circled mass on the mass spectrum matches the peptide overlayed on the graph.

Low abundance detected ~ 4.5 * 10^5^

In solution with trypsin,

APS, TEMED

In solution with trypsin

ExM

Mass spectrum of peak

Mass spectrum of peak

LC/Q-ToF Total ion chromatogram – C18 column, method 1

**Condition**

**Oxidation – histidine results**

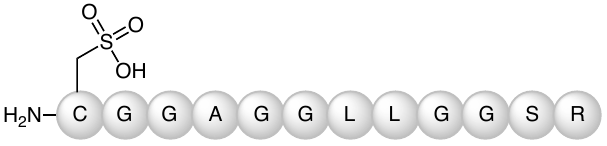

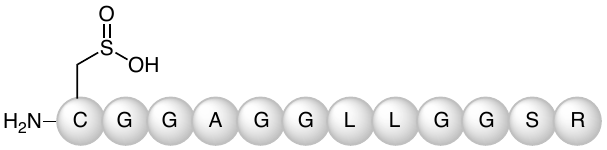

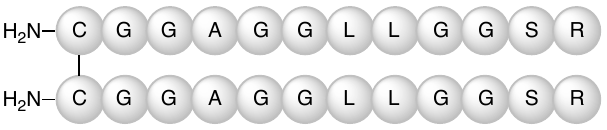

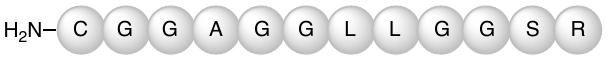

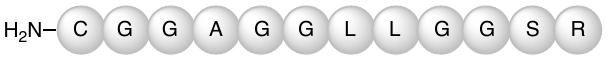

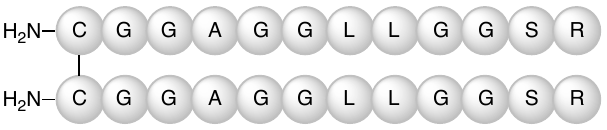

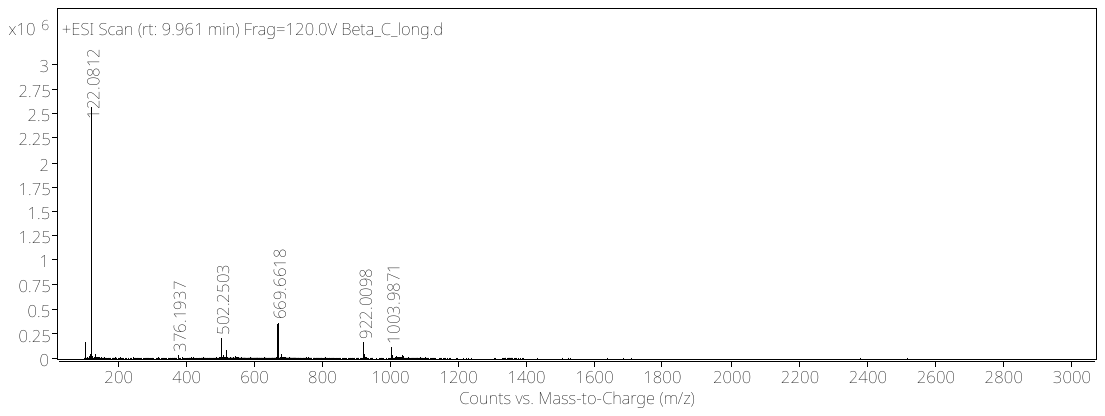

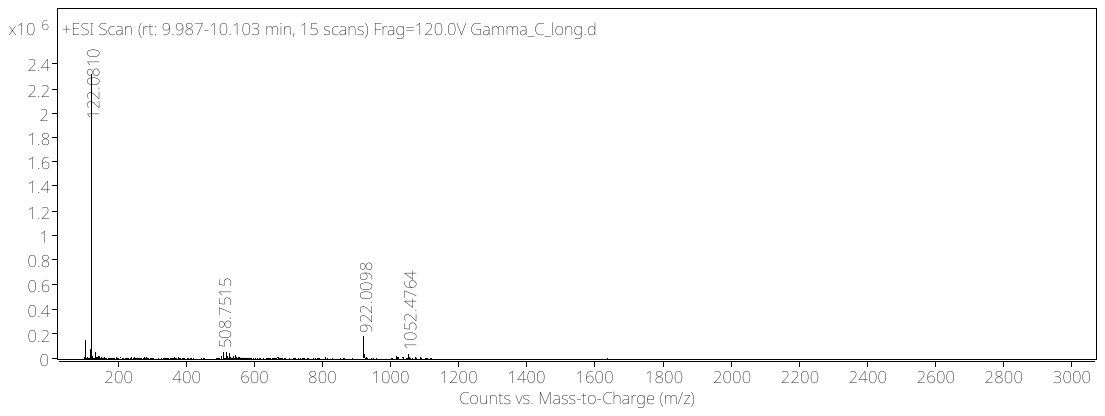

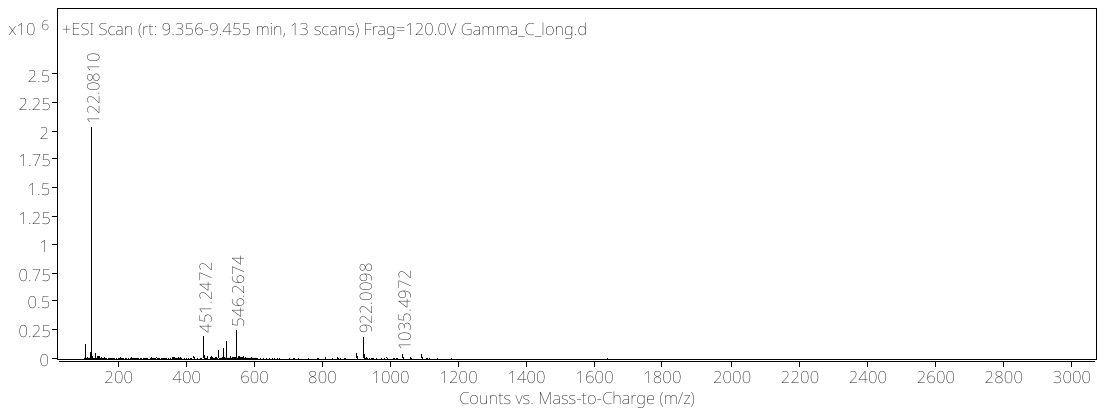

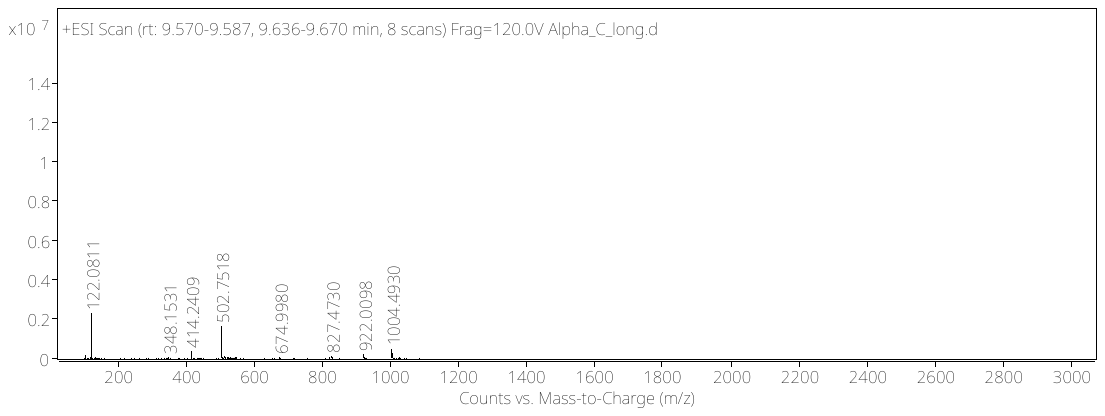

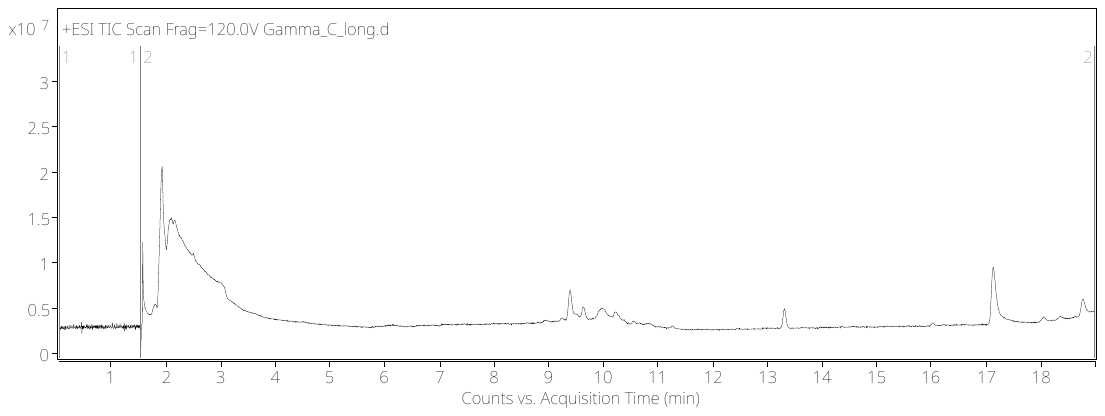

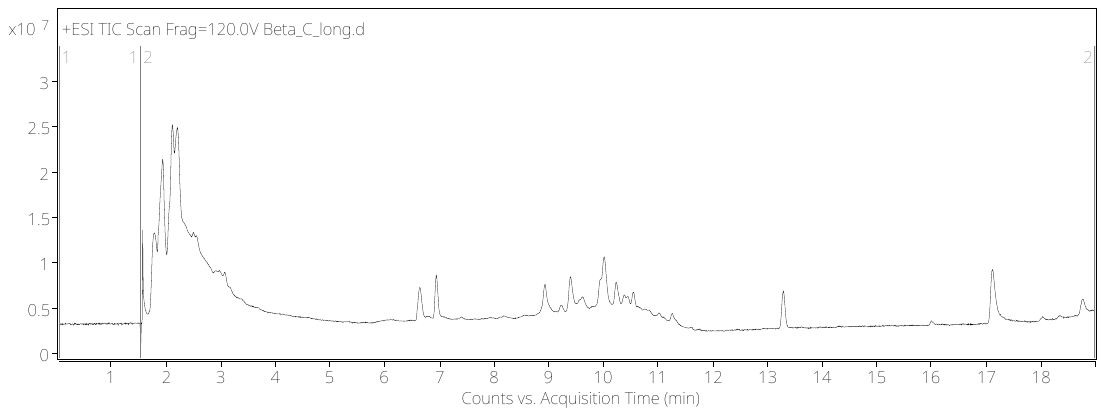

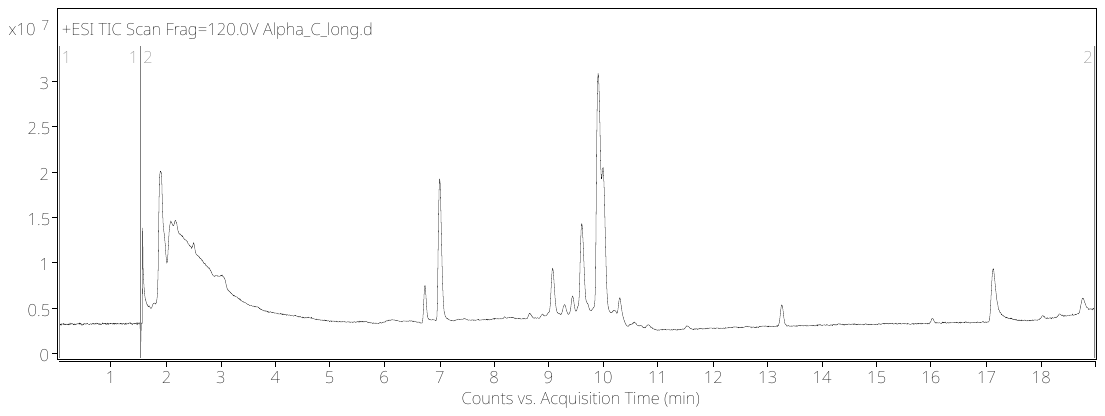

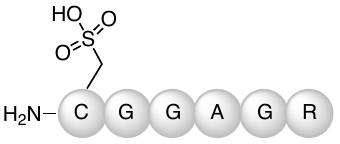

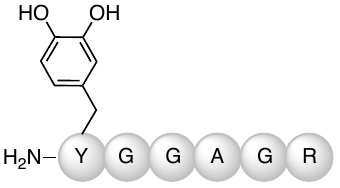

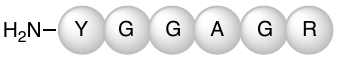

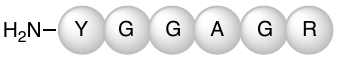

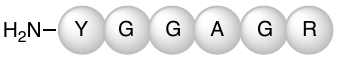

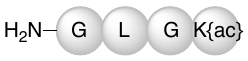

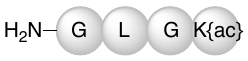

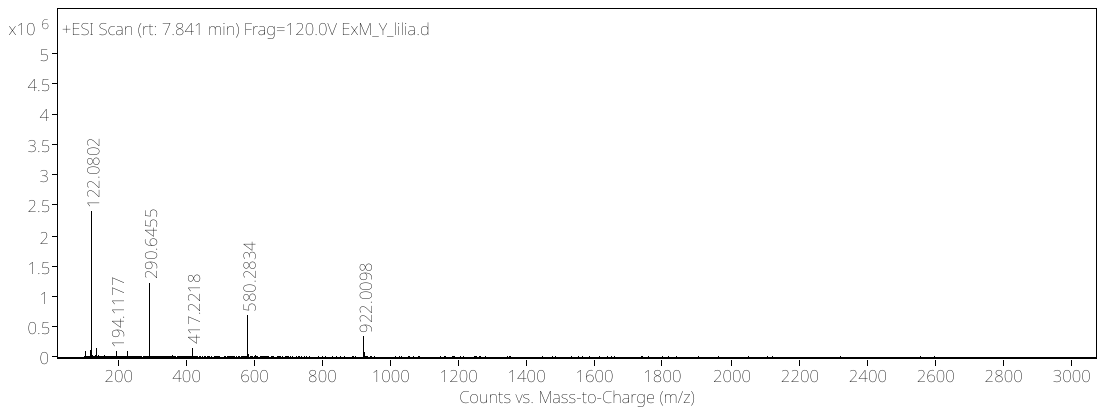

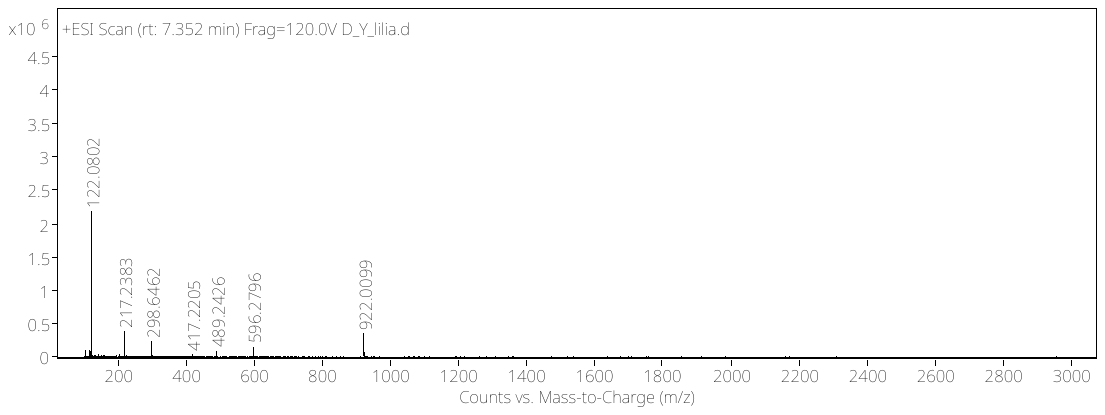

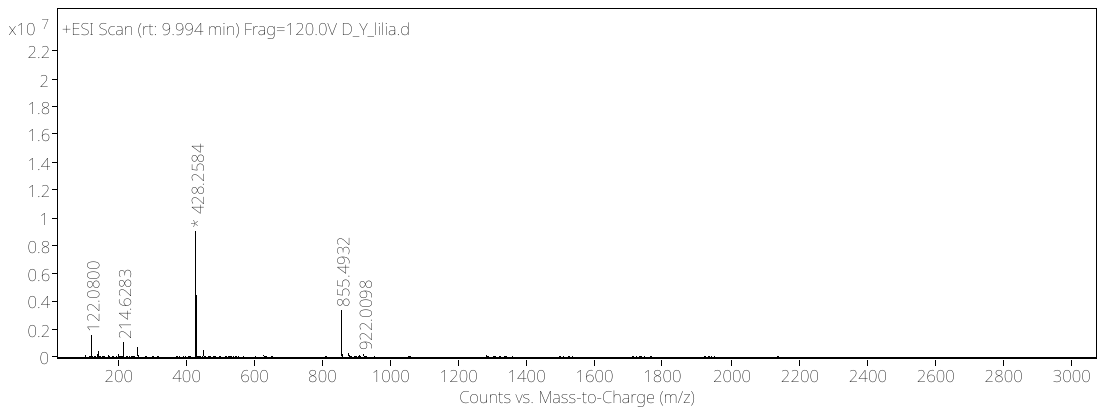

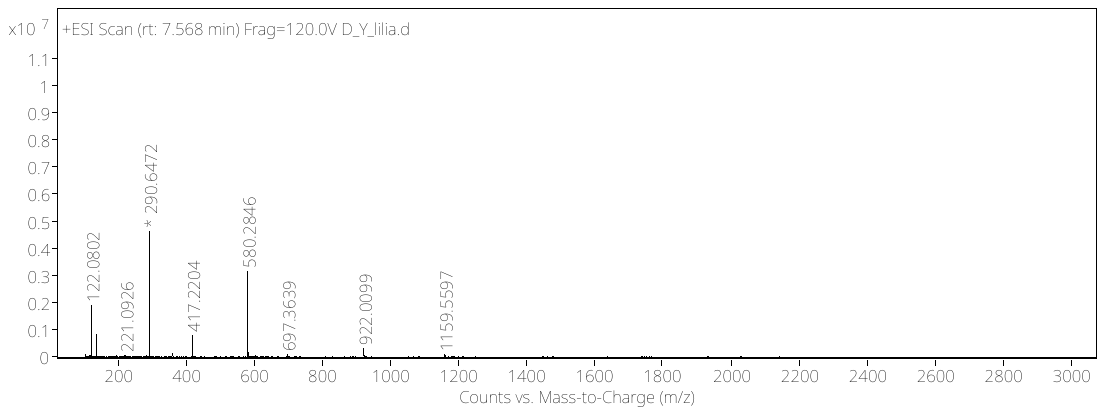

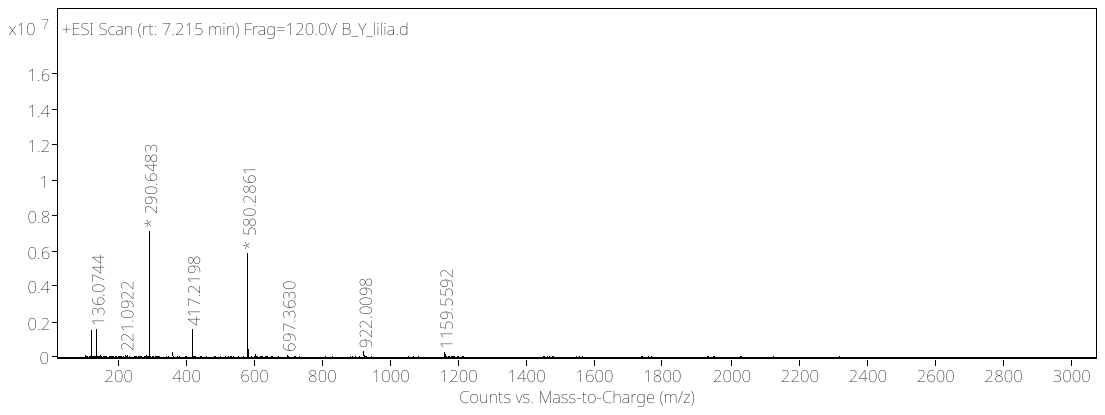

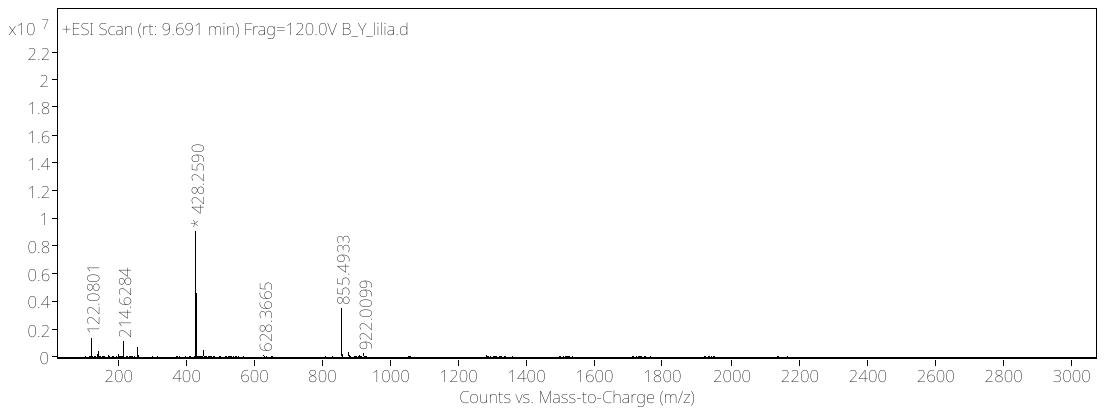

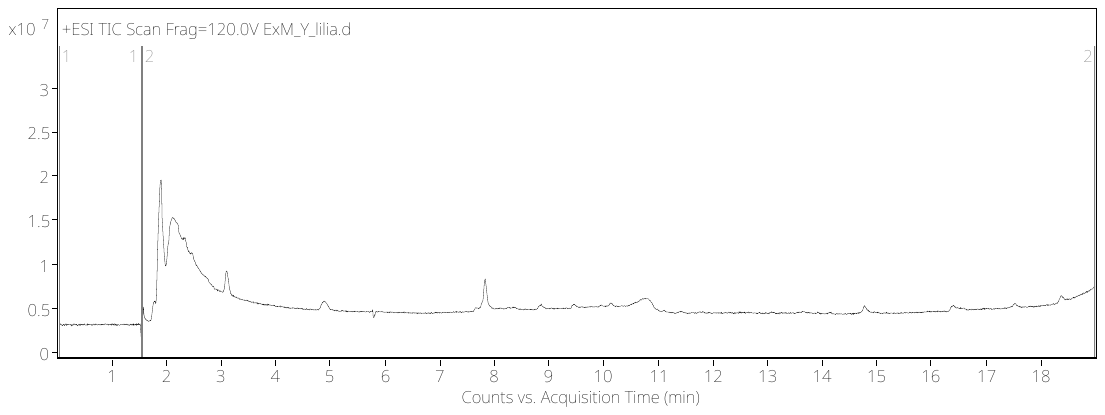

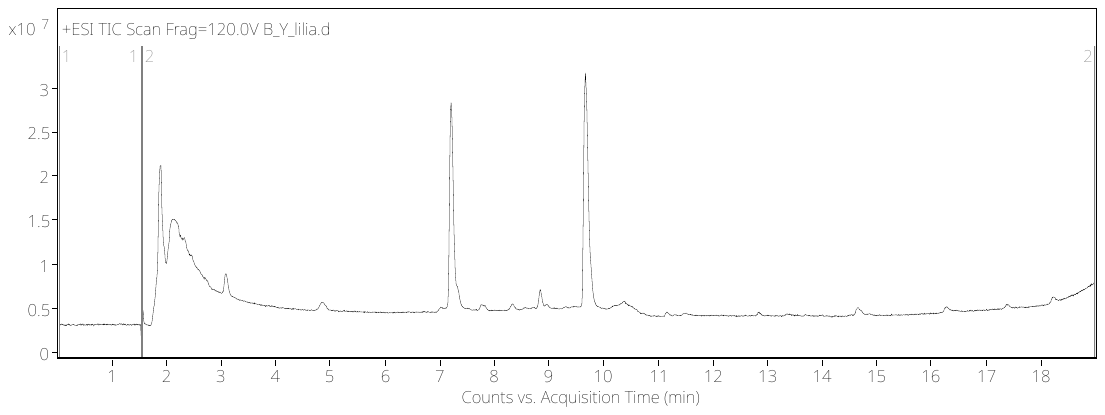

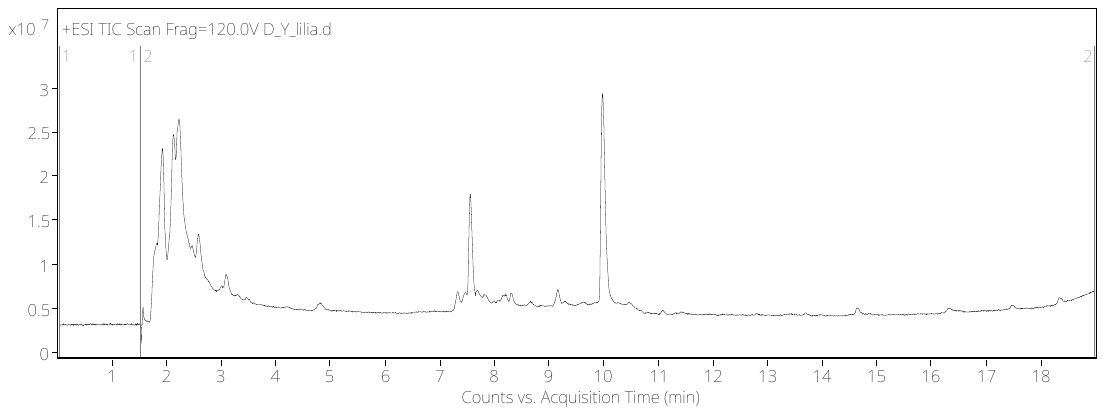

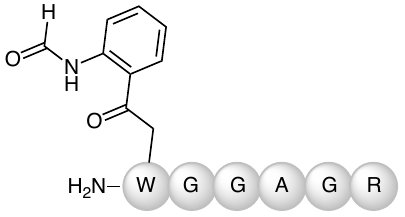

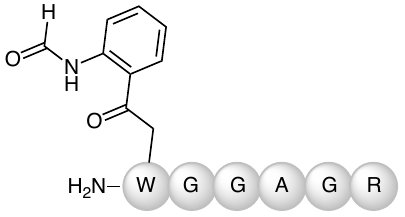

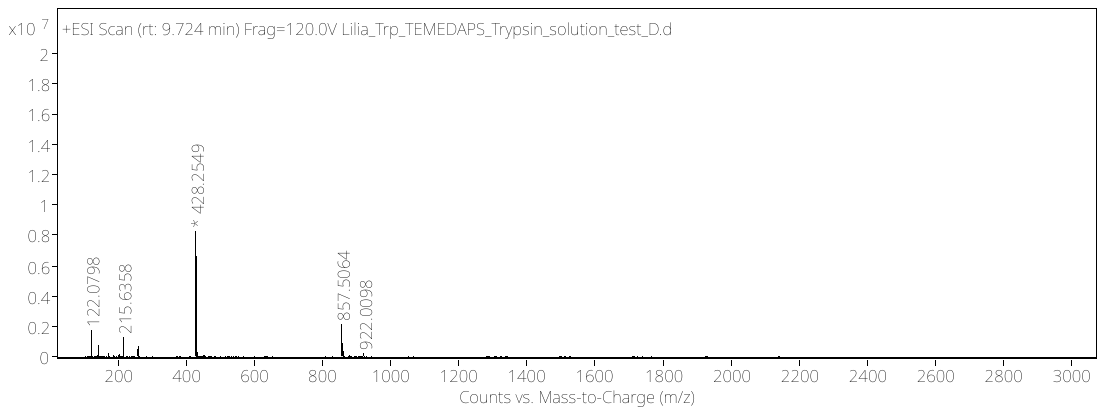

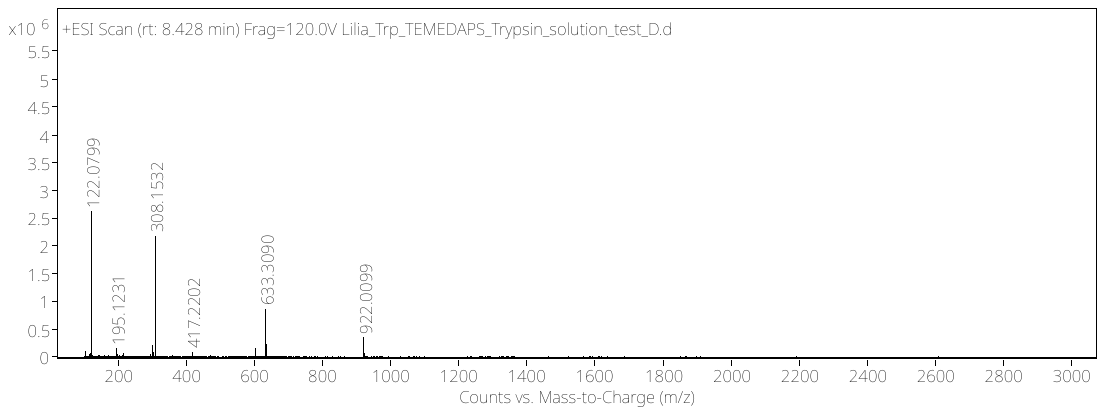

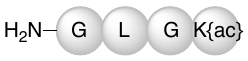

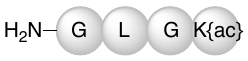

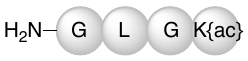

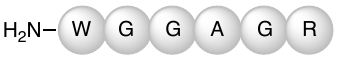

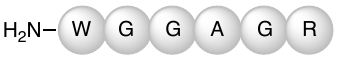

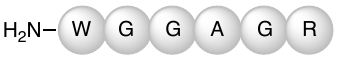

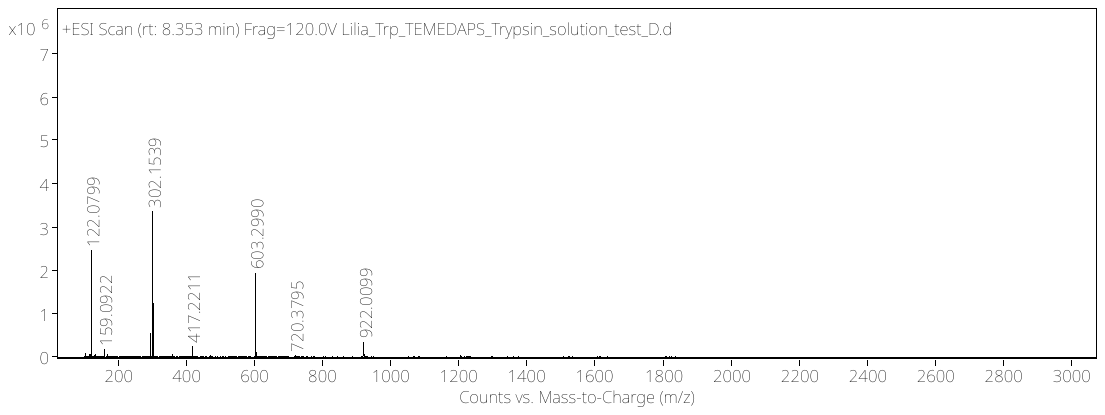

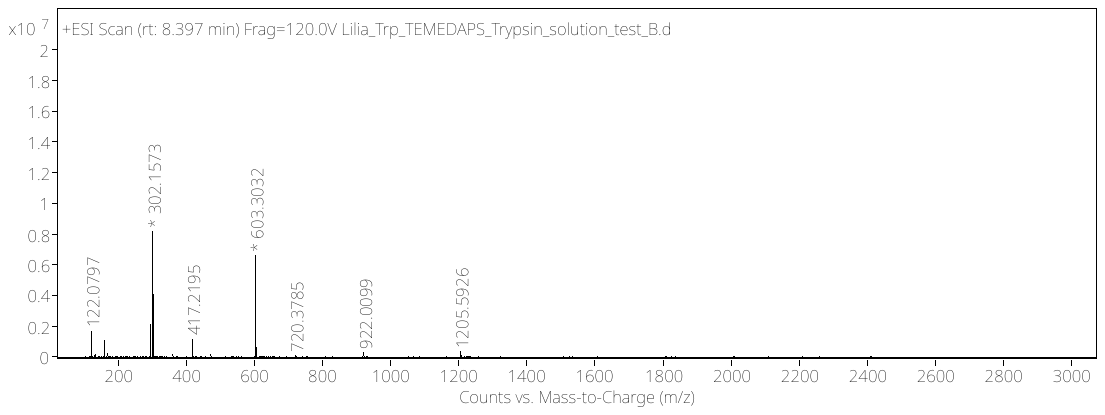

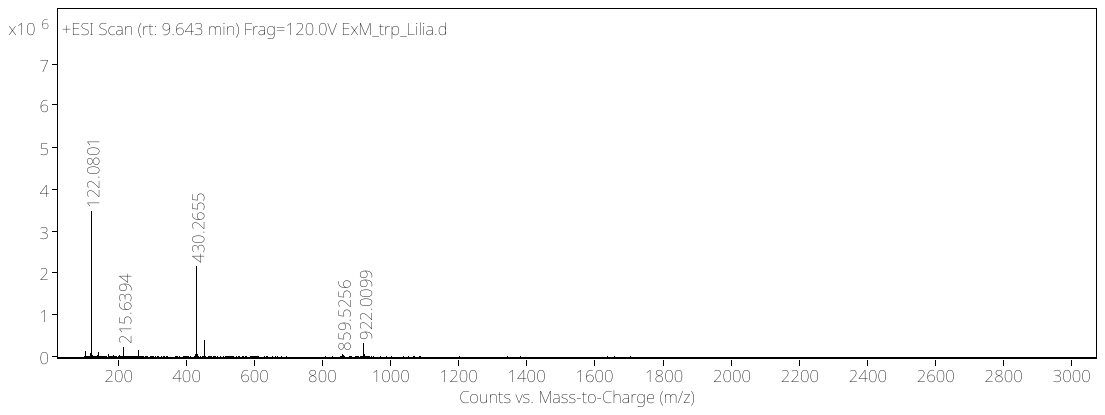

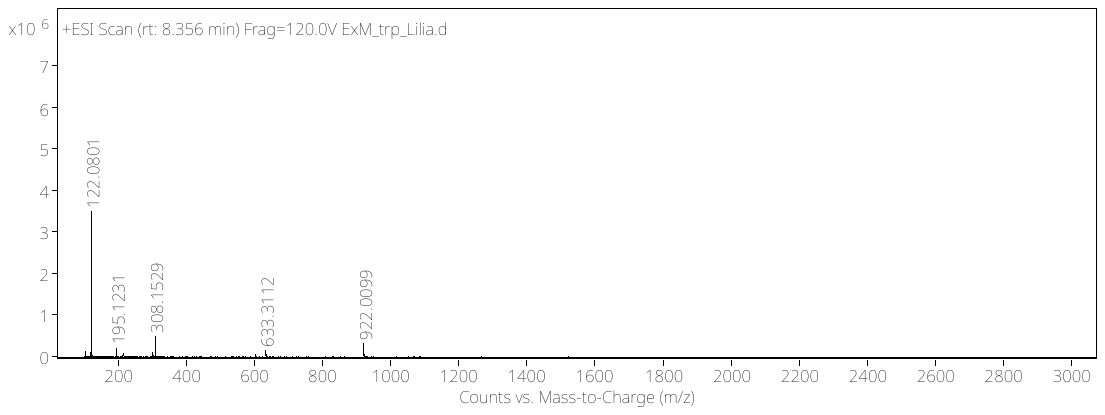

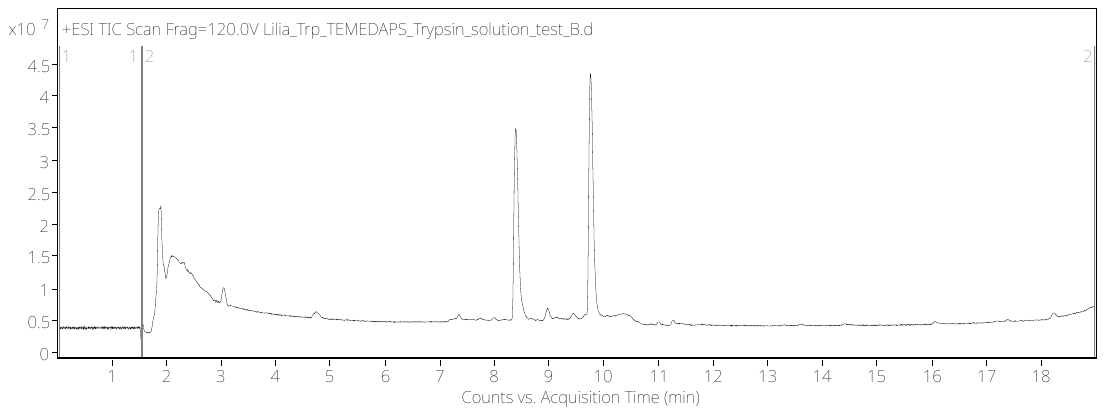

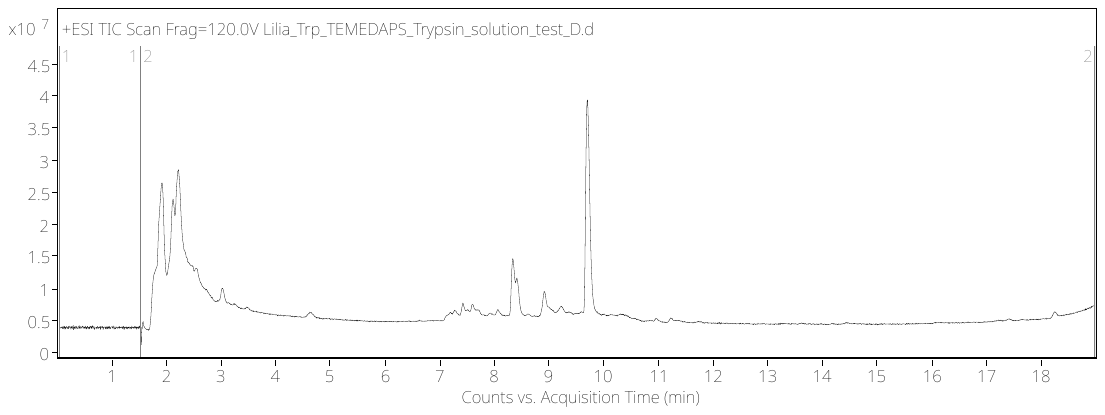

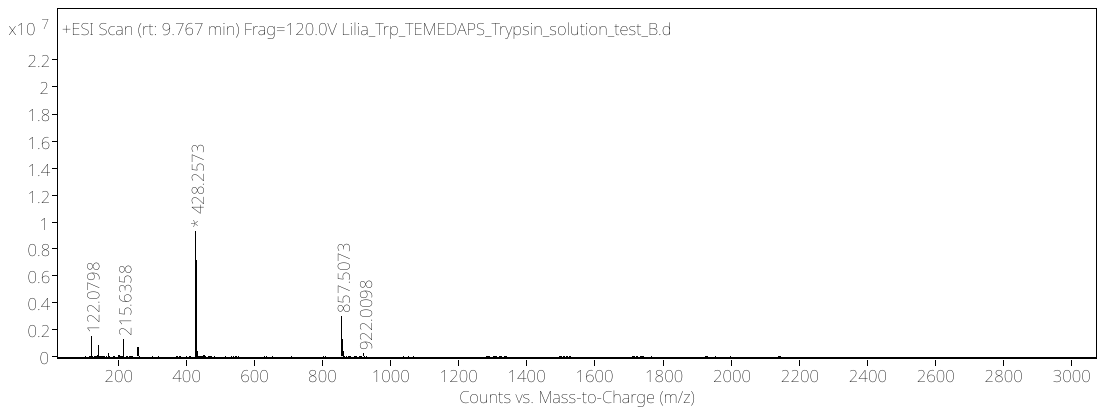

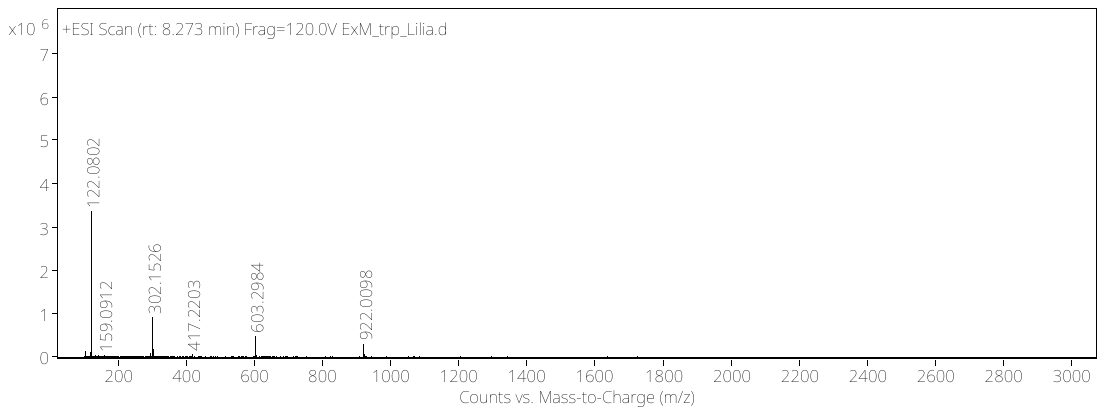

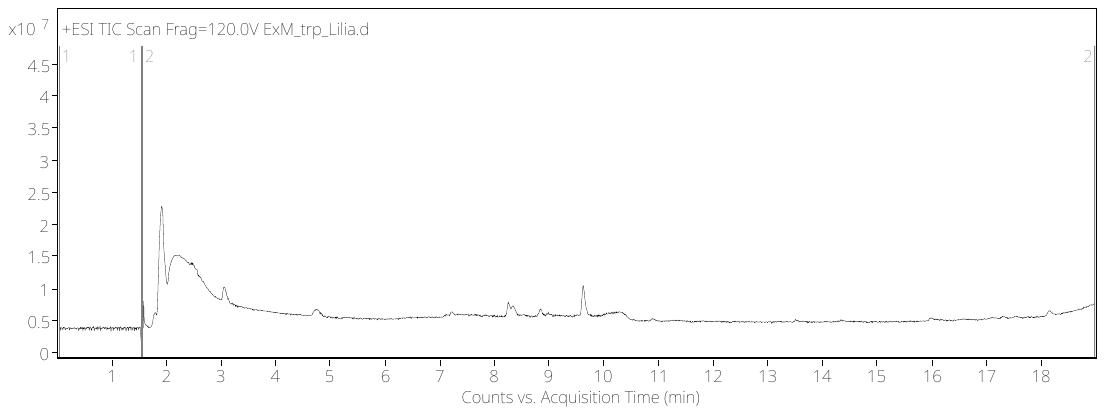

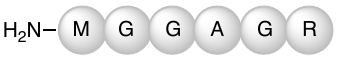

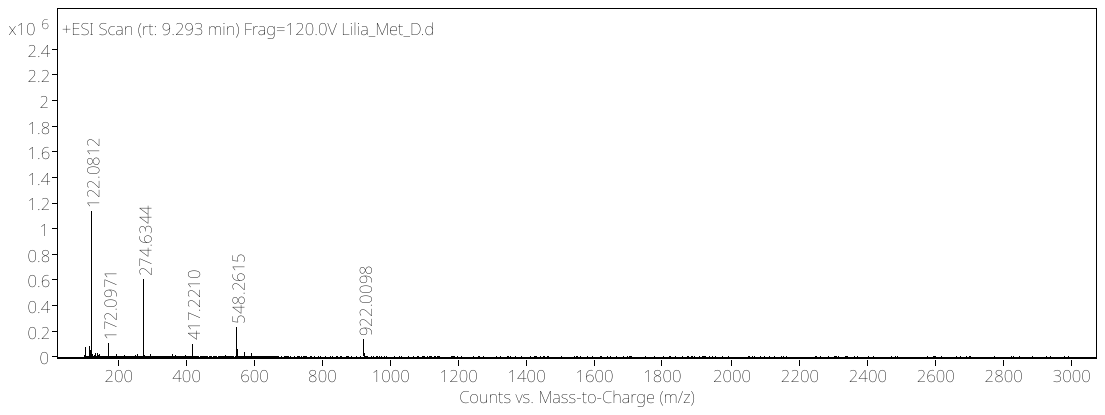

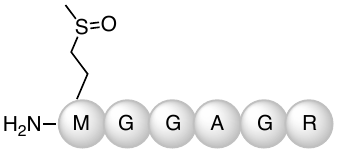

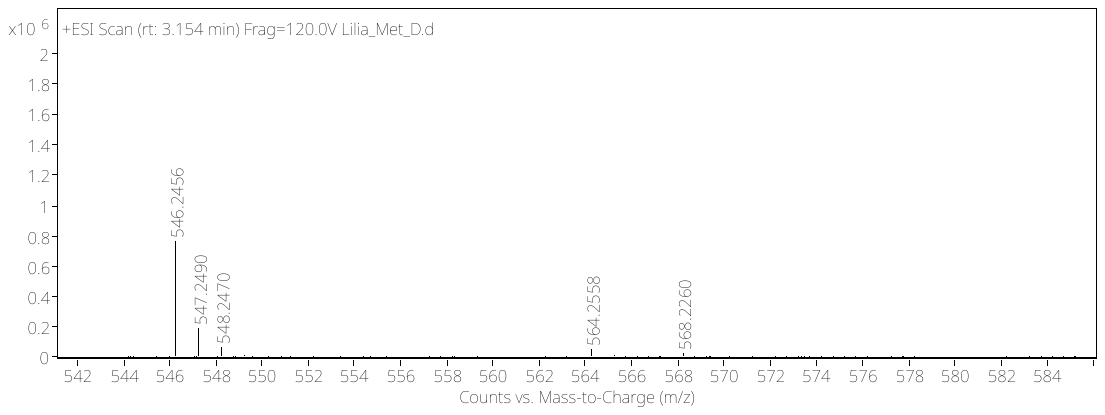

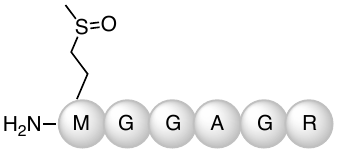

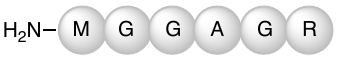

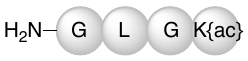

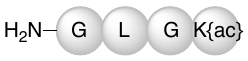

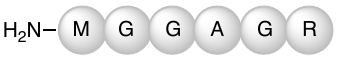

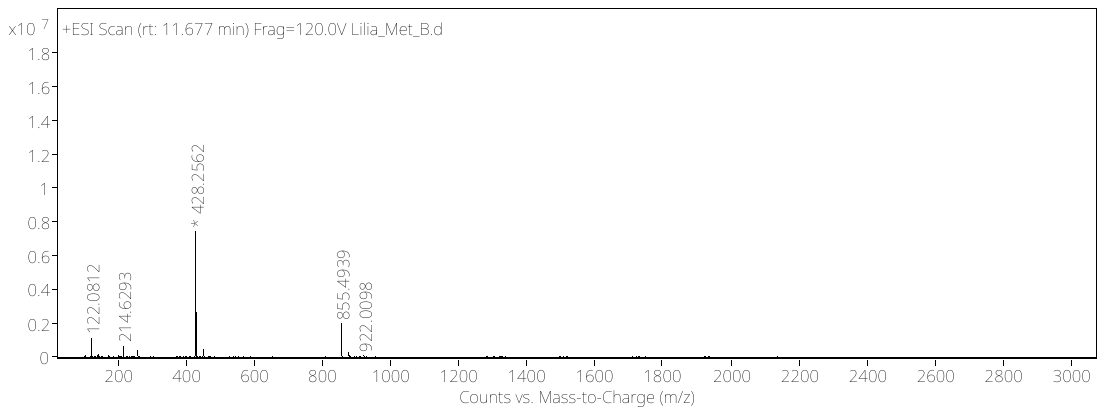

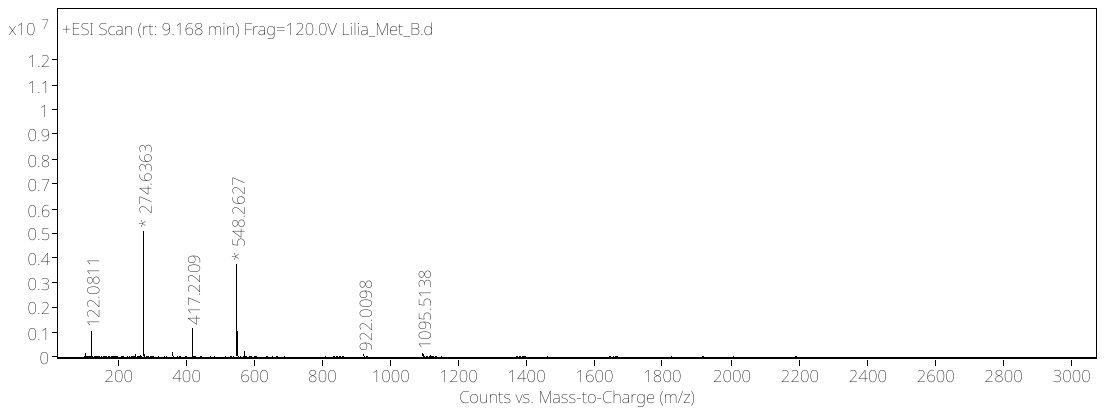

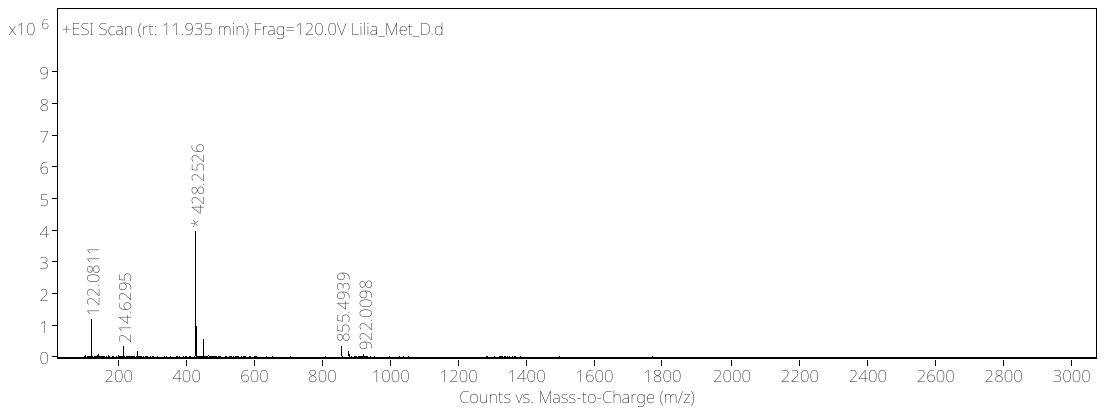

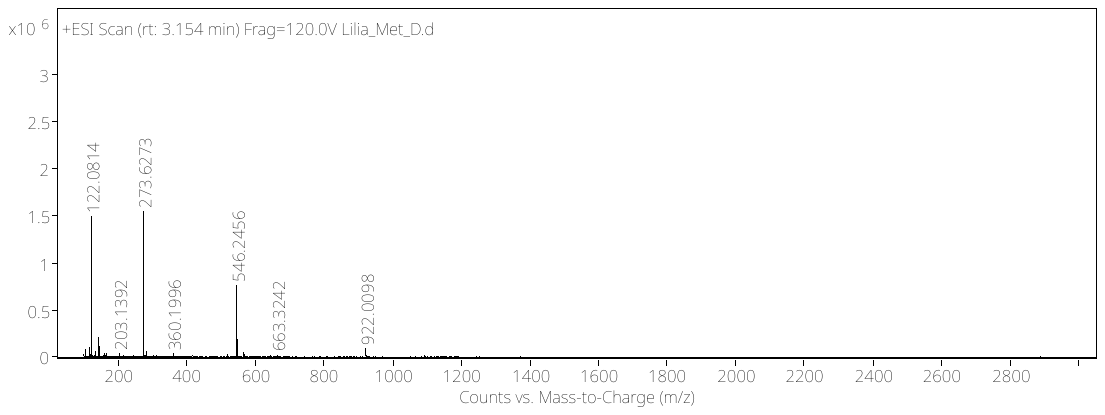

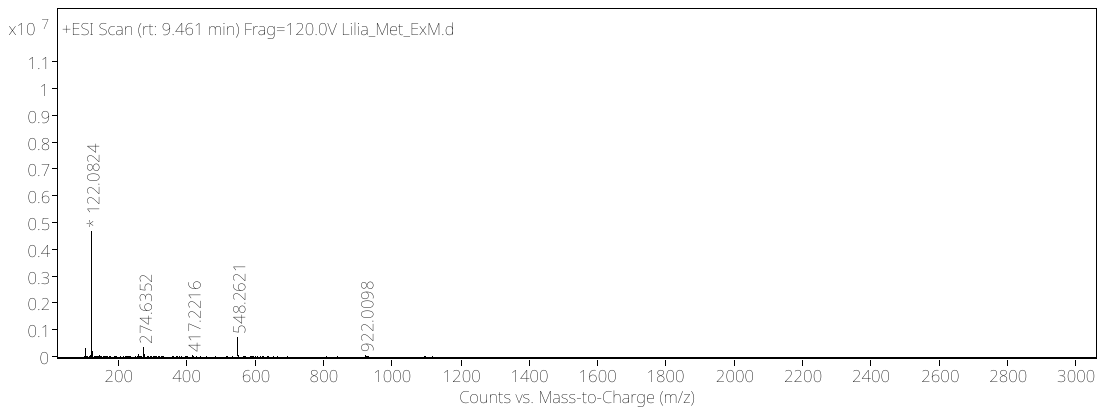

and/or

and/or

**Source Data - Supplementary Fig. 8b:**

The three conditions tested for oxidation with C_1_ peptide (CGGAGGLLGGSRGGK{Acrylic Acid}): ExM (top row), in solution with trypsin (middle row), in solution with trypsin and APS, TEMED (bottom row). LC/Q-ToF total ion chromatogram for each condition (center) and arrows pointing towards mass spectrum of trypsinized N-terminal fragment: cysteine peptide CGGAGGLLGGSR (exact mass: 1003.4869), cystine peptide (exact mass: 2004.9582), sulfinate peptide (exact mass: 1035.4767), sulfonate peptide (exact mass: 1051.4717) and trypsinized C-terminal fragment GLGK{acylic acid} (exact mass: 314.1590). The circled mass on the mass spectrum matches the peptide overlayed on the graph.

**Oxidation – cysteine results**

In solution with trypsin,

APS, TEMED

In solution with trypsin

ExM

Mass spectrum of peak

Mass spectrum of peak

LC/Q-ToF Total ion chromatogram – C18 column, method 1

**Condition**

**Source Data - Supplementary Fig. 8c:**

The three conditions tested for oxidation with Y_1_ peptide (YGGAGRGLGK{acrylic acid}): ExM (top row), in solution with trypsin (middle row), in solution with trypsin and APS, TEMED (bottom row). LC/Q-ToF total ion chromatogram for each condition (center) and arrows pointing towards mass spectrum of trypsinized N-terminal fragment: tyorisine peptide YGGAGR (exact mass: 579.2765), oxidized (DOPA or DOPA quinone) tyrosine peptide (exact mass: 595.2714), and trypsinized C-terminal fragment GLGK{acylic acid} (exact mass: 427.2431). The circled mass on the mass spectrum matches the peptide overlayed on the graph.

LC/Q-ToF Total ion chromatogram – C18 column, method 1

In solution with trypsin,

APS, TEMED

In solution with trypsin

ExM

Mass spectrum of peak

Mass spectrum of peak

**Condition**

**Oxidation – tyrosine results**

**Source Data - Supplementary Fig. 8e:**

The three conditions tested for oxidation with W_1_ peptide (WGGAGRGLGK{acrylic acid}): ExM (top row), in solution with trypsin (middle row), in solution with trypsin and APS, TEMED (bottom row). LC/Q-ToF total ion chromatogram for each condition (center) and arrows pointing towards mass spectrum of trypsinized N-terminal fragment: tryptophan peptide WGGAGR (exact mass: 602.2925), oxidized (NFK or DiOia) tryptophan peptide (exact mass: 634.2823), and trypsinized C-terminal fragment GLGK{acylic acid} (exact mass: 427.2431). The circled mass on the mass spectrum matches the peptide overlayed on the graph.

In solution with trypsin

In solution with trypsin,

APS, TEMED

**Oxidation – tryptophan results**

ExM

Mass spectrum of peak

Mass spectrum of peak

LC/Q-ToF Total ion chromatogram – C18 column, method 1

**Condition**

**Source Data - Supplementary Fig. 8a:**

The three conditions tested for oxidation with M_1_ peptide (MGGAGRGLGK{acrylic acid}): ExM (top row), in solution with trypsin (middle row), in solution with trypsin and APS, TEMED (bottom row). LC/Q-ToF total ion chromatogram for each condition (center) and arrows pointing towards mass spectrum of trypsinized N-terminal fragment: methionine peptide MGGAGR (exact mass: 547.2537), oxidized (methionine sulfoxide) methionine peptide (exact mass: 563.2486), and trypsinized C-terminal fragment GLGK{acylic acid} (exact mass: 427.2431). The circled mass on the mass spectrum matches the peptide overlayed on the graph. The thick arrow points to a zoomed in mass spectrum to show the mass of interest.

In solution with trypsin,

APS, TEMED

In solution with trypsin

**Oxidation – methionine results**

**Condition**

LC/Q-ToF Total ion chromatogram – C18 column, method 1

Mass spectrum of peak

Mass spectrum of peak

ExM

**Source Data - Supplementary Fig. 8h:**

The three conditions tested for oxidation with R_1_ peptide (RGGAGRGLGK{acrylic acid}): ExM (top row), in solution with trypsin (middle row), in solution with trypsin and APS, TEMED (bottom row). LC/Q-ToF total ion chromatogram for each condition (center) and arrows pointing towards mass spectrum of trypsinized N-terminal fragment: arginine peptide RGGAGR (exact mass: 572.3143), double trypsinized peptide: GGAGR (exact mass: 416.2132), and trypsinized C-terminal fragment GLGK{acylic acid} (exact mass: 427.2431). The circled mass on the mass spectrum matches the peptide overlayed on the graph.

**Oxidation – arginine results**

In solution with trypsin,

APS, TEMED

In solution with trypsin

**Condition**

LC/Q-ToF Total ion chromatogram – C18 column, method 1

Mass spectrum of peak

Mass spectrum of peak

ExM

**DMSO – LC/MS results**

**Condition**

LC/Q-ToF Total ion chromatogram – C18 column, method 1

Mass spectrum of peak

Mass spectrum of peak

Control peptide

Control peptide

Control peptide

DMSO

Control peptide

PITC:DMSO

**Source Data - Fig. 3c:**

In-gel Edman degradation with DMSO for conjugation buffer on “A15-peptide” (AGGAGGLLGGSRGGK{acrylic acid}) that is trypsinized from gel.

DMSO only (1^st^ row), 1:1000 ratio PITC to DMSO (2^nd^ row), DMSO then TFA (3^rd^ row), 1:1000 ratio PITC to DMSO followed by TFA (4^th^ row), trypsin and buffer only (5^th^ row). LC/Q-ToF total ion chromatogram for each condition (center) and arrows pointing towards mass spectrum of trypsinized fragments: original peptide: AGGAGGLLGGSR, peptide with PITC: *PTC*-AGGAGGLLGGSR, N-terminal cleaved: GGAGGLLGGSR, and control peptide: AGGAGK{acr}GLR.

Control peptide

DMSO +TFA

PITC:DMSO

+TFA

trypsin and control peptide only

**Formamide – LC/MS results**

**Condition**

LC/Q-ToF Total ion chromatogram – C18 column, method 1

Mass spectrum of peak

Mass spectrum of peak

Control peptide

Formamide

PITC:formamide

Control peptide

Control peptide

**Source Data - Fig. 3b:**

In-gel Edman degradation with formamide for conjugation buffer on “A15-peptide” (AGGAGGLLGGSRGGK{acrylic acid}) that is trypsinized from gel.

Formamide only (1^st^ row), 1:1000 ratio PITC to formamide (2^nd^ row), formamide then TFA (3^rd^ row), 1:1000 ratio PITC to formamide followed by TFA (4^th^ row), trypsin and buffer only (5^th^ row). LC/Q-ToF total ion chromatogram for each condition (center) and arrows pointing towards mass spectrum of trypsinized fragments: original peptide: AGGAGGLLGGSR, peptide with PITC: *PTC*-AGGAGGLLGGSR, N-terminal cleaved: GGAGGLLGGSR, and control peptide: AGGAGK{acr}GLR.

Formamide +TFA

Control peptide

PITC:formamide

+TFA

Control peptide

trypsin and control peptide only

Control peptide

Control peptide

Control peptide

**ClickP – LC/MS results**

**Condition**

ClickP:DMSO

DMSO

DMSO+TFA

ClickP:DMSO

+TFA

trypsin and control peptide only

LC/Q-ToF Total ion chromatogram – C18 column, method 1

Mass spectrum of peak

Mass spectrum of peak

Control peptide

**Source Data - Fig. 6c:**

In-gel Edman degradation with DMSO for conjugation buffer on “A15-peptide” (AGGAGGLLGGSRGGK{Acrylic Acid}) that is trypsinized from gel.

DMSO only (1^st^ row), 1:1000 ratio ClickP to DMSO (2^nd^ row), DMSO then TFA (3^rd^ row), 1:1000 ratio ClickP to DMSO followed by TFA (4^th^ row), trypsin and buffer only (5^th^ row). LC/Q-ToF total ion chromatogram for each condition (center) and arrows pointing towards mass spectrum of trypsinized fragments: original peptide: AGGAGGLLGGSR, peptide with ClickP: *ClickP*-AGGAGGLLGGSR, N-terminal cleaved: GGAGGLLGGSR, and control peptide: AGGAGK{acr}GLR.

Control peptide

Control peptide

**Condition**

Mass spectrum of peak

Mass spectrum of peak

LC/Q-ToF Total ion chromatogram – C18 column, method 1

**FITC – LC/MS results**

0.1M NaHCO_3_

Control peptide

FITC:0.1M NaHCO_3_

**Source Data - Fig. 3d:**

In-gel Edman degradation with 0.1 sodium bicarbonate pH 9 (NaHCO_3_) for conjugation buffer on “A15-peptide” (AGGAGGLLGGSRGGK{Acrylic Acid}) that is trypsinized from gel.

Sodium bicarbonate pH 8.5 (NaHCO_3_) only (1st row), 5.9 mM with 23:77 DMSO:0.1 M NaHCO_3_ pH 8.5 (2nd row), NaHCO_3_ pH 8.5 then TFA (3rd row), 5.9 mM with 23:77 DMSO:0.1 M NaHCO_3_ pH 8.5 followed by TFA (4th row), trypsin and buffer only (5th row). LC/Q-ToF total ion chromatogram for each condition (center) and arrows pointing towards mass spectrum of trypsinized fragments: original peptide: AGGAGGLLGGSR, peptide with FITC: *FTC*-AGGAGGLLGGSR, N-terminal cleaved: GGAGGLLGGSR, and control peptide: AGGAGK{acr}GLR.

Control peptide

0.1 MNaHCO3+TFA

FITC:0.1M NaHCO_3_

+TFA

Control peptide

trypsin and control peptide only

Control peptide

Control peptide

Side product of

*PTC*-AGGAGGLLGGSR

1 round, + PITC

DMSO

DMSO+PITC

DMSO +TFA

DMSO + PITC

+TFA

Mass spectrum of peak

Mass spectrum of peak

LC/Q-ToF Total ion chromatogram – C18 column, method 1

**Condition**

**Multiround DMSO – LC/MS results**

Side product of

*PTC*-GAGGLLGGSR

Zoom in

Side product of

*PTC*-GGAGGLLGGSR

LC/Q-ToF Total ion chromatogram – C18 column, method 1

Mass spectrum of peak

Mass spectrum of peak

Uncleaved

*PTC*-GAGGLLGGSR

Side product of

*PTC*-GGAGGLLGGSR

Side product of

*PTC*-GGAGGLLGGSR

2 rounds + PITC,

+ TFA

2 rounds + PITC

1 round, + PITC

+ TFA

**Condition**

**Condition**

LC/Q-ToF Total ion chromatogram – C18 column, method 1

Mass spectrum of peak

Mass spectrum of peak

**Source Data - Fig. 5b:**

Multiround in-gel Edman degradation with DMSO as conjugation buffer on “A15-peptide” (AGGAGGLLGGSRGGK{Acrylic Acid}) that is trypsinized from gel.

DMSO only (1^st^ row), 1:1000 ratio PITC to DMSO (2^nd^ row), DMSO then TFA (3^rd^ row), 1:1000 ratio PITC to DMSO followed by TFA (4^th^ row), 1 round of in-gel Edman degradation then 1:1000 ratio PITC to DMSO (5^th^ row), 1 round of in-gel Edman degradation then 1:1000 ratio PITC to DMSO followed by TFA (6^th^ row), 2 round of in-gel Edman degradation then 1:1000 ratio PITC to DMSO (7^th^ row), 2 round of in-gel Edman degradation then 1:1000 ratio PITC to DMSO followed by TFA (8^th^ row). LC/Q-ToF total ion chromatogram for each condition (center) and arrows pointing towards mass spectrum of trypsinized fragments: original peptide: AGGAGGLLGGSR, peptide with PITC: *PTC*-AGGAGGLLGGSR, peptide with N-terminal cleaved: GGAGGLLGGSR, peptide with N-terminal cleaved with PITC: *PTC*-GGAGGLLGGSR, peptide with 2 N-termini cleaved: GAGGLLGGSR, peptide with 2 N-termini cleaved with PITC: *PTC*-GAGGLLGGSR, peptide with 3 N-termini cleaved: AGGLLGGSR, and control peptide: AGGAGK{acr}GLR.

trypsin and control peptide only
