## Supplementary material for "Principles of *in situ* protein sequencing: expansion microscopy-adapted Edman degradation and amino acid recognition": Supp Figs, Notes, and tables

- Supplementary Materials
  - Supplementary Figures
    - Supplementary Figure 1
    - Supplementary Figure 2
    - Supplementary Figure 3
    - Supplementary Figure 4
    - Supplementary Figure 5
    - Supplementary Figure 6
    - Supplementary Figure 7
    - Supplementary Figure 8
    - Supplementary Figure 9
    - Supplementary Figure 10
    - Supplementary Figure 11
    - Supplementary Figure 12
    - Supplementary Figure 13
    - Supplementary Figure 14
    - Supplementary Figure 15
    - Supplementary Figure 16
    - Supplementary Figure 17
    - Supplementary Figure 18
    - Supplementary Figure 19
  - Supplementary Notes
    - Supplementary Note 1
    - Supplementary Note 2
    - Supplementary Note 3
    - Supplementary Note 4
    - Supplementary Note 5
    - Supplementary Note 6
    - Supplementary Note 7
    - Supplementary Note 8
    - Supplementary Note 9
    - Supplementary Note 10
  - Supplementary Tables
    - Supplementary Table 1
    - Supplementary Table 2

Supplementary Table 3
Supplementary Table 4
Supplementary Table 5
Supplementary Table 6
Supplementary Table 7
Supplementary Table 8
Supplementary Table 9
Supplementary Table 10
Supplementary Table 11
Supplementary Table 12

**Supplementary Figures**

#### Supplementary Figure 1

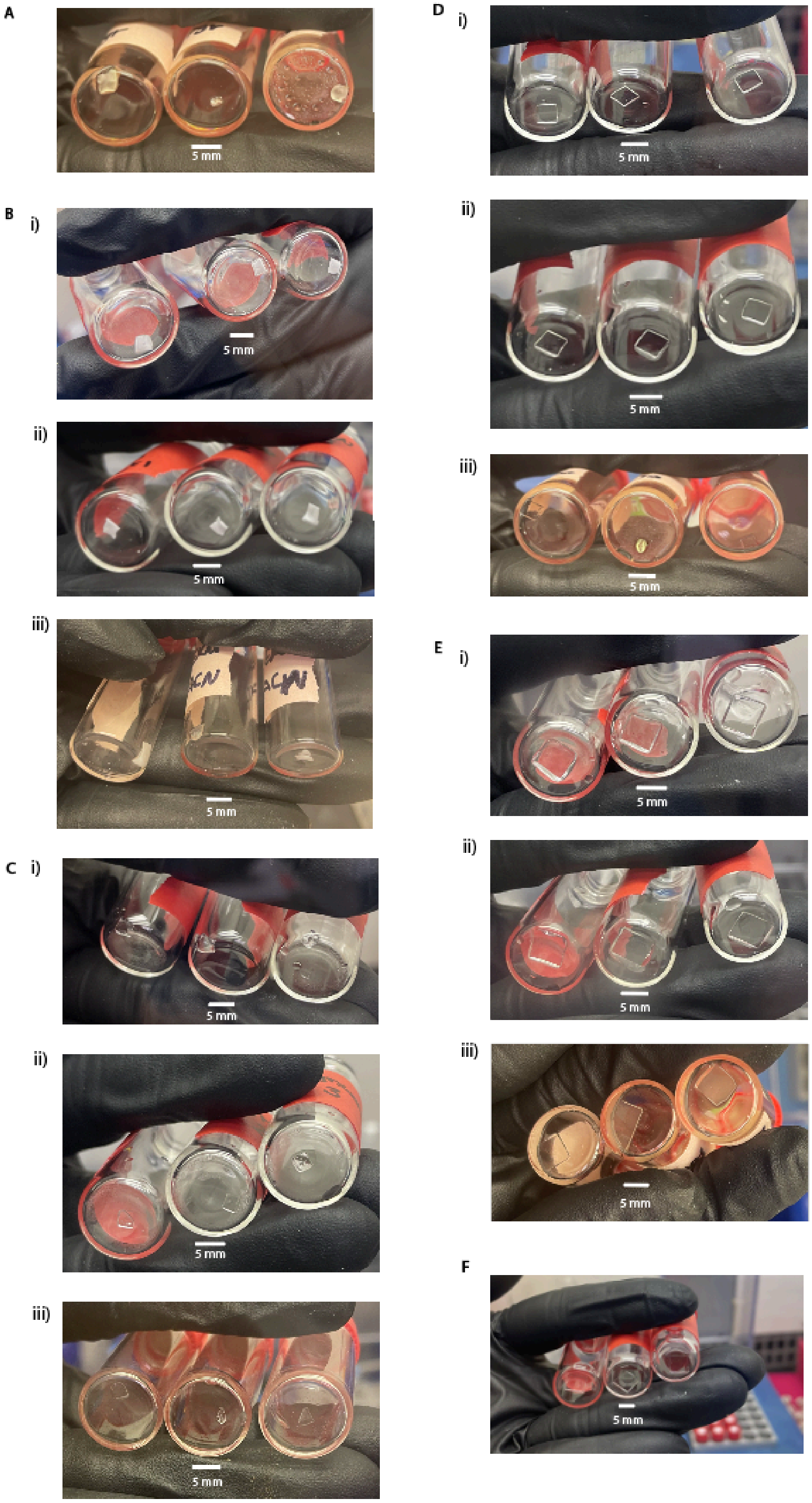

G

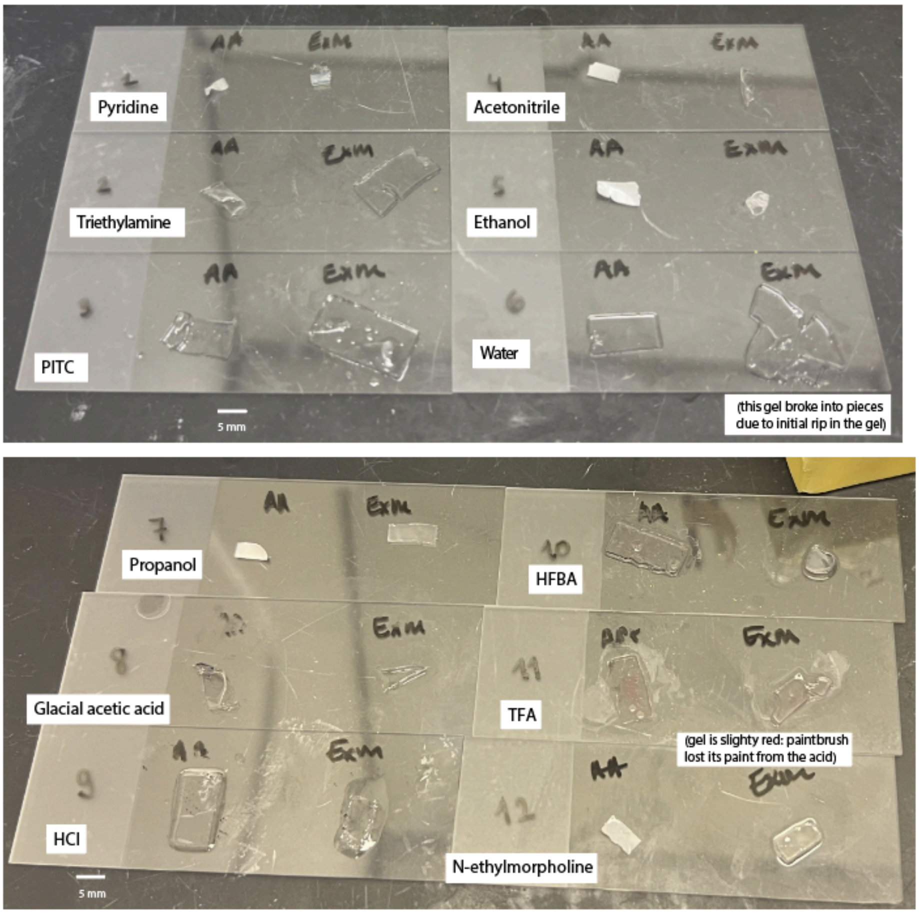

H

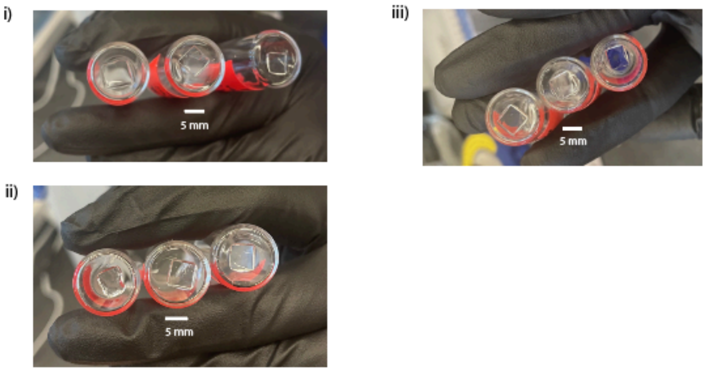

**Supplementary Figure 1**

Images of various gels (acrylamide, ExM, ExMre) after solvent with/without PITC. All gels were cut to 5 mm x 5 mm (the height depended on the gel type, ~0.3 mm for acrylamide, ~0.9 mm for ExM, ~0.7 mm for ExMre, i.e., a final gel volume of 7.8 µL, 22.8 µL and 17.5 µL, respectively) and washed 3 times with 1X PBS at room temperature (RT). ExM gels were made using 12:4:2:1:1 of StockX:water:2% BIS (w/v):10% APS (w/v):10% TEMED (v/v), as depicted in Fig. 2**B** (see Methods: ExM gel: making the empty gel gelation mixture for details), unless otherwise stated (all ratios throughout are of volumes added, unless otherwise indicated). For the gels treated with PITC to solvent (1:9 ratio PITC:solvent), they were submerged 30 min in 270 µL of solvent at 50 °C, and another 30 min at 50 °C with an additional 30 µL of PITC without removing the original solvent. All solution was then removed, and gels were imaged in glass vials, from the bottom (Supplementary Figure 1**A, Biii, Ciii, Diii, Eiii**). For all the other gels, 300 µL was added to the gels at 50 °C for 30 min unless otherwise noted, and the gel flat/top side surface size was measured through the glass vial (Supplementary Figure 1**Bi, Ci, Di, Ei, F, Hi-iii**); for the PITC ratio 1:1000 PITC:solvent case, the previous solution was then removed, and then 300 µL of a fresh solution was then added to the same gel, with PITC at a final ratio of 1:1000 ratio PITC:solvent (again, at 50 °C for 30 min). The solution was removed, and gels were similarly measured through the bottom of the glass vial after removal (Supplementary Figure 1**Bii, Cii, Dii, Eii**). In more detail, the gel flat/top side surface size was measured through the bottom of the glass vial after solution removal. In cases where the gel tore, the edges were carefully pushed back together prior to measurement. When the gel folded into a solid form and could not be unfolded, measurements were taken from the accessible edges by rotating the vial to obtain the most representative size and calculate the flat/top surface size as if it were unfolded (Supplementary Figure 1**A, Ciii, Diii)** . Flat/top surface sizes of gels are plotted in Supplementary Figure 2. Scale bar in white (using the central vial from the image as the reference for Supplementary Figure 1**A-F, H**; using the bottom leftmost microscope slide as reference for Supplementary Figure 1**G**; please note that given the 3-d nature of the vials and specimens, this means the scale bar is approximate for the other specimens): 5 mm.

(A) Gels submerged in pyridine followed by PITC to pyridine (1:9 ratio PITC:pyridine). Different gels are in three separate glass vials: a 9% acrylamide gel (left), ExM gel (middle) and ExMre gel (right).

(B) Gels submerged in acetonitrile (ACN) with/without PITC (1:9 and 1:1000 ratios PITC:ACN) (i) ExMre gels submerged in ACN without PITC. 3 different ExMre gels in separate glass vials, from different gelation solutions. (ii) Same gels as (i), after removal of solution and further submerging the same gels in fresh solution of PITC to ACN (1:1000 ratio PITC:ACN). (iii) 3 different gel types, 9% acrylamide gel (left), ExM gel (middle) and ExMre gel (right), treated with ACN followed by PITC to ACN(1:9 ratio PITC:ACN).

(C) As in B, but with 1:1 pyridine to water as solvent. For panel (iii) placement of the gels differs from (B): ExMre gel (left), ExM gel (middle) and a 9% acrylamide gel (right).

(D) As in B, but with DMSO as solvent. For panel (iii): a 9% acrylamide gel (left), ExM gel (middle) and ExMre gel (right).

(E) As in B, but with formamide as solvent (note: the gels imaged for (i) and (ii) are not the same gels that were used for gel size change results in Fig. 2**Civ**): a 9% acrylamide gel (left), ExM gel (middle) and ExMre gel (right).

(F) ExMre gels submerged in TFA. 3 different ExMre gels in separate glass vials from different gelation solutions.

(G) Other solvent testing, based on protocols listed in Supplementary Table 2, with various solvents used in conjugation and cleavage. 9% acrylamide gels and ExM gels (note: the gelation formula for ExM gels was different here, with 48:1:1 of StockX:10% TEMED (v/v):10% APS (w/v), see Methods: ExM gel: making the empty gel gelation mixture for StockX formula) tested in various solvents including pyridine, triethylamine, PITC, acetonitrile, ethanol, water, propanol, glacial acetic acid, hydrochloric acid (abbreviated: HCl), heptafluorobutyric acid (abbreviated: HFBA), trifluoroacetic acid (TFA), and N-ethylmorpholine. Each gel was washed twice with 1x phosphate buffered saline (PBS for 10 min) and then submerged in organic solvent for 10 min.

(H) ExMre gels submerged in various aqueous buffers. 3 different ExMre gels in separate glass vials from different gelation solutions. (i) 0.1 M sodium bicarbonate pH 8.5 at 50 °C. (ii) 1M Tris pH 8 at room temperature (RT). (iii) 1M Tris pH 9.5 at RT.

### Supplementary Figure 2

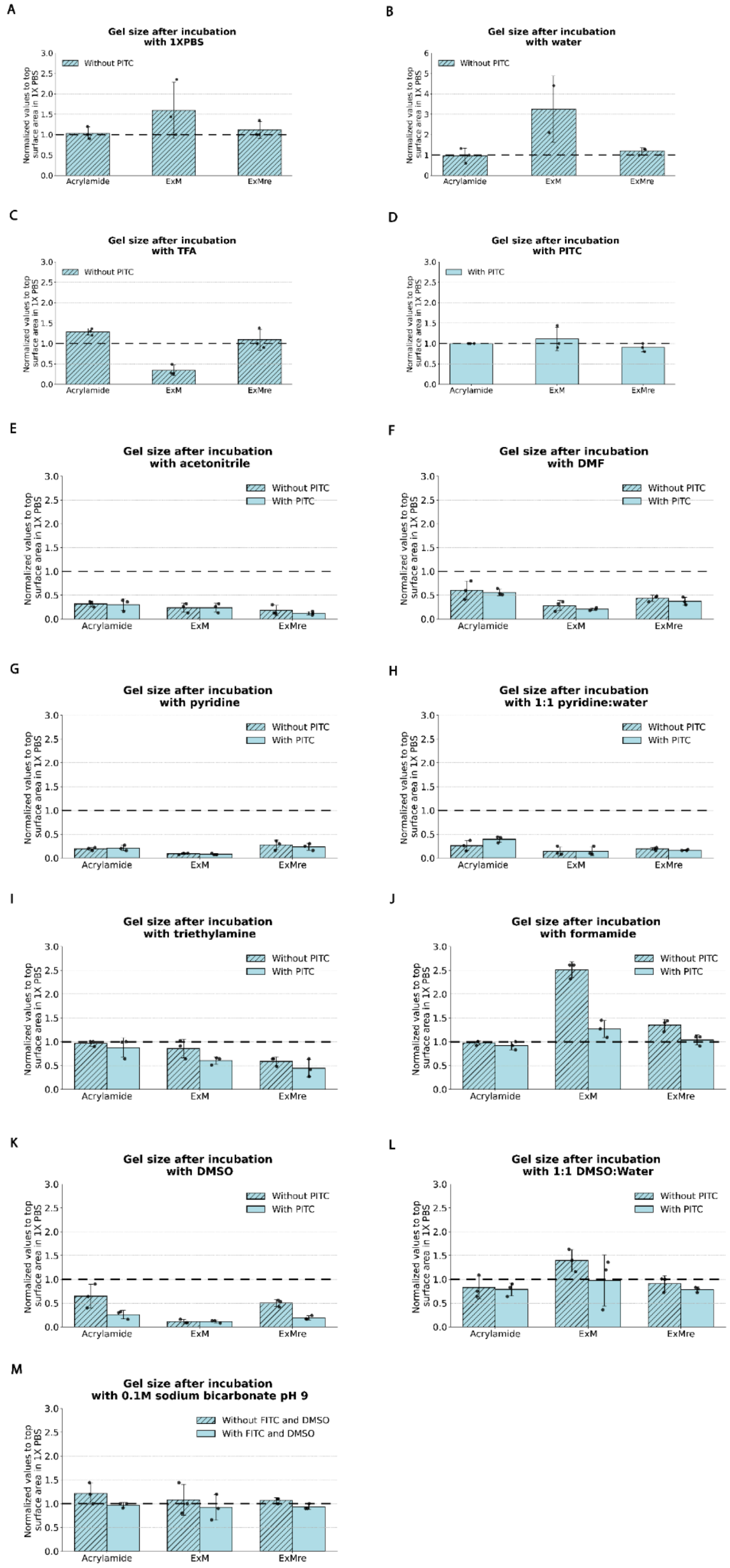

**Supplementary Figure 2: Surface flat/top side size of acrylamide, ExM, and ExMre gels in various solvents with/without PITC or FITC.** Surface size of flat/top side surface size of acrylamide, ExM, and ExMre gels when placed in various solutions (some previously documented for Edman degradation, as in **Supplementary Table 2**), normalized to surface size (again, flat/top side surface size) of the gel in 1X PBS before various solution treatment. All gels were cut to 5 mm x 5 mm (the height depended on the gel type, ~0.3 mm for acrylamide, ~0.9 mm for ExM, ~0.7 mm for ExMre, i.e., a final gel volume of 7.8 µL, 22.8 µL and 17.5 µL, respectively) and washed 3 times with 1X PBS at room temperature (RT). For gels treated with 1X PBS, water, TFA, PITC, they were submerged 30 min with 300 µL of solution for 30 min at 50 °C (**Supplementary Figure 2A-D**). For the gels treated with PITC (1:9 ratio PITC:solvent), they were submerged 30 min in 270 µL of solvent at 50 °C, measured, then another 30 min at 50 °C with 30 µL of PITC in the original 270 µL solvent. All solution was then removed, and gels were measured again (**Supplementary Figure 2E-J**). For the gels treated with FITC, they were first submerged in 231 µL of 0.1 M sodium bicarbonate pH 8.5, at 50 °C, measured, then another 30 min at 50 °C with 69 µL of 10 mg/mL FITC in DMSO. (A) 1X PBS, (B) water, (C) TFA, (D) 100% PITC, (E) acetonitrile (ACN), (F) dimethylformamide (DMF), (G) 100% pyridine, (H) 1:1 pyridine to water, (I) 100% triethylamine, (J) 100% formamide, (K) dimethylsulfoxide (DMSO), (L) 1:1 DMSO to water, (M) 0.1 M sodium bicarbonate pH 8.5. “With PITC” is PITC added (1:9 ratio PITC:solution) in the given solution (unlike **Figure 2C**, where PITC is added at 1:1000 ratio PITC:solvent. (Dashed black line: surface size of the flat side of the gel in 1X PBS; error bar: standard deviation; black dots, individual experiments; n=3 gels from the same starting gelation solution).

### Supplementary Figure 3

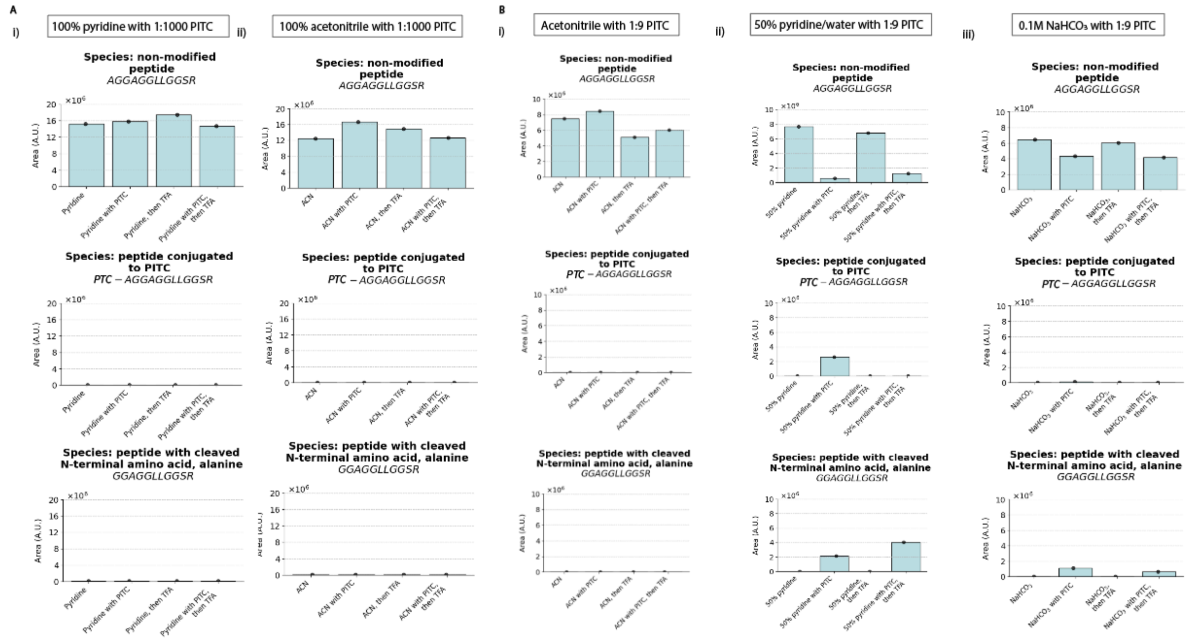

**Supplementary Figure 3:**

(A) In-gel Edman degradation with A15-peptide (as used in **Fig. 3A-D**) in ExMre gels. Here 100% pyridine or acetonitrile (ACN) were tested as conjugation solvent, PITC as Edman reagent (1:1000 ratio PITC:solvent). Bar graphs representing the relative abundance (arbitrary units, a.u.; all samples were processed with spiked-in controls; see **Methods**) of different peptide ion species detected on the LC/QToF. The bar graphs are obtained from comparing the area under the curve (AUC) of the chromatogram of various species (extracted based on the exact mass, see **Analysis of LC/QToF data**). The separate conditions compared include solvent only ("pyridine" or "ACN"), PITC to solvent (1:1000 ratio PITC:solvent) for 1 hour at 50 °C (e.g., "pyridine with PITC"), TFA for 30 min at 50 °C (e.g., "pyridine, then TFA"), PITC to solvent (1:1000 ratio PITC:solvent) for 1 hour at 50 °C followed by TFA for 30 min at 50 °C (e.g., "pyridine with PITC, then TFA"). The graphs in (i) were performed with 100% pyridine, and in (ii) were performed with ACN, as solvent for in-gel Edman conjugation. The relative abundance of the ion species: (top) non-modified peptide (AGGAGGLLGGSR), (middle) peptide conjugated to PITC, phenylthiocarbamoyl (PTC)-peptide (*PTC*-AGGAGGLLGGSR), and (bottom) peptide with cleaved N-terminal amino acid (GGAGGLLGGSR), were reported throughout the in-gel Edman degradation process in the various conditions (black dots, individual experiments; blue bar, mean; n=1 gelation solution).

(B) In-gel Edman degradation, as in (A), but with (i) ACN, (ii) 1:1 pyridine to water, and (iii) 0.1 M $NaHCO_3$ pH 8.5 as conjugation buffer, PITC as Edman reagent (PITC 1:9 ratio PITC:buffer). See **Methods** for further details on this protocol (black dots, individual experiments; blue bar, mean; error bar: standard deviation; n=1 gelation solution).

#### Supplementary Figure 4

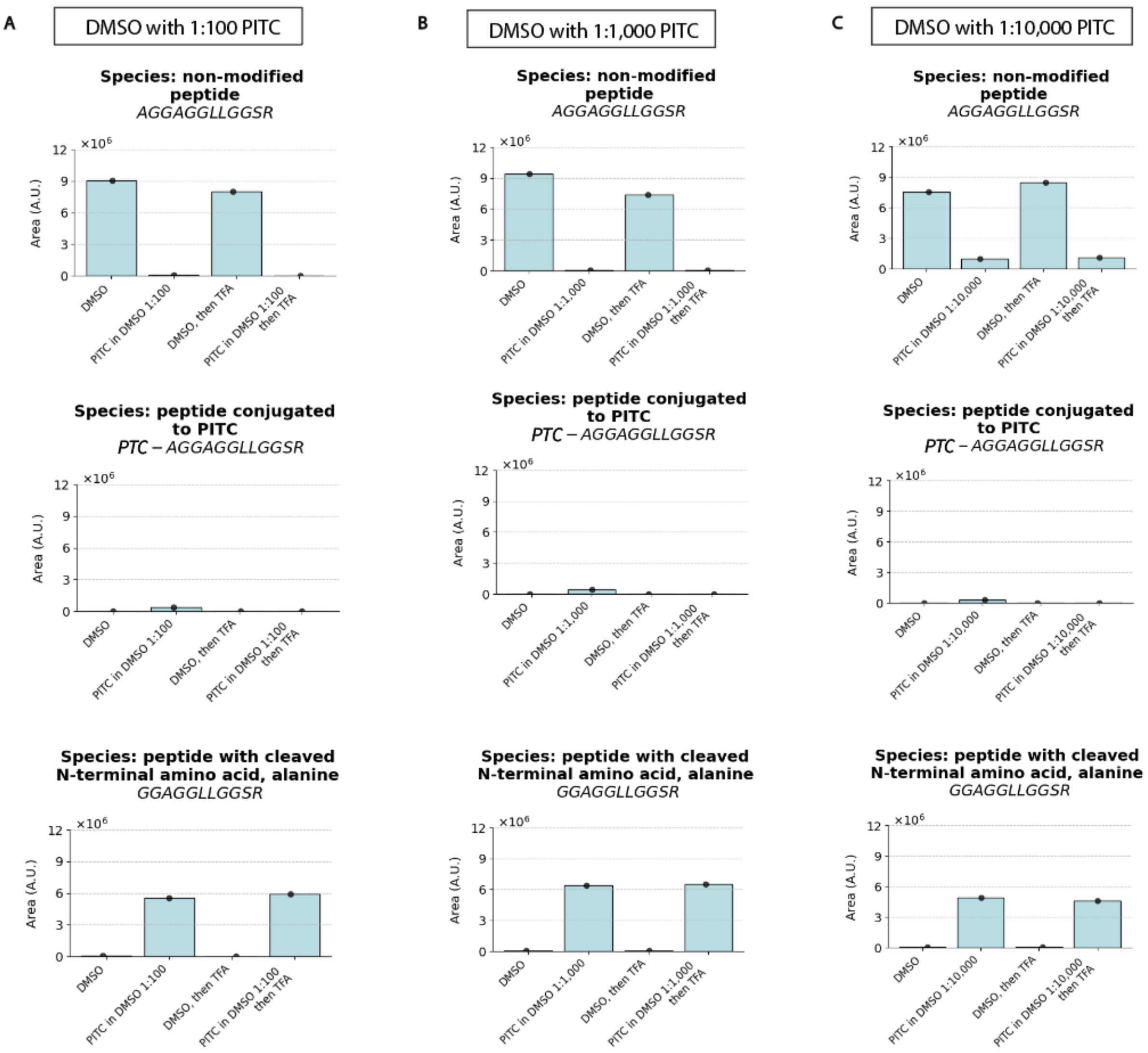

**Supplementary Figure 4**

(A) In-gel Edman degradation with A15 peptide in ExMre gels. Here dimethylsulfoxide (DMSO) is tested as a conjugation solvent, PITC as an Edman reagent at different ratios in the solvent. Bar graphs representing the relative abundance (arbitrary units, a.u.; all samples were processed with spiked-in controls; see Methods) of different peptide ion species detected on the LC/QToF. The bar graphs are obtained from comparing the area under the curve (AUC) of the chromatogram of various species (extracted based on the exact mass, see Analysis of LC/QToF data). The separate conditions compared include DMSO only ("DMSO"), PITC to solvent (1:100 ratio PITC:DMSO) for 1 hour at 50 °C ("PITC to DMSO 1:100"), TFA for 30 min at 50 °C (on the graph, denoted as "DMSO, then TFA"), (4) PITC to DMSO (1:100 ratio PITC:DMSO) for 1 hour at 50 °C followed by TFA for 30 min at 50 °C ( "PITC to DMSO 1:100, then TFA"). The relative abundance of the ion species: (top) non-modified peptide (AGGAGGLLGGSR), (middle) peptide conjugated to PITC (*PTC*-AGGAGGLLGGSR), and (bottom) peptide with cleaved N-terminal amino acid (GGAGGLLGGSR), were reported throughout the in-gel Edman degradation process in the various conditions (black dots, individual experiments; blue bar, mean; error bar: standard deviation; n=1 gelation solution).

(B) As in A, but with PITC to DMSO (1:1,000 ratio PITC:DMSO) .

(C) As in A, but with PITC to DMSO (1:10,000 ratio PITC:DMSO) .

**Supplementary Figure 5**

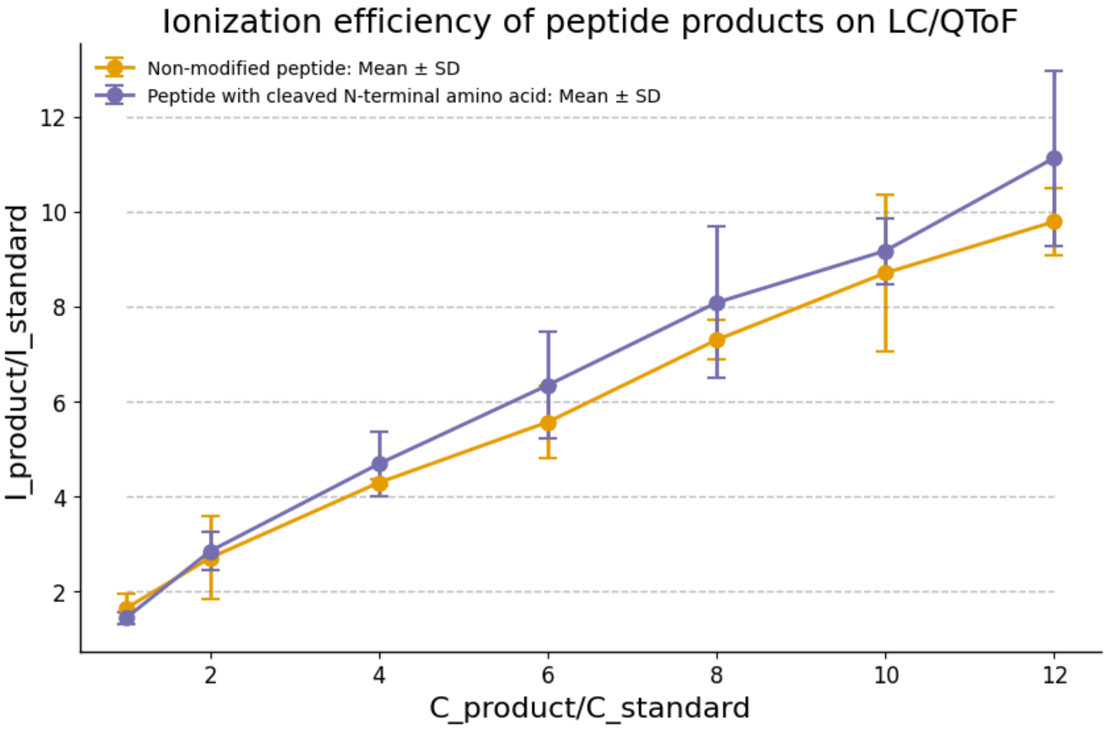

**Supplementary Figure 5: Control curve of the non-modified peptide and the peptide with cleaved N-terminal amino acid ionization efficiencies with increasing concentration.** Peptide fragments derived from the peptide denoted A15 (AGGAGGLLGGSRGGK{acr}) expected after trypsinization from the gel (denoted "product"). Two products are shown as separate control curves: the first, "non-modified peptide" (AGGAGGLLGGSR; mean value across replicates: orange dots; orange vertical lines: standard deviation of given concentration, n=3 separate peptide vials) and "peptide with cleaved N-terminal amino acid" (GGAGGLLGGSR; mean value across replicates: purple dots; purple vertical lines: standard deviation of given concentration, n=3 separate peptide vials). They are prepared in solution at various "C_product" concentrations from 5 µM to 60 µM (5, 10, 20, 30, 40, 50, 60 µM) and the area under the curve (AUC) of the extracted ion chromatogram of the two species is recorded as "I_product". The A9-peptide is denoted "standard" (AGGAGK{acr}GLR) and was used as the standard with a constant "C_standard" of 5 µM and the AUC of the extracted ion chromatogram is recorded as "I_standard" (extracted based on the exact mass, see Analysis of LC/QToF data). The statistical significance of the two means were tested with two-sided Welch's t-test for n=3 for the two conditions (non-modified peptide and peptide with cleaved N-terminal amino acid) at each "C_product/C_standard" level. None of the conditions were statistically significant at the 95% confidence level ("C_product/C_standard": 1, t-statistic: 0.847, p-value: 0.470; "C_product/C_standard": 2, t-statistic: -0.208, p-value: 0.849; "C_product/C_standard": 4, t-statistic: -0.816, p-value: 0.499; "C_product/C_standard": 6, t-statistic: -0.808, p-value: 0.470; "C_product/C_standard": 8, t-statistic: -0.672, p-value: 0.563; "C_product/C_standard": 10, t-statistic: -0.372, p-value: 0.737; "C_product/C_standard": 12, t-statistics: -0.961, p-value: 0.418). After adjusting for multiple-comparison testing, using Holm-Bonferroni, p-values are corrected to 1.0 at each concentration level.

#### Supplementary Figure 6

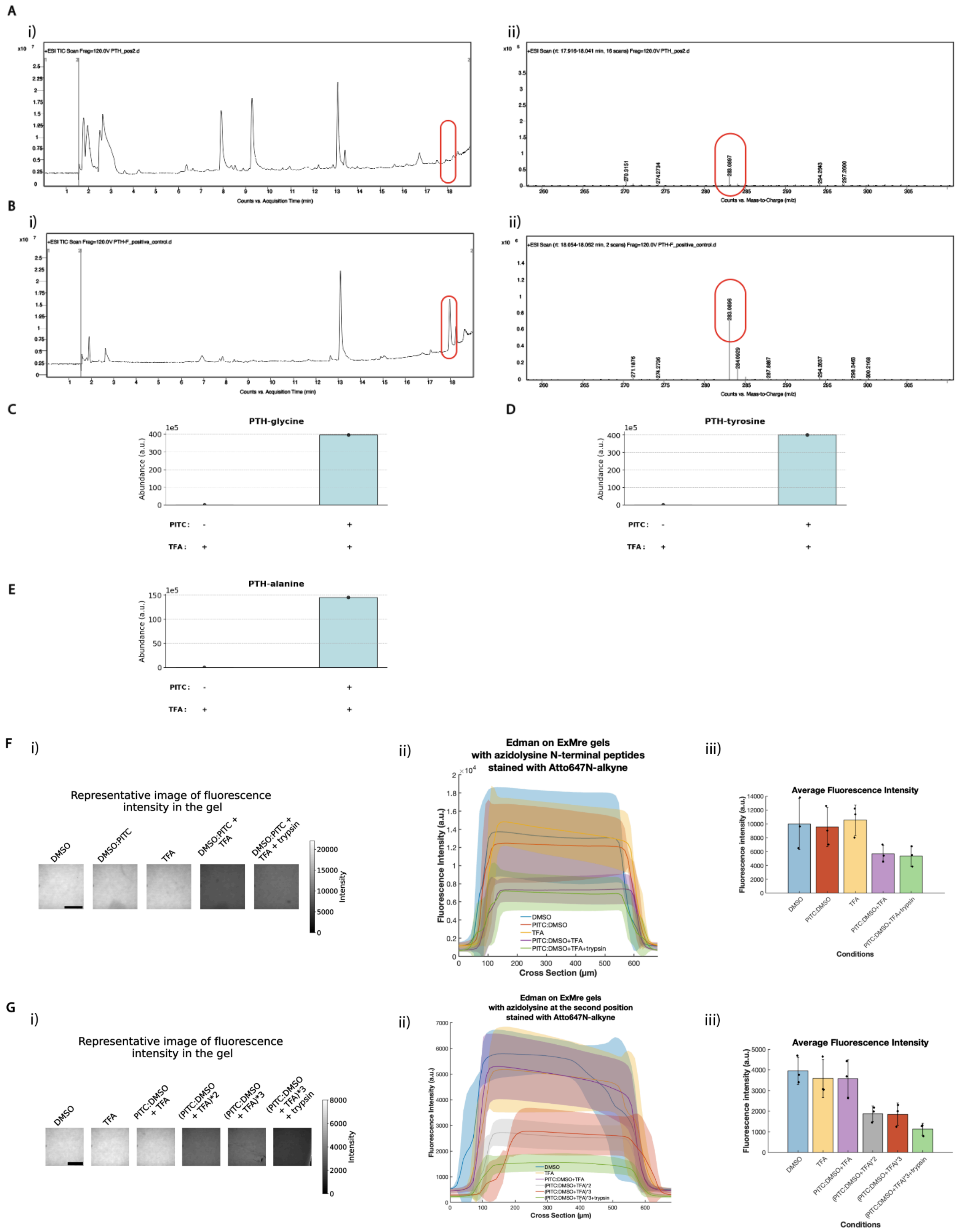

**Supplementary Figure 6:** Phenylthiohydantoin (PTH) detection assay from Fig. 3**H** for $F_1$ peptide (FGGAGRGLGK{acr}) embedded in ExMre gels

(A) (i) Total ion chromatogram (TIC) of one of the three replicates for PTH-phenylalanine (PTH-F) detection from ExMre gels containing $F_1$ peptide. Peptides in ExMre gels cut 5 mm x 5 mm x 0.35 mm (8.8 μL) with ~50 μM peptide are Edman degraded, and resuspended in 1:1

acetonitrile to water before injection in the LC/QToF (y axis: abundance in arbitrary units, x axis: retention time in minutes, red oval at ~18 min when PTH-F elutes). Considering expected (from earlier experiments) ~70% cleavage of PTH-F from the peptide, ~2 pmol is injected into the LC/QToF. (ii) The associated mass spectrum of the TIC at ~18 min showing presence of PTH-F (exact mass: 282.0827, and $[M+H]^+$: 283.0905; y axis: abundance in arbitrary units, x axis: mass to charge ratio, red oval showing detection of this species).

(B) (i) TIC of the positive control with pure PTH-F in 1:1 acetonitrile to water before injection in the LC/QToF (y axis: abundance in arbitrary units, x axis: retention time in minutes, red oval at ~18 min when PTH-F elutes). About ~1 nmol is injected into the LC/QToF, compared to ~2 pmol in (a). (ii) The associated mass spectrum of the TIC at ~18 min showing presence of PTH-F (exact mass: 282.0827, and $[M+H]^+$: 283.0905; y axis: abundance in arbitrary units, x axis: mass to charge ratio).

(C) PTH-glycine (PTH-G) detection of GGGAGRGLGK{acr} (abbreviated G-peptide) embedded in 9% acrylamide gels. Separate gels were subjected to various Edman conditions. The conditions included TFA only for 30 min at 50 °C, and PITC toDMSO (1:9 ratio PITC:DMSO) for 1 hour at 50 °C followed by TFA for 30 min at 50 °C. Subsequently, TFA was removed from the gels and they were immersed in 50 µL of 1:1 acetonitrile:water and agitated. Read-out was then performed by injecting the supernatant into LC/QToF using **LC Method (see Methods)**. Analysis of PTH-G abundance was performed using PTH-G exact mass, 193.0435 ± 0.0039 Da (see **Methods for Edman degradation and PTH detection** for details). (blue bar, mean; black dots, individual experiments, n=1 gelation solution).

(D) As in (C), but with PTH-Y detection of YGGAGRGLGK{acr} (abbreviated Y-peptide) embedded in 9% acrylamide gels. Analysis of PTH-Y abundance was performed using PTH-Y exact mass, 299.0854 ± 0.0060 Da.

(E) As in (C), but with PTH-A detection of AK{$N_3$}GAGGLLGGSRGGK{acr} embedded in ExMre gels. Analysis of PTH-A abundance was performed using PTH-A exact mass, 207.0592 ± 0.0041 Da.

(F) In-gel Edman degradation on azidolysine peptides over multiple rounds, when first staining peptides with alkyne-Atto647N prior to degradation. See **Methods** for **Yield calculations**. See **Supplementary Table 5G** for raw data and statistics. (i) A representative image of the fluorescence of the gels is depicted for each condition of in-gel Edman degradation for the K{N3}15-peptide, taking the 30th slice of the Z-stack for each (~300 µm deep into the gel, with 10 µm z steps). Imaging performed on a confocal microscope (~680 µm thick Z-stack). Scale bar, 500 µm. (ii) The gels in different conditions are compared in fluorescence intensity throughout their cross-section for the K{N3}15-peptide (line, mean; shaded area, standard deviation; n=3, different gelation solutions). (iii) Average fluorescence intensity of the ExMre gels in the different conditions for the K{N3}15-peptide by averaging across the whole volume imaged (black dots, individual experiments; colored bar, mean; error bar, standard deviation, n=3, different gelation solutions).

(G) As in (F), but for AK{N3}15-peptide and results with in-gel Edman degradation over 3 rounds. See **Supplementary Table 5H** for raw data and statistics.

#### Supplementary Figure 7

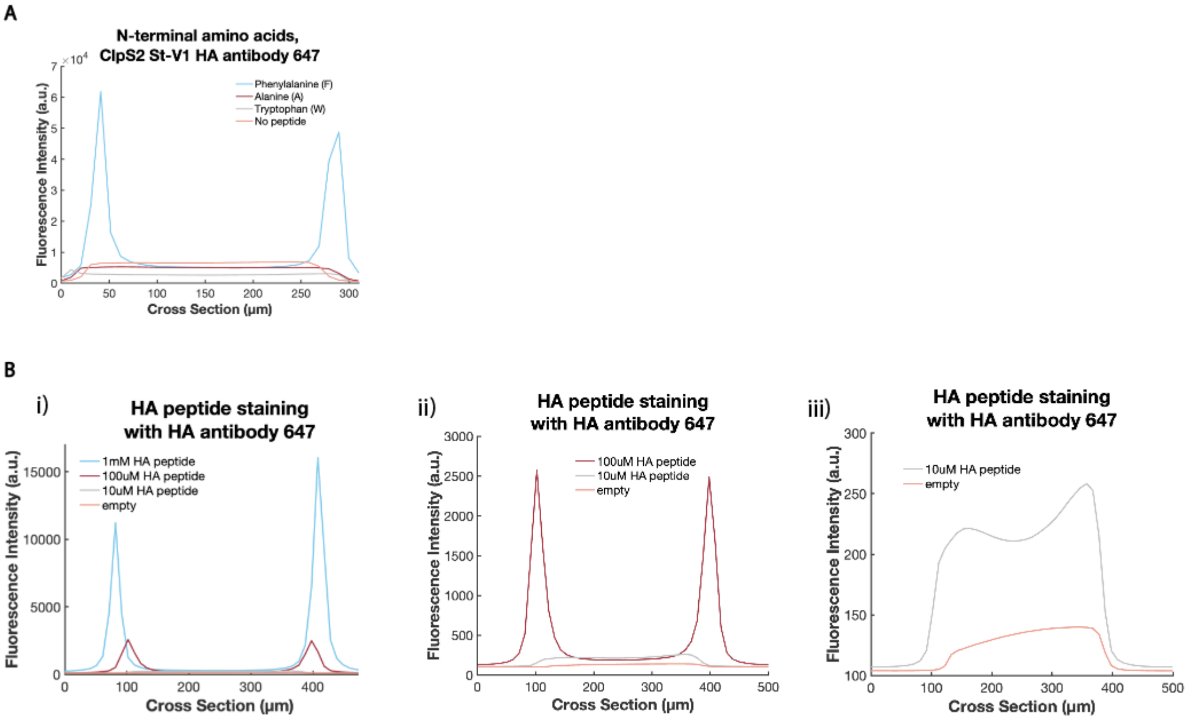

**Supplementary Figure 7:**

(A) 9% acrylamide gels were cast with different N-terminal peptides (XaaGGAGRGLGK{acr}, where Xaa was: F, A, W) at a concentration of ~5 mM or with no peptides embedded (labeled: "No peptide"). The gels (each with a volume of 1.2 µL) are compared for their fluorescence intensity across the thickness of the gel after staining with 20 µM ClpS2 St-V1 for 1 hour at RT, then washed for ~1 min, prior to incubating with 200 µg/mL anti hemagglutinin (HA) tag antibody 647 (HA-antibody 647) overnight (O/N) in 100 µL of solution at 4 °C in the dark. Imaging was performed using a confocal microscope with 10X objective, 100% laser power, 400 ms exposure time, 10 µm Z-steps.

(B) 9% acrylamide gels were cast with HA-tag peptide (YPYDVPDYAK{acr}) at a concentration of ~1 mM, ~100 µM and ~10 µM, and with no peptides embedded (labeled: "empty"). The gels are compared for their fluorescence intensity across the thickness of the gel after staining with 20 µM of ClpS2 St-V1 for 1 hour at RT and incubating with 20 µg/mL HA-antibody 647 O/N at 4 °C in the dark. Imaging was performed using a confocal microscope with 10X objective, 100% laser power, 100 ms exposure time, 10 µm Z-steps. (i) Shows all 4 conditions tested (1 mM, 100 µM, 10 µM, and empty). (ii) As in (i) but shows only 3 conditions (100 µM, 10 µM, and empty) with an adjusted y-axis. (iii) As in (ii) but shows only 2 conditions (10 µM and empty) with an adjusted y -axis.

**Supplementary Figure 8 Analysis of amino acid oxidation in ExM gels. Different N-termini peptides were embedded in ExM gels (XaaGGAGRGLGK{acr}, with Xaa: M, C, Y, W, F, H, P, R).**

**a)**

**i)** A)

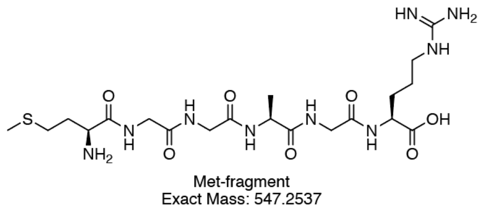

B)

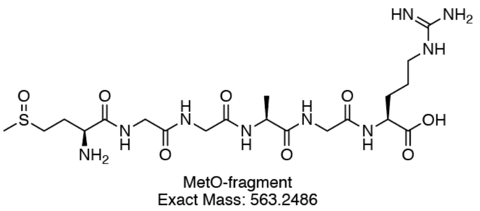

C)

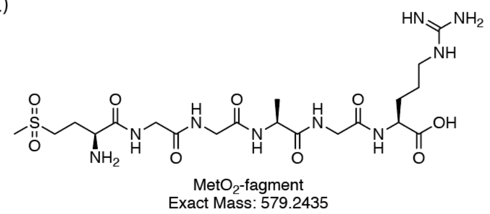

**ii)**

| **Methionine** | A) Met-fragment | B) MetO-fragment | C) $MetO_2$-fragment |
|---|---|---|---|
| *Exact mass* | *547.2537* | *563.2486* | *579.2435* |
| Condition 1: trypsin | 19,007,365 | 183,009.5 | 366.81 |
| Condition 2: TEMED+APS+trypsin | 1,021,971 | 2,801,300 | 262.52 |
| Condition 3: TEMED+APS+trypsin in ExM gel | 3,628,434.7 | 13,817,063 | 5,788.13 |

**iii)**

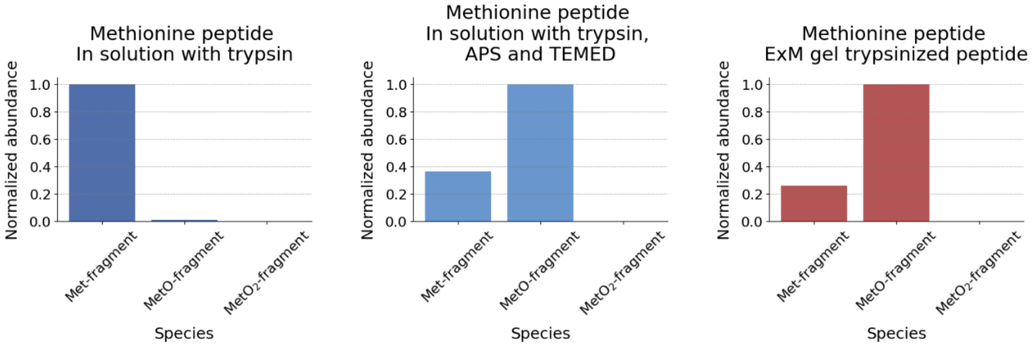

**Supplementary Figure 8a** Methionine and post-translational oxidation.
**i)** Structure of the various fragments resulting from enzymatic cleavage with trypsin from the parent peptide (sequence of parent peptide: MGGAGRGLGK{acr}), (A) “Methionine fragment” (MGGAGR, abbreviated: “Met-fragment”) resulting from trypsin cleavage, (B) modification of the fragment in (a) with known methionine oxidation products, notably methionine sulfoxide, to form “Methionine sulfoxide fragment” (abbreviated: “MetO-fragment”) (C), modification of the fragment in (a) with known methionine oxidation products, notably “Methionine sulfone fragment” (abbreviated: “$MetO_2$-fragment”) [1].

**ii)** Table detailing the exact mass, and absolute abundance recorded for the different products (from A-C peptide fragments from **Supplementary Figure 8ai**) in the different conditions (Condition 1: in solution with trypsin; Condition 2: in solution with trypsin, ammonium persulfate (APS) and N,N,N',N'-Tetramethylethylenediamine (TEMED); Condition 3: ExM gel trypsinized peptide). AUC for each species in the extracted ion chromatogram was obtained as in **Methods: Analysis of LC/QToF data** and the raw total ion chromatograms (TIC) located in **Source Data**. The dash ('-') means the species was not detected when using search parameters (the extracted ion chromatogram was generated using the $[M+H]^+$ ion of the species, with a mass tolerance of ±20 ppm, with all detected peaks integrated into the final value).
**iii)** Normalized abundance of methionine N-terminal peptide and oxidation products in different conditions. Normalized abundance of the 3 different species (Met-fragment, MetO-fragment, $MetO_2$-fragment; as detailed in **Supplementary Figure 8ai**) in: condition 1, after trypsin cleavage in solution (left); condition 2, after addition of APS and TEMED with trypsin cleavage in solution (middle); and condition 3, after trypsin cleavage of peptides from ExM gels (condition 3; right). AUC for each species in the extracted ion chromatogram was obtained as in **Methods: Analysis of LC/QToF data** and the raw total ion chromatograms (TIC) located in **Source Data**. The abundance of each species was normalized with max absolute scaling for plotting on the bar graph (n=1 gelation solution for each condition).

**b)**

i)

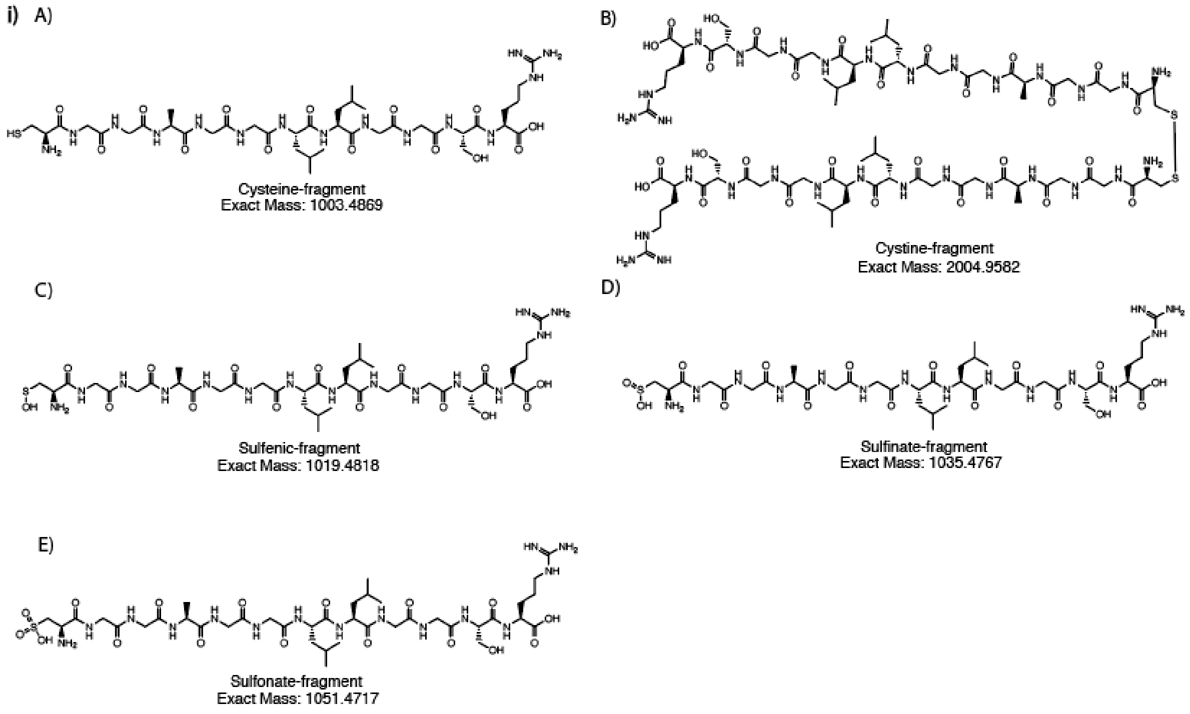

ii)

| **Cysteine** | A) Cys-fragment | C) Sulfenic-fragment | D) Sulfinate-fragment | E) Sulfonate-fragment |
|---|---|---|---|---|
| Exact mass | *1003.4869* | *1019.4818* | *1035.4767* | *1051.4717* |
| Condition 1: trypsin | 11,412,844 | 162,653.3 | 96,467.32 | 132,344.9 |
| Condition 2: TEMED+APS+trypsin | 354,068.38 | 102,825.1 | 457,772.2 | 1,327,463 |
| Condition 3: TEMED+APS+trypsin in ExM gel | 54,427.54 | 10,445.3 | 299,666.7 | 268,362.1 |

iii)

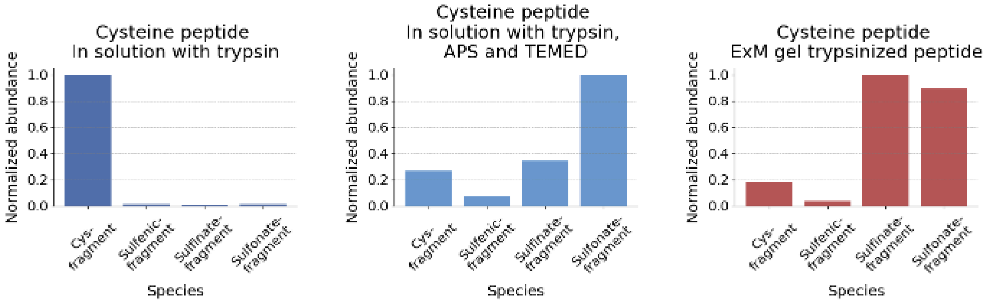

**Supplementary Figure 8b** Cysteine and post-translational oxidation.

**i)** Structure of the various fragments resulting from enzymatic cleavage with trypsin from the parent peptide (sequence of parent peptide: CGGAGGLLGGSRGGK{acr}), (A) "Cysteine fragment" (CGGAGGLLGGSR, abbreviated: "Cys-fragment") resulting from trypsin cleavage, (B) modification of the fragment in (A) with known cysteine oxidation products, notably cystine, to form "Cystine fragment" (abbreviated: "Cystine-fragment"), (C) modification of the fragment in (A) with known cysteine oxidation products, notably "Sulfenic acid fragment" (abbreviated: "Sulfenic-fragment"), (D) modification of the fragment in (A) with known cysteine oxidation products, notably "Sulfinate fragment" (abbreviated: "Sulfinate-fragment"), (E) modification of the fragment in (a) with known cysteine oxidation products, notably "Sulfonate fragment" (abbreviated: "Sulfonate-fragment") [2].

**ii)** Table detailing the exact mass, and absolute abundance recorded for the different products (from A-E peptide fragments from **Supplementary Figure 8bi;** note that Cysteine-fragment is omitted due to $[M+2]^{2+}$ mass having nearly identical mass to Cys-fragment - only the mass of Cys-fragment is plotted but may include contributions from Cysteine-fragment with one carbon-13 atom) in the different conditions (Condition 1: in solution with trypsin; Condition 2: in solution with trypsin, APS and TEMED; Condition 3: ExM gel trypsinized peptide). AUC for each species in the extracted ion chromatogram was obtained as in **Methods: Analysis of LC/QToF data** and the raw total ion chromatograms (TIC) located in **Source Data**. The dash ('-') means the species was not detected when using search parameters (the extracted ion chromatogram was generated using the $[M+H]^+$ ion of the species, with a mass tolerance of ±20 ppm, with all detected peaks integrated into the final value).
**iii**) Normalized abundance of cysteine N-terminal peptide and oxidation products in different conditions. Normalized abundance of the 5 different species (Cys-fragment, Cystine-fragment, Sulfenic-fragment, Sulfinate-fragment, Sulfonate-fragment; as detailed in **Supplementary Figure 8bi**) in: condition 1, after trypsin cleavage in solution (left); condition 2, after addition of APS and TEMED with trypsin cleavage in solution (middle); and condition 3, after trypsin cleavage of peptides from ExM gels (condition 3; right). AUC for each species in the extracted ion chromatogram was obtained as in **Methods: Analysis of LC/QToF data** and the raw total ion chromatograms (TIC) located in **Source Data**. The abundance of each species was normalized with max absolute scaling for plotting on the bar graph (n=1 gelation solution for each condition).

**c)**

i)

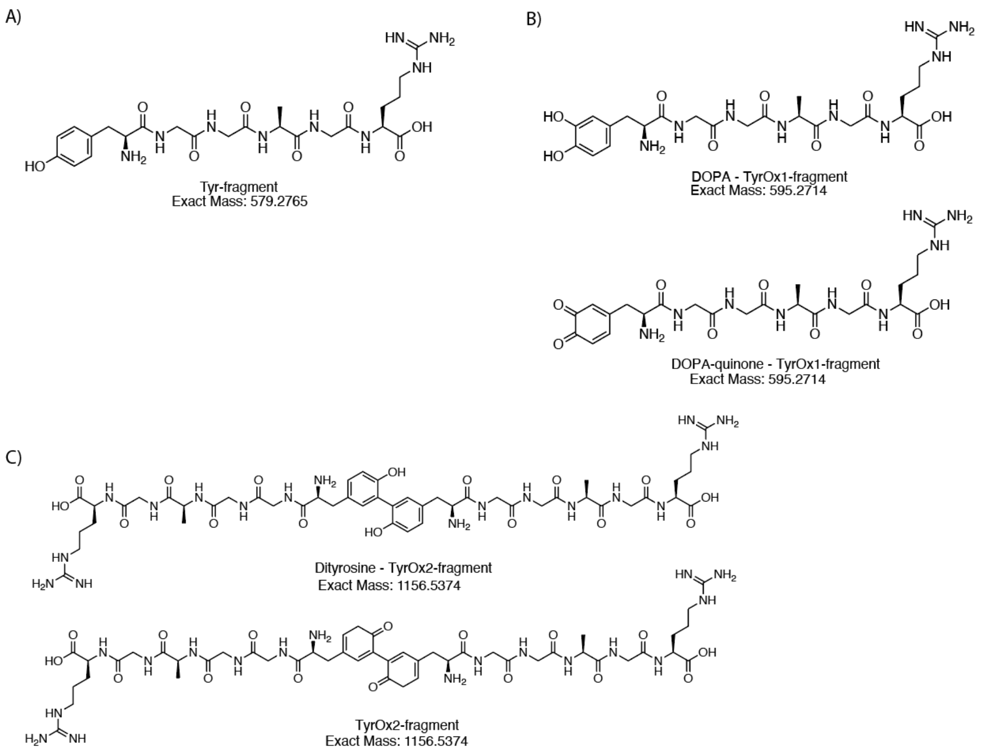

ii)

| **Tyrosine** | A) Tyr-fragment | B) TyrOx1-fragment | C) TyrOx2-fragment |
|---|---|---|---|
| *Exact mass* | *579.2765* | *595.2714* | *1156.5374* |
| Condition 1: trypsin | 28,020,395 | 18,620.58 | 240.48 |
| Condition 2: TEMED+APS+trypsin | 13,289,453 | 1,054,274 | 71,283.85 |
| Condition 3: TEMED+APS+trypsin in ExM gel | 2,509,423.3 | 30,242.02 | 255.59 |

iii)

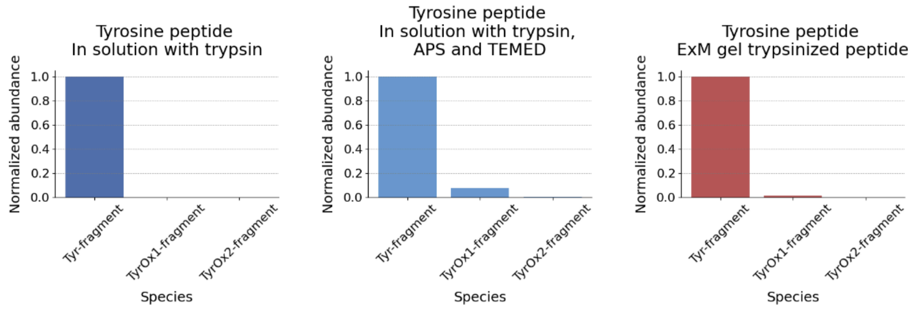

**Supplementary Figure 8c** Tyrosine and post-translational oxidation

**i)** Structure of the various fragments resulting from enzymatic cleavage with trypsin from the parent peptide (sequence of parent peptide: YGGAGRGLGK{acr}), (A) "Tyrosine fragment" (YGGAGR, abbreviated: "Tyr-fragment") resulting from trypsin cleavage, (B) modification of the fragment in (A) with known tyrosine oxidation products, notably "3,4-Dihydroxyphenlylanine (DOPA) and/or dopamine quinone", to form "Tyrosine Oxidation products 1 fragment" (abbreviated: "TyrOx1-fragment"), (C) modification of the fragment in (A) with known tyrosine oxidation products, notably "Dityrosine fragment" (abbreviated: "TyrOx2-fragment") [3].

**ii)** Table detailing the exact mass, and absolute abundance recorded for the different products (from A-C peptide fragments from **Supplementary Figure 8ci**) in the different conditions (Condition 1: in solution with trypsin; Condition 2: in solution with trypsin, APS and TEMED; Condition 3: ExM gel trypsinized peptide). AUC for each species in the extracted ion chromatogram was obtained as in **Methods: Analysis of LC/QToF data** and the raw total ion chromatograms (TIC) located in **Source Data**. The dash ('-') means the species was not detected when using search parameters (the extracted ion chromatogram was generated using the $[M+H]^+$ ion of the species, with a mass tolerance of ±20 ppm, with all detected peaks integrated into the final value).
**iii)** Normalized abundance of tyrosine N-terminal peptide and oxidation products in different conditions. Normalized abundance of the 3 different species (Tyr-fragment, TyrOx1-fragment, TyrOx2-fragment; as detailed in **Supplementary Figure 8ci**) in: condition 1, after trypsin cleavage in solution (left); condition 2, after addition of APS and TEMED with trypsin cleavage in solution (middle); and condition 3, after trypsin cleavage of peptides from ExM gels (condition 3; right). AUC for each species in the extracted ion chromatogram was obtained as in **Methods: Analysis of LC/QToF data** and the raw total ion chromatograms (TIC) located in **Source Data**. The abundance of each species was normalized with max absolute scaling for plotting on the bar graph (n=1 gelation solution for each condition).

**d)**

**i)**

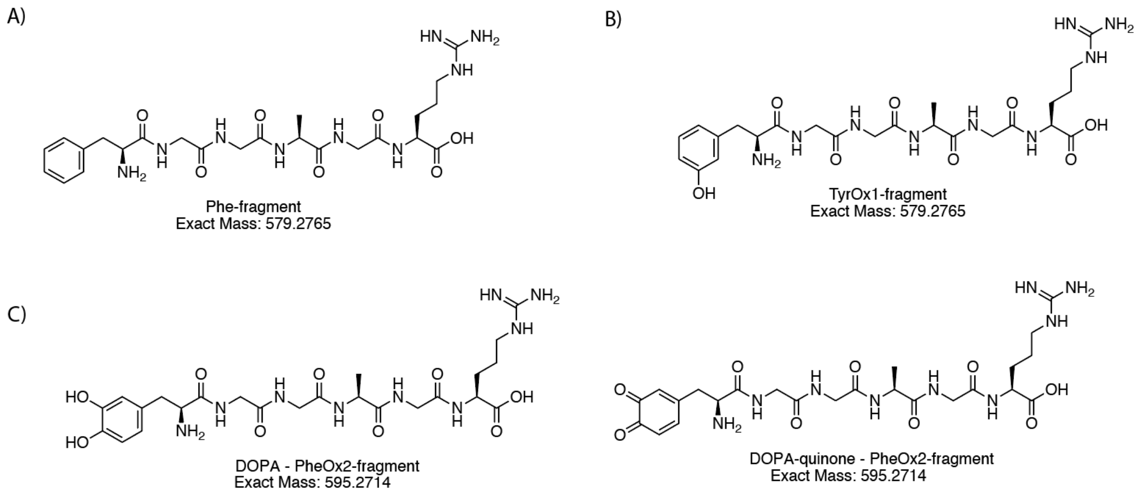

**ii)**

| **Phenylalanine** | A) Phe-fragment | B) PheOx1-fragment | C) PheOx2-fragment |
|---|---|---|---|
| *Exact mass* | *563.2816* | *579.2765* | *595.2714* |
| Condition 1: trypsin | 39,692,812 | 230.34 | 502.13 |
| Condition 2: TEMED+APS+trypsin | 27,486,226 | 128,252.7 | 2,134.02 |
| Condition 3: TEMED+APS+trypsin in ExM gel | 18,738,042 | 1,166.65 | - |

**iii)**

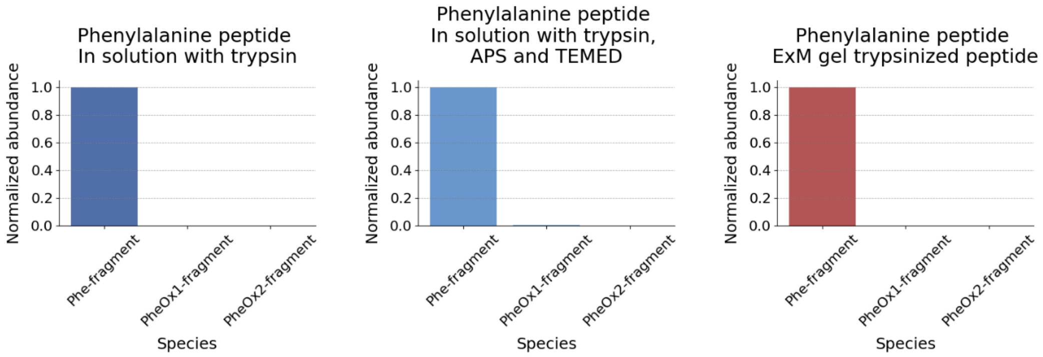

**Supplementary Figure 8d** Phenylalanine and post-translational oxidation
**i)** Structure of the various fragments resulting from enzymatic cleavage with trypsin from the parent peptide (sequence of parent peptide: FGGAGRGLGK{acr}), (A) "Phenylalanine fragment" (FGGAGR, abbreviated: "Phe-fragment") resulting from trypsin cleavage, (B) modification of the fragment in (A) with known phenylalanine oxidation products, notably "Meta- or ortho-tyrosine", to form "Oxidation products 1 fragment" (abbreviated: "PheOx1-fragment"), (C) modification of the fragment in (A) with known phenylalanine oxidation products, notably "Dihydroxyphenylalanine (DOPA) and/or dopamine quinone fragment" (abbreviated: "PheOx2-fragment") [4].
**ii)** Table detailing the exact mass, and absolute abundance recorded for the different products (from A-C peptide fragments from **Supplementary Figure 8di**) in the different conditions (Condition 1: in solution with trypsin; Condition 2: in solution with trypsin, APS and TEMED; Condition 3: ExM gel trypsinized peptide). AUC for each species in the extracted ion chromatogram was obtained as in **Methods: Analysis of LC/QToF data** and the raw total ion chromatograms (TIC) located in **Source Data**. The dash ('-') means the species was not detected when using search parameters (the extracted

ion chromatogram was generated using the $[M+H]^+$ ion of the species, with a mass tolerance of ±20 ppm, with all detected peaks integrated into the final value).
**iii )** Normalized abundance of phenylalanine N-terminal peptide and oxidation products in different conditions. Normalized abundance of the 3 different species (Phe-fragment, PheOx1-fragment, PheOx2-fragment; as detailed in **Supplementary Figure 8di**) in: condition 1, after trypsin cleavage in solution (left); condition 2, after addition of APS and TEMED with trypsin cleavage in solution (middle); and condition 3, after trypsin cleavage of peptides from ExM gels (condition 3; right). AUC for each species in the extracted ion chromatogram was obtained as in **Methods: Analysis of LC/QToF data** and the raw total ion chromatograms (TIC) located in **Source Data**. The abundance of each species was normalized with max absolute scaling for plotting on the bar graph (n=1 gelation solution for each condition).

**e)**

**i)**

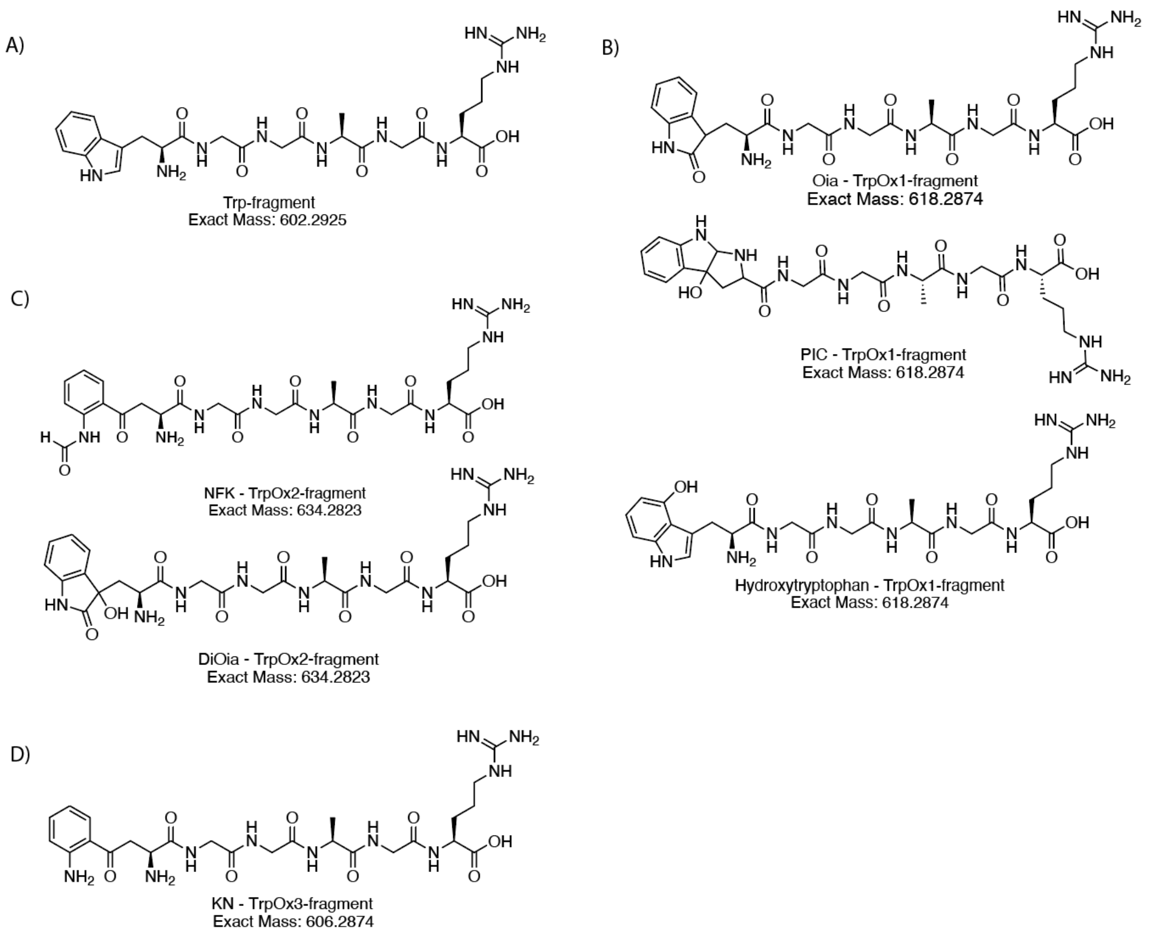

**ii)**

| **Tryptophan** | A) Trp-fragment | B) TrpOx1-fragment | C) TrpOx2-fragment | D) TrpOx3-fragment |
|---|---|---|---|---|
| *Exact mass* | *602.2925* | *618.2874* | *634.2823* | *606.2874* |
| Condition 1: trypsin | 37,058,997 | 91,994.91 | 9,823.04 | 20,127.28 |
| Condition 2: TEMED+APS+trypsin | 8,511,649.4 | 1,601,600 | 582,627 | 107,597.4 |
| Condition 3: TEMED+APS+trypsin in ExM gel | 2,119,325.3 | 108,509.7 | 97,603.25 | 11,329.72 |

**iii)**

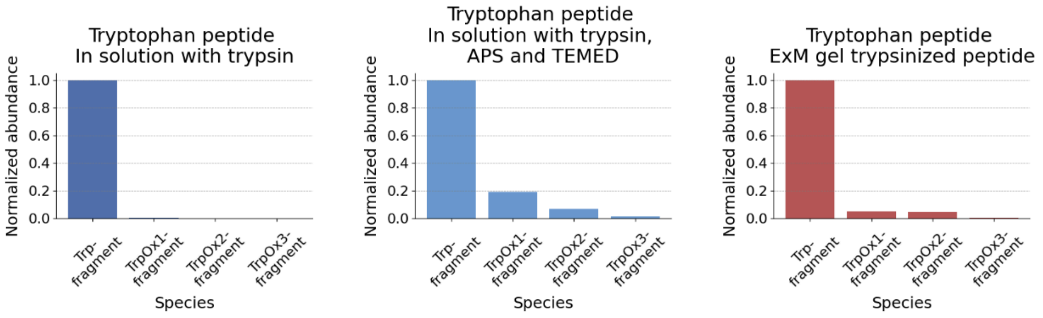

**Supplementary Figure 8e Tryptophan and post-translational oxidation.**

**i)** Structure of the various fragments resulting from enzymatic cleavage with trypsin from the parent peptide (sequence of parent peptide: WGGAGRGLGK{acr}), (A) "Tryptophan fragment"

(WGGAGR, abbreviated: “Trp-fragment”) resulting from trypsin cleavage, (B) modification of the fragment in (A) with known tryptophan oxidation products, notably “Oxindolylalanine (Oia), and/or 3-Hydroxypyrroloindole carboxylic acid (PIC), and/or hydroxytryptophan”, to form “Oxidation products 1 fragment” (abbreviated: “TrpOx1-fragment”), (C) modification of the fragment in (A) with known tryptophan oxidation products, notably “N-formylkynurenine (NFK) and/or dioxindolylalanine (DiOia)” (abbreviated: “TrpOx2-fragment”), (D) modification of the fragment in (A) with known tryptophan oxidation products, notably “Kynurenin (KN)” (abbreviated: “TrpOx3-fragment”) [5].
**ii)** Table detailing the exact mass, and absolute abundance recorded for the different products (from a-c peptide fragments from **Supplementary Figure 8ei**) in the different conditions (Condition 1: in solution with trypsin; Condition 2: in solution with trypsin, APS and TEMED; Condition 3: ExM gel trypsinized peptide). AUC for each species in the extracted ion chromatogram was obtained as in **Methods: Analysis of LC/QToF data** and the raw total ion chromatograms (TIC) located in **Source Data**. The dash (‘-’) means the species was not detected when using search parameters (the extracted ion chromatogram was generated using the $[M+H]^+$ ion of the species, with a mass tolerance of ±20 ppm, with all detected peaks integrated into the final value).
**iii)** Normalized abundance of tryptophan N-terminal peptide and oxidation products in different conditions. Normalized abundance of the 4 different species (Trp-fragment, TrpOx1-fragment, TrpOx2-fragment, TrpOx3-fragment; as detailed in **Supplementary Figure 8di**) in: condition 1, after trypsin cleavage in solution (left); condition 2, after addition of APS and TEMED with trypsin cleavage in solution (middle); and condition 3, after trypsin cleavage of peptides from ExM gels (condition 3; right). AUC for each species in the extracted ion chromatogram was obtained as in **Methods: Analysis of LC/QToF data** and the raw total ion chromatograms (TIC) located in **Source Data**. The abundance of each species was normalized with max absolute scaling for plotting on the bar graph (n=1 gelation solution for each condition).

**f)**

**i)** A)

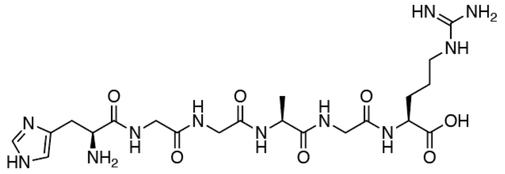

His-fragment
Exact Mass: 553.2721

B)

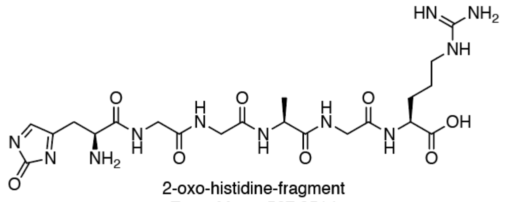

2-oxo-histidine-fragment
Exact Mass: 567.2514

C)

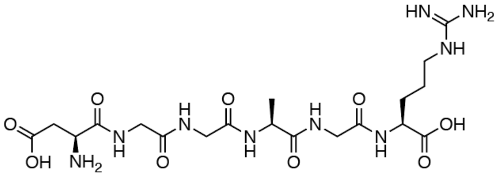

Aspartate-fragment
Exact Mass: 531.2401

D)

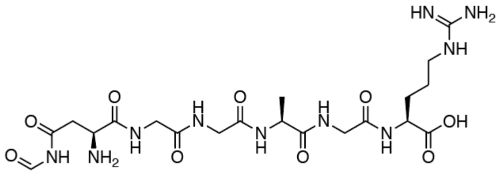

Formylasparagine-fragment
Exact Mass: 558.2510

E)

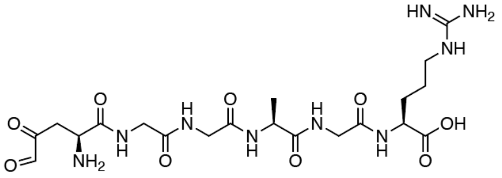

Aspartylurea-fragment
Exact Mass: 543.2401

**ii)**

| **Histidine** | A) His-fragment | B) 2-oxo-histidine-fragment | C) Aspartate-fragment | D) Formylasparagine-fragment | E) Aspartylurea-fragment |
|---|---|---|---|---|---|
| *Exact mass* | *553.2721* | *567.2514* | *531.2401* | *558.251* | *543.2401* |
| Condition 1: trypsin | 24,178,48.5 | 442.7 | 9,000.49 | 33,615.51 | 235.33 |
| Condition 2: TEMED+APS+trypsin | 1828127.9 | 22,679.29 | 44,921.53 | 19,516.84 | 28,212.79 |
| Condition 3: TEMED+APS+trypsin in ExM gel | 441,471.91 | - | 4,632.48 | 5,975.27 | 30,999.35 |

**iii)**

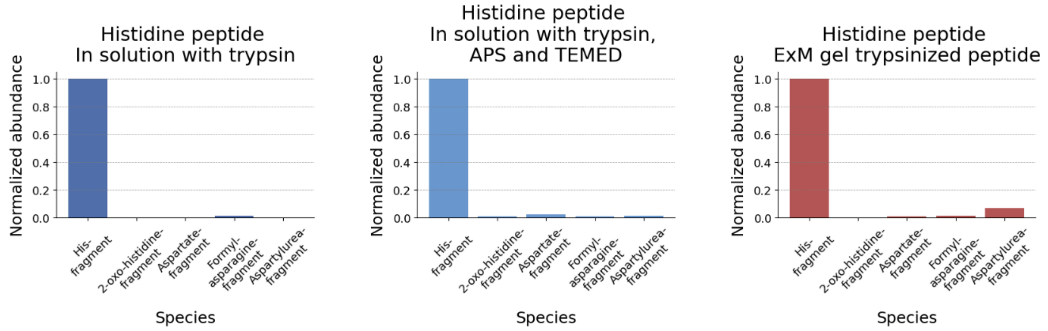

**Supplementary Figure 8f** Histidine and post-translational oxidation.

**i)** Structure of the various fragments resulting from enzymatic cleavage with trypsin from the parent peptide (sequence of parent peptide: HGGAGRGLGK{acr}), (A) "Histidine fragment" (HGGAGR, abbreviated: "His-fragment") resulting from trypsin cleavage, (B) modification of the fragment in (A) with known histidine oxidation products, notably 2-oxo-histidine, to form "2-oxo-histidine fragment" (abbreviated: "2-oxo-histidine-fragment"), (C) modification of the fragment in (A) with known histidine oxidation products, notably "Aspartate fragment" (abbreviated: "Aspartate-fragment"), (D) modification of the fragment in (A) with known histidine oxidation products, notably

“Formylasparagine fragment” (abbreviated: “Formylasparagine-fragment”), (E) modification of the fragment in (A) with known histidine oxidation products, notably “Aspartylurea fragment” (abbreviated: “Aspartylurea-fragment”) [6,7].

**ii)** Table detailing the exact mass, and absolute abundance recorded for the different products (from A-E peptide fragments from **Supplementary Figure 8fi**) in the different conditions (Condition 1: in solution with trypsin; Condition 2: in solution with trypsin, APS and TEMED; Condition 3: ExM gel trypsinized peptide). AUC for each species in the extracted ion chromatogram was obtained as in **Methods: Analysis of LC/QToF data** and the raw total ion chromatograms (TIC) located in **Source Data**. The dash (‘-’) means the species was not detected when using search parameters (the extracted ion chromatogram was generated using the $[M+H]^+$ ion of the species, with a mass tolerance of ±20 ppm, with all detected peaks integrated into the final value).

**iii)** Normalized abundance of histidine N-terminal peptide and oxidation products in different conditions. Normalized abundance of the 5 different species (His-fragment, 2-oxo-histidine-fragment, Aspartate-fragment, Formylasparagine-fragment, Aspartylurea-fragment; as detailed in **Supplementary Figure 8fi**) in: condition 1, after trypsin cleavage in solution (left); condition 2, after addition of APS and TEMED with trypsin cleavage in solution (middle); and condition 3, after trypsin cleavage of peptides from ExM gels (condition 3; right). AUC for each species in the extracted ion chromatogram was obtained as in **Methods: Analysis of LC/QToF data** and the raw total ion chromatograms (TIC) located in **Source Data**. The abundance of each species was normalized with max absolute scaling for plotting on the bar graph (n=1 gelation solution for each condition).

**g)**

**i)**

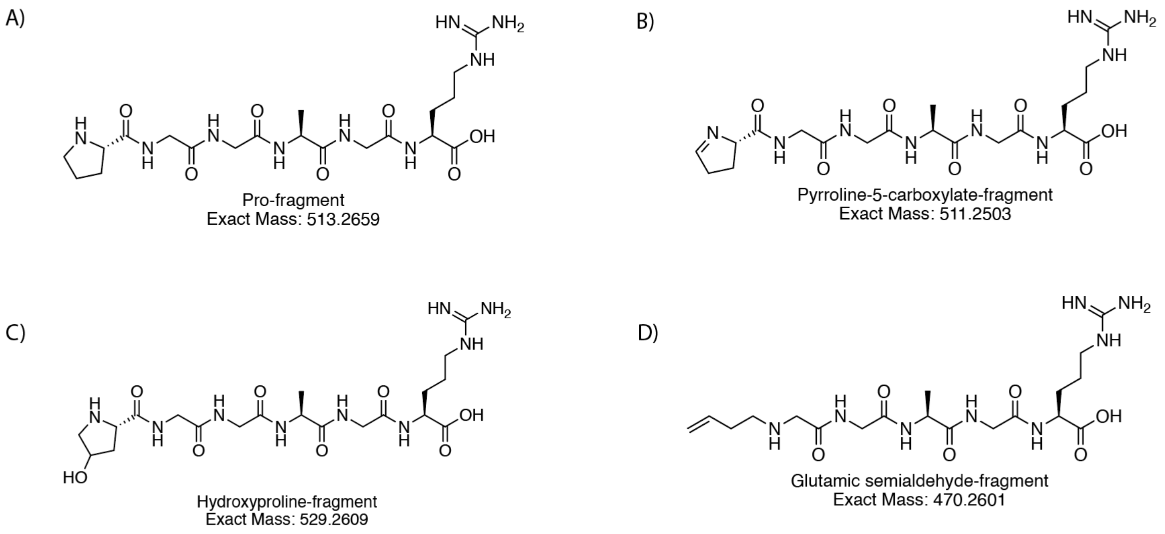

**ii)**

| **Proline** | A) Pro-fragment | B) Pyrroline-5-carboxylate-fragment | C) Hydroxyproline-fragment | D) Glutamic semialdehyde-fragment |
|---|---|---|---|---|
| *Exact mass* | *513.2659* | *511.2503* | *529.2609* | *527.2816* |
| Condition 1: trypsin | 37,903,603 | 371.23 | 6,882.07 | 5,223.48 |
| Condition 2: TEMED+APS+trypsin | 9,000,359.9 | 544,009.1 | 23,130.87 | 4,143.66 |
| Condition 3: TEMED+APS+trypsin in ExM gel | 5,514,152.5 | 294,169.1 | 3,102.69 | 1,426.29 |

**iii)**

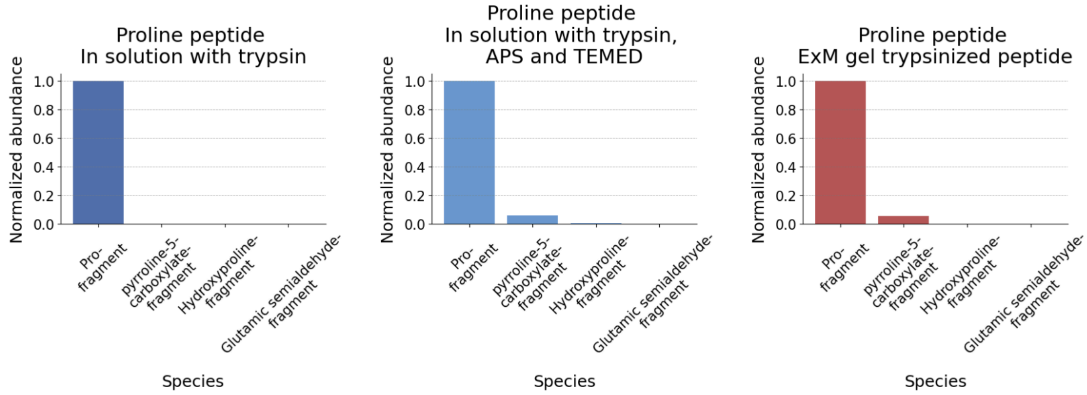

**Supplementary Figure 8g** Proline and post-translational oxidation.

**i)** Structure of the various fragments resulting from enzymatic cleavage with trypsin from the parent peptide (sequence of parent peptide: PGGAGRGLGK{acr}), (A) “Proline fragment” (sequence: PGGAGR, abbreviated: “Pro-fragment”) resulting from trypsin cleavage, (B) modification of the fragment in (A) with known proline oxidation products, notably “Pyrroline-5-carboxylate”, to form “Pyrroline-5-carboxylate fragment” (abbreviated: “Pyrroline-5-carboxylate-fragment”), (C) modification of the fragment in (A) with known proline oxidation products, notably “Hydroxyproline” (abbreviated: “Hydroxyproline-fragment”), (D) modification of the fragment in (A) with known proline oxidation products, notably “Glutamic semialdehyde” (abbreviated: “Glutamic semialdehyde-fragment”) [8].

**ii)** Table detailing the exact mass, and absolute abundance recorded for the different products (from A-D peptide fragments from **Supplementary Figure 8gi**) in the different conditions (Condition 1: in solution with trypsin; Condition 2: in solution with trypsin, APS and TEMED; Condition 3: ExM gel trypsinized peptide). AUC for each species in the extracted ion chromatogram was obtained as in **Methods: Analysis of LC/QToF data** and the raw total ion chromatograms (TIC) located in **Source Data**. The dash ('-') means the species was not detected when using search parameters (the extracted ion chromatogram was generated using the $[M+H]^+$ ion of the species, with a mass tolerance of ±20 ppm, with all detected peaks integrated into the final value).
**iii)** Normalized abundance of proline N-terminal peptide and oxidation products in different conditions. Normalized abundance of the 4 different species (Pro-fragment, Pyrroline-5-carboxylate-fragment, Hydroxyproline-fragment, Glutamic semialdehyde-fragment; as detailed in **Supplementary Figure 8gi**) in: condition 1, after trypsin cleavage in solution (left); condition 2, after addition of APS and TEMED with trypsin cleavage in solution (middle); and condition 3, after trypsin cleavage of peptides from ExM gels (condition 3; right). AUC for each species in the extracted ion chromatogram was obtained as in **Methods: Analysis of LC/QToF data** and the raw total ion chromatograms (TIC) located in **Source Data**. The abundance of each species was normalized with max absolute scaling for plotting on the bar graph (n=1 gelation solution for each condition).

**h)**

**i)**

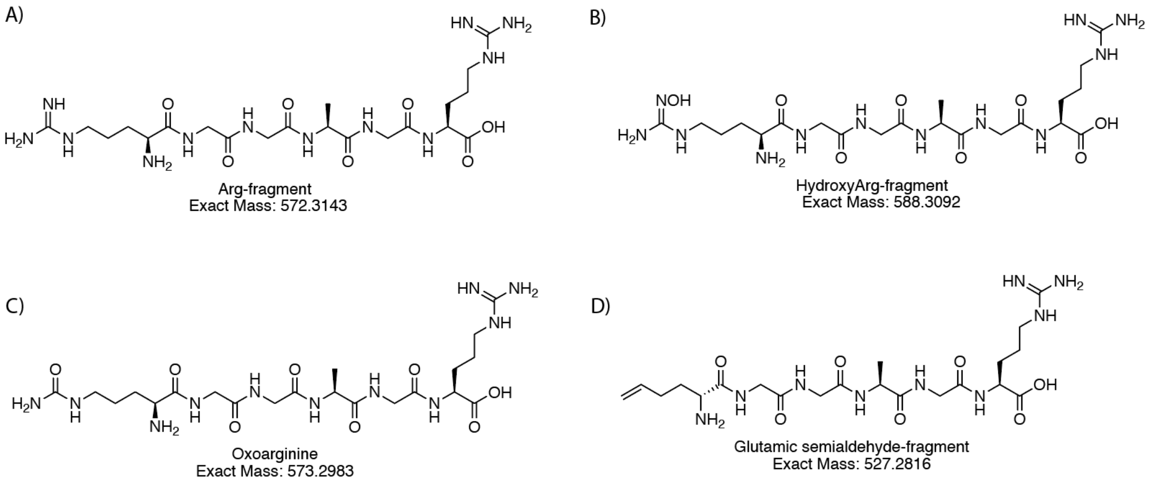

**ii)**

| **Arginine** | A) Arg-fragment | B) HydroxyArg-fragment | C) OxoArg-fragment | D) Glutamic semialdehyde-fragment |
|---|---|---|---|---|
| *Exact mass* | *572.3143* | *588.3092* | *573.2983* | *527.2816* |
| Condition 1: trypsin | 17,883,221 | 94,152.55 | 569.49 | 821.92 |
| Condition 2: TEMED+APS+trypsin | 7,767,661.7 | 51,065.82 | 13,029.76 | 1,481.44 |
| Condition 3: TEMED+APS+trypsin in ExM gel | 4,906,487.6 | 6,285.23 | 2,591.88 | 1,105.52 |

**iii)**

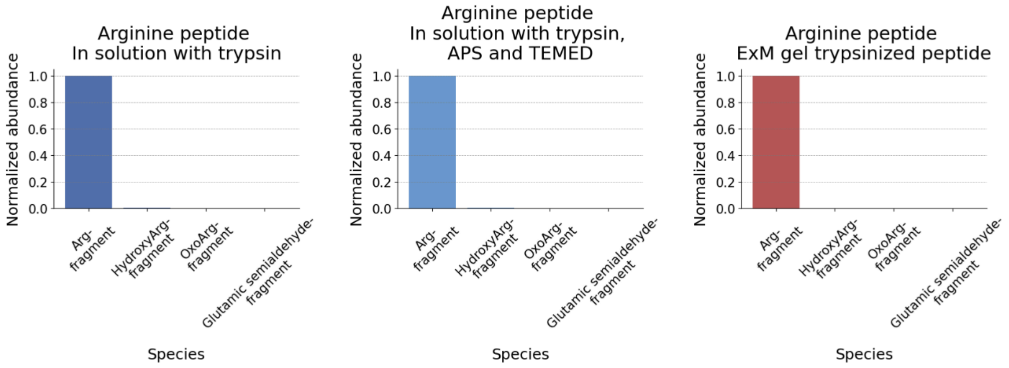

**Supplementary Figure 8h** Arginine and post-translational oxidation.

**i)** Structure of the various fragments resulting from enzymatic cleavage with trypsin from the parent peptide (sequence of parent peptide: RGGAGRGLGK{acr}), (A) "Arginine fragment" (RGGAGR, abbreviated: "Arg-fragment") resulting from trypsin cleavage, (B) modification of the fragment in (A) with known arginine oxidation products, notably "Hydroxyarginine", to form "Hydroxyarginine fragment" (abbreviated: "HydroxyArg-fragment"), C) modification of the fragment in (A) with known arginine oxidation products, notably "Oxoarginine" (abbreviated: "OxoArg-fragment"), (D) modification of the fragment in (A) with known arginine oxidation products, notably "Glutamic semialdehyde" (abbreviated: "Glutamic semialdehyde-fragment") [9,10].

**ii)** Table detailing the exact mass, and absolute abundance recorded for the different products (from A-D peptide fragments from **Supplementary Figure 3hi**) in the different conditions (Condition 1: in solution with trypsin; Condition 2: in solution with trypsin, APS and TEMED; Condition 3: ExM gel trypsinized peptide). AUC for each species in the extracted ion chromatogram was obtained as in **Methods: Analysis of LC/QToF data** and the raw total ion chromatograms (TIC) located in **Source Data**. The dash ('-') means the species was not detected when using search parameters (the extracted

ion chromatogram was generated using the $[M+H]^+$ ion of the species, with a mass tolerance of ±20 ppm, with all detected peaks integrated into the final value).
**iii)** Normalized abundance of arginine N-terminal peptide and oxidation products in different conditions. Normalized abundance of the 4 different species (Arg-fragment, HydroxyArg-fragment, OxoArg-fragment, Glutamic semialdehyde-fragment; as detailed in **Supplementary Fig. 3hi**) in: condition 1, after trypsin cleavage in solution (left); condition 2, after addition of APS and TEMED with trypsin cleavage in solution (middle); and condition 3, after trypsin cleavage of peptides from ExM gels (condition 3; right). AUC for each species in the extracted ion chromatogram was obtained as in **Methods: Analysis of LC/QToF data** and the raw total ion chromatograms (TIC) located in **Source Data**. The abundance of each species was normalized with max absolute scaling for plotting on the bar graph (n=1 gelation solution for each condition).

#### Supplementary Figure 9

A

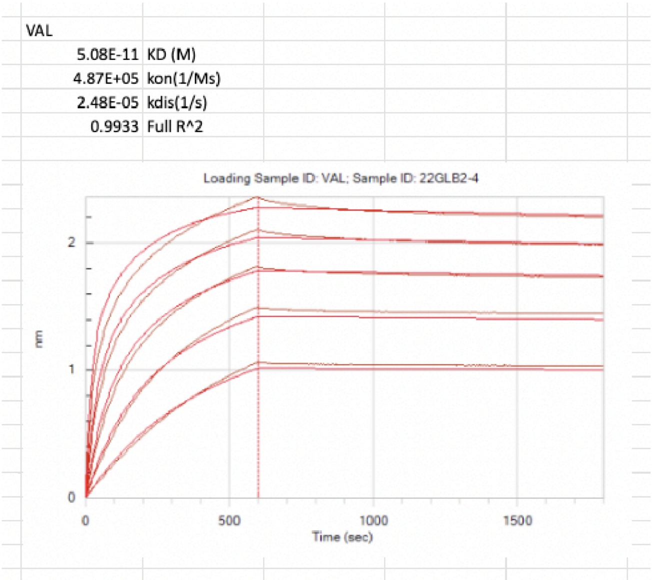

B

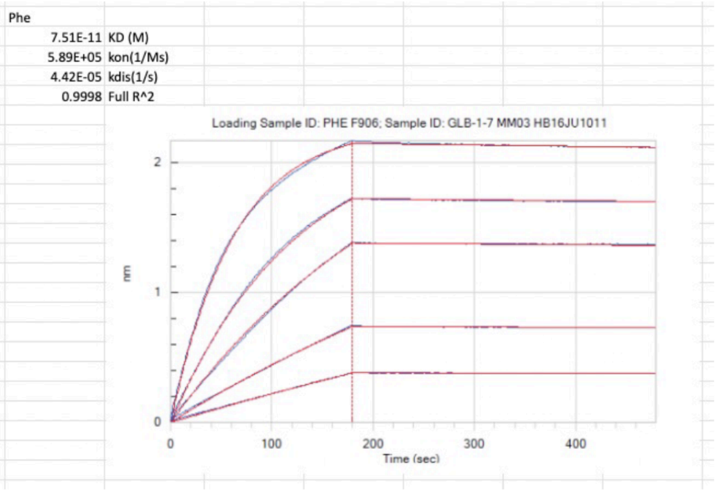

C

| | $K_d$ | $k_{on}$ | $k_{dis}$ |
|---|---|---|---|
| Glyphic K{ac} | 1.5 x $10^{-9}$ M | 3.45 x $10^5$ ($M^{-1}s^{-1}$) | 5.15 x $10^{-4}$ ($s^{-1}$) |
| Glyphic Ile | 5.85 x $10^{-12}$ M | 1.71 x $10^5$ ($M^{-1}s^{-1}$) | 1.00 x $10^{-6}$ ($s^{-1}$) |
| Glyphic Leu | 1.73 x $10^{-10}$ M | 6.73 x $10^5$ ($M^{-1}s^{-1}$) | 1.16 x $10^{-6}$ ($s^{-1}$) |

**Supplementary Figure 9:** Biolayer interferometry (BLI) curves for GFP-conjugated Glyphic antibody against ClickP-V (abbreviated: Glyphic V) and RFP-conjugated Glyphic antibody against ClickP-F (abbreviated: Glyphic F). (A) BLI curve obtained for Glyphic V against ClickP-valine antigen using five different antibody concentrations. The global fit was obtained using a bivalent fitting model and shows a dissociation constant of $K_d$ = 50.8 pM, a $k_{on}$ = 4.87 * $10^5$ $M^{-1}s^{-1}$, a $k_{dis}$ = 2.48 * $10^{-5}$ $s^{-1}$. (B) As in A, but for Glyphic F. $K_d$ = 75.1 pM, a $k_{on}$ = 5.89 * $10^5$ $M^{-1}s^{-1}$, a $k_{dis}$ = 4.42 * $10^{-5}$ $s^{-1}$. (C) Table showing the binding kinetic results for Glyphic antibodies against ClickP-acetyl-lysine (abbreviated: Glyphic K{ac}), ClickP-isoleucine (abbreviated: Glyphic Ile) and ClickP-leucine (Glyphic Leu).

#### Supplementary Figure 10

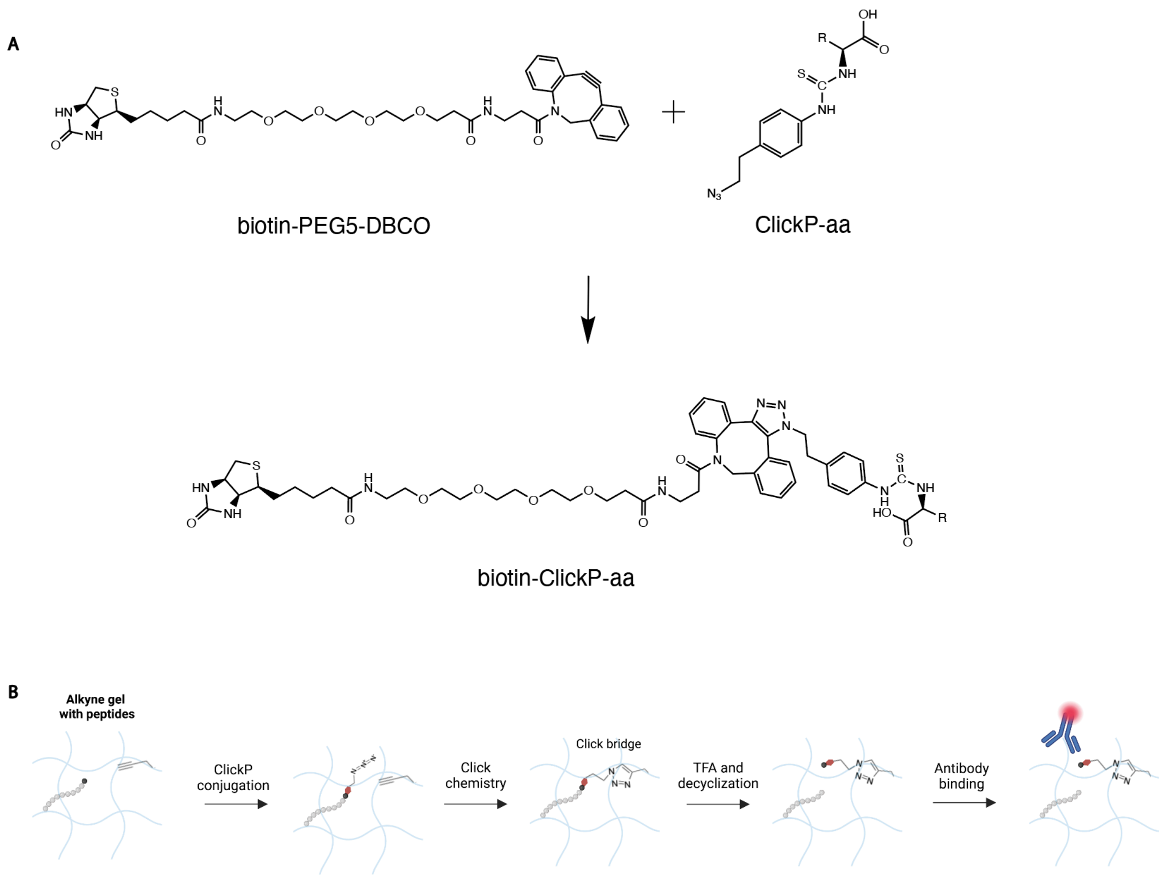

**Supplementary Figure 10:**

(A) Reaction of biotin-PEG5-DBCO (top left) with decyclized ClickP-amino acid (abbreviated: ClickP-aa) (top right) to form biotin-ClickP-aa (bottom). Biotin-ClickP-aa is the reagent used in **Figure 6D-I** for assessing specificity of binders in the ExMre gels. "R" in this case represents the variable group of the amino acid side chain.

(B) Conceptual extension to **Figure 6A**: peptides cast in an ExMre hydrogel conjugated with “clickable” groups (denoted: “Clickable gel”, e.g. with alkyne groups for copper-catalyzed chemistry). The bifunctional molecule, with an azide and isothiocyanate group, first reacts to primary amines at the N-terminus of peptides via the isothiocyanate group. Then, the copper-catalyzed click chemistry forms a bridge between the ClickP-conjugated N-terminus (4-(2-azidoethyl)phenylthiocarbamoyl-peptide) via the azide group with the alkyne group in the gel. TFA is then used to cleave the N-terminal amino acid from the peptide, and the covalent click reaction locally tethering the amino acid to the surrounding hydrogel network, prevents its diffusion out of the gel. The N-terminal amino acid is decyclized from the thiohydantoin or thiozolinone intermediates, from post-cleavage, into the thiocarbamoyl derivative with covalently tethered N-terminus and an available C-terminus (carboxyl group).

#### Supplementary Figure 11

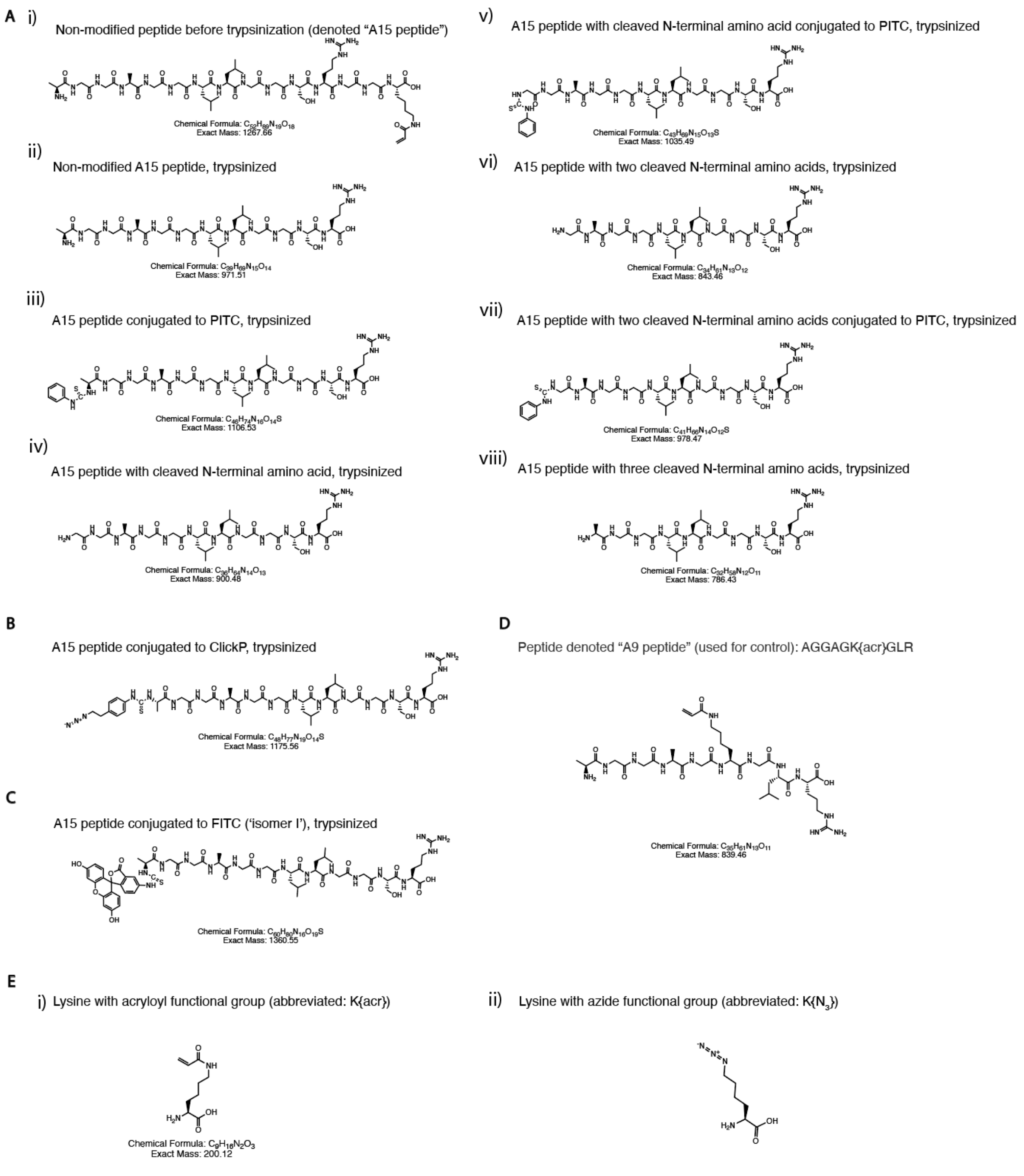

**Supplementary Figure 11:** Structure of various peptides (chemical formula and exact mass also detailed under each structure).

(A) (i) Structure of the “A15” peptide (i.e., AGGAGGLLGGSRGGK{acr}), (ii-viii) are structures of this peptide once it has been trypsinized from the gel (cleavage at arginine, R), (ii) the non-modified fragment trypsinized from the gel AGGAGGLLGGSR, (iii) the same fragment as (ii) but after conjugation with PITC, (i.e., *PTC*-AGGAGGLLGGSR), (iv) the same fragment as (ii) but with cleaved N-terminal amino acid (i.e., GGAGGLLGGSR), (v) the same fragment as (iv) but after conjugation with PITC (i.e., *PTC*-GGAGGLLGGSR), (vi) the same fragment as (ii) but with two cleaved N-terminal amino acids (i.e., GAGGLLGGSR),

(vii) the same fragment as (vi) but after conjugation with PITC (i.e., *PTC*-GAGGLLGGSR), (viii) the same fragment as (ii) but with three cleaved N-terminal amino acids (i.e., AGGLLGGSR).

(B) Structure of the same fragment as (ii) but after conjugation with ClickP (i.e., *ClickP*-AGGAGGLLGGSR).

(C) Structure of the same fragment as (ii) but after conjugation with FITC ('Isomer I') (*FITC*-AGGAGGLLGGSR).

(D) Structure of peptide denoted A9 peptide, used as ionization efficiency standard, with sequence AGGAGK{acr}GLR, where the acryloyl group on the side chain of lysine is abbreviated K{acr}.

(E) (i) Structure of the lysine amino acid modified with acryloyl group (K{acr}), (ii) structure of the lysine amino acid modified with azide group (abbreviated: K{$N_3$}).

#### Supplementary Figure 12

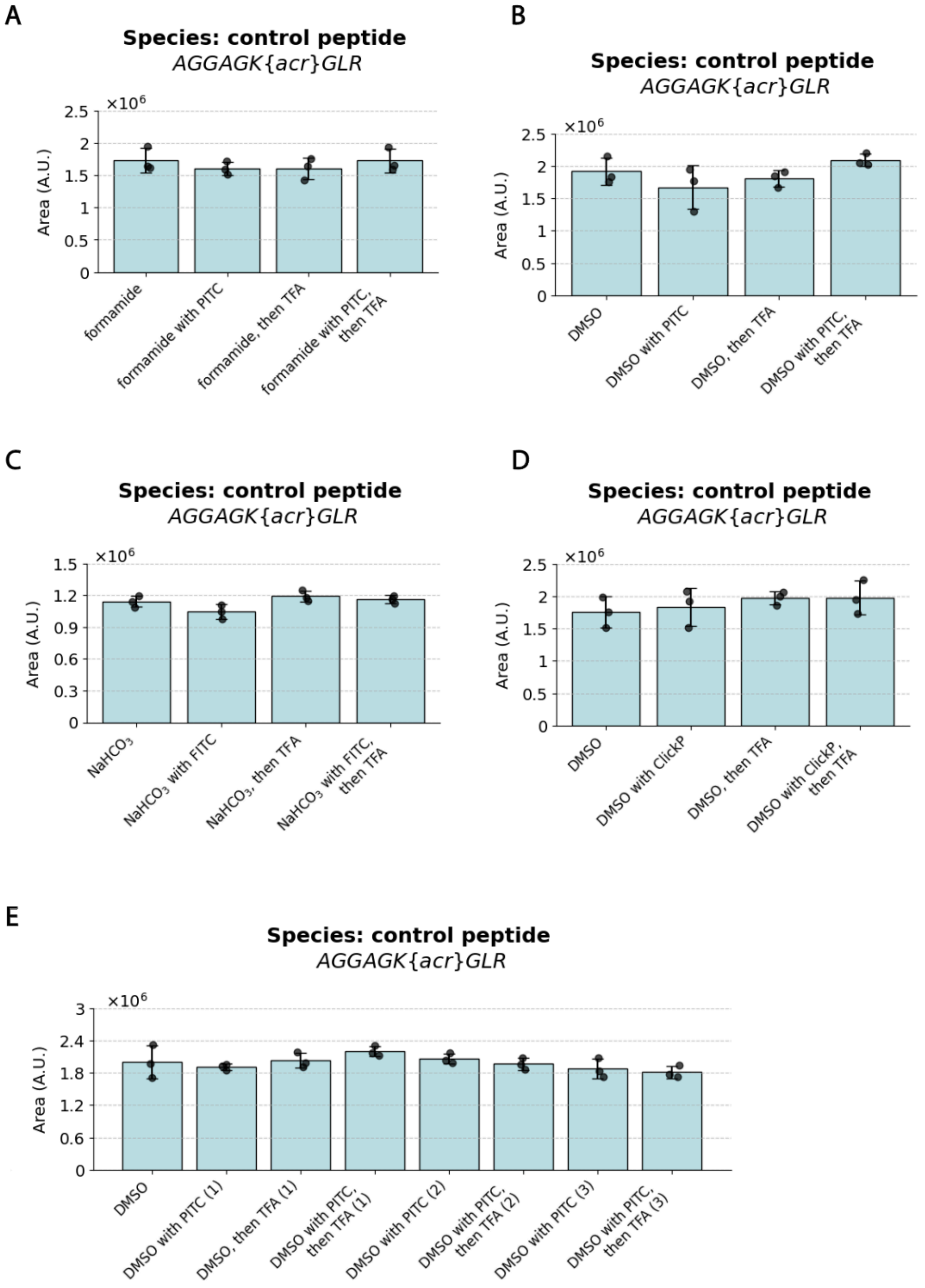

**Supplementary Figure 12:** Bar graphs representing the relative abundance (arbitrary units, a.u.) of A9-peptide (AGGAGK{acr}GLR) ion species detected on the LC/QToF. The bar graphs are obtained from comparing the area under the curve (AUC) of the chromatogram for the A9 peptide. It was spiked into the supernatant of the trypsinized gels, as an ionization efficiency standard, for downstream LC/QToF analysis in different in-gel Edman degradation conditions. A9 peptide abundance after the gel-anchored peptide (*not* the A9) was treated with: (A) PITC to formamide (1:1000 ratio PITC:formamide) conjugation solvent (as well as solvent only, TFA, and PITC followed by TFA; Figure 3**B**), (B) PITC to DMSO (1:1000 ratio PITC:DMSO) conjugation solvent (and related conditions, as in A; Figure 3**C**), (C) FITC at concentration 5.9 mM with 23:77 DMSO:0.1 M sodium bicarbonate ($NaHCO_3$) pH 8.5 (and related conditions, as in A; Figure 3**D**) (D) ClickP to DMSO

(1:1000 ratio ClickP:DMSO) as conjugation solvent (and related conditions, as in A; **Figure 6C**), (E) PITC to DMSO (1:1000 ratio PITC:DMSO) as conjugation solvent, over 3 rounds of in-gel Edman degradation (and related conditions, as in A; **Figure 5B**). Black dots, individual experiments; blue bar, mean; error bar: standard deviation; n=3 gelation solutions.

#### Supplementary Figure 13

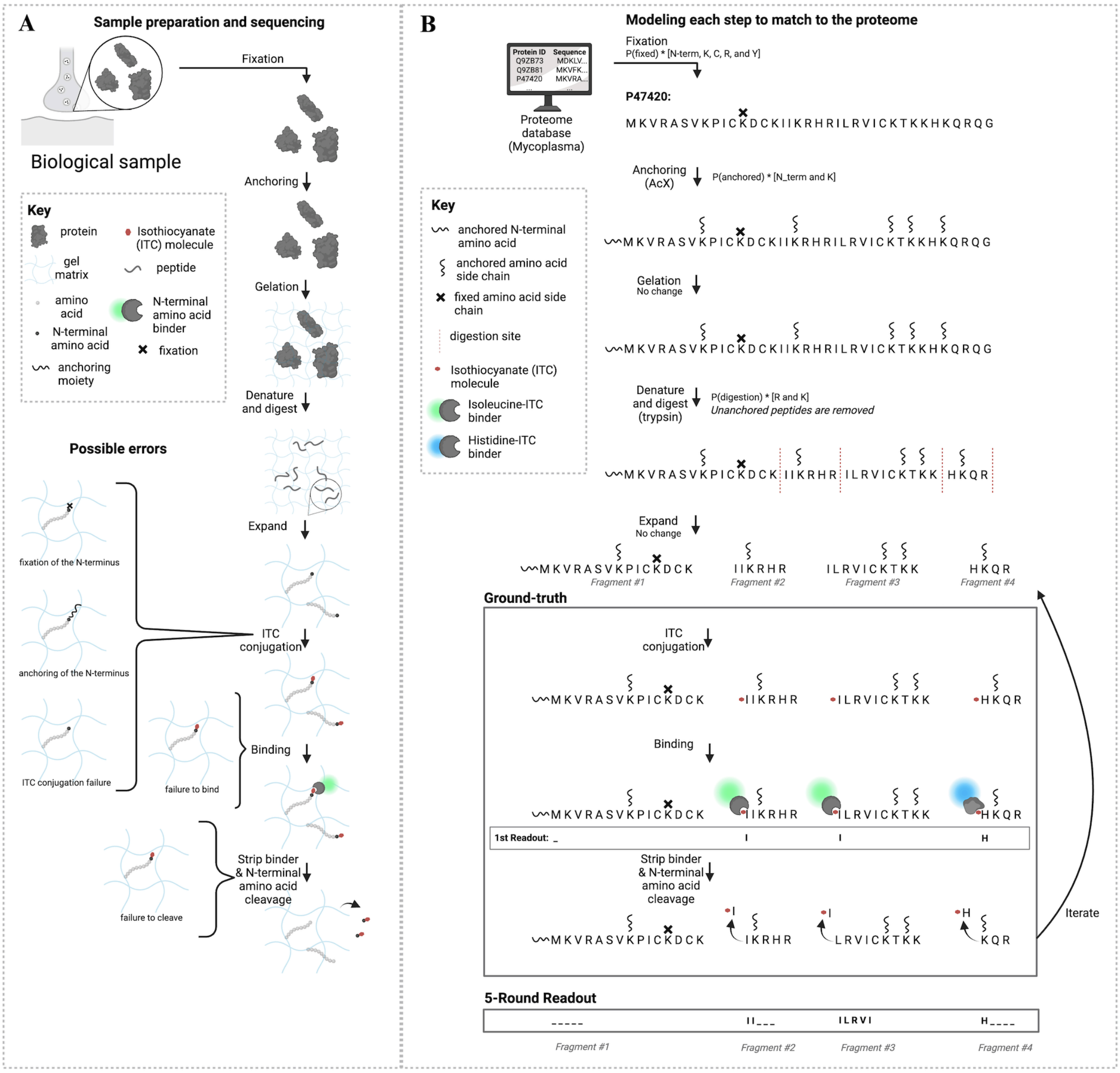

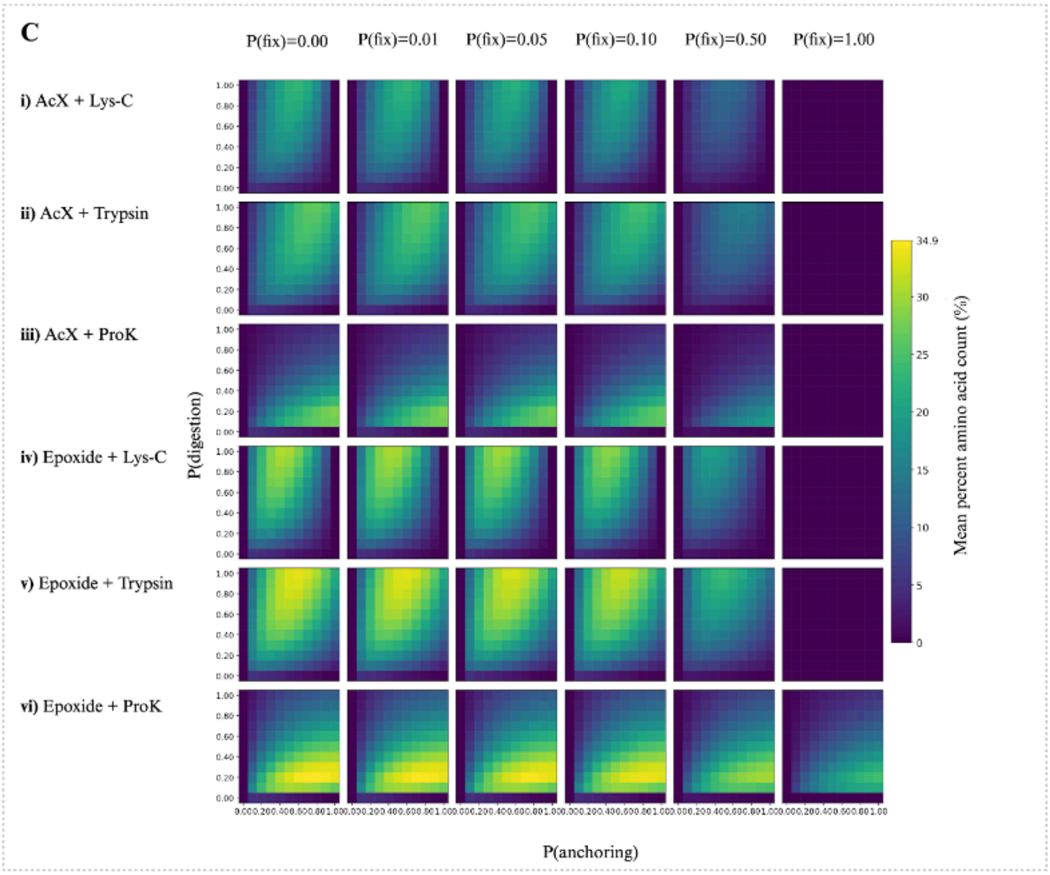
C
P(fix)=0.00
P(fix)=0.01
P(fix)=0.05
P(fix)=0.10
P(fix)=0.50
P(fix)=1.00
i) AcX + Lys-C
ii) AcX + Trypsin
iii) AcX + ProK
iv) Epoxide + Lys-C
v) Epoxide + Trypsin
vi) Epoxide + ProK
P(digestion)
P(anchoring)
Mean percent amino acid count (%)
34.9
30
25
20
15
10
5
0

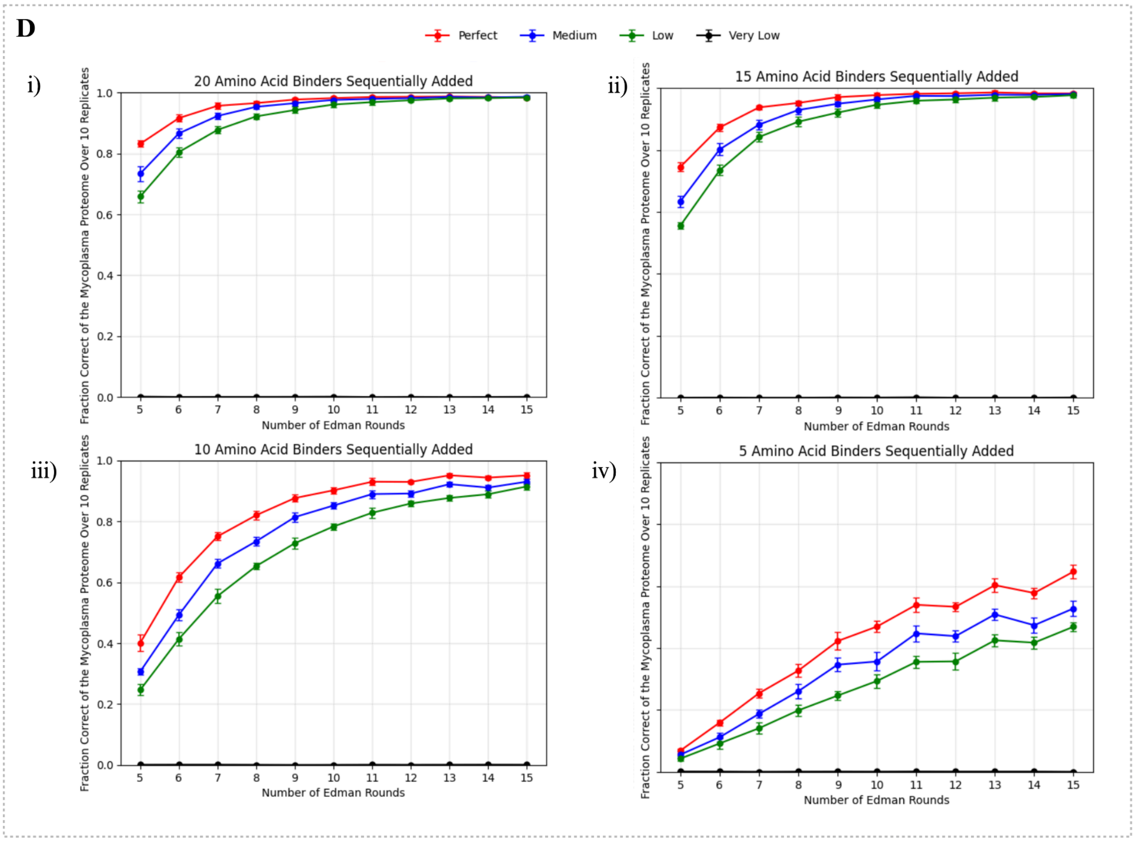
D
Perfect
Medium
Low
Very Low
i)
20 Amino Acid Binders Sequentially Added
ii)
15 Amino Acid Binders Sequentially Added
iii)
10 Amino Acid Binders Sequentially Added
iv)
5 Amino Acid Binders Sequentially Added
Fraction Correct of the Mycoplasma Proteome Over 10 Replicates
Number of Edman Rounds

**Supplementary Figure 13: A theoretical assessment of in situ protein sequencing.**

(A) Schematic of the experimental workflow being modeled. Starting from the biological sample, the chemical steps include fixation, anchoring, gelation, denature/digestion, expanding, and the sequencing chemistry. Major potential sequencing chemistry errors are depicted, including during fixation or anchoring of the N-terminal amino acid, ITC conjugation failure, failure for the NAAB to bind, and/or failure to cleave the N-terminal amino acid.

(B) Schematic of proteins in the pipeline of the computational model. Starting from modeling the upstream preparation steps, where fixation step modifies N-terminal (N-term), K, C, R, Y with probability P(fixed); the anchoring step that modifies N-term and K with probability P(anchored); gelation which does not lead to sequence modification; denature/digest step that cleaves peptides at R and K residues to expose new N-termini; and expansion which does not modify the sequences. Sequencing chemistry is performed with sequential addition of ITC to the N-termini of the peptide fragments, followed by NAAB binding and read-outs, and iteration of these two previous steps. The 5-round read-out shows the result of 5 iterations of those steps on 4 different fragments of the protein P47420 from the mycoplasma proteome, considering no errors (fragment #1 is anchored at the N-terminus and no reads are recorded; fragment #2 read-out is II for two isoleucine read-outs, abbreviated I, followed by loss of fragment from the gel after in-gel Edman degradation cleaves the fragment from the gel; fragment #3 read-out is ILRVI, where L is leucine, R is arginine, V is valine, assuming these binders are available; fragment #4 read-out is H for histidine followed by loss of fragment from the gel after in-gel Edman degradation cleaves the fragment from the gel).

(C) Heatmap of mean percent amino acid count in the *Mycoplasma genitalium* (referred to as "mycoplasma" hereafter) proteome across different types of anchoring conditions (AcX, epoxide) and types of digestion conditions (Lys-C, trypsin, Proteinase-K) at different probabilities of fixation (abbreviated: P(fix)), anchoring (abbreviated: P(anchoring)), and digestion (abbreviated: P(digestion)), with 15 in-gel Edman degradation rounds. Mean percent amino acid count is defined as the number of unmodified sequenceable amino acids (i.e. unfixed, unanchored, retained in gel after digestion or Edman degradation) divided by the length of the protein, averaged across all proteins in the proteome. We run this probabilistic process 10 times and average the resulting mean percentage amino acid count across trials for every combination of conditions.

(D) Fraction correct of the identified proteins out the whole mycoplasma proteome for very low (black line on the plots), low (green line on the plots), medium (blue line on the plots) and perfect (red line on the plots) binder specificities (see **Supplementary Table 12** for the kinetic values describing these specificity regimes) over increasing numbers of in-gel Edman degradation (from 5 to 15 rounds). Fraction correct is defined as the number of proteins identified correctly (i.e. not false positive and not uncertain; see Methods: Hidden Markov Model (HMM) based matching and fraction of proteome correctly identified for more details) divided by the size of the proteome. (i) assuming all 20 amino acid binders are sequentially added for recognition after each round, (ii) same as (i) but with 15 amino acid binders, against {A, N, D, E, Q, G, I, L, K, F, S, T, P, Y, V} (iii) 10 amino acid binders, against {A, N, D, E, Q, I, L, K, F, V} (iv) 5 amino acid binders, against {N, E, L, F, V} (dot: mean value of the fraction correct for that given condition; error bar: standard deviation for that given condition, n=10 simulations of the fixation, anchoring and digestion with 0.05 fixation, 0.8 AcX anchoring and 0.8 trypsin digestion, 0.1 Edman conjugation failure, 0.3 Edman cleaving failure).

#### Supplementary Figure 14

(A)

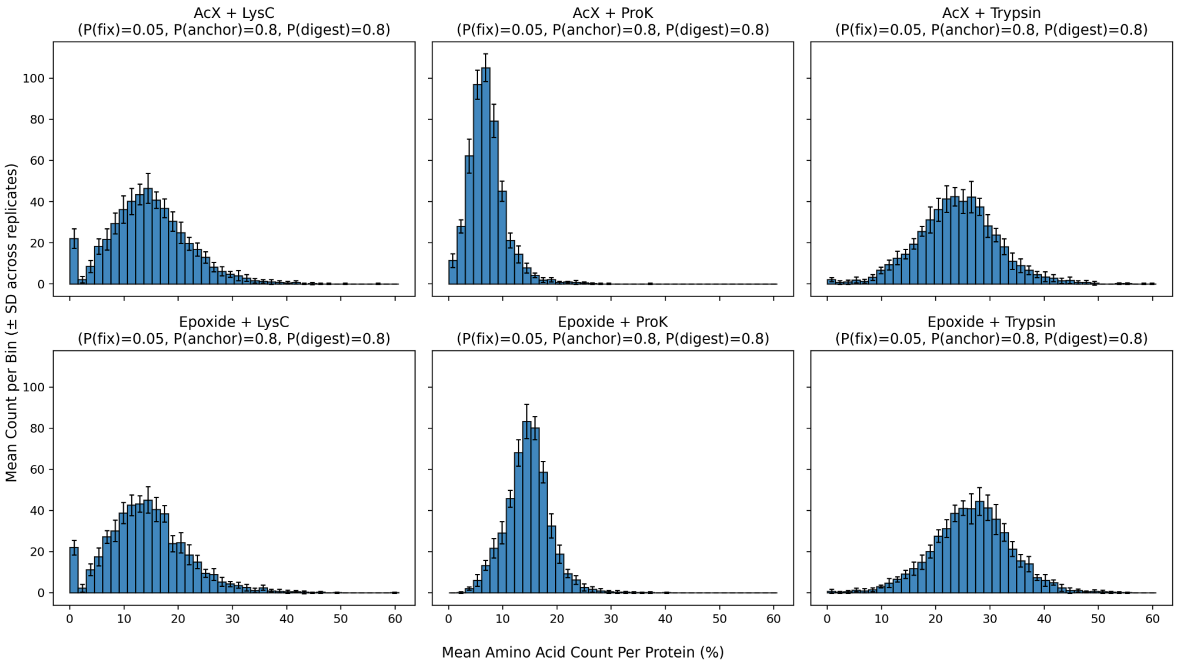

(B)

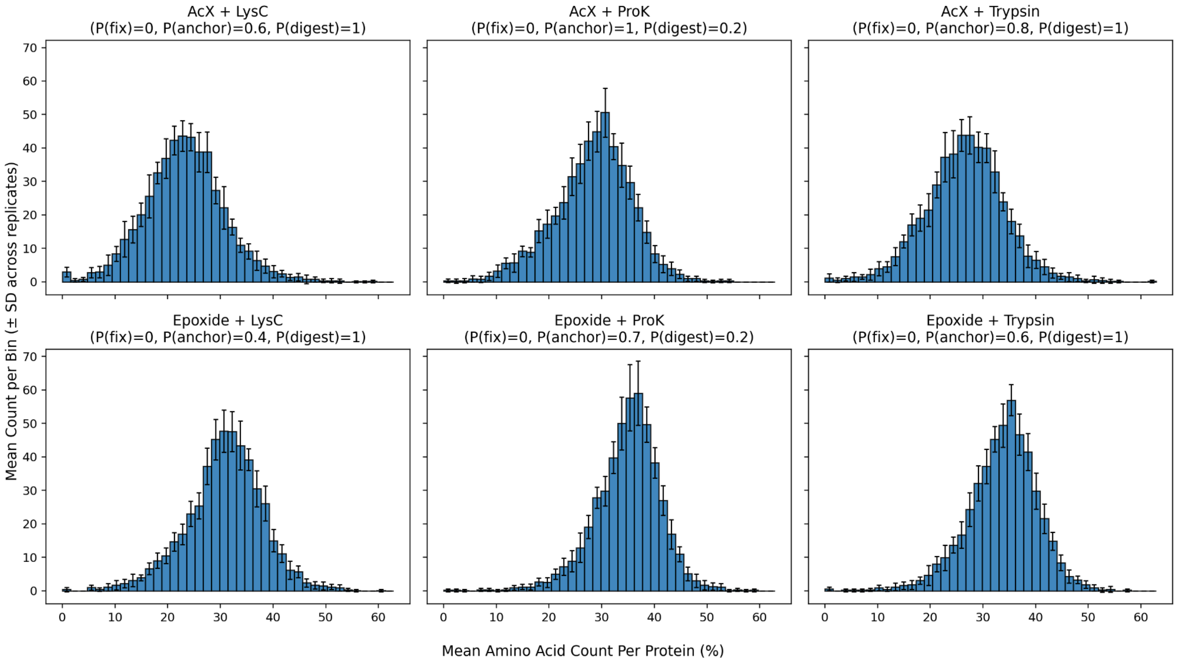

(C)

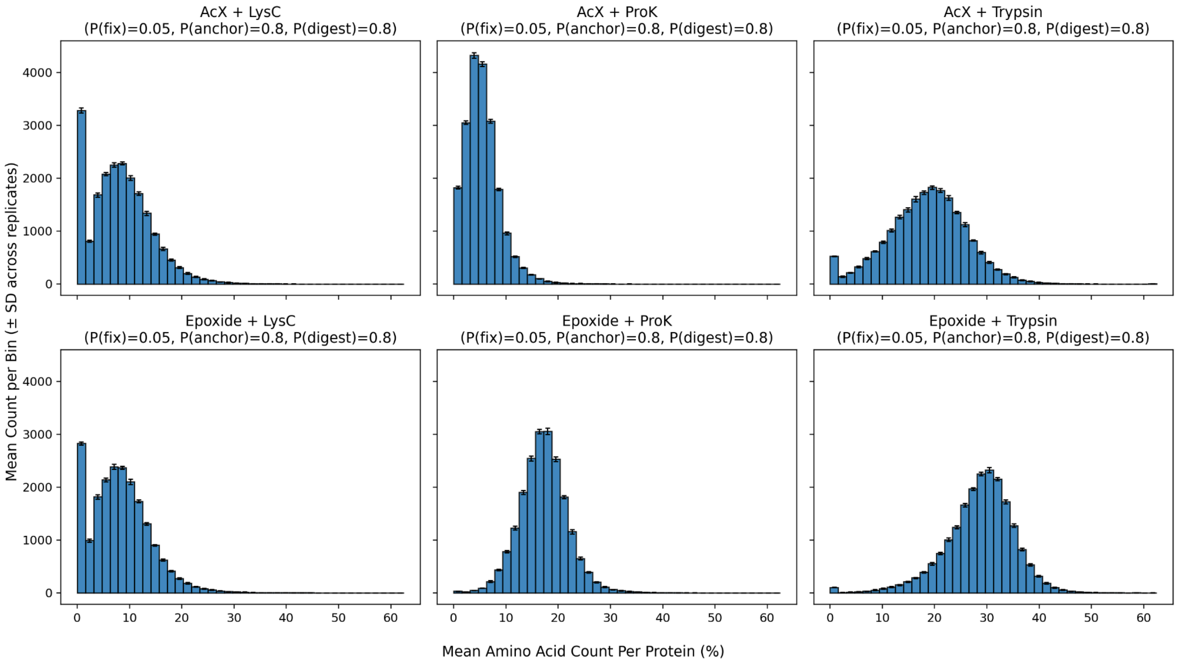

(D)

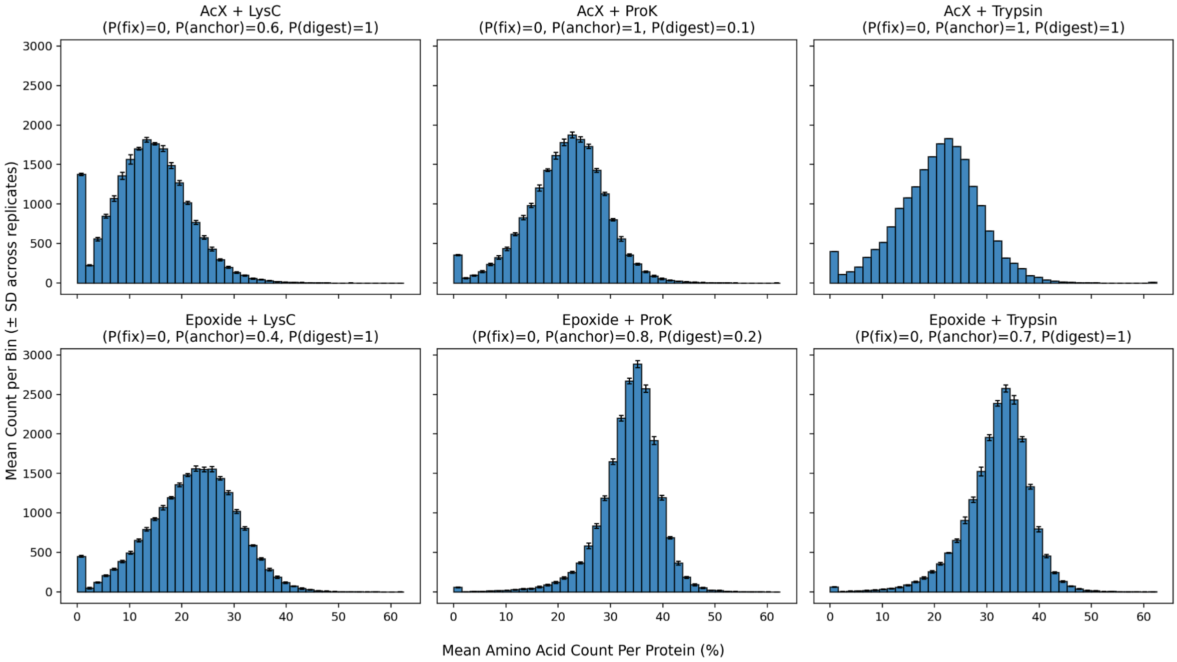

**Supplementary Figure 14:** Related to the theoretical assessment of in situ protein sequencing Supplementary Figure 13C (mycoplasma proteome) and Supplementary Figure 18 (human proteome) for the analysis of the distribution of the mean percent amino acid count for a certain set of parameters. Histogram distribution of the percent amino acid count for six different conditions of the chemistry: 6-((acryloyl)amino)hexanoic acid, succinimidyl ester (abbreviated: AcX) or epoxide anchoring in combination with endopeptidase Lys-C (abbreviated: Lys-C), proteinase K (abbreviated: ProK) or trypsin digestion (specific conditions are listed above each histogram). Mean protein count per bin is the frequency of proteins over the proteome (each bin has a range of 2.5% amino acid

count). Percent amino acid count is defined as the percentage of residues of a collection of fragments making up a protein that are not modified and remain accessible for read-out over 15 rounds of in-gel Edman degradation after the chemical steps. The simulation is run 10 times for fixation, anchoring and digestion and the resulting fragments are assessed for their percent amino acid count in the gel.
(A) The histogram results showing the distribution of percent amino acid count for all the proteins in the mycoplasma proteome (UniProt ID 243273, total of 483 proteins) for each condition with a fixed parametrization P(fixation)=0.05, P(anchoring)=0.80, P(digestion)=0.80 (abbreviated P(fix), P(anchor), P(digest), respectively). Blue bars: number of proteins with the given percent amino acid count; black error bars: standard deviation across the 10 simulations, throughout this figure.
(B) The histogram results showing the distribution of percent amino acid count for all the proteins in the mycoplasma proteome for each condition with the best parametrization maximizing the averaged percent amino acid count (highest parametrization for the given condition in **Supplementary Figure 13C**), where the best parametrization is labeled at the top of each histogram.
(C) The histogram results showing the distribution of percent amino acid count for all the proteins in the human proteome (UniProt ID 9606, total of 20,421 proteins) for each condition with a fixed parametrization, as in (A).
(D) The histogram results showing the distribution of percent amino acid count for all the proteins in the human proteome for each condition with the best parametrization maximizing the averaged percent amino acid count (highest parametrization for the given condition in **Supplementary Figure 18**), where the best parametrization is labeled at the top of each histogram.

#### Supplementary Figure 15

(A)

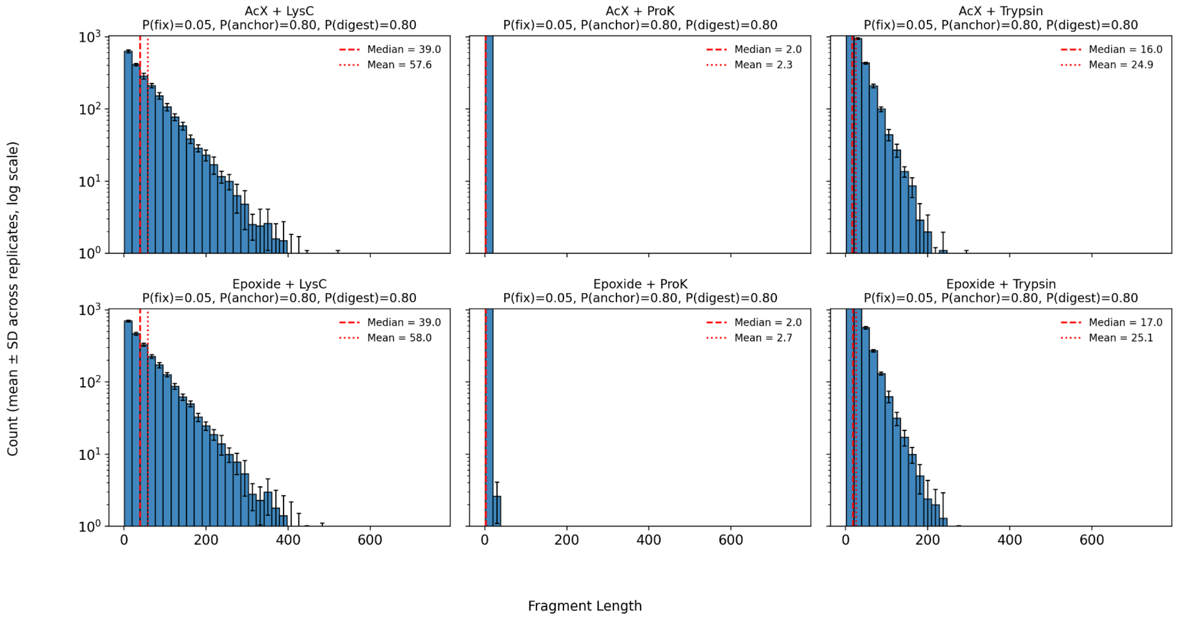

(B)

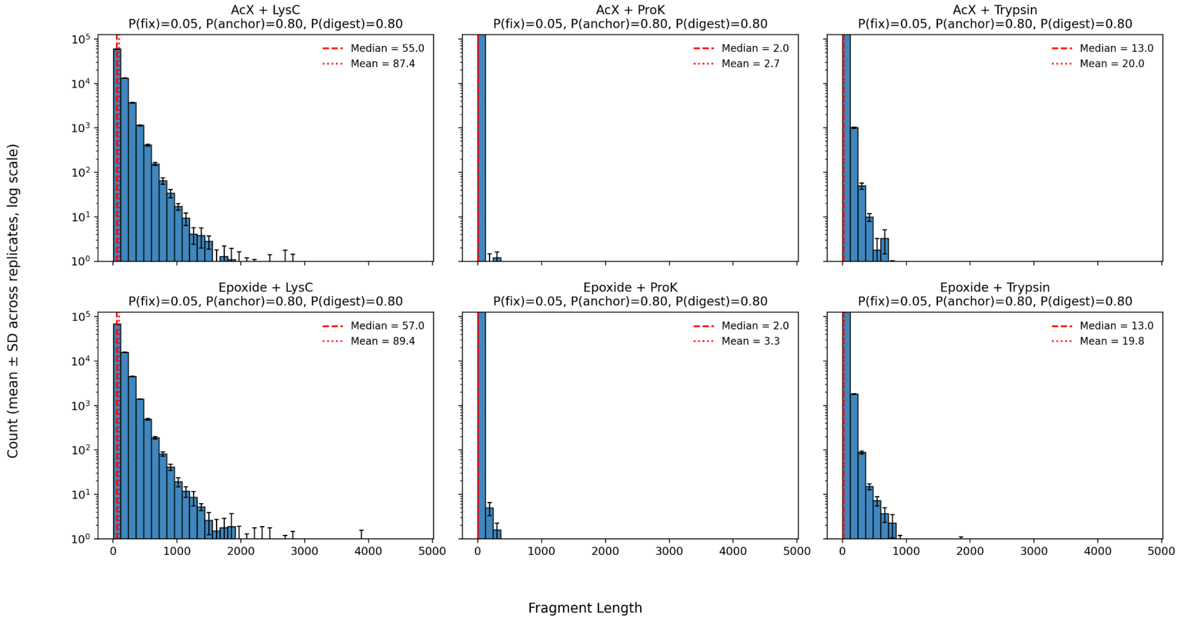

**Supplementary Figure 15:** Related to the theoretical assessment of *in situ* protein sequencing Supplementary Figure 13C (mycoplasma proteome) and Supplementary Figure 18 (human proteome), analyzing the fragment lengths under various conditions for fixation, anchoring and digestion. Fragment length is defined as the number of amino acids for a given fragment retained in the gel, but not considering amino acids that are located downstream (C-terminal) of the last anchoring amino acid of the fragment (since this fragment would be lost after in-gel Edman degradation). Histogram of the distribution of fragment lengths for different parametrization of the chemistry: AcX or epoxide anchoring with Lys-C, ProK or trypsin digestion (specific condition listed above each histogram). The simulation is run 10 times for fixation, anchoring and digestion and the resulting fragments are assessed for their fragment length.

(A) The results for the mycoplasma proteome (UniProt ID 243273, total of 483 proteins). Varying the anchoring conditions (AcX, or epoxide digestion conditions) and the digestion condition (trypsin, Lys-C, or ProK digestion conditions), where P(fixation)=0.05, P(anchoring)=0.80, P(digestion)=0.80 for all conditions (abbreviated: P(fix)=0.05, P(anchor)=0.80, P(digest)=0.80). (blue bars: number of proteins with the given fragment length; error bars: standard deviation for each bin across 10 simulations; vertical red dashed line with long dash: median fragment length; vertical red dashed line with short dash: mean fragment length).

(B) As in (A), but the results for the fixed parametrization (P(fix)=0.05, P(anchor)=0.80, P(digest)=0.80) for each condition for the human proteome (UniProt ID 9606, total of 20,421 proteins).

#### Supplementary Figure 16

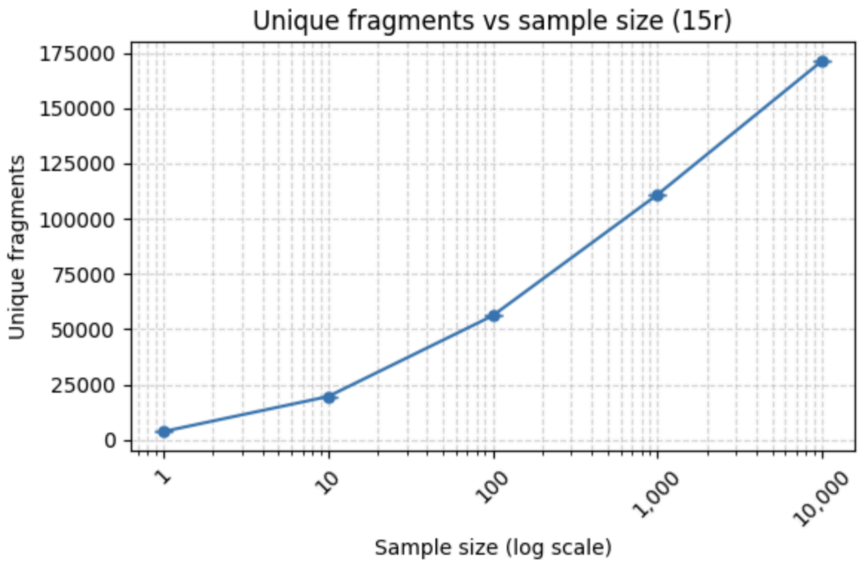

**Supplementary Figure 16:** Related to the theoretical assessment of *in situ* protein sequencing Supplementary Figure 13**B**, analyzing how increasing the number of simulations impacts the number of unique fragments stored in what we named the "reference fragment dataset" (defined as the database of the ground-truth sequences). Plot showing the number of unique fragments (over the first 15 amino acids from the N-terminus; relevant to 15 rounds of in-gel Edman degradation; abbreviated "15r") versus the number of simulations of fixation, anchoring and digestion chemistries (this is referred to as "Sample size" on the graph x-axis). The results for the mycoplasma proteome (UniProt 243273). For the true fragment space (i.e. no readout errors), we plot the number of unique fragments generated for 15 Edman rounds where the sample size is the number of times we perform the simulation of fixation, anchoring and digestion steps on the mycoplasma proteome.

#### Supplementary Figure 17

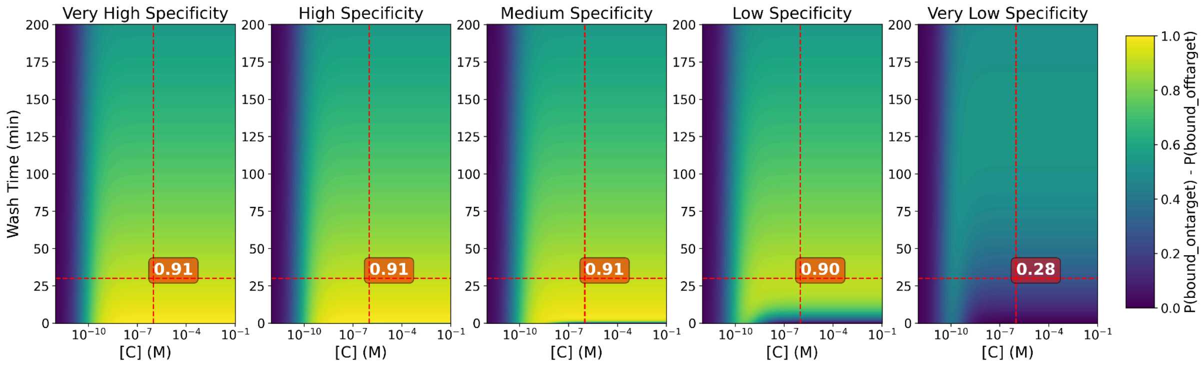

**Supplementary Figure 17:** Related to the theoretical assessment of *in situ* protein sequencing [Supplementary Figure 13](#)**D**, analyzing how wash time (min) and concentration of binder affects read-out. Heatmaps showing the difference between probability that a binder correctly binds its on-target and the probability that it incorrectly binds off-target sites when changing the wash time and the concentration of binder. This difference represents how much more likely an on-target site is to be occupied compared to an off-target site, but it does not reflect the overall binding probability across all possible targets, since it assumes equal site availability between on-target and off-target sites. These results are plotted across varying binder specificity regimes ("very high", "high", "medium", "low" and "very low"; where the kinetic values associated to these binder specificity regimes are detailed in Supplementary Table 12). The results assume that we add excess binder, and that binding reaches equilibrium (using the Langmuir equation for binding) such that after the washing, the probability of being bound to on-target and off-target is defined as:

$$p_{bound-ontarget} = \left(\frac{[C]}{[C] + K_d^{on-target}}\right) \cdot e^{-k_{off}^{on-target} t_{wash}}$$ and

$$p_{bound-offtarget} = \left(\frac{[C]}{[C] + K_d^{off-target}}\right) \cdot e^{-k_{off}^{off-target} t_{wash}}$$, where we plot

$p_{bound-ontarget} - p_{bound-offtarget}$. This does not consider any effects that the gel environment would have on the kinetics of binding, nor does it consider any non-specific background from binders bound non-specifically to the gel or not washed out. Here, $[C]$ is the concentration of binder, $K_d^{on-target}$ the equilibrium dissociation constant for the on-target, $K_d^{off-target}$ the equilibrium dissociation constant for the off-target, $k_{off}^{on-target}$ the dissociation rate for the on-target, and $k_{off}^{off-target}$ the dissociation rate for the off-target, and $t_{wash}$ is the time elapsed during the dissociation phase (during the washing). For $[C] = 1\ \mu M$ and $t_{wash} = 30\ min$, the probability difference is noted at the intersection of two dashed red lines for each binder specificity regime. These probabilities, for medium, low, and very low specificity, are used in the simulation of the sequencing in Supplementary Figure 13**D**.

#### Supplementary Figure 18

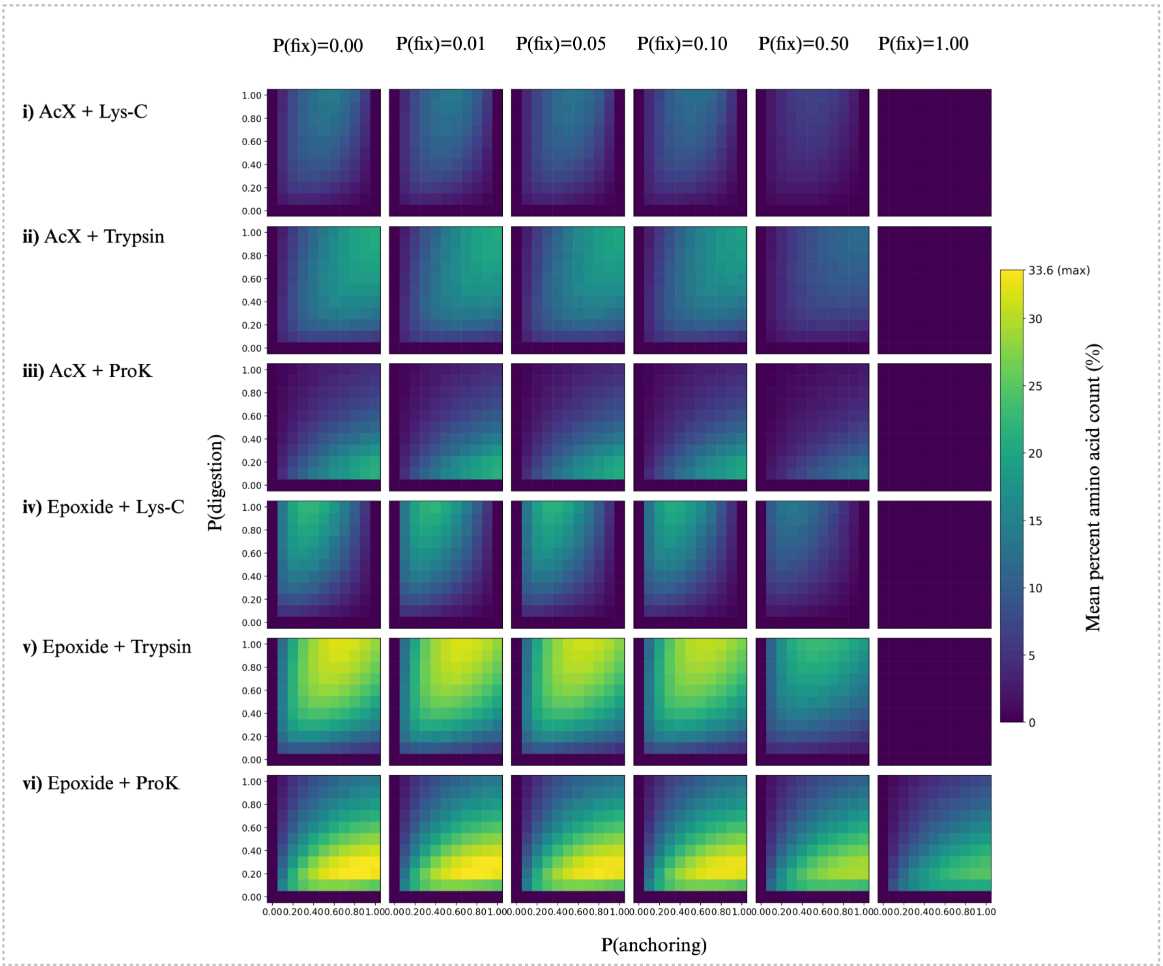

**Supplementary Figure 18:**

(A) Related to the theoretical assessment of in situ protein sequencing [Supplementary Figure 13](#)C, but rather than plotting the mycoplasma proteome results, plotting the human proteome results. Heatmap of mean percent amino acid count amino acid count (%) in the human proteome (UniProt ID 9606) across different types of anchoring conditions (AcX, epoxide) and types of digestion conditions (Lys-C, trypsin, Proteinase-K) at different probabilities of fixation, anchoring, and digestion, with 15 Edman rounds. Percent amino acid count is defined as the percentage of residues of a collection of fragments making up a protein that are not modified and remain accessible for read-out over 15 rounds of in-gel Edman degradation after the chemical steps. We run this probabilistic process 10 times and average the resulting mean percentage amino acid count across trials for every combination of conditions.

#### Supplementary Figure 19

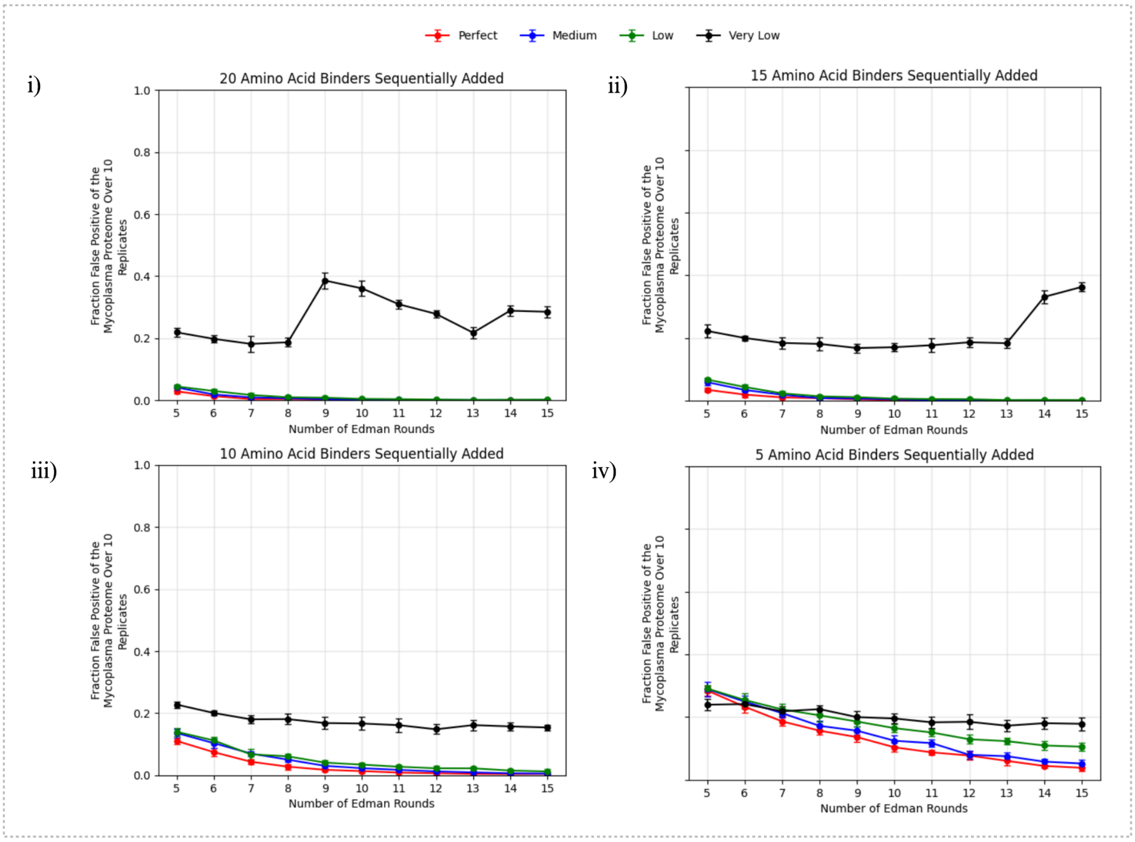

**Supplementary Figure 19:** Fraction false positive of the identified proteins out of the whole mycoplasma proteome for very low (black line on the plots), low (green line on the plots), medium (blue line on the plots) and perfect (red line on the plots) binder specificities (see Supplementary Table 12 for the kinetic values describing these specificity regimes) over increasing numbers of in-gel Edman degradation (from 5 to 15 rounds). (i) assuming all 20 amino acid binders are sequentially added for recognition after each round, (ii) same as (i) but with 15 amino acid binders, for {A, N, D, E, Q, G, I, L, K, F, S, T, P, Y, V} (iii) 10 amino acid binders, for {A, N, D, E, Q, I, L, K, F, V} (iv) 5 amino acid binders, for {N, E, L, F, V} (dot: mean value of the fraction correct for that given condition; error bar: standard deviation for that given condition, n=10 simulations of the fixation, anchoring and digestion with 0.05 fixation, 0.8 AcX anchoring and 0.8 trypsin digestion, 0.1 Edman conjugation failure, 0.3 Edman cleaving failure).

**Supplementary Figure 20**

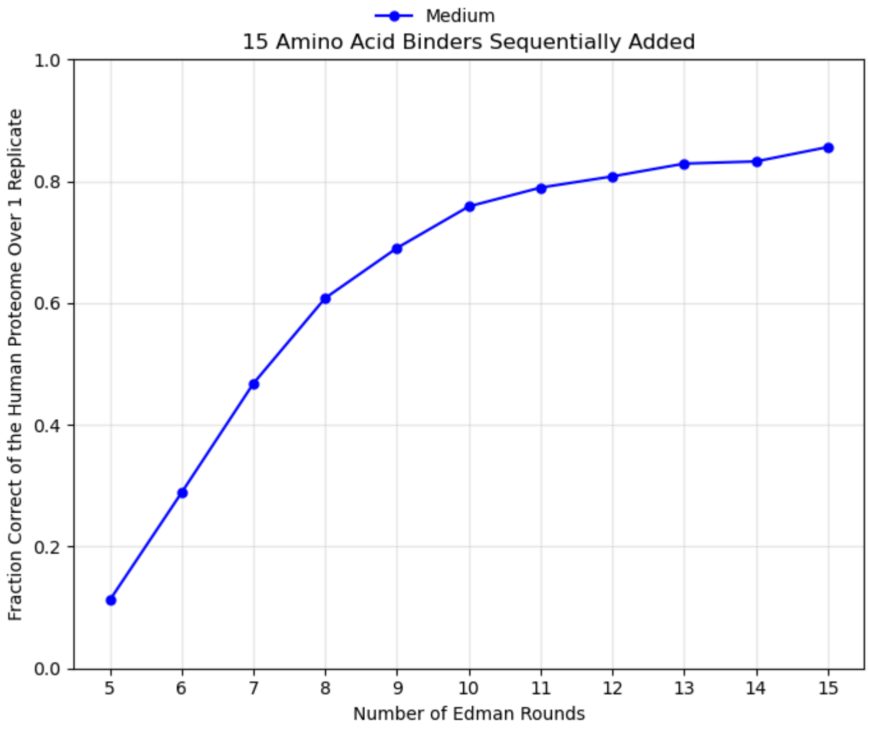

**Supplementary Figure 20**: Fraction correct of the identified proteins out the whole human proteome (Uniprot ID: 9606) for medium (blue line on the plots) binder specificities (see **Supplementary Table 12** for the kinetic values describing these specificity regimes) over increasing numbers of in-gel Edman degradation (from 5 to 15 rounds), with 15 amino acid binders sequentially added. Fraction correct is defined as the number of proteins identified correctly (i.e. not false positive and not uncertain; see **Methods**: Hidden Markov Model (HMM) based matching and fraction of proteome correctly identified for more details) divided by the size of the proteome (20,421 proteins). (i) assuming 15 amino acid binders are sequentially added for recognition after each round, against {A, N, D, E, Q, G, I, L, K, F, S, T, P, Y, V (dot: mean value of the fraction correct for that given condition; n=1 simulations of the fixation, anchoring and digestion with 0.05 fixation, 0.8 AcX anchoring and 0.8 trypsin digestion, 0.1 Edman conjugation failure, 0.3 Edman cleaving failure with 10 samples as part of reference fragment dataset).

**Supplementary Figure 21**

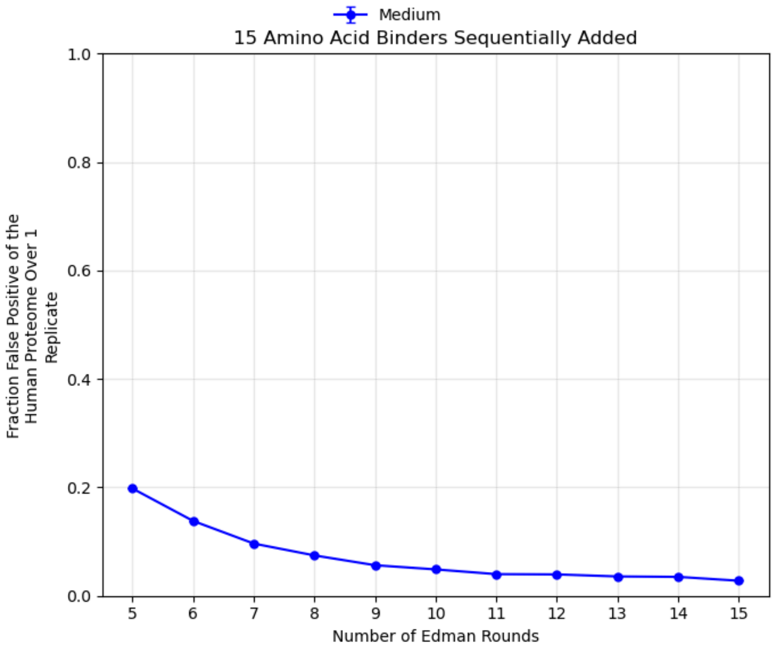

**Supplementary Figure 21**: Fraction false positive of the identified proteins out of the whole human proteome for medium (blue line on the plots) binder specificities (see **Supplementary Table 12** for the kinetic values describing these specificity regimes) over increasing numbers of in-gel Edman degradation (from 5 to 15 rounds). (i) assuming 15 amino acid binders are sequentially added for recognition after each round, for {A, N, D, E, Q, G, I, L, K, F, S, T, P, Y, V} (dot: mean value of the fraction correct for that given condition; n=1 simulations of the fixation, anchoring and digestion with 0.05 fixation, 0.8 AcX anchoring and 0.8 trypsin digestion, 0.1 Edman conjugation failure, 0.3 Edman cleaving failure with 10 samples as part of reference fragment dataset).

#### Supplementary Figure 22

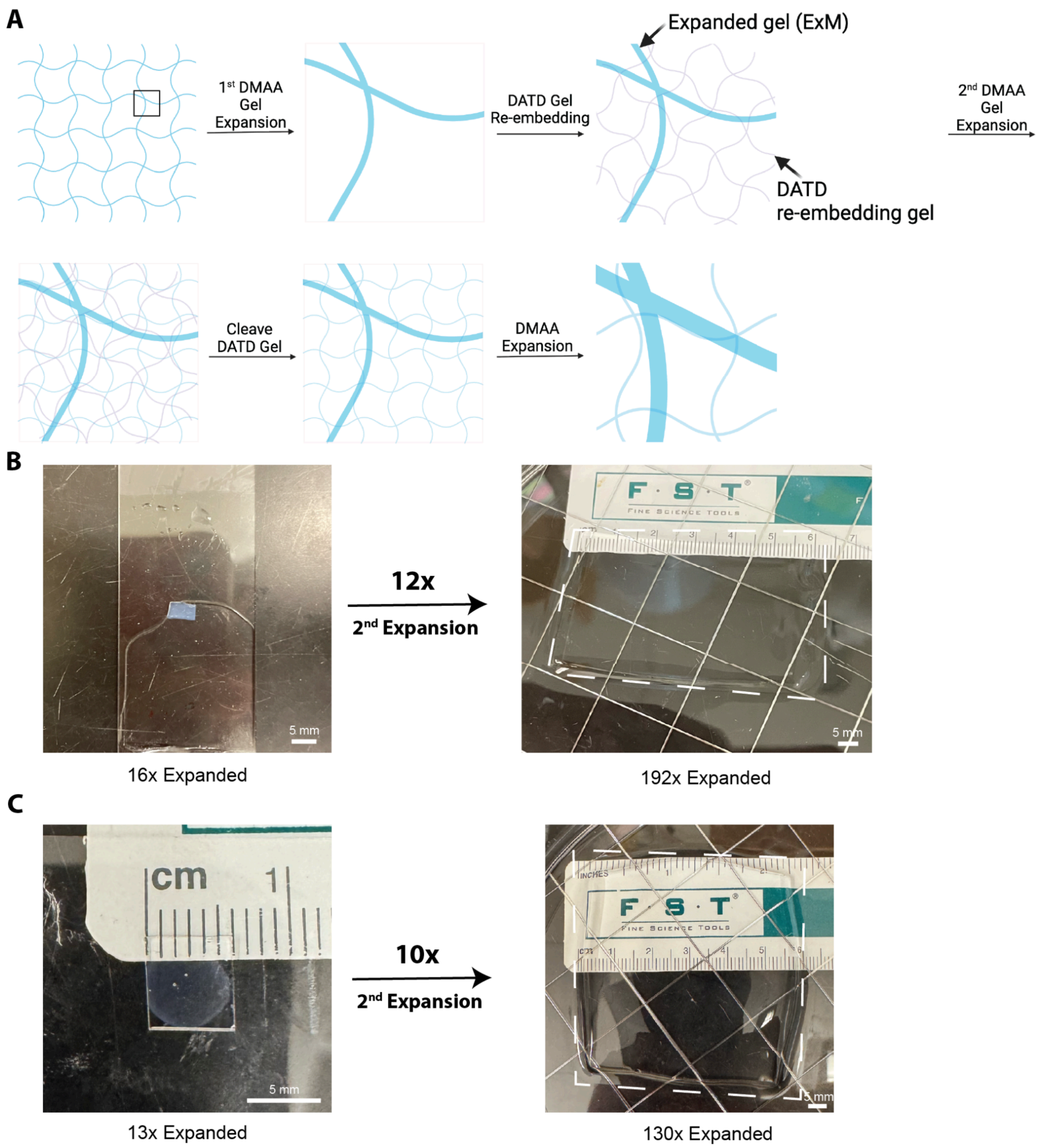

A) ~130-190× expansion workflow. The first N,N-dimethylacrylamide (DMAA)-based expansion gel is expanded ~16-fold using an established protocol[11]. The expanded DMAA gel is then re-embedded in a cleavable N,N'-Diallyl-L-tartardiamide (DATD) gel to preserve its expanded structure during subsequent processing. Next, a second DMAA expansion gel is cast throughout the re-embedded gel, followed by cleavage of the DATD re-embedding gel. The second DMAA gel is then expanded to achieve an overall expansion of ~130-190-fold (see Supplementary Note 11 for the protocol).

B) Physical expansion of the re-embedded gel during the final expansion step. Representative photographs of the gel before and after the final expansion. The gel expands ~12 fold during the second expansion step. The dashed outline marks the boundary of the expanded gel. Scale bars in white: 5 mm.

C) Another example of a gel, as in (B), with the gel expanding ~13-fold in the first expansion step, and ~10 fold during the second expansion step.

### Supplementary Notes

#### Supplementary Note 1: Information about previous solvents used in Edman degradation

This section is to add more background to the section **Solvent considerations for in-gel Edman degradation**. Edman degradation can determine the sequence of the first ~30 amino acids from the N-terminus (since over multiple rounds the cumulative yield decreases exponentially, with, for instance, over 30 rounds $0.99^{30}$ = 74% yield due to accumulated error) by detecting the cleaved phenylthiohydantoin amino acid (PTH-aa)[12,13]. Organic solvents solubilize PITC, a hydrophobic molecule, minimize its hydrolysis with water, and promote efficient isolation of the PTC-peptide product. Typically, pyridine and triethylamine maintain slightly alkaline conditions (e.g., pH 8-9), without adding additional buffers[14]. Acetonitrile (ACN), used in modern protocols[15–17], has advantages such as its higher polarity, volatility, reduced UV absorbance for chromatographic analysis, and reduced odor and handling concerns. Maintaining alkaline conditions in solvents such as ACN can be performed by adding triethylamine or pyridine[15,16,18–21]. After conjugation, PTC-peptides are recovered by evaporation, precipitation, or extraction, and are then cleaved with anhydrous acid (e.g., most commonly trifluoroacetic acid (TFA)). The differing solubilities of PTH-aa (hydrophobic) and peptide (hydrophilic) enable their separation by phase or solid-liquid partitioning. In the latter case, a non-polar solvent can precipitate the peptide, that is then separated from the soluble PTH-aa that is detected using paper chromatography[12]. Improvement in speed and sensitivity of the sequencing method include an automated version of this chemistry[13], peptide degradation on solid support[16], and detection of PTH-aa using HPLC[22], and/or coupled with mass spectrometry[23]. Despite these various modifications, all of these procedures rely on bulk PTH-aa extraction and detection[24].

In terms of the cleavage step, other solvents than TFA can enable cleavage (see **Supplementary Table 2** for a non-exhaustive list of Edman degradation protocol variations). Notably, boron trifluoride etherate in ACN[25,26] has been recently used for DNA-encoded Edman degradation. It also required chemically modified 7-deazapurine deoxynucleotides (c7dA and c7dG) for DNA resistance to depurination[27].

#### Supplementary Note 2: Information about ionic strength of different buffers and solutions used on ExM and ExMre gels

Different buffers have different ionic strengths, calculated below:

Ionic strength (I) = $\frac{1}{2}\sum_{i=1}^{n}(c_i * z_i^2)$, where $c_i$ is the molar concentration of an ion and $z_i$ is its charge, summed over all ions, $n$, present in the solution.

- For 1X PBS pH 7.4, three different types of salts: potassium phosphate monobasic (1.06 mM $KH_2PO_4$), sodium chloride (155.17 mM NaCl), sodium phosphate dibasic (2.97 mM $Na_2HPO_4$) contribute to ionic strength:

  $$I = \frac{1}{2}\sum_{i=1}^{n}(c_i * z_i^2) = \frac{1}{2}\{(c_{K^+} * z_{K^+}^2) + (c_{H_2PO_4^-} * z_{H_2PO_4^-}^2) + (c_{Na^+} * z_{Na^+}^2) + (c_{Cl^-} * z_{Cl^-}^2) + (c_{HPO_4^{2-}} * z_{PO_4^{2-}}^2)\}$$

  $$I = \frac{1}{2}\sum_{i=1}^{n}(c_i * z_i^2) = \frac{1}{2}\{(1.06 * 1^2) + (1.06 * 1^2) + (161.11 * 1^2) + (155.17 * 1^2) + (2.97 * 2^2)$$

}

  $I \simeq 165\, mM.$

- For re-embedding solution, two different types of salts: ammonium persulfate (0.075% APS i.e., 3.29 mM; $(NH_4)_2S_2O_8$) and Tris pH 8 (5 mM Tris-HCl) contribute to ionic strength:

  $$I = \frac{1}{2}\sum_{i=1}^{n}(c_i * z_i^2) = \frac{1}{2}\{(c_{NH_4^+} * z_{NH_4^+}^2) + (c_{S_2O_8^{2-}} * z_{S_2O_8^{2-}}^2) + (c_{Tris-H^+} * z_{Tris-H^+}^2) + (c_{Cl^-} * z_{Cl^-}^2)\}$$

  $$I = \frac{1}{2}\sum_{i=1}^{n}(c_i * z_i^2) = \frac{1}{2}\{(6.57 * 1^2) + (3.29 * 2^2) + (5 * 1^2) + (5 * 1^2)\}$$

  $I \simeq 15\, mM.$

The higher ionic strength of 1X PBS, compared to the re-embedding solution, contributes to why ExM gel shrinks more in 1X PBS than in the re-embedding solution.

After re-embedding is performed, the gel has a new internal structure. When placed in 1X PBS ExMre gels expand from hydration, and then can expand more in water.

#### Supplementary Note 3: Information about LC/QToF solvents and implications for Edman degradation read-out

The effect of formic acid and electrospray ionization (ESI) on read-out of PTC-peptides from in-gel Edman degradation

As mentioned in section **Testbed peptides anchored throughout ExMre gels for validation of in-gel Edman degradation**, peptides with a cleaved N-terminal amino acid can be detected using LC/QToF in the condition where only PITC has been added, without the second neat TFA cleavage step. This can be explained by the presence of 0.1% formic acid in the solutions used for liquid chromatography and ionization (see **Methods** section, under **LC/QToF analysis of synthetic peptides in the gel**, **LC Method**), as well as the use of ESI.

First, the acidic ~pH 2.7 of the aqueous solution could promote cleavage of the PTC-peptide. An aqueous acidic solution (i.e., hydrolytic solution) can also cleave peptide bonds within the peptide, but this requires high temperature, 100-160 °C, for 18-72 hours[28]. As such, peptide bond cleavage, beyond N-terminal amino acid cleavage via Edman degradation chemistry, is unlikely with the methods used here for LC/QToF detection of trypsinized fragments from the gel (0.1% formic acid for ~23 min at 30 °C). This would correspond to on-column cleavage where the peak of the peptide with cleaved N-terminal amino acid matches the retention time of the cleaved peptide in the PITC to DMSO (1:1000 ratio PITC:DMSO) followed by TFA condition (see **Source Data** for raw chromatograms and associated mass spectra).

Furthermore, we discovered that ESI can also promote the conversion of PTC-peptide to peptide with cleaved N-terminal amino acid from in-source fragmentation. Previously, ESI was shown to have an impact on interpretation of data[29]. In addition, previous reports show PTC-peptides were placed in a collision cell for cleavage[30]. Data here also demonstrates this occurs with PTC-peptides with ESI. In this case, unlike the on-column cleavage with formic acid, the peptide with cleaved N-terminal amino acid has a later retention time that matches the PTC-peptide retention time, since the cleavage happens after the elution, most likely inside the ESI source (see **Source Data** for the raw chromatograms and associated mass spectra).

Thus, both on-column cleavage and the in-source cleavage of the PTC-peptide could occur for samples in the mass spec instrument, either from the 0.1% formic acid in the solution for ionization and/or ESI.

Comparing the ionization efficiency of non-modified peptide (AGGAGGLLGGSR) and peptide with cleaved N-terminal amino acid (GGAGGLLGGSR)

The effect of formic acid and electrospray ionization on PTC-peptide precludes assessing the yield of the reaction by comparing the abundance of the non-modified peptide with the peptide with cleaved N-terminal amino acid. However, a control curve was still performed in order to assess the relative ionization efficiency of the non-modified peptide to the peptide with cleaved N-terminal amino acid. The ionization efficiencies were considered similar between the two species, as assessed with the control, which did not provide enough evidence to confidently say there's a difference in ionization efficiency between the two species (see **Supplementary Figure 5** for the control curve; in addition, since alanine is small and not ionizable, its loss after Edman degradation would suggest a similar peptide ionization efficiency to the parent peptide).

#### Supplementary Note 4: Conventional Edman degradation conditions in the gel

This Supplementary Note discusses conventional Edman degradation conditions that were tested using trypsinization-with-LC/QToF assay in the gel. We investigated two PITC-delivery solvents that we had shown to cause extreme gel shrinkage and opacity - pyridine and ACN - and found that PITC conjugation to gel-anchored peptide did not occur, even when PITC was administered at 1:9 in ACN (see **Supplementary Figure 3A** and **Supplementary Figure 3Bi** for results of peptide species detected on LC/QToF using these conditions; see **Supplementary Table 8** for full raw data and descriptive statistics related to **Supplementary Figure 3**). 1:1 pyridine:water, which caused comparable shrinkage but did not result in the visible opacity caused by pyridine and ACN, did result in PITC conjugation after adding 1:9 PITC:solvent, although with still some (~20%) peptide unreacted (top graph in **Supplementary Figure 3Bii**, n=1). In addition, to assess the possibility of performing PITC conjugation to the N-terminus of peptides in the gel using a purely aqueous solution, which is known to be compatible with ExMre gels, we tested 0.1 M sodium bicarbonate ($NaHCO_3$) pH 8.5 with PITC (1:9 PITC:buffer) for PTC-peptide formation. The results showed ~70% unreacted peptide (**Supplementary Figure 3Biii**, n=1). This suggested a partial Edman conjugation reaction in the ExMre gels in these last two cases. Although 0.1 M $NaHCO_3$ pH 8.5 did not shrink the gel, and did not lead to gel opacity (see **Supplementary Figure 1Hi** for representative gel images), the low yield of PITC conjugation in 0.1 M sodium bicarbonate pH 8.5 might be explained by the insoluble nature of PITC in aqueous buffer. In summary, Edman conjugation in the ExMre context correlates strongly with lack of shrinkage or opacity. In addition, it may also be affected by PITC solubility in the solvent at hand.

#### Supplementary Note 5: Phenylthiohydantoin amino acid (PTH-aa) detection

Calculation of the total dry weight of PTH-F recovered from the ExMre gels and injected into the LC/QToF:

Gel volume: 5*5*0.35 mm is ~9 µL

Several N-terminal amino acid peptides were tested for PTH-aa recovery after in-gel Edman degradation using 9% acrylamide gels (due to ease and speed of making and testing in these gels) and ExMre gels. We thought it useful to comment on these results, we note that our data should be regarded as preliminary (n=1 gelation solution for each condition).

PTH-tyrosine (PTH-Y), and PTH-glycine (PTH-G) were detected after PITC (1:9 ratio PITC:DMSO) conjugation reaction and TFA cleavage in 9% acrylamide gel (5 x 5 x 0.31 mm) containing N-terminal tyrosine or N-terminal glycine (**Xaa**GGAGRGLGK{acr}, where Xaa can be tyrosine or glycine), respectively. PTH-alanine was also detected after PITC (1:9 ratio PITC:DMSO) conjugation reaction and TFA cleavage in ExMre gel (5 x 5 x 0.7 mm) containing N-terminal alanine with the same sequence as above (i.e., AGGAGRGLGK{acr}). This included removal of TFA from the vial and resuspension in 1:1 acetonitrile to water, as in **Figure 3H**. These specific species, with exact mass (calculated with a mass accuracy within a 20 ppm window, see **Edman degradation and PTH detection** section in **Methods** for the relevant equation for calculating the mass accuracy window), matched the expected species. They sometimes exhibited different retention times to the PTH-aa positive control, which may be a result of the presence of an anilinothiozolinone (ATZ) intermediate in the experimental conditions. Indeed, PTH-aa and ATZ-aa have different chemical structures and properties, where treating with aqueous acid and heat can convert the ATZ isomer to PTH isomer. However, using this same protocol, PTH-tryptophan (PTH-W) was not detected, and may be due to differences in solubility in 1:1 acetonitrile to water, and/or differences in extraction of the PTH-aa from the gel after TFA due to distinct biochemical properties.

#### Supplementary Note 6: ClpS2 St-V1 thoughts on affinity and ClpS2 St-V1, tvClpS2 Q31H sequence information

This note serves to add information about ClpS2 St-V1 protein used in **Figure 4**.

1. Thoughts on affinity

The ClpS2 V1 binder has been previously used for non-in situ peptide sequencing[31] (sequence information for ClpS2 variants in **Supplementary Note 6.2**, below). ClpS2 V1 has a measured $k_{off}$ of ~0.1 $s^{-1}$ (or ~10 second dwell time) towards phenylalanine[32]; the variant ClpS2 St-V1 has improved thermostability while retaining the previously engineered higher affinity towards Phe[33]. However, with a dissociation rate on the order of seconds, this NAAB cannot be used for reliable in situ localization of single molecules when using downstream multi-step signal amplification strategies (requiring hours) employed in ExM protocols, which rely on enzymes or self-assembly cascades[34,35]. Here, to focus on the fundamental compatibility of protein sequencing technology components with the in situ milieu, we do not attempt single molecule imaging, and instead use bulk gel fluorescence to gauge whether NAAB binding can work on peptides in an ExM gel environment. As NAAB quality improves, an exciting next step will be to follow binding with signal amplification and then to do single molecule imaging - as has been done for other molecule types, e.g. RNA, in past expansion microscopy papers[34,35]. Our current experiments also allow an exploration of binder diffusion in the gel, important for any functioning in situ protein sequencing protocol.

2. Sequence information for ClpS2 St-V1

*Agrobacterium tumefaciens* ClpS2 was first engineered for higher specificity towards Phe (ClpS2 V1), and then for higher thermostability, named ClpS2 St-V1[32,33]. The ClpS2 St-V1 protein used in **Figure 4** has a C-terminal hemagglutinin (HA) tag. The sequence is as follows:

SSDSPVDLKPKPKVKPKLERPKLYKVMLLNDDYTP**M**SFVTEVLKAVFNMSEDQGRRVMMTA
HRFGSAVVGVSTRDIAETKAKQATDLAREAGFPLMFTTEPEE-*GGSGGSYPYDVPDYA**

where red highlighting indicates the proposed substrate contacts with phenylalaninamide[32,36], underlined and bolded residues represent the mutations of *A. tumefaciens* ClpS2 wild-type to result in ClpS2 V1 (R35M and E36S), and yellow highlighting indicates amino acids that are different between ClpS2 V1 and ClpS2 St-V1[33]. The ClpS2 V1 sequence is as follows [31]:

MSDSPVDLKPKPKVKPKLERPKLYKVMLLNDDYTP**MS**FVTVVLKAVFRMSEDTGRRVMMT
AHRFGSAVVVVCERDIAETKAKEATDLGKEAGFPLMFTTEPEE

The first amino acid "S" in the ClpS2 St-V1 sequence remains from cleaving off the N-terminal TEV cleavage site: (ENLYFQ/S) with Tobacco Etch Virus (TEV) protease.

3. Sequence information for tvClpS2 Q31H

*Thermosynechococcus vestitus* ClpS2 Q31H (tvClpS2 Q31H) mutant has the following sequence[37]:

*HHHHHH*-GSSGPSVVPQERQQVTRKHYPNYKVIVLNDDFNTF**H**HVAACLMKYIPNMTSDRA
WELTNQVHYEGQAIVWVGPQEQAELYHEQLLRAGLTMAPLEPE*

where the underlined and bolded residue represent the mutation of Q to H.

#### Supplementary Note 7

In the future, higher affinity NAABs will enable more precise measurements, and perhaps more quantitative characterizations. For our low affinity NAABs, we first conducted our exploration in a series of steps. We first assumed that if the binder is added in excess of peptide in the gel, and incubated long enough, it will reach equilibrium, with a binder-bound fraction (and by consequence, concentration) highest in a gel containing the peptide bearing the on-target N-terminal amino acid (see **Supplementary Note 8.2** for equations describing the kinetics of the binder in this condition against different targets). Then, washes to remove binder still in solution, on the order of a couple of minutes, would not have time to wash out all of the binder from the gel containing the on-target peptide, because of the nonzero time required for binder to diffuse out of the gel, even with a short dwell time, with an estimate of > ~70% binder remaining in the gel after a 1-3 min wash (see **Supplementary Note 8.3** for full calculations; this is framed as a lower bound because while we do take into account the slowing of diffusion from the gel itself[38], we do not take into account on-target binding, which would presumably keep more binder in the gel). Then, the gel, containing some fraction of binder within, which would be higher in the on-target gel, is immersed in a solution containing fluorescent antibody against HA-tag, keeping the overall volume small (i.e., a 1.4 µL gel is placed in 30 µL antibody solution), so that even as binder flows out from the gel into the solution, there is still a nonzero fraction of the binder in the volume of the gel labeled by the fluorescent antibody (see **Supplementary Note 8.4**). Finally, the external solution is removed, and the gel imaged. The net result is that some fraction of the NAAB will be retained within the gel, dependent on the initial high fraction of binder bound in the first step, despite the external solution exchanges, and that will result in a greater fluorescence in the gel, accordingly. The prediction is a final expected ~7 fold higher fluorescence intensity in gels with peptide bearing on-target Phe N-terminal amino acid vs. secondary substrates Tyr or Trp (presumably Ala would show no binding, which is why for the purposes of this calculation we compared to Tyr and Trp; see **Supplementary Note 8.4** for full calculations). As noted, the assumptions above are perhaps overestimates of loss of binders from the gel, and thus the fluorescences observed should be regarded as a lower bound on the actual amount; that said, the data should be regarded generally qualitatively, although of course comparisons of different conditions that use the same binder, should be possible.

Having established the assumptions, calculations, and bounds governing our experiment, we next sought to address practicalities - the actual properties of diffusion of binder and antibody in the gel, in relation to the peptide concentration in the gel, and other key experimental parameters like gel thickness. To this end, we tested a concentration of homogeneously distributed peptide of 5 mM, a concentration much higher than would be in an ultimate >100x expanded cell (indeed, it approximates the density in a living cell, to order of magnitude[39]), in 9% acrylamide gels, with a thickness of ~310 µm and volume of ~1.2 µL, and a binder concentration of 20 µM (well above the $K_d$ of 1.1 µM for Phe). The gel was incubated with 30 µL of the ClpS2 St-V1 binder for an hour at room temperature, followed by a short wash of ~1 min, then was incubated in 1.3 µM antibody overnight at 4 °C in 100 µL. Since wild-type ClpS2 binds N-terminal Phe with highest affinity, binds Trp and Tyr with reduced affinity, and does not bind N-terminal Ala[36], we chose to compare binding of this one binder to all four N-terminal amino acids in acrylamide gels. We cast gels with N-terminal Phe, Trp, Tyr and Ala peptides anchored to gels on their C-termini. The result of this experiment showed differential fluorescent intensity across N-termini peptides, confirming that this experimental design can observe population read-out differences in binding. However, the differential signal was only visible on the edges of the gel, and not in the middle (**Supplementary Figure 7A**): the edge-to-center ratio for Phe was ~12, but ~1 for Trp, Tyr and Ala. We hypothesized that, as in earlier studies of binder diffusion into dense samples[40], there was a lack of diffusion of ClpS2 St-V1 or HA-tag antibody into ExMre gels, because the dense amount of Phe-bearing peptide target on the edge of the gel soaked up all available tags (being in excess), before they could get into the center (which of course was not a problem if there was no binding, e.g. for Trp, Tyr, and Ala). We reasoned that this depletion could be ameliorated by reducing the concentration of peptides in the gels (similar to expanding a biological sample). Further, longer incubations of binder and antibody might also support protein diffusion into the core, as long as depletion was not an issue. Nevertheless, this experiment was still valuable since it was consistent with the NAAB binding to its target in an amino acid-specific way.

We next sought to explore more practical concentrations of peptide, e.g. such as might occur after expansion, and less than the binder concentration, to avoid binder depletion. The anti-HA tag antibody was the most likely to encounter a diffusion constraint, based on the Stokes-Einstein relation: larger molecules have larger diffusion coefficients, and slower diffusion flux (estimated ~2.5 fold higher diffusion coefficient for the binder compared to the antibody; see **Supplementary Note 8.1** for calculations). We tested HA-tag antibody diffusion on a set of acrylamide gels with varying HA tag concentrations, with fixed incubation time (O/N at 4 °C), and with an antibody concentration of 0.13 µM. Gels showed fluorescent intensity higher on the edges, than in the center, when the peptide concentration was >=100 µM, but at 10 µM concentration the signal was detected throughout (**Supplementary Figure 7B**). This suggests that the depletion of antibodies occurs on the edges of the gels due to an abundance of binding sites (peptides) when above ~10 µM. Based on this result, and considering the estimated difference in diffusion coefficient of ClpS2 St-V1, a ~4 hour incubation at room temperature might be expected to enable similar diffusion for the binder throughout the gel (since $t_{diffusion} \propto \frac{1}{D}$, see **Supplementary Note 8.1** for calculations). We next determined a finalized experimental protocol for ExMre gels. The final thickness of the ExMre gels was ~350 µm, and the final concentration of peptide ~50 µM. As a result, we increased the ClpS2 St-V1 incubation time to 6 hours at room temperature, and the antibody incubation to 7 days at 4 °C to let proteins diffuse throughout the gel. In addition, to save on protein binder material, the binder concentration was cut to 10 µM (still above the on-target $K_d$ value), which we estimated to enable a high fraction bound after equilibrium (calculated ~90% for on-target, see **Supplementary Note 8.2**). This protocol was used to test ClpS2 St-V1 and HA-tag antibody binding in the gels, comparing different N-terminal amino acids as before (**Supplementary Figure 7A**), but with the above modifications to observe signal throughout the gel.

#### Supplementary Note 8

Adapting from "The mathematics of diffusion"[41], we calculate an estimate as follows:

1. Protein radii (assuming spherical shape) and diffusion coefficients ($D$) of ClpS2 St-V1 and anti-HA antibody:

Assuming a protein density of ~1.35 g/cm$^3$[42] and molecular weight of 13,034 g/mol for ClpS2 St-V1 and ~150,000 g/mol for anti-HA antibody:

$$Volume_{ClpS2\ St-V1} = \frac{mass}{density} \simeq \frac{13{,}034\ g/mol \div 6.022*10^{23}\ molecules/mol}{1.35\ g/cm^3} \simeq 16.03\ nm^3$$

$Radius_{ClpS2\ St-V1} \simeq 1.56\ nm$ (using the exact result from the volume calculation)

$$Volume_{anti-HA\ antibody} = \frac{mass}{density} \simeq \frac{150{,}000\ g/mol \div 6.022*10^{23}\ molecules/mol}{1.35\ g/cm^3} \simeq 184.51\ nm^3$$

$Radius_{anti-HA\ antibody} \simeq 3.53\ nm$ (using the exact result from the volume calculation)

Abbreviations: diffusion coefficient ($D$ in m$^2$ s$^{-1}$), Boltzmann constant ($k_B$: 1.380649*10$^{-23}$m$^2$ kg s$^{-2}$ K$^{-1}$), temperature ($T$: 298 K for 25 °C and 277 K for 4 °C), dynamic viscosity (η: 0.89*10$^{-3}$ kg m$^{-1}$ s$^{-1}$ at 25 °C and 1.57*10$^{-3}$ kg m$^{-1}$ s$^{-1}$ at 4 °C) and Stokes radius ($r$ in m). We assume free diffusion in solution, in water:

$D_{ClpS2\ St-V1} = \frac{k_B T}{6\pi\eta r} \simeq 1.57 * 10^{-6}\ cm^2\ s^{-1}$ (calculated at room temperature, 25 °C, assuming $r$ = 1.56 * 10$^{-9}$ m)

$D_{anti-HA\ antibody} = \frac{k_B T}{6\pi\eta r} \simeq 3.66 * 10^{-7}\ cm^2\ s^{-1}$ (calculated at 4 °C, assuming $r$ = 3.53 * 10$^{-9}$ m)

$$Ratio_D = \frac{D_{ClpS2\ St-V1}}{D_{anti-HA\ antibody}} \simeq 4.3$$

Where $Ratio_D$ represents the ratio between the ClpS2 St-V1 and anti-HA antibody diffusion coefficients, which is directly proportional to their difference in rate of diffusion (for a fixed distance: $t \propto \frac{1}{D}$).

Thus, if O/N at 4 °C is needed for the antibody diffusion ( ~16 hours), then ~4 hours is needed for ClpS2 St-V1 binder diffusion at room temperature.

2. Kinetics of ClpS2 St-V1; fraction bound at equilibrium:

At equilibrium (i.e., when $t \to \infty$), the fraction of peptides bound ($f_{b,\ equilibrium}$) depends on the concentration of ClpS2 St-V1 (i.e., $[ClpS2\ St - V1\ binder]$) and its dissociation constant (i.e., $K_d$) towards the target:

$$f_{b,\ equilibrium} = \frac{[ClpS2\ St-V1\ binder]}{[ClpS2\ St-V1\ binder] + K_d}$$

This assumes that $[ClpS2\ St - V1\ binder]_{free} \simeq [ClpS2\ St - V1\ binder]_{total}$ since the binder is added in excess compared to the peptide (e.g., in **Fig. 4**, ~4.3x molar excess binder compared to peptide in the gel).

- For Phe – Assume $K_d$= 1.1μM (data for ClpS2 St-V1 against FRVECK-biotin) from [33]. Assume $k_{off} = 0.1\, s^{-1}$ (data for ClpS2 V1 against FGVECK-biotin) from [32].
- For Trp – Assume: $K_d$= 11.2 μM (data for ClpS2 St-V1 against WRVECK-biotin) from [33]. Assume $k_{off} = 0.76\, s^{-1}$ (data for ClpS2 V1 against WGVECK-biotin) from [32].
- For Tyr – Assume: $K_d$= 11.6 μM (data for ClpS2 V1 against YGVECK-biotin) from [32]. Assume $k_{off} = 0.5\, s^{-1}$ (data for ClpS2 V1 against YGVECK-biotin) from [32].

$$f_{b,\, equilibrium - Phe} = \frac{10\ \mu M}{10\ \mu M + 1.1\ \mu M} \simeq 90\%$$

$$f_{b,\, equilibrium - Trp} = \frac{10\ \mu M}{10\ \mu M + 11.2\ \mu M} \simeq 47\%$$

$$f_{b,\, equilibrium - Tyr} = \frac{10\ \mu M}{10\ \mu M + 11.6\ \mu M} \simeq 46\%$$

3. Diffusion of proteins out of the gel:

To calculate the percent of binders that diffuse out of the gel, we model the gel as a 3-d rectangular prism, where the binder is initially confined in the gel with dimensions 2 x 2 x 0.35 mm.

In a 1-d system, where the binder is confined in a region of $- h < x < + h$ and assuming a constant diffusion coefficient [41], this is described by the equation, from Fick's second law of diffusion:

$$C(x,t) = \frac{1}{2} C_0 \{erf \frac{h-x}{2\sqrt{Dt}} + erf \frac{h+x}{2\sqrt{Dt}}\}$$

where $erf$ is the error function defined as:

$$erf\, z = \frac{2}{\pi^{1/2}} \int_0^z exp(-\eta^2)\, d\eta$$

and where $C(x,t)$ is the concentration at point $x$ at time $t$.

In the 1-d case, the initial total amount of binder present in the gel before diffusion ($n_{start-1d}$), is given by:

$$n_{start-1d} = C_0 * 2h$$

where $C_0$ is the initial concentration.

The total amount of binder present in the gel, after diffusion from the wash is $n_{wash-1d}$, and the fraction of binder starting material still in the gel ($M_{1d}$) after a defined time, $t$ in seconds is then:

$$n_{wash-1d}(t) = \int_{-h}^{h} C(x,t)\, dx$$

$$M_{1d}(t) = \frac{n_{wash-1d}(t)}{n_{start-1d}}$$

Thus,

$$n_{wash-1d}(t) = \int_{-h}^{h} \frac{1}{2} C_0 \{erf \frac{h-x}{2\sqrt{Dt}} + erf \frac{h+x}{2\sqrt{Dt}}\} dx = 2C_0 \sqrt{Dt}\left(\frac{h}{\sqrt{Dt}} erf(\frac{h}{\sqrt{Dt}}) + \frac{e^{-h^2/Dt}}{\sqrt{\pi}} - \frac{1}{\sqrt{\pi}}\right)$$

$$M_{1d}(t) = \frac{n_{wash}(t)}{n_{start}} = \frac{\sqrt{Dt}}{h}\left(\frac{h}{\sqrt{Dt}} erf(\frac{h}{\sqrt{Dt}}) + \frac{e^{-h^2/Dt}}{\sqrt{\pi}} - \frac{1}{\sqrt{\pi}}\right)$$

The solution in three dimensions, for the gel, is the product of the solutions for each dimension:

For x and y dimensions; $h = 1\, mm$ with $- h < x < + h$ and $- h < y < + h$; and in the z dimension, $w = 0.175\, mm$ with $- w < z < + w$

Then:

$$n_{start-3d} = C_0 * 2h * 2h * 2w$$

$n_{wash-3d}(t) = \int_{volume} C(x, y, z, t)\, dx\, dy\, dz$ and $M_{3d}(t) = \frac{n_{wash-3d}(t)}{n_{start-3d}}$

Assuming diffusion process can be separable in Cartesian coordinates, where diffusion along x, y, and z is independent:

$$M_{3d}(t) = \left(\frac{\sqrt{Dt}}{h}\left(\frac{h}{\sqrt{Dt}} erf(\frac{h}{\sqrt{Dt}}) + \frac{e^{-h^2/Dt}}{\sqrt{\pi}} - \frac{1}{\sqrt{\pi}}\right)\right)^2 * \frac{\sqrt{Dt}}{w}\left(\frac{w}{\sqrt{Dt}} erf(\frac{w}{\sqrt{Dt}}) + \frac{e^{-w^2/Dt}}{\sqrt{\pi}} - \frac{1}{\sqrt{\pi}}\right)$$

Assuming a wash of 60 seconds to 180 seconds, with a diffusion coefficient ⅕ of $D_{ClpS2\, St-V1}$ to account for diffusion in the gel compared to diffusion in free solution, based on existing literature [38]. Then, the estimate becomes:

$$M_{3d}(60\, seconds) \simeq 82\%$$

$$M_{3d}(180\, seconds) \simeq 69\%$$

4. Calculations for ClpS2 St-V1 in the gel, after binding (a), after the wash (b), and after fluorescent antibody staining (c)

a) After, the first binding of ClpS2 St-V1: $f_{b,\, equilibrium\, -\, Phe} \simeq 90\%$, $f_{b,\, equilibrium\, -\, Trp} \simeq 47\%$ and $f_{b,\, equilibrium\, -\, Tyr} \simeq 46\%$. This means that in the 1.4 µL gel with 70 pmol peptide, there is then a number of moles ($n_{equilibrium}$) of ClpS2 St-V1 in the gel:

$n_{equilibrium - Phe} \simeq 63\, pmol$ , $n_{equilibrium - Trp} \simeq 33\, pmol$ and $n_{equilibrium - Tyr} \simeq 32\, pmol$

b) After the wash, assuming ~75% of binders remain in all of the gels (from calculation point 3, above): $n_{wash - Phe} \simeq 47\, pmol$ , $n_{wash - Trp} \simeq 25\, pmol$ and $n_{wash - Tyr} \simeq 24\, pmol$

c) In the 30 µL incubation in anti-HA antibody, the concentration of binder ($C_{binder}$) in solution, for each gel, is then: $C_{binder - Phe} \simeq 1.5\, \mu M$ , $C_{binder - Trp} \simeq 0.75\, \mu M$ and $C_{binder - Tyr} \simeq 0.73\, \mu M$ . Then, the fraction bound before imaging is defined by the more complex binding equation (since we cannot work under the $[ClpS2\, St - V1\, binder]_{free} \simeq [ClpS2\, St - V1\, binder]_{total}$ assumption in this case), where $[peptide]_{total} \simeq 2.3\, \mu M$ (treating the gel-bound ligand as if it were free and uniformly distributed since the system is equilibrated):

$$f_b = \frac{([peptide]_{total} + [ClpS2\, St{-}V1\, binder]_{total} + K_d) - \sqrt{\left([peptide]_{total} + [ClpS2\, St{-}V1\, binder]_{total} + K_d\right)^2 - 4*[peptide]_{total}*[ClpS2\, St{-}V1\, binder]_{total}}}{2*[peptide]_{total}}$$

Then:

$f_{b,\, imaging - Phe} \simeq 37\%$, $f_{b,\, imaging - Trp} \simeq 5\%$, and $f_{b,\, imaging - Tyr} \simeq 5\%$.

So approximately ~7 fold higher intensity signal expected in the Phe gels compared to the Trp and Tyr gels.

#### Supplementary Note 9

This Supplementary Note serves to give more information on the discussion and analysis of oxidation of amino acids in the process of free-radical polymerization, as part of a broader discussion on N-terminal amino acid binders for **Fig. 4**.

As proteins being attached to polymerized ExM gels are experiencing a free-radical filled environment, we examined the susceptibility of amino acid side chains to oxidation as a result of free-radical polymerization. Methionine and cysteine, sulfur-containing side chains, are most prone to free radical oxidation; aromatic amino acid side chains (Tyr, Trp, Phe), and some others (His, Pro, Lys, Arg) can also be oxidized, although they are less prone[43,44]. Thus, the amino acid side chains of Met, Cys, Tyr, Phe, Trp, His, Pro, Arg, at the N-terminus of an otherwise identical peptide chain, were tested for known oxidation products after free-radical polymerization in ExM gels. We here aimed only to gauge the presence or absence of an oxidation product, and, thus, analyzed only one batch of gels and one peptide aliquot of each N-terminal peptide. Each peptide and its ratio to oxidation products was obtained by analyzing relative abundances via LC/QToF (see **Source Data** for raw traces and spectra). The known oxidation products of each amino acid side chain were determined and listed in **Supplementary Figure 8A-Hi**. The raw area under the curve for each species (oxidized and non-oxidized) was documented for the following three conditions: (1) in solution with trypsin, a non-oxidative environment; (2) in solution with trypsin and ammonium persulfate (APS) and tetramethylethylenediamine (TEMED), which should capture the key environment that causes oxidation (since APS decomposes to sulfate radicals, and TEMED accelerates the decomposition of APS); (3) and ExM gel followed by trypsinization, which should similarly recapitulate the oxidative environment. Finally, we plotted the max-normalized area under the curve of the chromatogram, representing the relative abundance of each of the species. Note that the abundance of the various products cannot be compared directly, due to possible differences in ionization efficiency. However, the ratio of oxidized/unoxidized peptide (for a given oxidation product) can be estimated.

In summary: after ExM polymerization, the ratio of methionine sulfoxide/original methionine peptide went from ~0 (in non-free radical solution) to ~4 (in ExM gels) (**Supplementary Figure 8A**). The ratio of sulfinate/original cysteine peptide went from ~0 (in non-free-radical solution) to ~6 (in ExM gels) (**Supplementary Figure 8B**). On the other hand, all other amino acid side chains had a ratio of oxidation product/original peptide below ~0.07, for all oxidation products analyzed (**Supplementary Figure 8C-H**; see **Supplementary Table 11** for descriptive values). Since, in an *in situ* peptide sequencing workflow, amino acid oxidation might prevent recognition of binder reagents to the N-terminus amino acid (assuming the binders are specific to the non-modified side chain - of course, if an oxidation product is reliably produced in an ExM gel, one could simply make binders specific to the modified amino acid), side reactions, such as oxidation, might need to be regulated. Strategies might be adopted to prevent or recover oxidized residues. For instance, methionine sulfoxide can in principle be converted back to methionine by methionine sulfoxide reductase. For cysteine, reducing agents (i.e., TCEP, or DTT) can reduce disulfide bridges and thioredoxin can convert cysteine sulfenic acid (RSOH) back to a thiol group[43]. Nevertheless, some oxidation states of cysteine (i.e., sulfinate $RSO_2$, or sulfonate $RSO_3$) are likely irreversible[45]. Mitigation of side chain modifications from oxidation may benefit from the use of alternative gel polymerization strategies. Indeed, expansion microscopy has been demonstrated using a monomer that forms the ExM polymer network through click chemistry, which would be bioorthogonal to the amino acid detection questions at hand[46,47].

#### Supplementary Note 10 - A theoretical assessment of *in situ* protein sequencing

For our theoretical assessment of *in situ* protein sequencing, we decided to focus on the case of N-terminal binding to an amino acid while still attached to a peptide, as a conservative case for a binding event. Previous modeling approaches have explored how well a given *ex situ* single molecule protein sequencing or fingerprinting approach would perform in terms of proteomic coverage, using various proxies or metrics[48–54]; we here sought to develop such a framework for *in situ* protein sequencing.

*In situ* protein sequencing, using ExM chemistries and in-gel Edman degradation, involves stochastic chemical steps that need to be modeled as accurately as possible in order to assess the feasibility of mapping empirical protein sequences to the actual proteome (Supplementary Figure 13**A** for schematic of the whole experimental workflow) - or to be modeled pessimistically, to set a bound on future performance. Common alignment techniques, like BLASTP, are useful for homology detection between proteins, but are not suitable for *in situ* protein sequencing modeling. This is because the error profiles (from mismatches, gaps and deletions) of BLASTP do not consider the chemistries at hand (e.g., fixation, anchoring and digestion) and the read-out errors that are expected (i.e., in-gel Edman degradation reaction efficiencies, number of binders and their specificities), nor the complexity of sequencing multiple fragments from one protein, as expected in *in situ* protein sequencing. We thus set out to build our own model specifically for *in situ* protein sequencing (Supplementary Figure 13**B** for an outline of our modeling strategy).

We chose to perform these analyses on *Mycoplasma genitalium* and human proteomes. The former (referred to as "mycoplasma" hereafter) has the smallest proteome (483 Swiss-Prot reviewed proteins; UniProt ID 243273: by searching "taxonomy_id:243273" in UniProt), and thus could be a useful testbed for *in situ* protein sequencing. The human proteome (20,421 Swiss-Prot reviewed proteins, UniProt ID 9606: by searching "taxonomy_id:9606" in UniProt) is of course key to confronting human disease. We explored how varying the type and success rate of each chemical step (e.g., different anchoring chemistries, different digestion enzymes; % fixation, % anchoring, % digestion) affects the fidelity of the peptide fragments retained in the gel. In particular, we defined "percent amino acid count" as the percentage of residues of a protein that are not modified, and remain accessible for read-out, for 15 rounds of in-gel Edman degradation after chemical preprocessing (these results are contained in the section below, **Theoretical assessment Part 1: results regarding fixation, anchoring, gelation, and digestion**).

For a given parametrization of the chemical steps explored above (i.e., probabilities of 0.05 for fixation, 0.8 for anchoring with AcX, and 0.8 for digestion with trypsin), we subsequently simulated possible outcomes of the series of steps many times, and stored the results in a dataset containing the list of all such peptide fragments (which we call the "reference fragment dataset", representing the ground-truth sequences). Then, we simulated the rest of the workflow, modeling the errors expected with in-gel Edman chemistry, and molecular recognition with NAABs. This led to output reads for every fragment retained in gel, which we call the "error-prone fragment dataset". We assumed prior knowledge of which fragments originated from each parent protein, with no mixing or exchange occurring between neighboring proteins. (Note well, this assumption may not hold for protein complexes that are densely packed, e.g. many receptors that are multimeric, or the ribosome, or other structures - a full treatment of the problem will require additional work.) These assumptions were inspired by recent findings in the literature showing that it is possible to reach ~1 nm resolution, and to identify protein shape with ExM[55]. Fragment read-outs were then independently evaluated for their match to proteins, using the reference fragment dataset previously mentioned. A final assignment to a protein was performed if the number of fragments matching a protein was at least double that of the next best matching protein, a somewhat arbitrary choice (these results are contained in the section below, **Theoretical assessment Part 2: results for NAAB binding and Edman degradation**).

##### Theoretical assessment Part 1: considerations for fixation, anchoring, gelation, and digestion

The first step for a biological specimen involves fixation (Supplementary Figure 13**A**), which stabilizes the cellular architecture and the spatial relationships between proteins[56]. It is often performed with formaldehyde, a cross-linking agent that forms chemical bonds between amino acids, in ExM[11,57–60]. Chemical fixation via formaldehyde involves primarily amine (e.g., N-termini (or N-term for short) and lysines) and thiol groups (e.g. cysteine) forming methylol groups, and a subsequent step involving crosslinking via methylene bridges which can include other amino acid side chains (e.g., arginine, tyrosine, and to a lesser extent asparagine, glutamine, histidine and tryptophan)[56,61]. As such, our model was constructed to modify residues most likely to be fixed (e.g. N-term, lysine, cysteine, arginine, tyrosine), under the assumption that they are modified equally (arbitrarily, since we had no ground-truth data to support our decision), at a given probability (**Supplementary Figure 13B**). Alternative fixation strategies exist that rely on organic solvents, such as methanol or ethanol, which do not form chemical bonds between amino acids on proteins. Such fixation has been demonstrated with ExM[62]. However, this comes with a likelihood of poorer soluble-protein retention[63], and thus higher protein loss. Protein loss is a concern for any fixation strategy, and various fixation strategies and their relative protein retention have been studied, with for instance a decrease from ~65% to ~8% protein loss via one form of hydrogel embedding[64]. These results, and others[65,66], suggest that hydrogel-tethering can retain the majority of proteins under the right conditions. For this reason, we chose not to account for protein loss in the chemical fixation step. This also allowed us to isolate the percent amino acid count (defined above) from fixation parameters that could affect protein loss. It is important to note that protein loss beyond the fixation step was accounted for in our model, in subsequent steps such as anchoring, digestion and in-gel Edman degradation.

Anchoring is the step after fixation (**Supplementary Figure 13A**), and included in ExM to create covalent linkages between proteins and the hydrogel network. For anchoring chemistries, we compared two strategies previously reported in ExM studies, using the bifunctional molecules AcX and acrylate epoxide (which we abbreviate “epoxide”). AcX harbors an NHS (N-hydroxy succinimide) ester moiety that reacts with primary amines (e.g., N-terminus and lysine side chains). We assumed for AcX anchoring that all primary amine groups were modified with equal probability (**Supplementary Figure 13B**). On the other hand, epoxide was assumed to react with multiple side chains (e.g., N-term, lysine, cysteine, histidine, tyrosine aspartic acid, and glutamic acid)[67], leading to different outcomes in fragment retention in the gel. For epoxide anchoring we assumed that all side chains, listed above, were modified with equal probability (not depicted on **Supplementary Figure 13B**). In general, we assumed that protein anchoring to the gel occurs independently for each molecule. We also assume, for the purposes of this model, that amino acids are anchored at the same probability regardless of where they are within the tertiary structure of the protein; of course, this may not be the case in real life. (Note well - in practice we can denature proteins, and/or iteratively anchor and expand, to compensate for such issues.)

After a sample is anchored, the gel is cast. During the gelation step we assumed no change in primary protein sequence. Finally, for digestion (**Supplementary Figure 13A**), we wanted to compare enzymatic cleavage using trypsin (targeting K and R sidechains as shown in **Supplementary Figure 13B**), endoproteinase Lys-C (Lys-C; targeting lysine sidechain), and proteinase K (ProK; targeting aliphatic, aromatic and hydrophobic side chains including Y, W, E, L, V, A, I, F, and T[68–70]). These digestion enzymes are commonly used in ExM to chemically soften the sample and allow for expansion to occur[11,55,59,71,72]. Alternatively, ExM can be performed with a denaturation step (e.g., using beta-mercaptoethanol, SDS and/or high temperature, i.e., 95 °C)[59,73,74]. We did not explore denaturation further in our modeling, since fixation and anchoring, without a downstream digestion step, does not lead to fresh N-termini for sequencing via in-gel Edman degradation. Digestion enzymes expose fresh N-terminal sites that can be targeted by in-gel Edman degradation, generating several fragments of a single protein that can be sequenced in parallel. Sequencing many fragments in parallel would lead to many shots on goal. This, of course, assumes that the peptide fragments derived from the digestion of a single protein are sufficiently separated to be distinguished as discrete fluorescent puncta. We assumed prior knowledge of which fragments originated from each parent protein, with no mixing or exchange occurring between neighboring proteins. During this evaluation, at a given probability of digestion, it was assumed that all amino acids targeted by the enzyme were equally accessible for digestion, unless the residue was anchored to the gel, in which case the enzyme was modeled to be unable to cleave the downstream amide bond (due to specific side-chain recognition loss for the various enzymes[68–70,75–77]). All other amino acids were assumed available given that proteins can be denatured and linearized into peptide fragments prior to digestion and that we had no other ground-truth data to support our decision. Simulation of digestion is depicted in **Supplementary Figure 13B** for trypsin cleavage.

Importantly, we note that all of the chemical parameters for fixation, anchoring, and digestion can be independently modified (not just the probability values of the parameters, but also the reagent, thus affecting amino acid specificity at each step) within our code (**Methods: Theoretical assessment of in situ protein sequencing** section for code) to test alternatives.

**Theoretical assessment Part 1: results regarding fixation, anchoring, gelation, and digestion**

As mentioned above, we defined "percent amino acid count" as a metric to assess the number of amino acids that remain accessible for downstream sequencing for a given set of values in the chemical parameter space. We used this metric to determine the mean percent amino acid count, which is the percent amino acid count averaged over all proteins of the proteome, for several combinations of values in the parameter space. In this analysis, we assumed that for the human proteome, the first fragments of proteins were inaccessible for sequencing, since N-terminal acetylation affects ~70-80% of proteins[78], thus preventing the initiation of Edman degradation on original N-termini. However, this was not assumed for the mycoplasma proteome, since, although the extent of this post-translational modification (PTM) is not well characterized in *Mycoplasma genitalium*, N-terminal acetylation is significantly less common in bacterial and archaeal proteomes (10-29% N-terminal acetylation for *Mycobacterium tuberculosis* and *Pseudomonas aeruginosa PA14*)[79].

We conducted a systematic overview of how different fixation rates, anchoring rates for two anchoring methods (AcX, epoxide), and digestion rates for three digestion methods (Lys-C, trypsin, ProK), would affect the mean percent amino acid count in the mycoplasma and human proteomes. We varied the probability of reactivity towards amino acid targets from 0 to 1 (where 0 is no reaction, and 1 is complete reaction, assessed independently at each amino acid in the sequence). The results of this grid search are visualized in a heatmap (**Supplementary Figure 13C** and **Supplementary Figure 18**, for mycoplasma and human proteome results, respectively; raw data CSV files are available in our code on GitHub in **Methods: Theoretical assessment of in situ protein sequencing**). We also analyzed the variability in the percent amino acid count for certain parameters (discussed more below; with distributions plotted in **Supplementary Figure 14**).

Some patterns emerged: increased fixation generally reduced the number of residues available for anchoring to the gel network and led to decreased percent amino acid count in the gel, with the caveat, mentioned above, that we did not model protein loss at low fixation (**Supplementary Figure 13C**, going from left to right; **Supplementary Figure 18** for the same results in the human proteome). The results also showed differences in mean percent amino acid count when comparing AcX and epoxide (**Supplementary Figure 13Ci-iii** for AcX, **Supplementary Figure 13Civ-vi** for epoxide for the mycoplasma proteome; **Supplementary Figure 18i-iii** for AcX, **Supplementary Figure 18iv-vi** for epoxide for the human proteome). For AcX anchoring, fixation directly competed with anchoring since both processes target amines. Thus, as fixation probability increased, anchoring efficiency with AcX decreased, leading to greater fragment loss and less mean percent amino acid count (**Supplementary Figure 13ci-iii, Supplementary Figure 18i-iii**). For epoxide, the effect was not as pronounced because epoxide targets additional residues not modeled to be reactive to fixative (**Supplementary Figure 13Civ-vi, Supplementary Figure 18iv-vi**).

Some anchoring was essential, as no anchoring resulted in complete fragment loss (**Supplementary Figure 13c** and **Supplementary Figure 18**). On the other hand, complete anchoring eliminated all Lys-C cleavage sites and prevented digestion entirely (**Supplementary Figure 13Ci** and **Supplementary Figure 13civ** and **Supplementary Figure 18i** and **Supplementary Figure 18iv**). The best anchoring probability to maximize mean percent amino acid count from our model, represented a balance between these competing effects, with different ideal conditions (based on our assumptions) for AcX and epoxide due to their different amino acid selective reactivities. For instance, for AcX with Lys-C digestion for P(fix)=0.1, the best option was P(anchor)=0.6. However, for epoxide with Lys-C the best condition was P(anchor)=0.4 (i.e, shifted to the left, towards lower anchoring probability). This can be explained since, at a similar reaction anchoring efficiency rate, more residues are modified with epoxide than AcX. Our results suggested that epoxide, on average, was able to retain more mean percent amino acid count in its best conditions than AcX in its best conditions, at least in this theoretical assessment.

We next compared the effects of the three proteases (trypsin, Lys-C and ProK) on the change in percent amino acid count of fragments, and overall fragment lengths. Fragment length was defined as the number of amino acids of a fragment retained in the gel, not considering amino acids that are located downstream (C-terminal) of the last anchoring amino acid of the fragment (since this fragment would be lost after in-gel Edman degradation), but unlike the percent amino acid metric, was not limited to the first 15 amino acids of the fragment. For ProK, a P(digest)=0.2 resulted in the highest mean percent amino acid count. Higher probabilities, above 0.2, resulted in higher fragment loss and digestion leading to an overall lower mean percent amino acid count (Supplementary Figure 13**Ciii** and Supplementary Figure 13**Cvi** for mycoplasma proteome, Supplementary Figure 18**iii** and Supplementary Figure 18**vi** for the human proteome). However, for both trypsin and Lys-C conditions, a maximal probability of digestion, P(digest)=1.0, enabled the highest mean percent amino acid count (Supplementary Figure 13**Ci-ii** and Supplementary Figure 13**Civ-v; **Supplementary Figure 18**i-ii** and Supplementary Figure 18**iv-v**). This is consistent with the fact that ProK has broad specificity against amino acids (9 targets;[68–70]), thus generating many more fragments with free N-termini at a given P(digest) compared to trypsin (2 targets) or Lys-C (1 target) (see fragment lengths in Supplementary Figure 15**A-B** for mycoplasma proteome and human proteome, respectively). Taken together, these results suggested ProK could reach its highest mean percent amino acid count by tuning its probability of digestion to P(digest)=0.2, but above this value would start to lead to fragment loss and very short fragments (e.g. ~2 amino acids in median length with P(digest)=0.8) leading to a reduction in mean percent amino acid count. For Lys-C, the median fragment length was above 15 amino acids with P(digest)=0.8, suggesting incomplete coverage of the retained fragments with only 15 rounds of in-gel Edman degradation. In addition, for Lys-C and trypsin, given that their best condition was at their maximal digestion capabilities (P(digest)=1.0), both of these enzymatic strategies may not be reaching ideal fragment lengths for maximal coverage over 15 rounds of in-gel Edman degradation.

Finally, we looked at the distribution of percent amino acid count in the gel for every protein for a given set of the chemical parameters, to have a sense of the variability of this metric. In the case of digestion with Lys-C with either AcX or epoxide anchoring (P(fix)=0.05, P(anchor)=0.8 with AcX or epoxide, and P(digest)=0.8 with Lys-C), there were 4% and 11% of the proteins in the mycoplasma and human proteomes, respectively, with <2.5% amino acid count in the gel (Supplemental Figure 14**A,** two leftmost panels, for the mycoplasma proteome and Supplemental Figure 14**C,** for the human proteome). This percentage of proteins was therefore mostly lost in the process of anchoring or digestion. On the other hand, with the same P(fix), P(anchor) and P(digest) values, epoxide with trypsin led to the highest median percent amino acid count in the gel, where close to all proteins showed a percent amino acid count >2.5% (Supplemental Figure 14**A,** bottom rightmost panel, where mycoplasma has ~1.2% of proteins and Supplemental Figure 14**C** human has 0.6% of proteins with <2.5% amino acid count in the gel). We also analyzed the distribution for the percent amino acid count for the best parametrization (i.e., the one that maximized the mean percent amino acid count) for each condition in Supplemental Figure 14**B** and **D** (mycoplasma, human). In this case, all conditions' distributions were shifted towards higher mean percent amino acid count and fewer proteins had <2.5% amino acid count. However, for the human proteome, unlike the mycoplasma proteome, Lys-C digestion with AcX or epoxide still led to >2% of proteins with <2.5% amino acid count (the mycoplasma proteome showed <1% proteins with <2.5 % amino acid count in the same conditions). This discrepancy arises from modeling human proteins with capped N-termini from N-terminal acetylation, unlike those in the mycoplasma proteome, combined with the limited ability of Lys-C digestion to generate sufficient new free N-termini (Supplemental Figure 14**B** for mycoplasma, and Supplemental Figure 14**D** for human).

**Theoretical assessment Part 2: considerations for NAAB binding and Edman degradation**

For simulating *in situ* protein sequencing with in-gel Edman degradation and NAAB sequence read-out, where we assume that NAABs bind PITC-modified N-terminal amino acids. We assumed this since this approach would yield a read-out similar to that of NAABs targeting native N-terminal amino acids, except that PITC conjugation at the N-terminus could enhance NAAB dwell times required for reliable imaging detection. We selected probability values for each of the fixation, anchoring and digestion steps explored above. Specifically, we selected 0.05 for fixation, AcX at 0.8 probability of anchoring and 0.8 probability of digestion with trypsin. We made the 0.05 fixation selection, since amino acid side chain modifications on the order of ~3-22% conversion have been observed when incubating peptides with 50 times excess formaldehyde, at pH 7.2 and 35 °C for 48 h[56,61]. (Again, we did not consider protein loss related to incomplete fixation.) AcX is the most commonly used anchoring chemistry used in ExM protocols; although the true percentage of amine groups modified by AcX is unknown, NHS esters react selectively and efficiently with primary amines. Due to potential limitations on reaction probability by tertiary protein structure in a cell, we considered P(anchor)=0.8 when modeling the expected probability of AcX reaction to amines. Finally, for digestion, although the true value of digestion efficiency is unknown in-gel, it has been shown that trypsin digestion efficiency is ~80% in samples from yeast total protein extract analyzed with mass spectrometry[80]. Thus, we considered P(digest)=0.8. Since epoxide and ProK, rather than AcX and trypsin, showed the strongest performance (these results are contained in **Theoretical assessment Part 1: results regarding fixation, anchoring, gelation, and digestion**), the model might be expected to perform less well with AcX and trypsin than with epoxide and ProK, when considering their respective best parametrization.

Using this selection, we constructed an indexing structure (trie) that captures the space of different fragments that could be generated and retained in the gel, which we also call our reference fragment dataset (Supplementary Figure 13**B**). We simulated the experiment 1,000 times (to be conservative; varying the number of runs from 1 to 10,000 did increase the number of unique fragments that were added to the reference fragment dataset; Supplementary Figure 16). Consequently, selecting only 1,000 simulations constrained the reference fragment dataset, likely underestimating our ability to map back to the proteome. Using this approach, we sought to determine whether it was still possible to map a significant portion of the proteome.

In terms of modeling errors with in-gel Edman chemistry, we modeled conjugation with 10% PITC failure, and cleavage of N-terminal amino acid with TFA as 30% failure (Figure 3**G** suggesting ~70% yield). In addition, we assumed that in-gel Edman degradation can proceed past fixed or anchored amino acids. Indeed, previous work has demonstrated Edman degradation proceeding through bulky side-chains, including covalent modification of side-chains with fluorophores[17], arginylation of the side chain of acidic amino acids[81] and a 30 amino-acid polyproline linker between the side chain and the fluorophore[52], suggesting that this may be a realistic assumption.

In terms of the read-out with the NAABs, we assumed that binders cannot bind to fixed, anchored, or unconjugated (i.e. not bound to PITC) amino acids. To model the binding of NAABs for read-out, we evaluated how increasing the number of NAABs (targeting 5, 10, 15 or 20 amino acids), and increasing the specificity of the NAABs, impacted protein identification. PTMs, present on certain amino acid side chains (most abundantly lysine, tyrosine, asparagine, threonine, and serine), were not taken into consideration. However, since ~0.2% of side chains carry a PTM (~300k experimentally validated modifications out of ~190M amino acids, as found for the Swiss-Prot database[82]), we reasoned that this assumption would not alter the main results of our model. Modifications to amino acid side-chains by free-radical polymerization were not considered since results suggest that oxidation may remain minimal on most sidechains, except cysteine and methionine (Supplementary Figure 8**A-H**), and also because alternative chemistries exist that avoid such side reactions, such as tetra-gel ExM, which does not rely on free-radical polymerization[46,47]. Binders were assumed to bind PITC-conjugated N-terminal amino acids (with each having a specificity towards one amino acid). To simulate a range of specificities and affinities, we constructed a binding affinity matrix shown in Supplementary Table 12, for “perfect”, “very high”, “high”, “medium”, “low” and “very low” binding profiles. Specifically, perfect binding was used as a theoretical limit of the read-out, modeled as a binary outcome for on-target versus off-target. The medium binding scenario was modeled as a binder with 500x longer dwell time against the on-target compared to the off-target, where the on-target binding affinity has $K_d$ ~50 pM , $k_{off}$ ~5 * $10^{-5}$ $s^{-1}$ (as found for some high-affinity antibodies[83]), whereas off-target amino acid affinity was modeled as $K_d$ ~25 nM, $k_{off}$ ~2.5 * $10^{-2}$ $s^{-1}$. The low binding scenario was modeled with on-target kinetics as before, but only assumed a 50x longer dwell time for on-target compared to off-target binding ($k_{off}$~2.5 * $10^{-3}$ $s^{-1}$). Finally, for very low affinity, we similarly modeled on-target kinetics as before, but only assumed a 5x longer dwell time for on-target compared to off-target binding ($k_{off}$~2.5 * $10^{-4}$ $s^{-1}$). During the simulation, we performed sequential binder read-out (where the binders are not added simultaneously, but one at a time). We assumed that for each binder, the kinetics of binding reached equilibrium conditions prior to washing. In addition, we did not consider any specific effects that the gel environment would have on the kinetics of binding, nor any non-specific background from binders bound non-specifically to the gel or not washed out. To understand how binder concentration and wash time could impact read-out under these assumptions, we plotted heatmaps showing how different conditions influence difference in probability of on-target binding and off-target binding for very high, high, medium, low and very low specificity binders (**Supplementary Figure 17** for these heatmaps). Assuming an excess of binder compared to targets, and from these results in **Supplementary Figure 17**, we selected a 1 μM binder concentration because it fell in the range of values with a large difference between probability of on versus off-target binding (large differences could be noted for concentrations between 10 nM and 1 mM). Also since the binder is assumed to have high affinity, 1 μM is well above its $K_d$ value of 50 pM. In addition, even if the concentration of target (i.e. an N-terminal amino acid) has a higher concentration than 1 μM in the gel, since the target is immobilized in the gel, the solution volume is assumed to be in large enough excess that the binder depletion is negligible for the Langmuir equation to hold (**Supplementary Figure 17** caption for the equations describing the binding kinetics). We selected a 30 min wash prior to imaging since this fell in the range of ~0.8 for the difference in probability for on versus off target binding and can be a realistic timing for a wash step (although diffusion may pose a constraint during a 30 min wash, we did not model this further). For the sequential binder read-out, we assumed that we can successfully wash out the binders from their targets by stripping them from the gel, and perform a second round of binding and imaging (as performed previously[73]).

Using the parameters described above, we generated the error-prone fragment dataset (in contrast to the reference fragment dataset that did not account for readout errors) through Monte Carlo simulation. Monte Carlo simulation captures the stochastic realization of fragments generated from imperfect chemical steps, and was a natural way to generate synthetic data for the error prone fragment dataset. Building on previous single-molecule sequencing studies that have used Hidden Markov Models (HMMs)[49,51], we mapped the resulting fragments back to the reference fragment dataset using a HMM. HMMs in this context predict the likelihood that a sequence of hidden states (the true amino acids of a protein) produce the observed data (noisy amino acid readout from correct plus incorrect binding). We first remove any “uninformative fragments”, here defined as sequences with more than half unknown reads (“X”), from both the error-prone fragment and reference fragment database. Then, in the first stage of the HMM, we applied the Viterbi algorithm to each error-prone sequence to efficiently identify the single most likely peptide sequence given the observed binder readouts and modeled failure modes; this choice maximized computational speed and paralleled the pruning strategies used by Smith *et al.* for rapid path selection[51]. This dynamic programming approach traces the most likely sequence of hidden states, which we then use to remove candidate sequences whose log-likelihoods are less than about 1/150 as probable as the top sequence. In the second stage of the HMM, we used the forward algorithm to evaluate the top candidate alignments. Unlike Viterbi, which returned only the most likely path, the forward algorithm summed over almost all possible paths to compute the overall likelihood of the observed fragment from the reference fragments. This provided a probabilistic score that captured uncertainty across alternative fragment assignments and allowed us to rank candidate proteins. Unlike previous implementations, our HMM explicitly integrated binding‑specificity parameters and chemically derived error rates at each degradation cycle, relevant for the unique challenges of our *in situ* protein sequencing platform, which requires evaluation of how experimental variables influence the overall protein identification accuracy (i.e., including the chemical steps: fixation, anchoring and digestion and Edman degradation and readout inefficiencies). We chose to run the simulation with in-gel Edman degradation from 5 to 15 cycles, as the median length of a fragment is ~15 with the same parameters described above (Supplementary Figure 15**A-B**). The specific amino acid binders that were selected for smaller subsets (5, 10 and 15 binders) were chosen arbitrarily and the list of binders in those subsets are noted in the captions of Supplementary Figure 13**D**.

For the final assignment of fragments to a parent protein, a fragment was assigned only if the highest-scoring protein, computed as the sum of the weighted log-likelihoods across the top ten candidate fragments, had a total score at least 1.5x greater than that of the second-highest-scoring protein, otherwise it was "uncertain". At the protein level, a parent protein was considered identified if the total number of its assigned fragments was at least 1.5x the number of fragments supporting the next-highest-scoring protein, otherwise it was "uncertain". We recorded if each identification was "correct", "false positive", or "uncertain". See **Methods: Hidden Markov Model (HMM) based matching and fraction of proteome correctly identified** for more details.

**Theoretical assessment Part 2: results for NAAB binding and Edman degradation**

In Supplementary Figure 13**D**, we quantified the percent identified proteins from the mycoplasma proteome as the fraction of proteins for which our model's top candidate matched the true source protein. We plotted the "fraction correct" of the total mycoplasma proteome, defined as the number of proteins identified as correct (see above, i.e. not false positive and not uncertain; see **Methods: Hidden Markov Model (HMM) based matching and fraction of proteome correctly identified** for more details) divided by the size of the proteome, averaged across 10 iterations. Each of the four panels shows how this fraction correct changes as we increase the number of Edman rounds, under four binder‑specificity regimes (perfect, medium, low, and very low) and four different binder library sizes (20, 15, 10, and 5 amino acid binders). From Supplementary Figure 17 and Supplementary Table 12, we note that high and very high specificity binders under chosen wash times of 30 minutes and concentrations of 1 μM resulted in similar on and off-target binding probabilities, and thus were not plotted and concluded to be comparable to medium specificity. With 20 amino acid binders, perfect, medium, and low specificity converged to >90% accuracy by round 8, and reached 98-99% accuracy by round 12, whereas very low specificity remained at almost zero. Dropping to 15 amino acid binders looked very similar, with convergence of perfect, medium, and low specificity at >90% accuracy by round 8, and reaching ~98% accuracy by round 12. At 10 amino acid binders, perfect and medium specificity binders required 12 rounds to hit ~90%, while low specificity was roughly ~85% at 12 rounds. Finally, with only 5 amino acid binders, even perfect specificity binders struggled to pass ~60% identified proteins after 15 rounds, and medium and low specificity remained below ~60%.

We observed a plateau for perfect binding even with all 20 amino acid binders, with ~99% rather than 100% correctly identified proteins. This plateau occurred since some proteins have a low number of lysine residues that can be anchored to the gel matrix, resulting in lost proteins and/or few retained fragments. An example is protein Uniprot accession P47377 in the mycoplasma proteome with only 2 lysines. Thus, increasing the number of amino acid binders may yield diminishing returns beyond 15 distinct amino acid targets, where > 90% of proteins can be correctly identified after 8 in-gel Edman rounds for both 15 and 20 amino acid binders across perfect, medium, and low specificity binders (Supplementary Figure 13**Di-ii**). We reiterate that in this model, all on-target binding is considered to have a $k_{off}$ of $5 * 10^{-5}\ s^{-1}$ irrespective of the specificity (Supplementary Table 12). Moreover, the magnitude of the $k_{off}$ difference between on-target and off-target binding was critical: a 50-fold difference enabled correct protein identification, whereas a 5-fold difference did not, as reflected in the loss of identified proteins at very low specificity. There were diminishing returns in higher specificities, as noted by the similarity of curves in medium (500x difference between on-target and off-target in $k_{off}$) and low (50x difference) specificity (Supplementary Figure 13**Di-iv**). Increasing the number of in-gel Edman rounds resulted in higher percentages of correctly identified proteins. Finally, with the smaller binder library sizes (5 and 10 NAABs), increasing the number of Edman rounds to 15 did not fully rescue the percent proteins correctly identified, reaching ~90% for low specificity binders with 10 NAABs, and ~50% for medium specificity binders with 5 NAABs.

Finally, we also wanted to look at the percent identified proteins from the human proteome when performing rounds of degradation and detection. To do so, we tested medium specificity binders with 15 NAABs over 5 to 15 rounds of in-gel Edman degradation with 10% PITC failure, and cleavage of N-terminal amino acid with TFA as 30% failure. We did this modeling once, after performing 10 simulations to build the reference fragment dataset, and these results suggested ~80% correctly identified proteins after 11 rounds of degradation (see **Supplementary Figure 20** for this plot, **Supplementary Figure 21** for the false positive rate across rounds).

Together, these curves map out an estimate of how certain characteristics of *in situ* protein sequencing impact the percentage of correctly identified proteins, under the assumptions of our model. The results could inform experimental directions, such as which chemical steps require the most attention, in the future. The most critical development awaiting, of course, is the creation of new binders. As underlying chemistries are updated, to achieve more realistic simulations, we will need to update the assumptions of our model, to better constrain the parameters by empirical data. Once we do have sufficient empirical data, a more straightforward strategy would be to use standard machine learning techniques.

#### Supplementary Note 11

**Iterative 130-190x Expansion Protocol**

This protocol enables 150x iterative 2-round expansion with N,N-dimethylacrylamide (DMAA) gel

A) First DMAA Expansion ~16x

**Manufacture hydrophobic glass slides and coverslips**

Steps 1-4 are performed in a chemical fume hood with proper PPE at room temperature (~24 ˚C).

1. Add 20 µL trichloro(octadecyl)silane to 10 mL hexane.
2. Immerse glass slides and coverslips in the solution for 90 seconds.
3. Remove glass slides and coverslips from the solution with a tweezer.
4. Rinse the glass with 70% isopropanol and $ddH_2O$ sequentially.
5. Place glass inside a 37˚C incubator to dry.
6. Wipe off white residual reactants (expected) with a dry kimwipe.
   *Note: Hydrophobic glass slides and coverslips can, if properly handled, be reused at least 15 times. After each use, wash hydrophobic glass with $ddH_2O$ and gently wipe with kimwipe. These steps do not need to be repeated for every gelation.*

**Set up glove bag**

1. Place the glove bag on a bench.
2. Connect the glove bag to a tube attached to a compressed nitrogen cylinder nearby.
3. Seal the connection between the tube and the glove bag with tape.
4. Fill the glove bag with nitrogen. Then turn off the nitrogen and observe if the glove bag is slowly deflating. If that is the case, the glove bag is not airtight, and a small flow of nitrogen can be provided to keep the bag inflated.

**Gelation**

5. Gelation solution: dissolve 0.522 g sodium acrylate in 1 mL acidified Tris buffer (10% (v/v) 1 M Tris-HCl pH 8 buffer, 20% (v/v) 1.2 M HCl in ddH2O) in a 20-mL glass vial. Vortex.
6. Add 10 µL TEMED to 90 µL $ddH_2O$ in a 1.5-mL centrifuge tube. Vortex.
7. Add 7.5 µL 10% TEMED solution to the 20-mL glass vial.
8. Add 900 µL DMAA to the 20-mL glass vial. Vortex.
   *Note 1: DMAA and subsequent gelation solution are viscous. To ensure accurate volume, pre-wetting the pipet tip is required.*
   *Note 2: The gelation solution should be colorless and non-cloudy. Otherwise the sodium acrylate and DMAA quality is low or has degraded.*
9. Initiator solution: dissolve 45 mg potassium persulfate in 1 mL $ddH_2O$ in a 1.5-mL centrifuge tube to make the initiator solution. Vortex for 2 minutes.
   *Note: potassium persulfate takes time to fully dissolve and will precipitate from the solution if placed on ice. Do not place initiator solution on ice.*
10. Place the gelation solution on ice.
11. Construct the airtight humidified chamber: place a damp towel in the bottom of the airtight chamber. Place a platform on top of the damp towel.
12. Construct the gelation chamber: wrap parafilm strips with size ~4.5 cm × 0.2 cm around the hydrophobic glass slide with 0.4-cm gap.
13. Place the gelation chambers into the airtight humidified chamber. Place a hydrophobic glass slide on each chamber.
14. In a chemical fume hood, connect gas dispersion tube to a compressed nitrogen cylinder.
    *Note 1: The nitrogen flow needs to be kept minimal. Otherwise, the gelation solution will evaporate rapidly and freeze. For first-time users, please practice controlling the nitrogen flow in a 20-mL glass vial filled with 5 mL water to determine the minimal nitrogen flow required to generate bubbles.*
    *Note 2: The sponge head of the gas dispersion tube needs to be fully wetted to generate bubbles.*
15. Immerse gas dispersion tube with flowing nitrogen in gelation solution for 50 seconds in a chemical fume hood.

16. Cap the vial quickly after removing the gas dispersion tube to minimize oxygen exposure. *Note: The 20-mL glass vial is not airtight. Minimize the time between the completion of oxygen removal and placing the vial in the nitrogen-filled glove bag.*
17. Move pipets (P1000, P200, P20), pipet tips, transfer pipets, a tweezer, airtight humidified chamber with gelation chambers, DNA oligos solution, two 1.5-mL centrifuge tubes, gelation solution, initiator solution, and hydrophobic glass coverslips into the glove bag.
    *Note 1: No ice or cold block is needed. All subsequent steps are performed at room temperature. The tissue gelation protocol has been optimized to not gelate for at least 45 minutes upon the addition of initiator solution.*
    *Note 2: Putting pipet tips onto pipets before moving them into the glove bag can reduce tasks performed inside the glove bag.*
18. Turn on the nitrogen flow. Purge the glove bag by filling the glove bag with nitrogen then pushing on top to remove most of the gas within. Repeat purging three times.
    *Note: Ensure airtight chamber is not capped during purges.*
19. Seal the glove bag and turn off the nitrogen flow.
    **Steps 17–22 are performed inside the glove bag**
20. Inside the glove bag, add 411 µL gelation solution and 15 µL initiator solution in a 1.5-mL centrifuge tube. Flip the tube upside down five times for mixing.
21. Slowly add the solution into gelation chambers.
22. Place gelation chambers in the sealed airtight humidified chamber.
23. Remove the airtight humidified chamber from the glove bag. Place it in the dark at room temperature for 2 hours.
24. After incubation, cut out sections of gel into 0.5 x 0.5 cm pieces. For gels containing DNA origami, incubate the gel at 85 ˚C in 1x PBS for 10 minutes.
25. Place sections in a 20-cm petri dish containing 1x PBS. Measure expansion factor. Fully expand sections in $ddH_2O$. Measure expansion factor. Shrink the gel in 1x PBS, cut into 0.5 x 0.5 cm pieces.
26. Store in 1x PBS for up to 1 month.

B) Re-embedding (shrinks linearly to 80% its original size)

1. Place hydrophobic glass slides at the bottom of a 4-well plate.
2. Fully expand a 0.5 × 0.5 cm gel piece in the 4-well plate using $ddH_2O$.
3. Incubate the expanded gel in re-embedding solution (5% (w/v) acrylamide, 1% (w/v) N,N'-Diallyl-L-tartardiamide (DATD), 0.05% (v/v) TEMED, 0.025% (w/v) APS) for 20 minutes with shaking at 50 rpm.
4. Aspirate the solution and replace with fresh re-embedding solution.
5. Use multiple layers of Parafilm as spacers on the glass slide, then cap with a glass coverslip, taking care not to compress or deform the gel. Purge with $N_2$ and incubate at 37°C for 2 hours in a humidified, airtight chamber.
6. Cut the re-embedded gel into 0.5 × 0.5 cm pieces.
7. Store in 1× PBS for up to 1 month (inclusive of any prior storage time).

C) Second DMAA Expansion ~10–12x

The second DMAA gelation is in essence identical to the first gelation. Prepare one standard volume for each gel.

**Gelation for 1 standard volume**

1. Place each gel in 6-well plate without any liquid.
2. Gelation solution: dissolve 0.522 g sodium acrylate in 1 mL acidified Tris buffer (10% (v/v) 1 M Tris-HCl pH 8 buffer, 20% (v/v) 1.2 M HCl in ddH2O) in a 20-mL glass vial. Vortex.
3. Add 10 µL TEMED to 90 µL $ddH_2O$ in a 1.5-mL centrifuge tube. Vortex.
4. Add 7.5 µL 10% TEMED solution to the 20-mL glass vial.
5. Add 900 µL DMAA to the 20-mL glass vial. Vortex.

6. Initiator solution: dissolve 45 mg potassium persulfate in 1 mL $ddH_2O$ in a 1.5-mL centrifuge tube to make the initiator solution. Vortex for 2 minutes.
7. Place the gelation solution on ice.
8. Construct the airtight humidified chamber: place a damp towel in the bottom of the airtight chamber. Place a platform on top of the damp towel.
9. Construct the gelation chamber: wrap parafilm strips with size ~4.5 cm × 0.2 cm around the hydrophobic glass slide with 0.4-cm gap. Use 10 parafilm strips stacked on each side to accommodate the thickness of re-embedded gel.
10. Place the gelation chambers into the airtight humidified chamber.
11. In a chemical fume hood, connect gas dispersion tube to a compressed nitrogen cylinder.
    *Note 1: The nitrogen flow needs to be kept minimal. Otherwise, the gelation solution will evaporate rapidly and freeze. For first-time users, please practice controlling the nitrogen flow in a 20-mL glass vial filled with 5 mL water to determine the minimal nitrogen flow required to generate bubbles.*
    *Note 2: The sponge head of the gas dispersion tube needs to be fully wetted to generate bubbles.*
12. Immerse gas dispersion tube with flowing nitrogen in gelation solution for 50 seconds in a chemical fume hood.
13. Cap the vial quickly after removing the gas dispersion tube to minimize oxygen exposure.
    *Note: The 20-mL glass vial is not airtight. Minimize the time between the completion of oxygen removal and placing the vial in the nitrogen-filled glove bag.*
14. Move pipets (P1000, P200, P20), pipet tips, transfer pipets, a tweezer, airtight humidified chamber with gelation chambers, DNA oligos solution, two 1.5-mL centrifuge tubes, gelation solution, initiator solution, and hydrophobic glass coverslips into the glove bag.
    *Note 1: No ice or cold block is needed. All subsequent steps are performed at room temperature. The tissue gelation protocol has been optimized to not gelate for at least 45 minutes upon the addition of initiator solution.*
    *Note 2: Putting pipet tips onto pipets before moving them into the glove bag can reduce tasks performed inside the glove bag.*
15. Turn on the nitrogen flow. Purge the glove bag by filling the glove bag with nitrogen then pushing on top to remove most of the gas within. Repeat purging three times.
    *Note: Ensure airtight chamber is not capped during purges.*
16. Seal the glove bag and turn off the nitrogen flow.
    **Steps 17–18 are performed inside the glove bag**
17. Inside the glove bag, add 822 µL gelation solution and 40 µL initiator solution in a 1.5-mL centrifuge tube. Flip the tube upside down five times for mixing.
18. Add 800 µL active gelation solution to each gel in the 6-well plate. Seal the airtight chamber.
19. Place the airtight chamber on a shaker at 50 rpm for 20 minutes
20. Turn on the nitrogen flow. Purge the glove bag by filling the glove bag with nitrogen then pushing on top to remove most of the gas within. Repeat purging three times.
21. Seal the glove bag and turn off the nitrogen flow.
    **Step 22-26 are performed inside the glove bag**
22. Transfer the gel to the gelation chamber with a paintbrush
23. Add 411 µL gelation solution and 20 µL initiator solution in a 1.5-mL centrifuge tube. Flip the tube upside down five times for mixing.
24. Add 100 µL active gelation solution to the gel. Cap with hydrophobic glass coverslip.
25. Add more active gelation if necessary to surround the gel.
26. Place gelation chambers in the sealed airtight humidified chamber.
27. Remove the airtight humidified chamber from the glove bag. Place it in the dark at room temperature overnight (12–16 hours).
28. After incubation, cut out sections of gel into 0.5 x 0.5 cm pieces.

D) Cleave the DATD re-embedded gel and expand

1. Cleave the DATD re-embedded gel with 30 mL 50mM sodium metaperiodate in 0.1M sodium acetate buffer (pH 5) for 1 hour on the shaker.
2. Place sections in a 20-cm petri dish containing 1x PBS. Measure expansion factor. Fully expand sections in dd$H_2O$. Measure expansion factor.

### Supplementary Tables

#### Supplementary Table 1

Table documenting different in situ proteomic technologies compared based on spatial resolution, sensitivity, hardware and scale (‘omics’). The table categorizes the technologies based on general strategy (left-most column), spatial resolution, sensitivity, hardware, and scale.

| Comparison of spatial proteomic technologies | | | | | | | | |
|---|---|---|---|---|---|---|---|---|
| | | | | | Points for comparison | | | |
| | Author | Title | Year | Link | Spatial resolution (best) | Sensitivity | Hardware | Scale ('omics') |
| **Proximity labeling techniques** | Branon *et al.* | Efficient proximity labeling in living cells and organisms with TurboID | 2018 | https://www.nature.com/articles/nbt.4201 | Proteins will be labeled around a few nanometers of the enzyme. Averaged proteomic landscape across all the locations where the enzyme was active in the sample. | Limited to mass spectrometry sensitivity, which is at best, attomole sensitivity (approximately ~10^6 molecules needed for detection). | LC-MS/MS. | In theory MS can detect any protein, but is limited by: (1) Protein not in the right dynamic range to be detected (eg. protein abundance is too low) (2) Ionization efficiency of the proteins (eg. in highly complex samples, peptides can compete for ionization), (3) sample preparation considerations, (eg. membrane proteins may not solubilize, and degrade). |
| **Expansion Microscopy with Mass Spectrometry** | Drelich *et al.* | Toward High Spatially Resolved Proteomics Using Expansion Microscopy | 2021 | https://pubs.acs.org/doi/10.1021/acs.analchem.0c05372 | ~ 330 um lateral resolution. | | Manual microssection; LC-MS/MS. | |
| | Li *et al.* | Spatially resolved proteomics via tissue expansion | 2022 | https://www.nature.com/articles/s41467-022-34824-2 | ~ 160 um lateral resolution. | | Manual microssection; LC-ESI/MS. | |
| | Chan *et al.* | Gel-assisted mass spectrometry imaging enables sub-micrometer spatial lipidomics | 2024 | https://www.nature.com/articles/s41467-024-49384-w | ~ 1.3 um lateral resolution. | | MALDI-MSI. | |
| | Dong *et al.* | Spatial proteomics of single cells and organelles on tissue slides using filter-aided expansion proteomics | 2024 | https://www.nature.com/articles/s41467-024-53683-7 | ~ 73 um lateral resolution. | | LC-ESI/MS. | |
| | Zhang *et al.* | TEMI: tissue-expansion mass-spectrometry imaging | 2025 | https://www.nature.com/articles/s41592-025-02664-9 | ~ 20 um lateral resolution. | | MALDI-MSI. | |
| | Wang *et al.* | iPEX enables micrometre-resolution deep spatial proteomics via tissue expansion | 2025 | https://www.nature.com/articles/s41586-025-09734-0 | ~ 1-5 um lateral resolution. | | MALDI-MSI. | |
| **Tissue clearing techniques / Expansion Microscopy with multiplexed antibody staining** | Chung *et al.* | Structural and molecular interrogation of intact biological systems | 2013 | https://www.nature.com/articles/nature12107 | No expansion of tissue. Diffraction limit of light resolution ~ 300-700 nm. | At this resolution, not single molecule. Antibody read-out is dependent on affinity, and specificity. Also depends on fluorophore brightness, signal amplification and detection strategies. | Confocal microscope, (Leica SP5), single-photon and two-photon imaging. | Three rounds of antibody staining (3 colors per round). Total of 9 protein targets. |
| | Murray *et al.* | Simple, Scalable Proteomic Imaging for High-Dimensional Profiling of Intact Systems | 2015 | https://www.cell.com/cell/comments/S0092-8674(15)01505-6 | No expansion of tissue. Diffraction limit of light resolution ~ 300-700 nm. | | Confocal microscope, and custom-built light-sheet microscope. | 22 rounds of antibody staining (3 colors per round). Total of 66 targets. total |
| | Ku et al. | Multiplexed and scalable super-resolution imaging of threedimensional protein localization in size-adjustable tissues | 2016 | https://www.nature.com/articles/nbt.3641 | Expansion factor of ~ 4 fold. States ~ 60 nm lateral resolution. | | Single-photon confocal laser scanning imaging (Olympus). | Seven rounds of antibody staining (3 colors per round). Total of 21 targets. |
| | Park *et al.* | Protection of tissue physicochemical properties using polyfunctional crosslinkers | 2018 | https://www.nature.com/articles/nbt.4281 | Expansion factor of ~ 3 fold. (resolution not stated, but perhaps approximately ~ 130 nm). | | Confocal microscope (Olympus Confocal FV1000MPE 1, Leica TCS SP8), light sheet microsope (SmartSPIM, LifeCanvas). | Five rounds of antibody staining (3 colors per round). Total of 15 targets. |
| | Park *et al.* | Epitope-preserving magnified analysis of proteome (eMAP) | 2021 | https://www.science.org/doi/10.1126/sciadv.abf6589 | Expansion factor of ~ 10 fold (resolution can be estimated as approximately ~ 40 nm). | | Confocal microscope (Leica TCS SP8). | Three rounds of antibody staining. |
| | Bai *et al.* | Expanded vacuum-stable gels for multiplexed high-resolution spatial histopathology | 2023 | https://www.nature.com/articles/s41467-023-39616-w | Expansion factor of ~ 3.7 fold (resolution can be estimated as approximately ~ 100 nm). | | Custom MIBI-TOF mass spectrometer equipped with an oxygen duoplasmatron ion gun (Alpha), a custom MIBI-TOF mass spectrometer (Betty) equipped with a xenon ion source (Hyperion, Oregon Physics), and a commercially available MIBIscope from Ionpath equipped with a xenon ion source (with MIBI software version v.1.7.0-0f60ffbc). | Detects > 40 targets total using isotope-conjugated antibodies. |
| | Park *et al.* | Integrated platform for multiscale molecular imaging and phenotyping of the human brain | 2024 | https://www.science.org/doi/10.1126/science.adh9979 | Expansion factor of ~ 4.5 fold (resolution can be estimated as ~ 90 nm). | | Confocal and MegaSPIM light-sheet microscope. | Seven rounds of antibody staining. |
| | Kang *et al.* | Multiplexed expansion revealing for imaging multiprotein nanostructures in healthy and diseased brain | 2024 | https://www.nature.com/articles/s41467-024-53729-w | Expansion factor of ~ 20 fold, and ~ 20 nm lateral resolution. | | Nikon CSU-W1 or SORA confocal microscope. | Seven rounds of antibody staining. |

### Supplementary Table 2

A table documenting a non-exhaustive list of Edman degradation variations and optimizations over the years (from 1950 to 1996), including a wide range of conjugation solvents and cleavage conditions. All of the strategies rely on phenylthiohydantoin amino acid (PTH-aa) detection to separate and extract the PTH-aa.

**Edman degradation protocols**

| Title | Author | Year | Conjugation condition | Cleavage condition | Link |
|---|---|---|---|---|---|
| Method for Determination of the Amino Acid Sequence in Peptides | Pehr Edman | 1950 | 50% pyridine in water at 40°C with NaOH to about pH 9. 4.8% PITC (v/v) was added to that solution for 30 minutes. The pH was maintained throughout the reaction by adding small portions of NaOH. | Anhydrous nitromethane saturted with hydrogen chloride, 15 minutes. | http://actachemscand.org/pdf/acta_vol_04_p0283-0293.pdf |
| A technique for stepwise degradation of proteins from the amino-end | H. Fraenkel-Conrat | 1954 | 20% PITC in peroxide-free dioxane incubation for 2-3h at 40 °C. | A mix of glacial acetic acid and 5.7N HCl incubation for 4-16h. | https://pubs.acs.org/doi/10.1021/ja01642a085 |
| Quantitative determination of N-terminal amino acids in some serum proteins. | Sten Eriksson and John Sjöquist | 1960 | 100:3:1 of pyridine:triethylamine:PITC incubation at 40°C for 1.5 hours. | Mix of 1 mL water and 2 mL of HCl saturated with acetic acid and kept for 2 hours at 40 °C. | https://www.sciencedirect.com/science/article/pii/0006300260914530?via%3Dihub |
| A protein sequenator | Pehr Edman and Geoffrey Begg | 1967 | 5% v/v PITC in heptane was added after prior incubation with basic solvent (Quadrol). Incubated at 55°C for 30 minutes. | Anydrous HFBA was repeatedly added at 50°C. | https://febs.onlinelibrary.wiley.com/doi/full/10.1111/j.1432-1033.1967.tb00047.x |
| Solid-Phase Edman Degradation. An Automatic Peptide Sequencer. | Richard A. Laursen | 1971 | 1.7 mL of buffer (3:2 mixture of pyridine and a N-methylmorpholiniumtrifluoroacetate buffer (pH 8.1)) and 0.7 mL 20% PITC in acetonitrile incubation for 20 minutes at 45°C. | TFA incubation for 30 minutes at 45 °C. | https://febs.onlinelibrary.wiley.com/doi/abs/10.1111/j.1432-1033.1971.tb01366.x |
| A manual method of sequential edman degradation followed by dansylation for the determination of protein sequences. | Maire E. Percy and Barbara M. Buchwald | 1972 | 500 µL of 50% pyridine and 63 µL of N-ethylmorpholine prior incubation before 25 µL PITC addition and bubbling with nitrogen for 1 minute.. Incubated for 1 hour at 37°C or 45°C. | TFA incubation for 20 minutes at 37°C. | https://www.sciencedirect.com/science/article/pii/0003269772900073 |
| Automatic solid-phase Edman degradation | Richard A. Laursen | 1972 | 0.7 mL 5% PITC in acetonitrile in 1.7 mL of 3:2 pyridine and N-methylmorpholinium trifluoroacetate buffer (pH 8.1) at 45°C for 25 min. | TFA incubation for 30 minutes at 45°C. | https://www.sciencedirect.com/science/article/pii/S0076687972250303?via%3Dihub |
| Rapid manual sequencing of multiple peptide samples in a nitrogen chamber. | Richard B. Meagher | 1975 | 7.5% v/v PITC in 50% pyridine with $10^{-5}$ M dithiothreitol (DTT) for 60-90 minutes at 45°C. | TFA incubation at 45°C for 15 minutes. | https://www.sciencedirect.com/science/article/pii/0003269775903127?via%3Dihub |
| A general procedure for the manual sequencing of small quantities of peptides. | George E. Tarr | 1975 | 30% pyridine, 25% aqueous trimethylamine (0.1% in ethanthiol), 10% PITC in pyridine added at 1:1:4 ratio v/v for 25-30 minutes at 50 °C. | TFA purging for 6 minutes at 50 °C. | https://www.sciencedirect.com/science/article/pii/0003269775903589?via%3Dihub |
| Improved manual sequential analysis of peptides. | Mark Boehnert and David H. Schlesinger | 1979 | Under nitrogen, 0.4M triethylamine in propanol:water (3:2) solvent previously adjusted to pH 9.5 with TFA added prior to PITC. Performed at 54 °C for 30 minutes. | Concentrated HCl at 22°C for 5 minutes. | https://www.sciencedirect.com/science/article/pii/0003269779906080 |
| A gas-liquid solid phase peptide and protein sequenator. | Hewick et al. | 1981 | 15% PITC in n-heptane at 42°C for ~ 8 minutes twice. | TFA at 42°C and 0.01% dithiothreitol (DTT) twice once for ~ 5 minutes, another ~ 7 minutes. | https://www.sciencedirect.com/science/article/pii/S0021925818433777 |
| Manual Edman Degradation Peptides. Methods in Molecular Biology. Humana Press | Per Klemm | 1984 | 2.5% PITC 50% pyridine for 20 minutes at 50°C. | TFA incubation for 10 minutes at 45 °C. | https://link.springer.com/protocol/10.1385/0-89603-062-8:243 |
| A manual sequencing method for identification of phosphorylated amino acids in phosphopeptides. | Sean Sullivan and Tai Wai Wong | 1991 | Methanol:water:triethylamine:PITC mix (7:1:1:1 v/v) incubation for 10 minutes at 50 °C. | TFA incubation at 50 °C for 6 minutes. | https://www.sciencedirect.com/science/article/pii/000326979190356X |
| Semi-automatic amino acid sequencing and D/L-configuration determination of peptides with detection of liberated *N*-terminal phenylthiocarbamoylamino acids | Lida et al. | 1998 | 5% v/v PITC in heptane, and 12.5% m/v trimethylamine. | 10-200 mol $dm^{-3}$ boron trifluoride etherate in acetonitrile at 48°C for 5 minutes. | https://pubs.rsc.org/en/content/articlelanding/1998/an/a806109b |
| Proton: a major factor for the racemization and the dehydration at the cyclization/cleavage stage in the Edman sequencing method | Matsunaga et al. | 1996 | 20 mM 7-[(N,N-dimethylamino)sulfonyl]-4-(2,1,3-benzoxadiazolyl) isothiocyanate (i.e., DBD-NCS) in 50% pyridine in water with 10 µL 10-100 µM dipeptide, mixed and heated at 50°C for 15 minutes. | 1% v/v boron trifluoride and 0.1% v/v ethanethiol in dichloroethane at 50°C for 5 minutes. | https://pubs.acs.org/doi/10.1021/ac951253r |

#### Supplementary Table 3

Table documenting the name of compounds, exact mass, and chemical formula. These values are used for searching in the Agilent MassHunter Software for automated AUC extraction of species from the LC/QToF data. Greyed out characters are not counted in formula and exact mass, and are fragments eliminated after trypsinization from the gel. Curly bracket after amino acid is to specify modifications that have occurred to that specific side chain (e.g., modified with acryloyl functional group "acr", specific post-translational modification from oxidation, such as P{hydroxyproline} etc.). Post-translational oxidation modifications for each amino acid are described in more detail in **Supplementary Figure 8** (abbreviations: MetO is methionine sulfoxide, $MetO_2$ is methionine sulfone, DOPA is 3,4-dihydroxyphenlylanine, Oia is oxindolylalanine, NFK is N-formylkynurenine, KN is kynurenin).

| Peptide species | Monoisotopic mass | Formula |
|---|---|---|
| *Peptide for LC/QToF trypsinization assay ("A15-peptide") with PITC* | | |
| AGGAGGLLGGSRGGK{acr} | 971.5148 | C39H69N15O14 |
| *PTC*-AGGAGGLLGGSRGGK{acr} | 1106.5291 | C46H74N16O14S |
| GGAGGLLGGSRGGK{acr} | 900.4777 | C36H64N14O13 |
| *PTC*-GGAGGLLGGSRGGK{acr} | 1035.4920 | C43H69N15O13S |
| GAGGLLGGSRGGK{acr} | 843.4563 | C34H61N13O12 |
| *PTC*-AGGLLGGSRGGK{acr} | 978.4705 | C41H66N14O12S |
| GGLLGGSRGGK{acr} | 786.4348 | C32H58N12O11 |
| *Peptide for LC/QToF trypsinization assay ("A15-peptide") with ClickP* | | |
| *ClickP*-AGGAGGLLGGSRGGK{acr} | 1175.5618 | C48H77N19O14S |
| *Peptide for SPAAC fluorescence assay ("K{$N_3$}15-peptide" and "AK{$N_3$}15-peptide")* | | |
| K{N3}GGAGGLLGGSRGGK{acr} | 1350.7116 | C52H92N22O17 |
| AK{N3}GAGGLLGGSRGGK{acr} | 1364.7273 | C56H96N22O18 |
| *Peptide for LC/QToF trypsinization assay ("A15-peptide") with FITC* | | |
| *FTC*-AGGAGGLLGGSRGGK{acr} | 1360.5506 | C60H80N16O19S |
| *Peptide for ionization efficiency standard ("A9-peptide")* | | |
| AGGAGK{acr}GLR | 839.4613 | C35H61N13O11 |
| *C-terminal, after arginine, of the N-terminal peptides* | | |
| XaaGGAGRGLGK{acr} | 427.2431 | C19H33N5O6 |
| *Methionine peptide ("M-peptide") and its oxidation products* | | |
| MGGAGRGLGK{acr} | 547.2537 | C20H37N9O7S |
| M{MetO}GGAGRGLGK{acr} | 563.2486 | C20H37N9O8S |
| M{$MetO_2$}GGAGRGLGK{acr} | 579.2435 | C20H37N9O9S |
| MGGAGRGLGK{acr} | 956.4862 | C39H68N14O12S |
| *Cysteine peptide ("C-peptide") and its oxidation products* | | |
| CGGAGRGLGK{acr} | 519.2224 | C18H33N9O7S |
| K{acr}GLGRGAGGC{Cystine} GGAGRGLGK{acr} | 1036.4291 | C36H64N18O14S2 |
| C{Sulfenic acid}GGAGRGLGK{acr} | 535.2173 | C18H33N9O8S |
| C{Sulfinate}GGAGRGLGK{acr} | 551.2122 | C18H33N9O9S |
| C{Sulfonate}GGAGRGLGK{acr} | 567.2071 | C18H33N9O10S |
| CGGAGRGLGK{acr} | 928.4549 | C37H64N14O12S |
| *Tyrosine peptide ("Y-peptide") and its oxidation products* | | |
| YGGAGRGLGK{acr} | 579.2765 | C24H37N9O8 |
| Y{DOPA}GGAGRGLGK{acr} | 595.2714 | C24H37N9O9 |
| K{acr}GLGRGAGGY{dityrosine} | 1156.5374 | C48H72N18O16 |

| | | |
|---|---|---|
| YGGAGRGLGK{acr} | | |
| YGGAGRGLGK{acr} | 988.509 | C43H68N14O13 |
| *Phenylalanine peptide ("$F_1$" peptide) and its oxidation products* | | |
| FGGAGRGLGK{acr} | 563.2816 | C24H37N9O7 |
| F{Meta-tyrosine}GGAGRGLGK{acr} | 579.2765 | C24H37N9O8 |
| F{DOPA}GGAGRGLGK{acr} | 595.2714 | C24H37N9O9 |
| FGGAGRGLGK{acr} | 972.5141 | C43H68N14O12 |
| *Tryptophan peptide ("W-peptide") and its oxidation products* | | |
| WGGAGRGLGK{acr} | 602.2925 | C26H38N10O7 |
| W{Oia}GGAGRGLGK{acr} | 618.2874 | C26H38N10O8 |
| W{NFK}GGAGRGLGK{acr} | 634.2823 | C26H38N10O9 |
| W{KN}GGAGRGLGK{acr} | 606.2874 | C25H38N10O8 |
| WGGAGRGLGK{acr} | 1011.525 | C45H69N15O12 |
| *Histidine peptide ("H-peptide") and its oxidation products* | | |
| HGGAGRGLGK{acr} | 553.2721 | C21H35N11O7 |
| H{2-oxo-histidine}GGAGRGLGK{acr} | 567.2514 | C21H33N11O8 |
| H{aspartate}GGAGRGLGK{acr} | 531.2401 | C19H33N9O9 |
| H{formylasparagine}GGAGRGLGK{acr} | 558.251 | C20H34N10O9 |
| H{aspartylurea}GGAGRGLGK{acr} | 543.2401 | C20H33N9O9 |
| HGGAGRGLGK{acr} | 962.5046 | C40H66N16O12 |
| *Proline peptide ("P-peptide") and its oxidation products* | | |
| PGGAGRGLGK{acr} | 513.2659 | C20H35N9O7 |
| P{pyrroline-5-carboxylate} GGAGRGLGK{acr} | 511.2503 | C20H33N9O7 |
| P{hydroxyproline}GGAGRGLGK{acr} | 529.2609 | C20H35N9O8 |
| P{glutamic semialdehyde} GGAGRGLGK{acr} | 527.2816 | C21H37N9O7 |
| PGGAGRGLGK{acr} | 922.4985 | C39H66N14O12 |
| *Arginine peptide ("R-peptide") and its oxidation products* | | |
| RGGAGRGLGK{acr} | 572.3143 | C21H40N12O7 |
| R{hydroxyarginine} GGAGRGLGK{acr} | 588.3092 | C21H40N12O8 |
| R{oxoarginine}GGAGRGLGK{acr} | 573.2983 | C21H39N11O8 |
| R{glutamic semialdehyde} GGAGRGLGK{acr} | 527.2816 | C21H37N9O7 |
| RGGAGRGLGK{acr} | 981.5468 | C40H71N17O12 |
| *PTH-amino acids* | | |
| PTH-phenylalanine | 282.0827 | C16H14N2OS |
| PTH-alanine | 206.0514 | C10H10N2OS |
| PTH-tyrosine | 298.0776 | C16H14N2O2S |
| PTH-glycine | 192.0357 | C9H8N2OS |

#### Supplementary Table 4

Tables documenting the results for all ExM and ExMre gels placed in various solvents plotted in Fig. 2**Ci-vii**. (i) Surface size (flat/top side of gel) of ExM and ExM re-embedded (ExMre) gels when placed in solvents used in Edman degradation, namely 1:1 pyridine to water, acetonitrile (ACN), dimethylsulfoxide (DMSO), formamide, 0.1 M sodium bicarbonate ($NaHCO_3$) pH 8.5, Trifluoroacetic acid (TFA), with and without Edman reagents (PITC, 1:1000; FITC, 5.9 mM in 23:77 DMSO:0.1 M sodium bicarbonate pH 8.5). (ii) Normalized values from (i) to flat/top surface size in 1X PBS. (iii) Mean percent shrinkage going from 1X PBS to various solvents based on values obtained in (ii).

Note: Some columns contain what appear to be repeated values - this is not an error. Instead, this often occurred because of the way we made and trimmed the gels, to a similar size of approximately ~0.5 x 0.5 cm and then measured exactly when washed 3x with 1X PBS 10 min at room temperature. Due to the limited values of length and width that could be measured with our ruler (gel length and width sizes between ~0.1 - 0.8 cm), we were only able to measure a small range of length and width values. This also limited the output values of the area when these two values were multiplied. As a result, since our gels had similar starting sizes prior to incubation in solvent or solvent or buffer with PITC or FITC, many gels ended up with similar or identical measured areas after treatment.

(i) Surface size of the flat/top side of ExM and ExMre gels, made from 3 separate gelation solutions, in various solvents in with/without PITC or FITC, except for pyridine which is 3 replicates from the same gelation solution.

| | Area of the gel ($cm^2$) | | | | | |
|---|---|---|---|---|---|---|
| **Gelation number** | **ExM 1X PBS** | **ExMre 1X PBS** | **ExM pyridine** | **ExMre pyridine** | **ExM pyridine with PITC** | **ExMre pyridine with PITC** |
| 1 (replicate 1) | 0.300 | 0.250 | 0.030 | 0.040 | 0.020 | 0.040 |
| 1 (replicate 2) | 0.200 | 0.300 | 0.020 | 0.090 | 0.020 | 0.090 |
| 1 (replicate 3) | 0.275 | 0.250 | 0.020 | 0.090 | 0.020 | 0.063 |
| **Gelation number** | **ExM 1X PBS** | **ExMre 1X PBS** | **ExM 1:1 pyridine to water** | **ExMre 1:1 pyridine to water** | **ExM 1:1 pyridine to water with PITC** | **ExMre 1:1 pyridine to water with PITC** |
| 1 | 0.275 | 0.250 | 0.040 | 0.075 | 0.040 | 0.040 |
| 2 | 0.330 | 0.250 | 0.050 | 0.075 | 0.040 | 0.040 |
| 3 | 0.300 | 0.250 | 0.040 | 0.063 | 0.040 | 0.040 |
| **Gelation number** | **ExM 1X PBS** | **ExMre 1X PBS** | **ExM ACN** | **ExMre ACN** | **ExM ACN with PITC** | **ExMre ACN with PITC** |
| 1 | 0.330 | 0.250 | 0.075 | 0.063 | 0.040 | 0.040 |
| 2 | 0.275 | 0.250 | 0.060 | 0.050 | 0.040 | 0.040 |
| 3 | 0.250 | 0.250 | 0.040 | 0.050 | 0.040 | 0.040 |
| **Gelation number** | **ExM 1X PBS** | **ExMre 1X PBS** | **ExM DMSO** | **ExMre DMSO** | **ExM DMSO with PITC** | **ExMre DMSO with PITC** |
| 1 | 0.330 | 0.250 | 0.060 | 0.123 | 0.090 | 0.123 |
| 2 | 0.275 | 0.250 | 0.075 | 0.123 | 0.075 | 0.123 |
| 3 | 0.275 | 0.250 | 0.075 | 0.123 | 0.075 | 0.123 |
| **Gelation number** | **ExM 1X PBS** | **ExMre 1X PBS** | **ExM formamide** | **ExMre formamide** | **ExM formamide with PITC** | **ExMre formamide with PITC** |
| 1 | 0.300 | 0.225 | 0.560 | 0.275 | 0.720 | 0.330 |
| 2 | 0.275 | 0.225 | 0.560 | 0.300 | 0.720 | 0.325 |
| 3 | 0.250 | 0.250 | 0.560 | 0.303 | 0.720 | 0.360 |
| **Gelation** | **ExMre 1X PBS** | **ExMre 0.1 M** | **ExMre 5.9 mM** | | | |

| number | | NaHCO3 pH 8.5 | FITC in 23:77 DMSO:0.1 M NaHCO3 pH 8.5 |
|---|---|---|---|
| 1 | 0.225 | 0.275 | 0.225 |
| 2 | 0.225 | 0.275 | 0.225 |
| 3 | 0.225 | 0.275 | 0.225 |
| **Gelation number** | **ExMre 1X PBS** | **ExMre TFA** | |
| 1 | 0.250 | 0.250 | |
| 2 | 0.250 | 0.250 | |
| 3 | 0.250 | 0.250 | |

(ii) Normalized values to top/flat surface size in 1X PBS.

| | ExM top/flat surface size normalized to 1X PBS | | ExMre top/flat surface size normalized to 1X PBS | |
|---|---|---|---|---|
| **Gelation number** | **pyridine** | **pyridine with PITC** | **pyridine** | **pyridine with PITC** |
| 1(replicate 1) | 0.100 | 0.067 | 0.160 | 0.160 |
| 1(replicate 2) | 0.100 | 0.100 | 0.300 | 0.300 |
| 1(replicate 3) | 0.072 | 0.072 | 0.360 | 0.250 |
| *Stdev* | *0.016* | *0.018* | *0.103* | *0.071* |
| | ExM top/flat surface size normalized to 1X PBS | | ExMre top/flat surface size normalized to 1X PBS | |
| **Gelation number** | **1:1 pyridine to water** | **1:1 pyridine to water with PITC** | **1:1 pyridine to water** | **1:1 pyridine to water with PITC** |
| 1 | 0.145 | 0.145 | 0.300 | 0.160 |
| 2 | 0.152 | 0.121 | 0.300 | 0.160 |
| 3 | 0.133 | 0.133 | 0.250 | 0.160 |
| *Stdev* | *0.009* | *0.012* | *0.029* | *0.000* |
| | ExM top/flat surface size normalized to 1X PBS | | ExMre top/flat surface size normalized to 1X PBS | |
| **Gelation number** | **ACN** | **ACN with PITC** | **ACN** | **ACN with PITC** |
| 1 | 0.227 | 0.121 | 0.250 | 0.160 |
| 2 | 0.218 | 0.145 | 0.200 | 0.160 |
| 3 | 0.160 | 0.160 | 0.200 | 0.160 |
| *Stdev* | *0.036* | *0.020* | *0.029* | *0.000* |
| | ExM top/flat surface size normalized to 1X PBS | | ExMre top/flat surface size normalized to 1X PBS | |
| **Gelation number** | **DMSO** | **DMSO with PITC** | **DMSO** | **DMSO with PITC** |
| 1 | 0.182 | 0.273 | 0.490 | 0.490 |
| 2 | 0.273 | 0.273 | 0.490 | 0.490 |
| 3 | 0.273 | 0.273 | 0.490 | 0.490 |
| *Stdev* | *0.052* | *0.000* | *0.000* | *0.000* |
| | ExM top/flat surface size normalized to 1X PBS | | ExMre top/flat surface size normalized to 1X PBS | |
| **Gelation number** | **Formamide** | **Formamide with PITC** | **Formamide** | **Formamide with PITC** |
| 1 | 1.867 | 2.400 | 1.222 | 1.467 |
| 2 | 2.036 | 2.618 | 1.333 | 1.444 |
| 3 | 2.240 | 2.880 | 1.210 | 1.440 |
| *Stdev* | *0.187* | *0.240* | *0.068* | *0.014* |
| | ExM top/flat surface size normalized to 1X PBS | | | |
| **Gelation number** | **0.1 M NaHCO3 pH 8.5** | **5.9 mM FITC in DMSO:0.1 M NaHCO3 pH 8.5** | | |
| 1 | 1.222 | 1.000 | | |

| | | |
|---|---|---|
| 2 | 1.222 | 1.000 |
| 3 | 1.222 | 1.000 |
| *Stdev* | *0.000* | *0.000* |

| | ExMre top/flat surface size normalized to 1X PBS |
|---|---|
| **Gelation number** | **TFA** |
| 1 | 1.000 |
| 2 | 1.000 |
| 3 | 1.000 |
| *Stdev* | *0.000* |

(iii) Percent shrinkage of ExM and ExMre gels when placed in different solvents with/without PITC or FITC.

| **pyridine** | **% shrinkage from a to b (a vs. b)** |
|---|---|
| ExM in 1X PBS vs. pyridine without PITC | 90.9 |
| ExM in 1X PBS vs. pyridine ExM with PITC (1:9 ratio PITC:pyridine) | 92.0 |
| ExM in pyridine without PITC vs. ExM in pyridine with PITC (1:9 ratio PITC:pyridine) | 1.1 |
| ExMre in 1X PBS vs. pyridine without PITC | 72.7 |
| ExMre in 1X PBS vs. pyridine with PITC (1:9 ratio PITC:pyridine) | 76.3 |
| ExMre in pyridine without PITC vs. ExMre in pyridine with PITC (1:9 ratio PITC:pyridine) | 3.7 |
| **1:1 pyridine to water** | **% shrinkage from a to b (a vs. b)** |
| ExM in 1X PBS vs. 1:1 pyridine to water without PITC | 85.7 |
| ExM in 1X PBS vs. 1:1 pyridine to water ExM with PITC (1:1000 ratio PITC to 1:1 pyridine to water) | 86.7 |
| ExM in 1:1 pyridine to water without PITC vs. ExM in 1:1 pyridine to water with PITC (1:1000 ratio PITC to 1:1 pyridine to water) | 1.0 |
| ExMre in 1X PBS vs. 1:1 pyridine to water without PITC | 71.7 |
| ExMre in 1X PBS vs. 1:1 pyridine to water with PITC (1:1000 ratio PITC to 1:1 pyridine to water) | 84.0 |
| ExMre in 1:1 pyridine to water without PITC vs. ExMre in 1:1 pyridine to water with PITC (1:1000 ratio PITC to 1:1 pyridine to water) | 12.3 |
| **Acetonitrile** | **% shrinkage from a to b (a vs. b)** |
| ExM in 1X PBS vs. acetonitrile without PITC | 79.8 |
| ExM in 1X PBS vs. acetonitrile ExM with PITC (1:1000 ratio PITC:acetonitrile) | 85.8 |
| ExM in acetonitrile without PITC vs. ExM in acetonitrile with PITC (1:1000 ratio PITC:acetonitrile) | 6.0 |
| ExMre in 1X PBS vs. acetonitrile without PITC | 78.3 |
| ExMre in 1X PBS vs. acetonitrile with PITC (1:1000 ratio PITC:acetonitrile) | 84.0 |
| ExMre in acetonitrile without PITC vs. ExMre in acetonitrile with PITC (1:1000 ratio PITC:acetonitrile) | 5.7 |
| **DMSO** | **% shrinkage from a to b (a vs. b)** |
| ExM in 1X PBS vs. DMSO without PITC | 75.8 |
| ExM in 1X PBS vs. DMSO ExM with PITC (1:1000 ratio PITC:DMSO) | 72.7 |
| ExM in DMSO without PITC vs. ExM in DMSO with PITC | -3.0 |

| | |
|---|---|
| (1:1000 ratio PITC:DMSO) | |
| ExMre in 1X PBS vs. DMSO without PITC | 51.0 |
| ExMre in 1X PBS vs. DMSO with PITC (1:1000 ratio PITC:DMSO) | 51.0 |
| ExMre in DMSO without PITC vs. ExMre in DMSO with PITC (1:1000 ratio PITC:DMSO) | 0.0 |
| **Formamide** | **% shrinkage from a to b (a vs. b)** |
| ExM in 1X PBS vs. formamide without PITC | -104.8 |
| ExM in 1X PBS vs. formamide ExM with PITC (1:1000 ratio PITC:formamide) | -163.3 |
| ExM in formamide without PITC vs. ExM in formamide with PITC (1:1000 ratio PITC:formamide) | -58.5 |
| ExMre in 1X PBS vs. formamide without PITC | -25.5 |
| ExMre in 1X PBS vs. formamide with PITC (1:1000 ratio PITC:formamide) | -45.0 |
| ExMre in formamide without PITC vs. ExMre in formamide with PITC (1:1000 ratio PITC:formamide) | -19.5 |
| **0.1 M NaHCO3 pH 8.5** | **% shrinkage from a to b (a vs. b)** |
| ExMre in 1X PBS vs. 0.1 M NaHCO3 pH 8.5 without FITC | -22.2 |
| ExMre in 1X PBS vs. 0.1 M NaHCO3 pH 8.5 with FITC in DMSO (23:77 ratio DMSO:0.1 M sodium bicarbonate buffer pH 8.5) | 0.0 |
| ExMre in 0.1 M NaHCO3 pH 8.5 without FITC vs. ExMre in 0.1 M NaHCO3 pH 8.5 with FITC in DMSO (23:77 ratio DMSO:0.1 M sodium bicarbonate buffer pH 8.5) | 22.2 |
| **TFA** | **% shrinkage from a to b (a vs. b)** |
| ExMre in 1X PBS vs. in TFA | 0.0 |

#### Supplementary Table 5

Values and analysis for the LC/QToF trypsinization assay from Fig. 3 and strain-promoted alkyne-azide cycloaddition (SPAAC) chemistry click chemistry fluorescence assay.

##### (A) Formamide with PITC

(i) Raw data values for the area under the curve (AUC) of the chromatogram of various species for PITC (1:1000 ratio PITC:formamide) conjugation (Fig. 3**B**) (extracted based on the exact mass, see Methods for more details, and Source Data for raw traces). Three tables for the three separate gelation solutions. The cells highlighted in grey are the ones used to calculate the conversion rate (1-(solvent with PITC[“A15-peptide”] / solvent [“A15-peptide”])). A15-peptide stands for AGGAGGLLGGSRGGK{acr}. Cells with ‘-’ means that the species was not detected in that sample using the automatic extraction.

| Conjugation with PITC to formamide (1:1000 ratio PITC:formamide) | **Gelation** | **Formamide** | **Formamide with PITC** | **Formamide, then TFA** | **Formamide with PITC, then TFA** | **Conversion rate** |
|---|---|---|---|---|---|---|
| **A15-peptide** | Gelation 1 | 12,117,368 | 394,015 | 15,178,608 | 680,108 | 96.75 |
| **PITC conjugated A15-peptide** | | - | 7,493,956 | - | - | |
| **A15-peptide with cleaved N-terminal amino acid** | | 54,993 | 6,919,759 | 87,591 | 14,344,037 | |
| **A9-peptide (control)** | | 1,638,492 | 1,583,580 | 1,423,334 | 1,583,069 | |

| Conjugation with PITC to formamide (1:1000 ratio PITC:formamide) | **Gelation** | **Formamide** | **Formamide with PITC** | **Formamide, then TFA** | **Formamide with PITC, then TFA** | **Conversion rate** |
|---|---|---|---|---|---|---|
| **A15-peptide** | Gelation 2 | 14,191,286 | 567,866 | 16,829,402 | 1,384,801 | 96.00 |
| **PITC conjugated A15-peptide** | | - | 9,085,305 | - | - | |
| **A15-peptide with cleaved N-terminal amino acid** | | 69,133 | 11,251,107 | 100,469 | 18,304,827 | |
| **A9-peptide (control)** | | 1,610,108 | 1,518,859 | 1,642,476 | 1,658,903 | |

| Conjugation with PITC to formamide (1:1000 ratio PITC:formamide) | **Gelation** | **Formamide** | **Formamide with PITC** | **Formamide, then TFA** | **Formamide with PITC, then TFA** | **Conversion rate** |
|---|---|---|---|---|---|---|
| **A15-peptide** | Gelation 3 | 11,261,653 | 472,423 | 12,264,577 | 555,385 | 95.81 |
| **PITC conjugated A15-peptide** | | - | 6999122 | - | - | |
| **A15-peptide with cleaved N-terminal amino acid** | | 50,898 | 8,048,591 | 77,366 | 11,057,572 | |
| **A9-peptide (control)** | | 1,950,392 | 1,713,521 | 1,750,512 | 1,927,406 | |

(ii) Conversion rate of A15-peptide to PITC conjugated A15-peptide, and % yield for conjugation with PITC to formamide (1:1000 ratio PITC:formamide) (Fig. 3**B**).

| Conjugation with PITC to formamide (1:1000 ratio PITC:formamide) **Average % yield (peptide with cleaved N-terminal amino acid / non-modified peptide * 100)** | 98.72 |
|---|---|
| **Average conversion rate (%)** | 96.18 |

(iii) Statistics: One-way Analysis of Variance (ANOVA) and Tukey's post-hoc Honestly Significant Difference (HSD) test on the abundance of peptide with cleaved N-terminal amino acid using PITC to formamide (1:1000 ratio PITC:formamide) for conjugation (Fig. 3**Biii**). In Tukey HSD results, conditions are abbreviated, A: formamide, B: formamide with PITC, C: formamide then TFA, D: formamide with PITC then TFA.

| **ANOVA Results** | | | | | |
|---|---|---|---|---|---|
| Test | Comparison | sum_sq | df | F | PR(>F) |
| ANOVA | group | 4.53E+14 | 3.00E+00 | 3.32E+01 | 7.31E-05 |
| ANOVA | Residual | 3.64E+13 | 8.00E+00 | | |

| **Tukey HSD Results** | | | | | | | |
|---|---|---|---|---|---|---|---|
| Test | group1 | group2 | meandiff | p-adj | lower | upper | reject |
| Tukey HSD | A | B | 8.68E+06 | 4.73E-03 | 3.10E+06 | 1.43E+07 | TRUE |
| Tukey HSD | A | C | 3.01E+04 | 1.00E+00 | -5.55E+06 | 5.61E+06 | FALSE |
| Tukey HSD | A | D | 1.45E+07 | 1.51E-04 | 8.93E+06 | 2.01E+07 | TRUE |
| Tukey HSD | B | C | -8.65E+06 | 4.83E-03 | -1.42E+07 | -3.07E+06 | TRUE |
| Tukey HSD | B | D | 5.83E+06 | 4.09E-02 | 2.49E+05 | 1.14E+07 | TRUE |
| Tukey HSD | C | D | 1.45E+07 | 1.53E-04 | 8.90E+06 | 2.01E+07 | TRUE |

(iv) Gel size changes throughout in-gel Edman degradation with PITC to formamide (1:1000 ratio PITC:formamide) (Fig. 3**Ei**).

| **Conjugation with PITC to formamide (1:1000 PITC:formamide) - Gel size changes in Edman steps** | **Gelation 1, area ($cm^2$)** | **Gelation 2, area ($cm^2$)** | **Gelation 3, area ($cm^2$)** | **Mean (normalized to surface area in 1M Tris pH 9.5) ($cm^2$)** | **Standard deviation ($cm^2$)** |
|---|---|---|---|---|---|
| 1. 3x wash with Tris 1M pH 9.5 | 0.330 | 0.303 | 0.250 | 1.000 | 0.000 |
| 2. 2x formamide wash | 0.300 | 0.360 | 0.250 | 1.031 | 0.143 |
| 3. PITC conjugation | 0.360 | 0.420 | 0.300 | 1.224 | 0.151 |
| 4. TFA | 0.330 | 0.360 | 0.330 | 1.156 | 0.161 |
| 5. 2x formamide wash | 0.360 | 0.360 | 0.300 | 1.156 | 0.060 |
| 6. 3x wash with Tris 1M pH 8 | 0.330 | 0.303 | 0.250 | 1.000 | 0.000 |

##### (B) DMSO with PITC

(i) Raw data values for the area under the curve (AUC) of the chromatogram of various species for PITC to DMSO (1:1000 ratio PITC:DMSO) for conjugation (Fig. 3C) (extracted based on the exact mass, see Methods for more details, and Source Data for raw traces). Three tables for the three separate gelation solutions. The cells highlighted in grey are the ones used to calculate the conversion rate (1-(solvent with PITC["A15-peptide"] / solvent ["A15-peptide"])). A15-peptide stands for AGGAGGLLGGSRGGK{acr}. Cells with '-' means that the species was not detected in that sample using the automatic extraction.

| Conjugation with PITC to DMSO (1:1000 ratio PITC:DMSO) | **Gelation** | **DMSO** | **DMSO with PITC** | **DMSO, then TFA** | **DMSO with PITC, then TFA** | **Conversion rate** |
|---|---|---|---|---|---|---|

| | | | | | | |
|---|---|---|---|---|---|---|
| **A15-peptide** | Gelation 1 | 15,459,666 | 276,139 | 15,338,350 | 323,062 | 98.21 |
| **PITC conjugated A15-peptide** | | | 10,904,256 | | 23,333 | |
| **A15-peptide with cleaved N-terminal amino acid** | | 84,704 | 7,448,340 | 103,752 | 14,365,449 | |
| **A9-peptide (control)** | | 1,835,948 | 1,773,090 | 1,663,375 | 2,047,203 | |

| Conjugation with PITC to DMSO (1:1000 ratio PITC:DMSO) | **Gelation** | **DMSO** | **DMSO with PITC** | **DMSO, then TFA** | **DMSO with PITC, then TFA** | **Conversion rate** |
|---|---|---|---|---|---|---|
| **A15-peptide** | Gelation 2 | 18,095,571 | 230,579 | 17,231,541 | 344,879 | 98.73 |
| **PITC conjugated A15-peptide** | | | 8,054,863 | | 29,359 | |
| **A15-peptide with cleaved N-terminal amino acid** | | 106,966 | 7,135,934 | 89,852 | 15,418,384 | |
| **A9-peptide (control)** | | 2,155,301 | 1,298,155 | 1,840,897 | 2,023,220 | |

| Conjugation with PITC to DMSO (1:1000 ratio PITC:DMSO) | **Gelation** | **DMSO** | **DMSO with PITC** | **DMSO, then TFA** | **DMSO with PITC, then TFA** | **Conversion rate** |
|---|---|---|---|---|---|---|
| **A15-peptide** | Gelation 3 | 14,680,209 | 118,927 | 14,148,945 | 329,337 | 99.19 |
| **PITC conjugated A15-peptide** | | | 8,729,040 | | 25,109 | |
| **A15-peptide with cleaved N-terminal amino acid** | | 87,737 | 4,869,072 | 77,527 | 14,377,025 | |
| **A9-peptide (control)** | | 1,763,566 | 1,950,219 | 1,915,314 | 2,201,666 | |

(ii) Conversion rate of A15-peptide to PITC conjugated A15-peptide, and % yield for PITC (1:1000 ratio PITC:DMSO) conjugation (Fig. 3C).

| | |
|---|---|
| Conjugation with PITC to DMSO (1:1000 ratio PITC:DMSO) **Average % yield (peptide with cleaved N-terminal amino acid / non-modified peptide * 100)** | 94.52 |
| **Average conversion rate (%)** | 98.71 |

(iii) Statistics: One-way Analysis of Variance (ANOVA) and Tukey's post-hoc Honestly Significant Difference (HSD) test on the abundance of peptide with cleaved N-terminal amino acid using PITC to DMSO (1:1000 ratio PITC:DMSO) for conjugation (Fig. 3Ciii). In Tukey HSD results, conditions are abbreviated, A: DMSO, B: DMSO with PITC, C: DMSO then TFA, D: DMSO with PITC then TFA.

**ANOVA Results**

| Test | Comparison | sum_sq | df | F | PR(>F) |
|---|---|---|---|---|---|
| ANOVA | group | 4.33E+14 | 3.00E+00 | 2.46E+02 | 3.24E-08 |

| ANOVA | Residual | 4.70E+12 | 8.00E+00 |
|---|---|---|---|

| **Tukey HSD Results** | | | | | | | |
|---|---|---|---|---|---|---|---|
| Test | group1 | group2 | meandiff | p-adj | lower | upper | reject |
| Tukey HSD | A | B | 6.39E+06 | 3.37E-05 | 4.39E+06 | 8.39E+06 | TRUE |
| Tukey HSD | A | C | -2.76E+03 | 1.00E+00 | -2.01E+06 | 2.00E+06 | FALSE |
| Tukey HSD | A | D | 1.46E+07 | 5.64E-08 | 1.26E+07 | 1.66E+07 | TRUE |
| Tukey HSD | B | C | -6.39E+06 | 3.36E-05 | -8.40E+06 | -4.39E+06 | TRUE |
| Tukey HSD | B | D | 8.24E+06 | 4.96E-06 | 6.23E+06 | 1.02E+07 | TRUE |
| Tukey HSD | C | D | 1.46E+07 | 5.63E-08 | 1.26E+07 | 1.66E+07 | TRUE |

(iv) Gel size changes throughout in-gel Edman degradation with PITC (1:1000 ratio PITC:DMSO) conjugation (Fig. 3**Eii**).

| **Conjugation with PITC to DMSO (1:1000 ratio PITC:DMSO) - Gel size changes in Edman steps** | **Gelation 1, area ($cm^2$)** | **Gelation 2, area ($cm^2$)** | **Gelation 3, area ($cm^2$)** | **Mean (normalized to surface are in 1M Tris pH 9.5) ($cm^2$)** | **Standard deviation ($cm^2$)** |
|---|---|---|---|---|---|
| 1. 3x wash with Tris 1M pH 9.5 | 0.300 | 0.300 | 0.275 | 1.000 | 0.000 |
| 2. 2x DMSO wash | 0.300 | 0.300 | 0.275 | 1.000 | 0.000 |
| 3. PITC conjugation | 0.300 | 0.275 | 0.275 | 0.971 | 0.048 |
| 4. TFA | 0.225 | 0.275 | 0.225 | 0.829 | 0.084 |
| 5. 2x DMSO wash | 0.180 | 0.225 | 0.180 | 0.669 | 0.076 |
| 6. 3x wash with Tris 1M pH 8 | 0.300 | 0.300 | 0.275 | 1.000 | 0.000 |

##### (C) $NaHCO_3$ with FITC

(i) Raw data values for the area under the curve (AUC) of the chromatogram of various species for 5.9 mM FITC conjugation in 23:77 DMSO:0.1 M sodium bicarbonate ($NaHCO_3$) pH 8.5 (Fig. 3**D**) (extracted based on the exact mass, see **Methods** for more details, and **Source Data** for raw traces). Three tables for the three separate gelation solutions. The cells highlighted in grey are the ones used to calculate the conversion rate (1-(solvent with FITC[“A15-peptide”] / solvent [“A15-peptide”])). A15-peptide stands for AGGAGGLLGGSRGGK{acr}. Cells with ‘-’ means that the species was not detected in that sample using the automatic extraction.

| Conjugation with FITC in 0.1 M $NaHCO_3$ pH 8.5 (5.9 mM FITC in 23:77 ratio DMSO:0.1 M $NaHCO_3$ pH 8.5) | **Gelation** | **0.1 M $NaHCO_3$** | **23:77 DMSO:0.1 M $NaHCO_3$ with FITC** | **0.1 M $NaHCO_3$, then TFA** | **23:77 DMSO:0.1 M $NaHCO_3$ with FITC, then TFA** | **Conversion rate** |
|---|---|---|---|---|---|---|
| **A15-peptide** | Gelation 1 | 11,073,221 | 2,942,798 | 11,874,162 | 2,485,449 | 73.42 |
| **FITC conjugated A15-peptide** | | - | 495304 | - | - | |
| **A15-peptide with cleaved N-terminal amino acid** | | 130,625 | 5,100,100 | 131,053 | 8,326,486 | |
| **A9-peptide (control)** | | 1,192,016 | 1,113,044 | 1,246,017 | 1,198,247 | |

| Conjugation with FITC in 0.1 M $NaHCO_3$ pH 8.5 (5.9 mM FITC in 23:77 ratio DMSO:0.1 M $NaHCO_3$ pH 8.5) | **Gelation** | **0.1 M $NaHCO_3$** | **23:77 DMSO:0.1 M $NaHCO_3$ with FITC** | **0.1 M $NaHCO_3$, then TFA** | **23:77 DMSO:0.1 M $NaHCO_3$ with FITC, then TFA** | **Conversion rate** |
|---|---|---|---|---|---|---|
| **A15-peptide** | Gelation 2 | 15,174,962 | 3,649,573 | 13,304,213 | 3,063,850 | 75.95 |

| | | | | | |
|---|---|---|---|---|---|
| **FITC conjugated A15-peptide** | | - | 638249 | - | - |
| **A15-peptide with cleaved N-terminal amino acid** | | 141,670 | 7,944,263 | 137,796 | 9,897,684 |
| **A9-peptide (control)** | | 1,089,661 | 976,075 | 1,146,393 | 1,124,571 |

| Conjugation with FITC in 0.1 M $NaHCO_3$ pH 8.5 (5.9 mM FITC in 23:77 ratio DMSO:0.1 M $NaHCO_3$ pH 8.5) | **Gelation** | **0.1 M $NaHCO_3$** | **23:77 DMSO:0.1 M $NaHCO_3$ with FITC** | **0.1 M $NaHCO_3$, then TFA** | **23:77 DMSO:0.1 M $NaHCO_3$ with FITC, then TFA** | **Conversion rate** |
|---|---|---|---|---|---|---|
| **A15-peptide** | | 11,107,184 | 2,580,088 | 11,099,956 | 2,328,726 | |
| **FITC conjugated A15-peptide** | | - | 443245 | - | - | 76.77 |
| **A15-peptide with cleaved N-terminal amino acid** | | 98,419 | 5,065,557 | 126,352 | 7,022,611 | |
| **A9-peptide (control)** | Gelation 3 | 1,142,427 | 1,047,343 | 1,180,805 | 1,163,367 | |

(ii) Conversion rate of A15-peptide to PITC conjugated A15-peptide, and % yield for 5.9 mM FITC conjugation in 23:77 DMSO:0.1 M sodium bicarbonate ($NaHCO_3$) pH 8.5 (Fig. 3**D**).

| | |
|---|---|
| Conjugation with FITC in 0.1 M $NaHCO_3$ pH 8.5 (5.9 mM FITC in 23:77 ratio DMSO:0.1 M $NaHCO_3$ pH 8.5) **Average % yield (peptide with cleaved N-terminal amino acid / non-modified peptide * 100)** | 69.59 |
| **Average conversion rate (%)** | 75.38 |

(iii) Statistics: One-way Analysis of Variance (ANOVA) and Tukey's post-hoc Honestly Significant Difference (HSD) test on the abundance of peptide with cleaved N-terminal amino acid using 5.9 mM FITC conjugation in 23:77 DMSO:0.1 M sodium bicarbonate ($NaHCO_3$) pH 8.5 (Fig. 3**Diii**). In Tukey HSD results, conditions are abbreviated, A: $NaHCO_3$, B: $NaHCO_3$ with FITC, C: $NaHCO_3$ then TFA, D: $NaHCO_3$ with FITC then TFA.

| **ANOVA Results** | | | | | |
|---|---|---|---|---|---|
| Test | Comparison | sum_sq | df | F | PR(>F) |
| ANOVA | group | 1.60E+14 | 3.00E+00 | 4.43E+01 | 2.49E-05 |
| ANOVA | Residual | 9.61E+12 | 8.00E+00 | | |

| **Tukey HSD Results** | | | | | | | |
|---|---|---|---|---|---|---|---|
| Test | group1 | group2 | meandiff | p-adj | lower | upper | reject |
| Tukey HSD | A | B | 5.91E+06 | 7.62E-04 | 3.05E+06 | 8.78E+06 | TRUE |
| Tukey HSD | A | C | 8.16E+03 | 1.00E+00 | -2.86E+06 | 2.87E+06 | FALSE |
| Tukey HSD | A | D | 8.29E+06 | 6.93E-05 | 5.43E+06 | 1.12E+07 | TRUE |
| Tukey HSD | B | C | -5.91E+06 | 7.69E-04 | -8.77E+06 | -3.04E+06 | TRUE |
| Tukey HSD | B | D | 2.38E+06 | 1.08E-01 | -4.86E+05 | 5.24E+06 | FALSE |
| Tukey HSD | C | D | 8.28E+06 | 6.98E-05 | 5.42E+06 | 1.12E+07 | TRUE |

(iv) Gel size changes throughout in-gel Edman degradation with using 5.9 mM FITC conjugation in 23:77 DMSO:0.1 M sodium bicarbonate ($NaHCO_3$) pH 8.5 (Fig. 3**Eiii**).

| **Conjugation with FITC in 23:77 DMSO:$NaHCO_3$ - Gel size changes in Edman steps** | **Gelation 1, area ($cm^2$)** | **Gelation 2, area ($cm^2$)** | **Gelation 3, area ($cm^2$)** | **Mean (normalized to top/flat surface size in 1M Tris pH 9.5) ($cm^2$)** | **Standard deviation ($cm^2$)** |
|---|---|---|---|---|---|
| 1. 3x wash with Tris 1M pH 9.5 | 0.303 | 0.360 | 0.303 | 1.000 | 0.000 |
| 2. 2x 0.1 M $NaHCO_3$wash | 0.303 | 0.303 | 0.303 | 0.940 | 0.092 |
| 3. FITC conjugation | 0.275 | 0.303 | 0.303 | 0.912 | 0.080 |
| 4. TFA | 0.250 | 0.303 | 0.250 | 0.832 | 0.008 |
| 5. 2x 2x 0.1 M $NaHCO_3$ wash | 0.303 | 0.303 | 0.250 | 0.886 | 0.096 |
| 6. 3x wash with Tris 1M pH 8 | 0.303 | 0.360 | 0.303 | 1.000 | 0.000 |

##### (D) Fluorescence assay with azidolysine peptide

(i) For Fig. 3**G**, raw average fluorescence intensity values for each gel, and average over the replicates.

| **Condition name and number** | **Average intensity ExMre gel 1** | **Average intensity ExMre gel 2** | **Average intensity ExMre gel 3** | **Average intensity over replicates** | **Standard deviation over replicates** |
|---|---|---|---|---|---|
| (1) DMSO | 25,068.05586 | 26,800.6509 | 22,135.4875 | **24,668.06475** | 2,358.16285 |
| (2) PITC:DMSO | 23,926.83914 | 22,876.96382 | 18,189.04312 | **21,664.28203** | 3,055.08161 |
| (3) DMSO+TFA | 24,913.99409 | 25,783.96689 | 22,445.86609 | **24,381.27569** | 1,731.63821 |
| (4) PITC:DMSO+TFA | 9,763.790698 | 9,629.696378 | 6,710.793017 | **8,701.426698** | 1,725.24264 |
| (5) PITC:DMSO+TFA +trypsin | 2,020.303932 | 1,890.965337 | 2,020.303932 | **1,977.191067** | 74.673673 |

(ii) % yield calculation based on the fluorescence azidolysine read-out. The "averageIntensities(x)" represents the values obtained in Supplementary Table 5**Di** after averaging over the replicates of a given condition, where "x" represents the number associated with that condition.

| **% Yield** |
|---|
| Formula:<br>efficiencyAvg = (1 - ((averageIntensities(4) - averageIntensities(5))/(averageIntensities(1) - averageIntensities(5)))) * 100 |
| **70.366** |

(iii) One-way Analysis of Variance (ANOVA) and Tukey's post-hoc Honestly Significant Difference (HSD) test (FWER = 0.05) on the abundance of average fluorescence intensity in the gel (Fig. 3**G**). In Tukey HSD results, conditions are abbreviated, A: DMSO, B: PITC:DMSO, C: DMSO+TFA, D: PITC:DMSO+TFA, E: PITC:DMSO+TFA+trypsin

##### ANOVA Results

| | Sum of Squares | df | F | p-value |
|---|---|---|---|---|
| Group | 1.281e+09 | 4.0 | 76.7038 | 1.830e-07 |
| Residual | 4.175e+07 | 10.0 | — | — |

##### Tukey HSD Results

| Group 1 | Group 2 | Mean Diff | p-adj | Lower | Upper | Reject |
|---|---|---|---|---|---|---|
| A | B | −3003.78 | 0.4236 | −8494.41 | 2486.85 | False |
| A | C | −286.79 | 0.9998 | −5777.42 | 5203.84 | False |
| A | D | −15966.64 | 0.0 | −21457.27 | −10476.01 | True |
| A | E | −22690.87 | 0.0 | −28181.50 | −17200.25 | True |
| B | C | 2716.99 | 0.5133 | −2773.63 | 8207.62 | False |
| B | D | −12962.86 | 0.0001 | −18453.48 | −7472.23 | True |
| B | E | −19687.09 | 0.0 | −25177.72 | −14196.46 | True |
| C | D | −15679.85 | 0.0 | −21170.48 | −10189.22 | True |
| C | E | −22404.08 | 0.0 | −27894.71 | −16913.46 | True |
| D | E | −6724.24 | 0.0159 | −12214.86 | −1233.61 | True |

##### (E) Phenylthiohydantoin-phenylalanine (PTH-F) detection

For Fig. 3**H**. Analysis of PTH-F abundance was performed using PTH-F exact mass, 282.0827 ± 0.0056 Da (see **Methods:** **In-gel Edman degradation with PTH-aa detection** for details on extracting the abundance; see Supplementary Figure 6**A-B** for raw traces and mass spectra). The abundance extracted from the chromatograms are documented in the table below.

| | **Abundance (area) gel 1** | **Abundance (area) gel 2** | **Abundance (area) gel 3** | **Average abundance over replicates** | **Standard deviation across replicates** |
|---|---|---|---|---|---|
| **PTH-F negative control (DMSO+TFA)** | 787.54 | 1,196.55 | 235.46 | 739.85 | 482.31 |
| **PTH-F experiment (PITC:DMSO+TFA)** | 197,004.58 | 246,786.47 | 63,878.04 | 169,223.03 | 94,566.031 |
| **PTH-F positive control** | - | - | - | 35,445,968.85 | - |

##### PTH-neg vs PTH-F (Welch's t-test)

| Test | t | p-value |
|---|---:|---:|
| Welch's t-test | −3.086 | 0.0909 |

##### (F) Fluorescence assay with azidolysine at the second position of the peptide

For **Fig. 3I**.

| **Condition name and number** | **Average intensity ExMre gel 1** | **Average intensity ExMre gel 2** | **Average intensity ExMre gel 3** | **Average intensity over replicates** | **Standard deviation over replicates** |
|---|---|---|---|---|---|
| (1) DMSO | 3547.70033 | 5011.18561 | 2883.10230 | **3,813.99608** | 1,088.74685 |

| | | | | | |
|---|---|---|---|---|---|
| (2) TFA | 3061.03983 | 4875.80575 | 3002.57022 | **3,646.47193** | 1,065.03563 |
| (3) PITC:DMSO+TFA | 2632.65886 | 3488.24249 | 2482.75595 | **2,867.8857** | 542.44775 |
| (4) (PITC:DMSO+TFA)*2 | 848.31575 | 1728.20360 | 1051.92558 | **1,209.48164** | 460.61762 |
| (5) (PITC:DMSO+TFA)*3 | 1414.43833 | 1505.74522 | 950.63879 | **1,290.27411** | 297.65475 |
| (6) (PITC:DMSO+TFA)*3+trypsin | 628.27463 | 903.62195 | 530.42240 | **687.43966** | 193.50672 |

(ii) % yield calculation based on the fluorescence azidolysine read-out. The "averageIntensities(x)" represents the values obtained in **Supplementary Table 5Fi** after averaging over the replicates of a given condition, where "x" represents the number associated with that condition.

| **% Yield** |
|---|
| Formula:<br>efficiencyAvg = (1 - ((averageIntensities(4) - averageIntensities(6))/(averageIntensities(1) - averageIntensities(6)))) * 100 |
| 83.303 (cumulative over two rounds); 91.270 (per-round) |

(iii) One-way Analysis of Variance (ANOVA) and Tukey's post-hoc Honestly Significant Difference (HSD) test (FWER = 0.05) on the abundance of average fluorescence intensity in the gel (Fig. 3I). In Tukey HSD results, conditions are abbreviated, A: DMSO, B: DMSO+TFA, C: PITC:DMSO+TFA, D: (PITC:DMSO+TFA)*2, E: (PITC:DMSO+TFA)*3, F: (PITC:DMSO+TFA)*3+trypsin.

**ANOVA Results**

| | **Sum of Squares** | **df** | **F** | **p-value** |
|---|---|---|---|---|
| Group | 2.767e+07 | 5.0 | 11.2474 | 0.00034 |
| Residual | 5.904e+06 | 12.0 | — | — |

**Tukey HSD Results**

| **Group 1** | **Group 2** | **Mean Diff** | **p-adj** | **Lower** | **Upper** | **Reject** |
|---|---|---|---|---|---|---|
| A | B | −167.52 | 0.9996 | −2091.26 | 1756.22 | False |
| A | C | −946.11 | 0.5836 | −2869.85 | 977.63 | False |
| A | D | −2604.51 | 0.0068 | −4528.25 | −680.77 | True |
| A | E | −2523.72 | 0.0086 | −4447.46 | −599.98 | True |
| A | F | −3126.56 | 0.0016 | −5050.30 | −1202.82 | True |
| B | C | −778.59 | 0.7487 | −2702.33 | 1145.15 | False |
| B | D | −2436.99 | 0.011 | −4360.73 | −513.25 | True |
| B | E | −2356.20 | 0.014 | −4279.94 | −432.46 | True |
| B | F | −2959.03 | 0.0025 | −4882.77 | −1035.29 | True |
| C | D | −1658.40 | 0.1072 | −3582.14 | 265.34 | False |
| C | E | −1577.61 | 0.1342 | −3501.35 | 346.13 | False |

| | | | | | | |
|---|---|---|---|---|---|---|
| C | F | −2180.45 | 0.0235 | −4104.19 | −256.71 | True |
| D | E | 80.79 | 1.0 | −1842.95 | 2004.53 | False |
| D | F | −522.04 | 0.9361 | −2445.78 | 1401.70 | False |
| E | F | −602.83 | 0.8907 | −2526.57 | 1320.91 | False |

**(G) Fluorescence assay with azidolysine peptide stained with Atto647N-alkyne**

For Supplementary Figure 6F.

| Condition name and number | Average intensity ExMre gel 1 | Average intensity ExMre gel 2 | Average intensity ExMre gel 3 | Average intensity over replicates | Standard deviation over replicates |
|---|---|---|---|---|---|
| (1) DMSO | 13805.07789 | 9646.95120 | 6495.15907 | **9982.39605** | 3666.48616 |
| (2) PITC:DMSO | 12597.40173 | 9010.19420 | 7036.96407 | **9548.18667** | 2818.98806 |
| (3) DMSO+TFA | 12204.50065 | 11381.35733 | 8016.12142 | **10533.99313** | 2219.04252 |
| (4) PITC:DMSO+TFA | 5404.21221 | 7003.63563 | 4582.32606 | **5663.39130** | 1231.28606 |
| (5) PITC:DMSO+TFA+trypsin | 6741.01583 | 5498.30417 | 3833.47506 | **5357.59835** | 1458.86837 |

(ii) % yield calculation based on the fluorescence azidolysine read-out. The "averageIntensities(x)" represents the values obtained in Supplementary Table 5G after averaging over the replicates of a given condition, where "x" represents the number associated with that condition.

| **% Yield** |
|---|
| Formula:<br>efficiencyAvg = (1 - ((averageIntensities(4) - averageIntensities(5))/(averageIntensities(1) - averageIntensities(5)))) * 100 |
| **93.388** |

(iii) One-way Analysis of Variance (ANOVA) and Tukey's post-hoc Honestly Significant Difference (HSD) test (FWER = 0.05) on the abundance of average fluorescence intensity in the gel (Fig. 6F). In Tukey HSD results, conditions are abbreviated, A: DMSO, B: PITC:DMSO, C: DMSO+TFA, D: PITC:DMSO+TFA, E: PITC:DMSO+TFA+trypsin.

**ANOVA Results (Fig. 6F)**

| | **Sum of Squares** | **df** | **F** | **p-value** |
|---|---|---|---|---|
| Group | 7.486e+07 | 4.0 | 3.1236 | 0.065583 |
| Residual | 5.992e+07 | 10.0 | — | — |

**Tukey Results (Fig. 6F)**

| **Group 1** | **Group 2** | **Mean Diff** | **p-adj** | **Lower** | **Upper** | **Reject** |
|---|---|---|---|---|---|---|
| A | B | −434.21 | 0.9994 | −7011.80 | 6143.38 | False |
| A | C | 551.60 | 0.9985 | −6025.99 | 7129.19 | False |
| A | D | −4319.00 | 0.2683 | −10896.60 | 2258.59 | False |

| | | | | | | |
|---|---|---|---|---|---|---|
| A | E | −4624.80 | 0.2173 | −11202.39 | 1952.79 | False |
| B | C | 985.81 | 0.9862 | −5591.78 | 7563.40 | False |
| B | D | −3884.80 | 0.3561 | −10462.39 | 2692.80 | False |
| B | E | −4190.59 | 0.2924 | −10768.18 | 2387.00 | False |
| C | D | −4870.60 | 0.1824 | −11448.19 | 1706.99 | False |
| C | E | −5176.39 | 0.1458 | −11753.99 | 1401.20 | False |
| D | E | −305.79 | 0.9999 | −6883.38 | 6271.80 | False |

**(H) Fluorescence assay with azidolysine at the second position of the peptide with Atto647N-alkyne**

For Supplementary Figure 6G.
(i)

| Condition name and number | Average intensity ExMre gel 1 | Average intensity ExMre gel 2 | Average intensity ExMre gel 3 | Average intensity over replicates | Standard deviation over replicates |
|---|---|---|---|---|---|
| (1) DMSO | 3768.32012 | 4689.31041 | 3394.27779 | **3950.63607** | 666.48838 |
| (2) TFA | 3033.06153 | 4647.50800 | 3078.45588 | **3586.34181** | 919.27713 |
| (3) PITC:DMSO+TFA | 3690.77265 | 4428.29530 | 2627.86793 | **3582.31196** | 905.10081 |
| (4) (PITC:DMSO+TFA)*2 | 1978.84341 | 2178.77923 | 1446.19021 | **1867.93762** | 378.67762 |
| (5) (PITC:DMSO+TFA)*3 | 2296.46891 | 1997.76895 | 1244.29027 | **1846.17604** | 542.22254 |
| (6) (PITC:DMSO+TFA)*3+trypsin | 1305.20325 | 1283.27603 | 789.55444 | **1126.01124** | 291.58632 |

(ii) % yield calculation based on the fluorescence azidolysine read-out. The "averageIntensities(x)" represents the values obtained in Supplementary Table 5**Hi** after averaging over the replicates of a given condition, where "x" represents the number associated with that condition.

| **% Yield** |
|---|
| Formula:<br>efficiencyAvg = (1 - ((averageIntensities(4) - averageIntensities(6))/(averageIntensities(1) - averageIntensities(6)))) * 100 |
| 73.734 (cumulative over 2 rounds); 85.868 (per-round) |

(iii) One-way Analysis of Variance (ANOVA) and Tukey's post-hoc Honestly Significant Difference (HSD) test (FWER = 0.05) on the abundance of average fluorescence intensity in the gel (Supplementary Figure 6G). In Tukey HSD results, conditions are abbreviated, A: DMSO, B: DMSO+TFA, C: PITC:DMSO+TFA, D: (PITC:DMSO+TFA)*2, E: (PITC:DMSO+TFA)*3, F: (PITC:DMSO+TFA)*3+trypsin.

**ANOVA Results (Fig. 6G)**

| | Sum of Squares | df | F | p-value |
|---|---|---|---|---|
| Group | 2.105e+07 | 5.0 | 9.6021 | 0.000707 |

| | | | | |
|---|---|---|---|---|
| Residual | 5.262e+06 | 12.0 | — | — |

**Tukey HSD Results (Fig. 6G)**

| Group 1 | Group 2 | Mean Diff | p-adj | Lower | Upper | Reject |
|---|---|---|---|---|---|---|
| A | B | −364.29 | 0.9817 | −2180.36 | 1451.77 | False |
| A | C | −368.32 | 0.9808 | −2184.39 | 1447.74 | False |
| A | D | −2082.70 | 0.0218 | −3898.76 | −266.63 | True |
| A | E | −2104.46 | 0.0203 | −3920.53 | −288.39 | True |
| A | F | −2824.62 | 0.0023 | −4640.69 | −1008.56 | True |
| B | C | −4.03 | 1.0 | −1820.10 | 1812.04 | False |
| B | D | −1718.40 | 0.0676 | −3534.47 | 97.66 | False |
| B | E | −1740.17 | 0.0632 | −3556.23 | 75.90 | False |
| B | F | −2460.33 | 0.0067 | −4276.40 | −644.27 | True |
| C | D | −1714.37 | 0.0684 | −3530.44 | 101.69 | False |
| C | E | −1736.14 | 0.064 | −3552.20 | 79.93 | False |
| C | F | −2456.30 | 0.0068 | −4272.37 | −640.24 | True |
| D | E | −21.76 | 1.0 | −1837.83 | 1794.30 | False |
| D | F | −741.93 | 0.7418 | −2557.99 | 1074.14 | False |
| E | F | −720.16 | 0.7633 | −2536.23 | 1095.90 | False |

**Supplementary Table 6**
Values and analysis for the LC/QToF trypsinization assay from Fig. 5B.

(i) Raw data values for the area under the curve (AUC) of the chromatogram of various species for PITCto DMSO (1:1000 ratio PITC:DMSO) for conjugation over multiple rounds (Fig. 5B) (extracted based on the exact mass, see Methods for more details, and Source Data for raw traces). Three tables for the three separate gelation solutions. The cells highlighted in grey are the ones used to calculate the conversion rate (1-(solvent with PITC["A15-peptide"] / solvent ["A15-peptide"])). A15-peptide stands for AGGAGGLLGGSRGGK{acr}. Cells with '-' means that the species was not detected in that sample using the automatic extraction.

| Gelation 1 | **DMSO** | **DMSO with PITC** | **DMSO, then TFA** | **DMSO with PITC, then TFA** | **1 round, then DMSO with PITC** | **1 round, then DMSO with PITC, then TFA** | **2 rounds, then DMSO with PITC** | **2 rounds, then DMSO with PITC, then TFA** |
|---|---|---|---|---|---|---|---|---|
| **A15-peptide** | 14,152,440 | 234,022 | 16,895,166 | 241,708 | 55,744 | 55,988 | 127,394 | 105,796 |
| **PITC conjugated A15-peptide** | - | 8,528,288 | - | 31,819 | 94,071 | - | 42,671 | - |
| **A15-peptide with cleaved N-terminal amino acid** | 76,571 | 7,159,311 | 101,167 | 11,042,843 | 110,995 | 302,012 | 69,181 | 103,014 |
| **PITC conjugated A15-peptide with cleaved N-terminal amino acid** | - | 354,635 | - | 23,086 | 11,642,582 | 605,347 | 937,425 | - |
| **A15-peptide with two cleaved N-terminal amino acids** | 73,330 | 21,890 | 62,400 | 187,758 | 624,556 | 6,975,236 | 214,828 | 611,031 |
| **PITC conjugated A15-peptide with two cleaved N-terminal amino acid** | 64,524 | 30,557 | 78,959 | 4,345 | 271,111 | 39,892 | 11,299,778 | 1,992,023 |
| **A15-peptide with three cleaved N-terminal amino acids** | 78,578 | 5,505 | 10,341 | 29,603 | 26,503 | 83,453 | 444,363 | 2,076,406 |
| **A9-peptide (control)** | 1,717,854 | 1,950,484 | 1,916,119 | 2,305,651 | 1,992,306 | 2,085,178 | 1,726,787 | 1,770,300 |

| Gelation 2 | **DMSO** | **DMSO with PITC** | **DMSO, then TFA** | **DMSO with PITC, then TFA** | **1 round, then DMSO with PITC** | **1 round, then DMSO with PITC, then TFA** | **2 rounds, then DMSO with PITC** | **2 rounds, then DMSO with PITC, then TFA** |
|---|---|---|---|---|---|---|---|---|

| | | | | | | | | |
|---|---|---|---|---|---|---|---|---|
| **A15-peptide** | 19,664,727 | 168,662 | 18,026,845 | 382,515 | 61,289 | 73,152 | 23,823 | 175,778 |
| **PITC conjugated A15-peptide** | - | 10,588,346 | - | 37,486 | 119,421 | - | 39,940 | - |
| **A15-peptide with cleaved N-terminal amino acid** | 114,996 | 9,038,666 | 101,342 | 16,031,614 | 186,135 | 323,530 | 67,534 | 159,065 |
| **PITC conjugated A15-peptide with cleaved N-terminal amino acid** | - | 464,449 | - | 28,326 | 18,012,585 | 679,072 | 939,101 | - |
| **A15-peptide with two cleaved N-terminal amino acids** | 97,726 | 74,181 | 72,097 | 270,900 | 973,920 | 7,341,481 | 225,736 | 952,458 |
| **PITC conjugated A15-peptide with two cleaved N-terminal amino acid** | 93,023 | 45,738 | 82,455 | 6,666 | 426,174 | 71,780 | 9,759,509 | 2,432,709 |
| **A15-peptide with three cleaved N-terminal amino acids** | 13,299 | - | 13,090 | 43,111 | 29,064 | 152,920 | 421,886 | 2,545,115 |
| **A9-peptide (control)** | 2,329,190 | 1,932,427 | 1,994,133 | 2,169,287 | 2,175,041 | 1,959,964 | 2,075,030 | 1,728,591 |

| Gelation 3 | **DMSO** | **DMSO with PITC** | **DMSO, then TFA** | **DMSO with PITC, then TFA** | **1 round, then DMSO with PITC** | **1 round, then DMSO with PITC, then TFA** | **2 rounds, then DMSO with PITC** | **2 rounds, then DMSO with PITC, then TFA** |
|---|---|---|---|---|---|---|---|---|
| **A15-peptide** | 13,935,909 | 283,570 | 12,028,033 | 271,999 | 48,147 | 128,729 | 113,952 | 113,403 |
| **PITC conjugated A15-peptide** | - | 5,677,587 | - | 16,631 | 102,012 | - | 37,689 | - |
| **A15-peptide with cleaved N-terminal amino acid** | 77,417 | 6,630,107 | 72,205 | 11,187,479 | 141,646 | 344,558 | 82,374 | 114,200 |
| **PITC conjugated A15-peptide with cleaved N-terminal amino acid** | - | 318,844 | - | 18,965 | 14,851,035 | 786,992 | 1,140,599 | - |
| **A15-peptide with two cleaved N-terminal amino acids** | 68,099 | 20,399 | 44,486 | 207,266 | 816,241 | 6,151,386 | 49,806 | 561,106 |
| **PITC conjugated A15-peptide with two cleaved N-terminal amino acid** | 71,146 | 31,762 | 71,623 | 3,814 | 332,970 | 56,436 | 9,159,739 | 1,296,611 |
| **A15-peptide with three** | 6,102 | - | 6,064 | 35,149 | 23,037 | 80,978 | 392,714 | 1,781,781 |

| | | | | | | | | |
|---|---|---|---|---|---|---|---|---|
| **cleaved N-terminal amino acids** | | | | | | | | |
| **A9-peptide (control)** | 1,974,781 | 1,848,514 | 2,181,989 | 2,129,434 | 2,027,558 | 1,856,479 | 1,833,510 | 1,938,633 |

(ii) Conversion rate of A15-peptide to PITC conjugated A15-peptide, and % yield for conjugation with PITC to DMSO (1:1000 ratio PITC:DMSO) over multiple rounds (Fig. 5**B**).

| | |
|---|---|
| **First round average % yield (peptide with cleaved N-terminal amino acid / non-modified peptide * 100)** | 82.43 |
| **First round average conversion rate from A15-peptide to PITC conjugated to A15-peptide (%)** | 98.48 |
| **Second round average conversion rate from A15-peptide with cleaved N-terminal amino acid to PITC conjugated to A15-peptide with cleaved N-terminal amino acid (%)** | 98.86 |
| **Third round average conversion rate from A15-peptide with two cleaved N-terminal amino acids to PITC conjugated to A15-peptide with two cleaved N-terminal amino acids (%)** | 97.68 |

(iii) One-way Analysis of Variance (ANOVA) and Tukey's post-hoc Honestly Significant Difference (HSD) test (FWER = 0.05) on the abundance of average fluorescence intensity in the gel (Fig. 5**B**). In Tukey HSD results, conditions are abbreviated, A: DMSO, B: DMSO:PITC, C: DMSO+TFA, D: DMSO:PITC+TFA (1), E: DMSO:PITC (2), F: DMSO:PITC+TFA (2), G: DMSO:PITC (3), H: DMSO:PITC+TFA (3).

**ANOVA results Fig. 5Bi, non-modified peptide**

| | **Sum of Squares** | **df** | **F** | **p-value** |
|---|---|---|---|---|
| Group | 1.100e+15 | 7.0 | 60.7059 | 2.459e-10 |
| Residual | 4.143e+13 | 16.0 | — | — |

**Tukey HSD results Fig. 5Bi, non-modified peptide**

| **Group 1** | **Group 2** | **Mean Diff** | **p-adj** | **Lower** | **Upper** | **Reject** |
|---|---|---|---|---|---|---|
| A | B | −15688940.67 | 0.0 | −20237873.02 | −11140008.32 | True |
| A | C | −267677.33 | 1.0 | −4816609.68 | 4281255.02 | False |
| A | D | −15618951.33 | 0.0 | −20167883.68 | −11070018.98 | True |
| A | E | −15862632.0 | 0.0 | −20411564.35 | −11313699.65 | True |
| A | F | −15831735.67 | 0.0 | −20380668.02 | −11282803.32 | True |
| A | G | −15829302.33 | 0.0 | −20378234.68 | −11280369.98 | True |
| A | H | −15786033.0 | 0.0 | −20334965.35 | −11237100.65 | True |
| B | C | 15421263.33 | 0.0 | 10872330.98 | 19970195.68 | True |

| | | | | | | |
|---|---|---|---|---|---|---|
| B | D | 69989.33 | 1.0 | −4478943.02 | 4618921.68 | False |
| B | E | −173691.33 | 1.0 | −4722623.68 | 4375241.02 | False |
| B | F | −142795.0 | 1.0 | −4691727.35 | 4406137.35 | False |
| B | G | −140361.67 | 1.0 | −4689294.02 | 4408570.68 | False |
| B | H | −97092.33 | 1.0 | −4646024.68 | 4451840.02 | False |
| C | D | −15351274.0 | 0.0 | −19900206.35 | −10802341.65 | True |
| C | E | −15594954.67 | 0.0 | −20143887.02 | −11046022.32 | True |
| C | F | −15564058.33 | 0.0 | −20112990.68 | −11015125.98 | True |
| C | G | −15561625.0 | 0.0 | −20110557.35 | −11012692.65 | True |
| C | H | −15518355.67 | 0.0 | −20067288.02 | −10969423.32 | True |
| D | E | −243680.67 | 1.0 | −4792613.02 | 4305251.68 | False |
| D | F | −212784.33 | 1.0 | −4761716.68 | 4336148.02 | False |
| D | G | −210351.0 | 1.0 | −4759283.35 | 4338581.35 | False |
| D | H | −167081.67 | 1.0 | −4716014.02 | 4381850.68 | False |
| E | F | 30896.33 | 1.0 | −4518036.02 | 4579828.68 | False |
| E | G | 33329.67 | 1.0 | −4515602.68 | 4582262.02 | False |
| E | H | 76599.0 | 1.0 | −4472333.35 | 4625531.35 | False |
| F | G | 2433.33 | 1.0 | −4546499.02 | 4551365.68 | False |
| F | H | 45702.67 | 1.0 | −4503229.68 | 4594635.02 | False |

**ANOVA results Fig. 5Bii, peptide conjugated to PITC**

| | Sum of Squares | df | F | p-value |
|---|---|---|---|---|
| Group | 1.783e+14 | 7.0 | 33.4994 | 2.126e-08 |
| Residual | 1.216e+13 | 16.0 | — | — |

**Tukey HSD results Fig. 5Bii, peptide conjugated to PITC**

| Group 1 | Group 2 | Mean Diff | p-adj | Lower | Upper | Reject |
|---|---|---|---|---|---|---|
| A | B | 8264740.33 | 0.0 | 5800105.60 | 10729375.06 | True |
| A | C | 0.0 | 1.0 | −2464634.73 | 2464634.73 | False |
| A | D | 28645.33 | 1.0 | −2435989.40 | 2493280.06 | False |
| A | E | 105168.0 | 1.0 | −2359466.73 | 2569802.73 | False |
| A | F | 0.0 | 1.0 | −2464634.73 | 2464634.73 | False |
| A | G | 40100.0 | 1.0 | −2424534.73 | 2504734.73 | False |
| A | H | 0.0 | 1.0 | −2464634.73 | 2464634.73 | False |
| B | C | −8264740.33 | 0.0 | −10729375.06 | −5800105.60 | True |
| B | D | −8236095.0 | 0.0 | −10700729.73 | −5771460.27 | True |

| | | | | | | |
|---|---|---|---|---|---|---|
| B | E | −8159572.33 | 0.0 | −10624207.06 | −5694937.60 | True |
| B | F | −8264740.33 | 0.0 | −10729375.06 | −5800105.60 | True |
| B | G | −8224640.33 | 0.0 | −10689275.06 | −5760005.60 | True |
| B | H | −8264740.33 | 0.0 | −10729375.06 | −5800105.60 | True |
| C | D | 28645.33 | 1.0 | −2435989.40 | 2493280.06 | False |
| C | E | 105168.0 | 1.0 | −2359466.73 | 2569802.73 | False |
| C | F | 0.0 | 1.0 | −2464634.73 | 2464634.73 | False |
| C | G | 40100.0 | 1.0 | −2424534.73 | 2504734.73 | False |
| C | H | 0.0 | 1.0 | −2464634.73 | 2464634.73 | False |
| D | E | 76522.67 | 1.0 | −2388112.06 | 2541157.40 | False |
| D | F | −28645.33 | 1.0 | −2493280.06 | 2435989.40 | False |
| D | G | 11454.67 | 1.0 | −2453180.06 | 2476089.40 | False |
| D | H | −28645.33 | 1.0 | −2493280.06 | 2435989.40 | False |
| E | F | −105168.0 | 1.0 | −2569802.73 | 2359466.73 | False |
| E | G | −65068.0 | 1.0 | −2529702.73 | 2399566.73 | False |
| E | H | −105168.0 | 1.0 | −2569802.73 | 2359466.73 | False |
| F | G | 40100.0 | 1.0 | −2424534.73 | 2504734.73 | False |
| F | H | 0.0 | 1.0 | −2464634.73 | 2464634.73 | False |
| G | H | −40100.0 | 1.0 | −2504734.73 | 2424534.73 | False |

**ANOVA results Fig. 5Biii, peptide with cleaved N-terminal amino acid**

| | Sum of Squares | df | F | p-value |
|---|---|---|---|---|
| Group | 4.934e+14 | 7.0 | 58.3295 | 3.334e-10 |
| Residual | 1.934e+13 | 16.0 | — | — |

**Tukey HSD Fig. 5Biii, peptide with cleaved N-terminal amino acid**

| Group 1 | Group 2 | Mean Diff | p-adj | Lower | Upper | Reject |
|---|---|---|---|---|---|---|
| A | B | 7519700.0 | 0.0 | 4412080.30 | 10627319.70 | True |
| A | C | 1910.0 | 1.0 | −3105709.70 | 3109529.70 | False |
| A | D | 12664317.33 | 0.0 | 9556697.64 | 15771937.03 | True |
| A | E | 56597.33 | 1.0 | −3051022.36 | 3164217.03 | False |
| A | F | 233705.33 | 1.0 | −2873914.36 | 3341325.03 | False |
| A | G | −16631.67 | 1.0 | −3124251.36 | 3090988.03 | False |
| A | H | 35765.0 | 1.0 | −3071854.70 | 3143384.70 | False |
| B | C | −7517790.0 | 0.0 | −10625409.70 | −4410170.30 | True |
| B | D | 5144617.33 | 0.0006 | 2036997.64 | 8252237.03 | True |

| | | | | | | |
|---|---|---|---|---|---|---|
| B | E | −7463102.67 | 0.0 | −10570722.36 | −4355482.97 | True |
| B | F | −7285994.67 | 0.0 | −10393614.36 | −4178374.97 | True |
| B | G | −7536331.67 | 0.0 | −10643951.36 | −4428711.97 | True |
| B | H | −7483935.0 | 0.0 | −10591554.70 | −4376315.30 | True |
| C | D | 12662407.33 | 0.0 | 9554787.64 | 15770027.03 | True |
| C | E | 54687.33 | 1.0 | −3052932.36 | 3162307.03 | False |
| C | F | 231795.33 | 1.0 | −2875824.36 | 3339415.03 | False |
| C | G | −18541.67 | 1.0 | −3126161.36 | 3089078.03 | False |
| C | H | 33855.0 | 1.0 | −3073764.70 | 3141474.70 | False |
| D | E | −12607720.0 | 0.0 | −15715339.70 | −9500100.30 | True |
| D | F | −12430612.0 | 0.0 | −15538231.70 | −9322992.30 | True |
| D | G | −12680949.0 | 0.0 | −15788568.70 | −9573329.30 | True |
| D | H | −12628552.33 | 0.0 | −15736172.03 | −9520932.64 | True |
| E | F | 177108.0 | 1.0 | −2930511.70 | 3284727.70 | False |
| E | G | −73229.0 | 1.0 | −3180848.70 | 3034390.70 | False |
| E | H | −20832.33 | 1.0 | −3128452.03 | 3086787.36 | False |
| F | G | −250337.0 | 1.0 | −3357956.70 | 2857282.70 | False |
| F | H | −197940.33 | 1.0 | −3305560.03 | 2909679.36 | False |
| G | H | 52396.67 | 1.0 | −3055223.03 | 3160016.36 | False |

**ANOVA results Fig. 5Biv, peptide with cleaved N-terminal amino acid**

| | Sum of Squares | df | F | p-value |
|---|---|---|---|---|
| Group | 5.576e+14 | 7.0 | 62.6499 | 1.933e-10 |
| Residual | 2.034e+13 | 16.0 | — | — |

**Tukey HSD Fig. 5Biv, peptide with cleaved N-terminal amino acid**

| Group 1 | Group 2 | Mean Diff | p-adj | Lower | Upper | Reject |
|---|---|---|---|---|---|---|
| A | B | 379309.33 | 0.9999 | −2808281.58 | 3566900.25 | False |
| A | C | 0.0 | 1.0 | −3187590.91 | 3187590.91 | False |
| A | D | 23459.0 | 1.0 | −3164131.91 | 3211049.91 | False |
| A | E | 14835400.67 | 0.0 | 11647809.75 | 18022991.58 | True |
| A | F | 690470.33 | 0.9936 | −2497120.58 | 3878061.25 | False |
| A | G | 1005708.33 | 0.9494 | −2181882.58 | 4193299.25 | False |
| A | H | 0.0 | 1.0 | −3187590.91 | 3187590.91 | False |
| B | C | −379309.33 | 0.9999 | −3566900.25 | 2808281.58 | False |
| B | D | −355850.33 | 0.9999 | −3543441.25 | 2831740.58 | False |

| | | | | | | |
|---|---|---|---|---|---|---|
| B | E | 14456091.33 | 0.0 | 11268500.42 | 17643682.25 | True |
| B | F | 311161.0 | 1.0 | −2876429.91 | 3498751.91 | False |
| B | G | 626399.0 | 0.9964 | −2561191.91 | 3813989.91 | False |
| B | H | −379309.33 | 0.9999 | −3566900.25 | 2808281.58 | False |
| C | D | 23459.0 | 1.0 | −3164131.91 | 3211049.91 | False |
| C | E | 14835400.67 | 0.0 | 11647809.75 | 18022991.58 | True |
| C | F | 690470.33 | 0.9936 | −2497120.58 | 3878061.25 | False |
| C | G | 1005708.33 | 0.9494 | −2181882.58 | 4193299.25 | False |
| C | H | 0.0 | 1.0 | −3187590.91 | 3187590.91 | False |
| D | E | 14811941.67 | 0.0 | 11624350.75 | 17999532.58 | True |
| D | F | 667011.33 | 0.9948 | −2520579.58 | 3854602.25 | False |
| D | G | 982249.33 | 0.9551 | −2205341.58 | 4169840.25 | False |
| D | H | −23459.0 | 1.0 | −3211049.91 | 3164131.91 | False |
| E | F | −14144930.33 | 0.0 | −17332521.25 | −10957339.42 | True |
| E | G | −13829692.33 | 0.0 | −17017283.25 | −10642101.42 | True |
| E | H | −14835400.67 | 0.0 | −18022991.58 | −11647809.75 | True |
| F | G | 315238.0 | 1.0 | −2872352.91 | 3502828.91 | False |
| F | H | −690470.33 | 0.9936 | −3878061.25 | 2497120.58 | False |
| G | H | −1005708.33 | 0.9494 | −4193299.25 | 2181882.58 | False |

**ANOVA results Fig. 5Bv, peptide with two cleaved N-terminal amino acids**

| | Sum of Squares | df | F | p-value |
|---|---|---|---|---|
| **Group** | 1.137e+14 | 7.0 | 282.0716 | 1.520e-15 |
| **Residual** | 9.210e+11 | 16.0 | — | — |

**Tukey HSD Fig. 5Bv, peptide with two cleaved N-terminal amino acids**

| Group 1 | Group 2 | Mean Diff | p-adj | Lower | Upper | Reject |
|---|---|---|---|---|---|---|
| A | B | −40895.0 | 1.0 | −719117.32 | 637327.32 | False |
| A | C | −20057.33 | 1.0 | −698279.65 | 658164.98 | False |
| A | D | 142256.33 | 0.9947 | −535965.98 | 820478.65 | False |
| A | E | 725187.33 | 0.0318 | 46965.02 | 1403409.65 | True |
| A | F | 6742982.67 | 0.0 | 6064760.35 | 7421204.98 | True |
| A | G | 83738.33 | 0.9998 | −594483.98 | 761960.65 | False |
| A | H | 628480.0 | 0.0799 | −49742.32 | 1306702.32 | False |
| B | C | 20837.67 | 1.0 | −657384.65 | 699059.98 | False |
| B | D | 183151.33 | 0.9775 | −495070.98 | 861373.65 | False |

| | | | | | | |
|---|---|---|---|---|---|---|
| B | E | 766082.33 | 0.0213 | 87860.02 | 1444304.65 | True |
| B | F | 6783877.67 | 0.0 | 6105655.35 | 7462099.98 | True |
| B | G | 124633.33 | 0.9976 | −553588.98 | 802855.65 | False |
| B | H | 669375.0 | 0.0544 | −8847.32 | 1347597.32 | False |
| C | D | 162313.67 | 0.9886 | −515908.65 | 840535.98 | False |
| C | E | 745244.67 | 0.0261 | 67022.35 | 1423466.98 | True |
| C | F | 6763040.0 | 0.0 | 6084817.68 | 7441262.32 | True |
| C | G | 103795.67 | 0.9993 | −574426.65 | 782017.98 | False |
| C | H | 648537.33 | 0.0662 | −29684.98 | 1326759.65 | False |
| D | E | 582931.0 | 0.1207 | −95291.32 | 1261153.32 | False |
| D | F | 6600726.33 | 0.0 | 5922504.02 | 7278948.65 | True |
| D | G | −58518.0 | 1.0 | −736740.32 | 619704.32 | False |
| D | H | 486223.67 | 0.2695 | −191998.65 | 1164445.98 | False |
| E | F | 6017795.33 | 0.0 | 5339573.02 | 6696017.65 | True |
| E | G | −641449.0 | 0.0708 | −1319671.32 | 36773.32 | False |
| E | H | −96707.33 | 0.9995 | −774929.65 | 581514.98 | False |
| F | G | −6659244.33 | 0.0 | −7337466.65 | −5981022.02 | True |
| F | H | −6114502.67 | 0.0 | −6792724.98 | −5436280.35 | True |
| G | H | 544741.67 | 0.1681 | −133480.65 | 1222963.98 | False |

**ANOVA results Fig. 5Bvi, peptide with two cleaved N-terminal amino acid conjugated to PITC**

| | Sum of Squares | df | F | p-value |
|---|---|---|---|---|
| Group | 2.564e+14 | 7.0 | 188.6475 | 3.647e-14 |
| Residual | 3.107e+12 | 16.0 | — | — |

**Tukey HSD Fig. 5Bvi, peptide with two cleaved N-terminal amino acid conjugated to PITC**

| Group 1 | Group 2 | Mean Diff | p-adj | Lower | Upper | Reject |
|---|---|---|---|---|---|---|
| A | B | −40212.0 | 1.0 | −1285870.33 | 1205446.33 | False |
| A | C | 1448.0 | 1.0 | −1244210.33 | 1247106.33 | False |
| A | D | −71289.33 | 1.0 | −1316947.66 | 1174368.99 | False |
| A | E | 267187.33 | 0.994 | −978470.99 | 1512845.66 | False |
| A | F | −20195.0 | 1.0 | −1265853.33 | 1225463.33 | False |
| A | G | 9996777.67 | 0.0 | 8751119.34 | 11242435.99 | True |
| A | H | 1830883.33 | 0.0022 | 585225.00 | 3076541.66 | True |
| B | C | 41660.0 | 1.0 | −1203998.33 | 1287318.33 | False |
| B | D | −31077.33 | 1.0 | −1276735.66 | 1214580.99 | False |

| | | | | | | |
|---|---|---|---|---|---|---|
| B | E | 307399.33 | 0.9864 | −938258.99 | 1553057.66 | False |
| B | F | 20017.0 | 1.0 | −1225641.33 | 1265675.33 | False |
| B | G | 10036989.67 | 0.0 | 8791331.34 | 11282647.99 | True |
| B | H | 1871095.33 | 0.0017 | 625437.00 | 3116753.66 | True |
| C | D | −72737.33 | 1.0 | −1318395.66 | 1172920.99 | False |
| C | E | 265739.33 | 0.9942 | −979918.99 | 1511397.66 | False |
| C | F | −21643.0 | 1.0 | −1267301.33 | 1224015.33 | False |
| C | G | 9995329.67 | 0.0 | 8749671.34 | 11240987.99 | True |
| C | H | 1829435.33 | 0.0022 | 583777.00 | 3075093.66 | True |
| D | E | 338476.67 | 0.9768 | −907181.66 | 1584134.99 | False |
| D | F | 51094.33 | 1.0 | −1194563.99 | 1296752.66 | False |
| D | G | 10068067.0 | 0.0 | 8822408.67 | 11313725.33 | True |
| D | H | 1902172.67 | 0.0015 | 656514.34 | 3147830.99 | True |
| E | F | −287382.33 | 0.9907 | −1533040.66 | 958275.99 | False |
| E | G | 9729590.33 | 0.0 | 8483932.00 | 10975248.66 | True |
| E | H | 1563696.0 | 0.0091 | 318037.67 | 2809354.33 | True |
| F | G | 10016972.67 | 0.0 | 8771314.34 | 11262630.99 | True |
| F | H | 1851078.33 | 0.0019 | 605420.00 | 3096736.66 | True |
| G | H | −8165894.33 | 0.0 | −9411552.66 | −6920236.00 | True |

**ANOVA results Fig. 5Bvii, peptide with three cleaved N-terminal amino acids**

| | Sum of Squares | df | F | p-value |
|---|---|---|---|---|
| Group | 1.137e+13 | 7.0 | 85.3655 | 1.791e-11 |
| Residual | 3.044e+11 | 16.0 | — | — |

**Tukey HSD Fig. 5Bvii, peptide with three cleaved N-terminal amino acids**

| Group 1 | Group 2 | Mean Diff | p-adj | Lower | Upper | Reject |
|---|---|---|---|---|---|---|
| A | B | −30824.67 | 1.0 | −420740.68 | 359091.35 | False |
| A | C | −22828.0 | 1.0 | −412744.02 | 367088.02 | False |
| A | D | 3294.67 | 1.0 | −386621.35 | 393210.68 | False |
| A | E | −6458.33 | 1.0 | −396374.35 | 383457.68 | False |
| A | F | 73124.0 | 0.9973 | −316792.02 | 463040.02 | False |
| A | G | 386994.67 | 0.0525 | −2921.35 | 776910.68 | False |
| A | H | 2101774.33 | 0.0 | 1711858.32 | 2491690.35 | True |
| B | C | 7996.67 | 1.0 | −381919.35 | 397912.68 | False |
| B | D | 34119.33 | 1.0 | −355796.68 | 424035.35 | False |

| | | | | | | |
|---|---|---|---|---|---|---|
| B | E | 24366.33 | 1.0 | −365549.68 | 414282.35 | False |
| B | F | 103948.67 | 0.979 | −285967.35 | 493864.68 | False |
| B | G | 417819.33 | 0.0313 | 27903.32 | 807735.35 | True |
| B | H | 2132599.0 | 0.0 | 1742682.98 | 2522515.02 | True |
| C | D | 26122.67 | 1.0 | −363793.35 | 416038.68 | False |
| C | E | 16369.67 | 1.0 | −373546.35 | 406285.68 | False |
| C | F | 95952.0 | 0.9866 | −293964.02 | 485868.02 | False |
| C | G | 409822.67 | 0.0358 | 19906.65 | 799738.68 | True |
| C | H | 2124602.33 | 0.0 | 1734686.32 | 2514518.35 | True |
| D | E | −9753.0 | 1.0 | −399669.02 | 380163.02 | False |
| D | F | 69829.33 | 0.998 | −320086.68 | 459745.35 | False |
| D | G | 383700.0 | 0.0554 | −6216.02 | 773616.02 | False |
| D | H | 2098479.67 | 0.0 | 1708563.65 | 2488395.68 | True |
| E | F | 79582.33 | 0.9955 | −310333.68 | 469498.35 | False |
| E | G | 393453.0 | 0.0471 | 3536.98 | 783369.02 | True |
| E | H | 2108232.67 | 0.0 | 1718316.65 | 2498148.68 | True |
| F | G | 313870.67 | 0.1664 | −76045.35 | 703786.68 | False |
| F | H | 2028650.33 | 0.0 | 1638734.32 | 2418566.35 | True |
| G | H | 1714779.67 | 0.0 | 1324863.65 | 2104695.68 | True |

(iv) Gel size changes throughout in-gel Edman degradation for PITC (1:1000 ratio PITC:DMSO) conjugation over multiple rounds (Fig. 5A).

| **Conjugation with PITC to DMSO (1:1000 ratio PITC:DMSO)- Gel size changes in Edman steps** | **Gelation 1, area ($cm^2$)** | **Gelation 2, area ($cm^2$)** | **Gelation 3, area ($cm^2$)** | **Mean (normalized to surface area in 1M Tris pH 9.5) ($cm^2$)** | **Standard deviation ($cm^2$)** |
|---|---|---|---|---|---|
| 1. 3x wash with Tris 1M pH 9.5 | 0.250 | 0.275 | 0.275 | 1.000 | 0.000 |
| 2. 2x DMSO wash | 0.250 | 0.303 | 0.275 | 1.034 | 0.058 |
| 3. PITC conjugation | 0.250 | 0.330 | 0.275 | 1.069 | 0.115 |
| 4. TFA | 0.250 | 0.275 | 0.225 | 0.938 | 0.105 |
| 5. 2x DMSO wash | 0.160 | 0.180 | 0.180 | 0.650 | 0.008 |
| 6. 3x wash with Tris 1M pH 9.5 | 0.250 | 0.275 | 0.275 | 1.000 | 0.000 |
| 7. 2x DMSO wash | 0.225 | 0.250 | 0.248 | 0.903 | 0.005 |
| 8. PITC conjugation | 0.250 | 0.330 | 0.248 | 1.034 | 0.153 |
| 9. TFA | 0.250 | 0.275 | 0.248 | 0.966 | 0.058 |
| 10. DMSO wash | 0.160 | 0.180 | 0.180 | 0.650 | 0.008 |
| 11. 3x wash with Tris 1M pH 9.5 | 0.250 | 0.275 | 0.275 | 1.000 | 0.000 |
| 12. 2x DMSO wash | 0.225 | 0.275 | 0.225 | 0.906 | 0.091 |
| 13. PITC conjugation | 0.225 | 0.250 | 0.225 | 0.875 | 0.050 |
| 14. TFA | 0.225 | 0.250 | 0.225 | 0.875 | 0.050 |
| 15. DMSO wash | 0.160 | 0.180 | 0.160 | 0.625 | 0.038 |
| 16. 3x wash with Tris 1M pH 8 | 0.250 | 0.275 | 0.225 | 0.938 | 0.105 |

**Repeated-measures ANOVA (Fig. 5A)**

| Effect | df | F | p-value |
|---|---|---|---|
| Time point | 1.67, 3.34 | 19.01 | 0.016 |

***Greenhouse-Geisser corrected; n = 3 gels.***

**Dunnett's Test — Each Time Point vs Control (t1) (Fig. 5A)**

| Comparison | Adjusted p | Significance |
|---|---|---|
| t1 vs t2 | 1.0000 | ns |
| t1 vs t3 | 0.9722 | ns |
| t1 vs t4 | 0.9868 | ns |
| t1 vs t5 | 0.0003 | *** |
| t1 vs t6 | 1.0000 | ns |
| t1 vs t7 | 0.8005 | ns |
| t1 vs t8 | 1.0000 | ns |
| t1 vs t9 | 1.0000 | ns |
| t1 vs t10 | 0.0002 | *** |
| t1 vs t11 | 1.0000 | ns |
| t1 vs t12 | 0.8230 | ns |
| t1 vs t13 | 0.5125 | ns |
| t1 vs t14 | 0.5123 | ns |
| t1 vs t15 | 0.0001 | *** |
| t1 vs t16 | 0.9867 | ns |

***Significance: *** p < 0.001, ** p < 0.01, * p < 0.05, ns = not significant.***

**Supplementary Table 7**
Values and analysis for the LC/QToF trypsinization assay from Fig. 6.

(i) Raw data values for the area under the curve (AUC) of the chromatogram of various species for ClickP (1:1000 ratio ClickP:DMSO) conjugation (Fig. 6C) (extracted based on the exact mass, see Methods for more details, and Source Data for raw traces). Three tables for the three separate gelation solutions. The cells highlighted in grey are the ones used to calculate the conversion rate (1-(solvent with ClickP["A15-peptide"] / solvent ["A15-peptide"])). A15-peptide stands for AGGAGGLLGGSRGGK{acr}. Cells with '-' means that the species was not detected in that sample using the automatic extraction.

| | Gelation | DMSO | DMSO with ClickP | DMSO, then TFA | DMSO with ClickP, then TFA | Conversion rate |
|---|---|---|---|---|---|---|
| **A15-peptide** | Gelation 1 | 13,053,858 | 208,055 | 11,854,227 | 214,082 | 98.41 |
| **PITC conjugated A15-peptide** | | - | 2,892,880 | - | - | |
| **A15-peptide with cleaved N-terminal amino acid** | | 86,430 | 3,028,163 | 84,602 | 5,280,782 | |
| **A9-peptide (control)** | | 1,990,306 | 1,513,936 | 2,062,392 | 1,734,325 | |

| | Gelation | DMSO | DMSO with ClickP | DMSO, then TFA | DMSO with ClickP, then TFA | Conjugation Efficiency |
|---|---|---|---|---|---|---|
| **A15-peptide** | Gelation 2 | 11,948,873 | 169,549 | 12,536,770 | 188,635 | 98.58 |
| **PITC conjugated A15-peptide** | | - | 2,317,192 | - | 13,628 | |
| **A15-peptide with cleaved N-terminal amino acid** | | 67,053 | 1,757,956 | 78,724 | 4,698,120 | |
| **A9-peptide (control)** | | 1,512,368 | 2,079,931 | 1,997,604 | 2,253,932 | |

| | Gelation | DMSO | DMSO with ClickP | DMSO, then TFA | DMSO with ClickP, then TFA | Conjugation Efficiency |
|---|---|---|---|---|---|---|
| **A15-peptide** | Gelation 3 | 9,167,841 | 108,656 | 9,047,914 | 197,729 | 98.81 |
| **PITC conjugated A15-peptide** | | - | 1,667,014 | - | 10,077 | |
| **A15-peptide with cleaved N-terminal amino acid** | | 61,582 | 1,218,684 | 63,182 | 5,427,226 | |
| **A9-peptide (control)** | | 1,764,034 | 1,930,392 | 1,859,683 | 1,953,816 | |

(ii) Conversion rate of A15-peptide to ClickP conjugated A15-peptide, and % yield for conjugation with ClickP (1:1000 ratio ClickP:DMSO) (**Fig. 6C**).

| | |
|---|---|
| **Average % yield (peptide with cleaved N-terminal amino acid / non-modified peptide * 100)** | 49.26 |
| **Average conversion rate (%)** | 98.60 |

(iii) Statistics: One-way Analysis of Variance (ANOVA) and Tukey's post-hoc Honestly Significant Difference (HSD) test on the abundance of peptide with cleaved N-terminal amino acid using ClickP (1:1000 ratio ClickP:DMSO) for conjugation (**Fig. 6Ciii**). In Tukey HSD results, conditions are abbreviated, A: DMSO, B: DMSO with ClickP, C: DMSO then TFA, D: DMSO with ClickP then TFA.

| **ANOVA Results** | | | | | |
|---|---|---|---|---|---|
| Test | Comparison | sum_sq | df | F | PR(>F) |
| ANOVA | group | 5.14E+13 | 3.00E+00 | 6.77E+01 | 5.01E-06 |
| ANOVA | Residual | 2.02E+12 | 8.00E+00 | | |

| **Tukey HSD Results** | | | | | | | |
|---|---|---|---|---|---|---|---|
| Test | group1 | group2 | meandiff | p-adj | lower | upper | reject |
| Tukey HSD | A | B | 1.93E+06 | 6.71E-03 | 6.15E+05 | 3.25E+06 | TRUE |
| Tukey HSD | A | C | 3.81E+03 | 1.00E+00 | -1.31E+06 | 1.32E+06 | FALSE |
| Tukey HSD | A | D | 5.06E+06 | 8.19E-06 | 3.75E+06 | 6.38E+06 | TRUE |
| Tukey HSD | B | C | -1.93E+06 | 6.79E-03 | -3.24E+06 | -6.11E+05 | TRUE |
| Tukey HSD | B | D | 3.13E+06 | 2.82E-04 | 1.82E+06 | 4.45E+06 | TRUE |
| Tukey HSD | C | D | 5.06E+06 | 8.24E-06 | 3.75E+06 | 6.38E+06 | TRUE |

**Supplementary Table 8**

Values and analysis for the LC/QToF trypsinization assay from **Supplementary Figures**.

**(A & B)** Raw data values for the area under the curve (AUC) of the chromatogram of various species for results from in-gel Edman degradation with 1:1000 ratio PITC:pyridine (Supplementary Table 8**Ai**), 1:1000 PITC ratio PITC:acetonitrile (Supplementary Table 8**Aii**) (see Supplementary Figure 3**Ai and** 3**Aii** for the plots of these results), 1:9 ratio PITC to acetonitrile) (Supplementary Table 8**Bi**), 1:9 ratio PITC to 1:1 pyridine and water (Supplementary Table 8**Bii**), 1:9 ratio PITC:0.1 M $NaHCO_3$ pH 8.5 (Supplementary Table 8**Biii**) (see Supplementary Figure 3**Bi and** 3**Bii and** 3**Biii** for the plots of these results) (extracted based on the exact mass, see Methods for more details, and Source Data for raw traces). One table is produced for the single gelation solution for each solvent. The cells highlighted in grey are the ones used to calculate the % yield (solvent with PITC and then TFA["A15-peptide"] / solvent and then TFA["A15-peptide"]). A15-peptide stands for AGGAGGLLGGSRGGK{acr}. Cells with '-' means that the species was not detected in that sample using the automatic extraction.

| **Table 8 A i) Conjugation with using PITC to pyridine (1:1000 ratio PITC:pyridine)** | **Gelation** | **pyridine** | **pyridine with PITC** | **pyridine, then TFA** | **pyridine with PITC, then TFA** | **% yield (peptide with cleaved N-terminal amino acid / non-modified peptide * 100)** |
|---|---|---|---|---|---|---|
| **A15-peptide** | Gelation 1 | 15,197,533 | 15,865,081 | 17,420,478 | 14,652,655 | 0.57 |
| **PITC conjugated A15-peptide** | | - | - | - | - | |
| **A15-peptide with cleaved N-terminal amino acid** | | 90,340 | 85,353 | 102,755 | 98,890 | |
| **A9-peptide (control)** | | 3,669,196 | 3,818,575 | 3,538,687 | 3,724,464 | |

| **Table 8 A ii) Conjugation with using PITC to acetonitrile (1:1000 ratio PITC:acetonitrile)** | **Gelation** | **Acetonitrile** | **acetonitrile with PITC** | **acetonitrile, then TFA** | **acetonitrile with PITC, then TFA** | **% yield (peptide with cleaved N-terminal amino acid / non-modified peptide * 100)** |
|---|---|---|---|---|---|---|
| **A15-peptide** | Gelation 1 | 12,415,553 | 16,650,344 | 14,832,564 | 12626395 | 0.46 |
| **PITC conjugated A15-peptide** | | - | - | - | - | |
| **A15-peptide with cleaved N-terminal amino acid** | | 60,749 | 99,444 | 69,906 | 67,941 | |
| **A9-peptide (control)** | | 2,350,910 | 3,277,849 | 3,766,113 | 3,979,891 | |

| **Table 8 B i) Conjugation using PITC to acetonitrile (1:9 ratio PITC:acetonitrile)** | **Gelation** | **Acetonitrile** | **acetonitrile with PITC** | **acetonitrile, then TFA** | **acetonitrile with PITC, then TFA** | **% yield (peptide with cleaved N-terminal amino acid / non-modified peptide * 100)** |
|---|---|---|---|---|---|---|
| **A15-peptide** | Gelation 1 | 7,493,808 | 8,463,312 | 5,112,020 | 6,006,358 | 0.27 |

| | | | | | |
|---|---|---|---|---|---|
| **PITC conjugated A15-peptide** | | - | - | - | - |
| **A15-peptide with cleaved N-terminal amino acid** | | 21,128 | 21,568 | 11,002 | 13,558 |
| **A9-peptide (control)** | | 2,150,273 | 1,945,023 | 1,902,023 | 2,216,702 |

| **Table 8 B ii) Conjugation using PITC in 1:1 pyridine to water (1:9 ratio PITC to 1:1 pyridine to water)** | **Gelation** | **1:1 pyridine to water** | **1:1 pyridine to water with PITC** | **1:1 pyridine to water, then TFA** | **1:1 pyridine to water with PITC, then TFA** | **% yield (peptide with cleaved N-terminal amino acid / non-modified peptide * 100)** |
|---|---|---|---|---|---|---|
| **A15-peptide** | Gelation 1 | 7,681,288 | 565,768 | 6,802,453 | 1,242,845 | 59.34 |
| **PITC conjugated A15-peptide** | | 5,431 | 2,594,118 | - | - | |
| **A15-peptide with cleaved N-terminal amino acid** | | 12,188 | 2,144,764 | 10,933 | 4,036,719 | |
| **A9-peptide (control)** | | 1,921,927 | 1,836,294 | 2,052,361 | 2,052,891 | |

| **Table 8 B iii) Conjugation using PITC 0.1 M $NaHCO_3$ pH 8.5 (1:9 ratio PITC:0.1 M $NaHCO_3$ pH 8.5)** | **Gelation** | **0.1 M $NaHCO_3$ pH 8.5** | **0.1 M $NaHCO_3$ pH 8.5 with PITC** | **0.1 M $NaHCO_3$ pH 8.5, then TFA** | **0.1 M $NaHCO_3$ pH 8.5 with PITC, then TFA** | **% yield (peptide with cleaved N-terminal amino acid / non-modified peptide * 100)** |
|---|---|---|---|---|---|---|
| **A15-peptide** | Gelation 1 | 6,429,218 | 4,290,786 | 6,016,772 | 4,163,519 | 10.55 |
| **PITC conjugated A15-peptide** | | - | 90,914 | 7,154 | 5,535 | |
| **A15-peptide with cleaved N-terminal amino acid** | | 24,410 | 1,089,746 | 22,675 | 634,809 | |
| **A9-peptide (control)** | | 1,940,911 | 2,125,041 | 1,879,287 | 2,120,299 | |

**(C)** Raw data values for the area under the curve (AUC) of the chromatogram of various species from in-gel Edman degradation with different ratios of PITC to DMSO (1:100, 1:1,000, 1:10,0000 ratios PITC:DMSO) (see **Supplementary Figure 4** for the plots of these results) (extracted based on the exact mass, see **Methods** for more details, and **Source Data** for raw traces). Three tables for the three separate ratios of PITC to DMSO (1:100, 1:1,000, 1:10,000 PITC:DMSO). The cells highlighted in grey are the ones used to calculate the conversion rate (1-(solvent with PITC[“A15-peptide”] / solvent [“A15-peptide”])). A15-peptide stands for AGGAGGLLGGSRGGK{acr}. Cells with ‘-’ means that the species was not detected in that sample using the automatic extraction.

| **Conjugation with PITC to DMSO (1:100 ratio PITC:DMSO)** | **Gelation** | **DMSO** | **DMSO with PITC** | **DMSO, then TFA** | **DMSO with PITC, then TFA** | **Conversion rate** |
|---|---|---|---|---|---|---|
| **A15-peptide** | Gelation 1 | 9,052,218 | 47,171 | 7,989,893 | 42,236 | 99.48 |

| | | | | | |
|---|---|---|---|---|---|
| **PITC conjugated A15-peptide** | | - | 392,164 | - | - |
| **A15-peptide with cleaved N-terminal amino acid** | | 48,959 | 5,573,096 | 40,151 | 5,936,781 |
| **A9-peptide (control)** | | 4,742,905 | 4,541,611 | 4,878,650 | 5,148,177 |

| **Conjugation with 1:1000 PITC to DMSO (1:1000 ratio PITC:DMSO)** | **Gelation** | **DMSO** | **DMSO with PITC** | **DMSO, then TFA** | **DMSO with PITC, then TFA** | **Conversion rate** |
|---|---|---|---|---|---|---|
| **A15-peptide** | Gelation 1 | 9,382,061 | 64,147 | 7,360,985 | 77,077 | 99.32 |
| **PITC conjugated A15-peptide** | | - | 438,161 | - | - | |
| **A15-peptide with cleaved N-terminal amino acid** | | 48,913 | 6,422,687 | 35,693 | 6,522,567 | |
| **A9-peptide (control)** | | 4,920,745 | 4,448,116 | 5,565,431 | 4,697,838 | |

| **Conjugation with 1:10,000 PITC to DMSO (1:10,000 ratio PITC:DMSO)** | **Gelation** | **DMSO** | **DMSO with PITC** | **DMSO, then TFA** | **DMSO with PITC, then TFA** | **Conversion rate** |
|---|---|---|---|---|---|---|
| **A15-peptide** | Gelation 1 | 7,542,918 | 959,140 | 8,492,660 | 1,109,025 | 87.28 |
| **PITC conjugated A15-peptide** | | - | 297842 | - | - | |
| **A15-peptide with cleaved N-terminal amino acid** | | 37,026 | 4,876,060 | 44,617 | 4,574,983 | |
| **A9-peptide (control)** | | 4,616,757 | 4,847,843 | 5,146,408 | 4,922,362 | |

**(D)** Raw data values for Supplementary Figure 5 representing the area under the curve (AUC) of the chromatogram of various peptide fragments derived from the peptide denoted A15 (AGGAGGLLGGSRGGK{acr}) expected after trypsinization from the gel (denoted "I_product"). Two products are shown in different tables: the first, "non-modified A15-peptide" (AGGAGGLLGGSR) and "A15-peptide with cleaved N-terminal amino acid" (GGAGGLLGGSR). The A9-peptide is denoted "standard" (AGGAGK{acr}GLR) and was used in all samples as the standard with a constant "C_standard" of 5 µM. TheAUC of the extracted ion chromatogram for the A9-peptide is recorded as "I_standard" (extracted based on the exact mass, see Analysis of LC/QToF data for details on method of extraction). The AUC of non-modified A15-peptide and A15-peptide with cleaved N-terminal amino acid were measured from 5 µM to 60 µM (in 5 µM increments; denoted: "C_product") with A9-peptide concentration always at 5 µM.

| **Replicate 1** | **5 µM peptide "C_product"** | **10 µM peptide "C_product"** | **20 µM peptide "C_product"** | **30 µM peptide "C_product"** | **40 µM peptide "C_product"** | **50 µM peptide "C_product"** | **60 µM peptide "C_product"** |
|---|---|---|---|---|---|---|---|

| | | | | | | | |
|---|---|---|---|---|---|---|---|
| **Non-modified A15-peptide "I_product"** | 3,799,448 | 6,085,960 | 11,319,222 | 14,253,684 | 19,809,211 | 23,947,646 | 25,596,532 |
| **A15-peptide with cleaved N-terminal amino acid "I_product"** | 2,525,298 | 5,020,612 | 10,220,593 | 13,724,111 | 17,279,065 | 19,271,410 | 23,143,493 |
| **A-9 control peptide "I_standard"** | 1,975,582 | 1,551,193 | 2,617,067 | 2,297,308 | 2,646,120 | 2,173,907 | 2,440,661 |
| **Non-modified A15-peptide "I_product/I_standard"** | 1.92 | 3.92 | 4.33 | 6.20 | 7.49 | 11.02 | 10.49 |
| **A15-peptide with cleaved N-terminal amino acid "I_product/I_standard"** | 1.28 | 3.24 | 3.91 | 5.97 | 6.53 | 8.86 | 9.48 |

| **Replicate 2** | **5 µM peptide "C_product"** | **10 µM peptide "C_product"** | **20 µM peptide "C_product"** | **30 µM peptide "C_product"** | **40 µM peptide "C_product"** | **50 µM peptide "C_product"** | **60 µM peptide "C_product"** |
|---|---|---|---|---|---|---|---|
| **Non-modified A15-peptide "I_product"** | 2,744,052 | 4,939,161 | 8,944,190 | 12,276,463 | 15,692,870 | 17,966,806 | 20,359,400 |
| **A15-peptide with cleaved N-terminal amino acid "I_product"** | 3,446,088 | 6,532,592 | 11,438,018 | 16,089,772 | 20,973,147 | 23,719,649 | 27,786,734 |
| **A-9 control peptide "I_standard"** | 2,302,553 | 2,156,882 | 2,052,737 | 2,045,822 | 2,041,539 | 2,341,466 | 2,028,886 |
| **Non-modified A15-peptide "I_product/I_standard"** | 1.19 | 2.29 | 4.36 | 6.00 | 7.69 | 7.67 | 10.03 |
| **A15-peptide with cleaved N-terminal amino acid "I_product/I_standard"** | 1.50 | 3.03 | 5.57 | 7.86 | 10.27 | 10.13 | 13.70 |

| **Replicate 3** | **5 µM peptide "C_product"** | **10 µM peptide "C_product"** | **20 µM peptide "C_product"** | **30 µM peptide "C_product"** | **40 µM peptide "C_product"** | **50 µM peptide "C_product"** | **60 µM peptide "C_product"** |
|---|---|---|---|---|---|---|---|
| **Non-modified A15-peptide "I_product"** | 4,068,182 | 4,683,734 | 8,511,458 | 11,651,232 | 15,479,768 | 17,802,635 | 19,996,525 |
| **A15-peptide with cleaved N-terminal amino acid "I_product"** | 3,455,652 | 5,597,608 | 9,316,407 | 13,460,858 | 17,144,374 | 20,441,281 | 23,081,247 |
| **A-9 control peptide "I_standard"** | 2,249,002 | 2,450,888 | 2,032,303 | 2,595,972 | 2,306,414 | 2,400,407 | 2,264,781 |
| **Non-modified A15-peptide "I_product/I_standard"** | 1.81 | 1.91 | 4.19 | 4.49 | 6.71 | 7.42 | 8.83 |

| | | | | | | | |
|---|---|---|---|---|---|---|---|
| **A15-peptide with cleaved N-terminal amino acid "I_product/I_standard"** | 1.54 | 2.28 | 4.58 | 5.19 | 7.43 | 8.52 | 10.19 |

| | **5 µM peptide "C_product"** | **10 µM peptide "C_product"** | **20 µM peptide "C_product"** | **30 µM peptide "C_product"** | **40 µM peptide "C_product"** | **50 µM peptide "C_product"** | **60 µM peptide "C_product"** |
|---|---|---|---|---|---|---|---|
| **A-9 control peptide "C_standard"** | 5 | 5 | 5 | 5 | 5 | 5 | 5 |
| **Mean "C_product/C_standard"** | 1 | 2 | 4 | 6 | 8 | 10 | 12 |

| **Concentration of peptide (µM) "C_product"** | **5 µM peptide "C_product"** | **10 µM peptide "C_product"** | **20 µM peptide "C_product"** | **30 µM peptide "C_product"** | **40 µM peptide "C_product"** | **50 µM peptide "C_product"** | **60 µM peptide "C_product"** |
|---|---|---|---|---|---|---|---|
| **Mean non-modified peptide "I_product/I_standard"** | 1.63 | 2.55 | 4.29 | 5.50 | 7.29 | 8.63 | 9.79 |
| **Standard deviation non-modified peptide** | 0.39 | 1.07 | 0.09 | 0.94 | 0.51 | 2.01 | 0.86 |
| **Mean peptide with cleaved N-terminal amino acid "I_product/I_standard"** | 1.44 | 2.78 | 4.62 | 6.241 | 7.92 | 9.17 | 10.99 |
| **Standard deviation peptide with cleaved N-terminal amino acid** | 0.14 | 0.50 | 0.84 | 1.38 | 1.95 | 0.85 | 2.26 |
| **T-statistic (Welch's t-test, n=3) comparing the two means of "I_product/I_standard" for non-modified A15-peptide and A15-peptide with cleaved N-terminal amino acid** | 0.847 | -0.208 | -0.816 | -0.808 | -0.672 | -0.372 | -0.961 |
| **P-value (Welch's t-test, n=3) comparing the two means of "I_product/I_standard" for non-modified A15-peptide and peptide with A15-peptide with cleaved N-terminal amino acid** | 0.470 | 0.849 | 0.499 | 0.470 | 0.563 | 0.737 | 0.418 |

#### Supplementary Table 9

**a) ClpS2 St-V1 assay with anti-HA antibody 488 comparing different N-terminal amino acids (phenylalanine, alanine, tryptophan, tyrosine)**

(i) Raw average fluorescence intensity values for each gel, average, and standard deviation, over the replicates for Fig. 4**D**.

| **Condition name** | **Average intensity ExMre gel 1** | **Average intensity ExMre gel 2** | **Average intensity ExMre gel 3** | **Average intensity over replicates** | **Standard deviation over replicates** |
|---|---|---|---|---|---|
| Phenylalanine (F) | 13,731 | 7,847 | 9,084 | **10,221** | 3102.31725 |
| Alanine (A) | 308 | 277 | 255 | **280** | 26.6270539 |
| Tryptophan (W) | 561 | 347 | 1084 | **664** | 379.142453 |
| Tyrosine (Y) | 1,741 | 873 | 1,179 | **1,264** | 440.2469 |

(ii) Fold difference between the average intensity read-out for Fig. 4**D** based on values documented in (i).

| **Fold change** | |
|---|---|
| F/A | 37 |
| F/W | 15 |
| F/Y | 8 |

(iii) One-way Analysis of Variance (ANOVA) and Tukey's post-hoc Honestly Significant Difference (HSD) test (FWER = 0.05) on the abundance of average fluorescence intensity in the gel (Fig. 4**A-D**). In Tukey HSD results, conditions are abbreviated, A: phenylalanine, B: alanine, C: tryptophan, D: tyrosine.

**ANOVA Results**

| | **Sum of Squares** | **df** | **F** | **p-value** |
|---|---|---|---|---|
| Group | 2.039e+08 | 3.0 | 27.2859 | 0.000149 |
| Residual | 1.993e+07 | 8.0 | — | — |

**Tukey HSD Results**

| **Group 1** | **Group 2** | **Mean Diff** | **p-adj** | **Lower** | **Upper** | **Reject** |
|---|---|---|---|---|---|---|
| A | B | −9940.67 | 0.0003 | −14067.16 | −5814.18 | True |
| A | C | −9556.67 | 0.0003 | −13683.16 | −5430.18 | True |
| A | D | −8956.33 | 0.0005 | −13082.82 | −4829.84 | True |

| | | | | | | |
|---|---|---|---|---|---|---|
| B | C | 384.0 | 0.9901 | −3742.49 | 4510.49 | False |
| B | D | 984.33 | 0.8684 | −3142.16 | 5110.82 | False |
| C | D | 600.33 | 0.9645 | −3526.16 | 4726.82 | False |

**b) ClpS2 St-V1 assay with anti-HA antibody 488 on $F_1$ peptide combined with in-gel Edman degradation.**

(i) Raw average fluorescence intensity values for each gel, average, and standard deviation, over the replicates for Fig. 4**Fiii**.

| **Condition name** | **Average intensity ExMre gel 1** | **Average intensity ExMre gel 2** | **Average intensity over replicates** | **Standard deviation over replicates** |
|---|---|---|---|---|
| (1) DMSO | 5,381 | 7,731 | **6,556** | 1,662 |
| (2) PITC:DMSO | 474 | 411 | **443** | 45 |
| (3) DMSO+TFA | 6,290 | 8,694 | **7,492** | 1,700 |
| (4) PITC:DMSO+TFA | 400 | 640 | **520** | 170 |
| (5) PITC:DMSO+TFA+trypsin | 250 | 606 | **428** | 252 |

(ii) Fold difference between the average intensity read-out for Fig. 4**Fiii** based on values documented in (i) (e.g., condition (1) is DMSO and (3) is TFA, as number in (i))

| **Fold change in average fluorescence intensity read-out** | Ratio |
|---|---|
| (1)/(2) | 14.82 |
| (1)/(3) | 0.88 |
| (1)/(4) | 12.61 |
| (1)/(5) | 15.32 |

| **Fold change in average fluorescence intensity read-out** | Ratio |
|---|---|
| (3)/(1) | 1.14 |
| (3)/(2) | 16.93 |
| (3)/(4) | 14.41 |
| (3)/(5) | 17.50 |

(iii) One-way Analysis of Variance (ANOVA) and Tukey's post-hoc Honestly Significant Difference

(HSD) test (FWER = 0.05) on the abundance of average fluorescence intensity in the gel (Fig. 4F). In Tukey HSD results, conditions are abbreviated, A: DMSO, B: DMSO:PITC, C: DMSO+TFA, D: DMSO:PITC+TFA, E: DMSO:PITC+TFA+trypsin.

**ANOVA Results (Fig. 4F)**

| | Sum of Squares | df | F | p-value |
|---|---|---|---|---|
| Group | 104182267.6 | 4.0 | 22.6680 | 0.002104 |
| Residual | 5745010.5 | 5.0 | — | — |

**Tukey HSD Results (Fig. 4F)**

| Group 1 | Group 2 | Mean Diff | p-adj | Lower | Upper | Reject |
|---|---|---|---|---|---|---|
| A | B | −6113.5 | 0.0121 | −10413.49 | −1813.51 | True |
| A | C | 936.0 | 0.8955 | −3363.99 | 5235.99 | False |
| A | D | −6036.0 | 0.0127 | −10335.99 | −1736.01 | True |
| A | E | −6128.0 | 0.0119 | −10427.99 | −1828.01 | True |
| B | C | 7049.5 | 0.0065 | 2749.51 | 11349.49 | True |
| B | D | 77.5 | 1.0 | −4222.49 | 4377.49 | False |
| B | E | −14.5 | 1.0 | −4314.49 | 4285.49 | False |
| C | D | −6972.0 | 0.0068 | −11271.99 | −2672.01 | True |
| C | E | −7064.0 | 0.0064 | −11363.99 | −2764.01 | True |
| D | E | −92.0 | 1.0 | −4391.99 | 4207.99 | False |

c) **ClpS2 St-V1 assay with anti-HA antibody 488 on $G_1F_2$ peptide combined with in-gel Edman degradation.**

(i) Raw average fluorescence intensity values for each gel, average, and standard deviation, over the replicates for Fig. 4**Giii**.

| **Condition name** | **Average intensity ExMre gel 1** | **Average intensity ExMre gel 2** | **Average intensity ExMre gel 3** | **Average intensity over replicates** | **Standard deviation over replicates** |
|---|---|---|---|---|---|
| (1) DMSO | 658 | 418 | 339 | **472** | 166 |
| (2) PITC:DMSO | 485 | 526 | 491 | **501** | 22 |
| (3) DMSO+TFA | 499 | 589 | 450 | **513** | 71 |
| (4) PITC:DMSO+TFA | 2,463 | 3,802 | 3,235 | **3167** | 672 |
| (5) PITC:DMSO+TFA +trypsin | 266 | 357 | 484 | **369** | 109 |

(ii) Fold difference between the average intensity read-out for Fig. 4**Giii** based on values documented in (i) (e.g., condition (4) is PITC:DMSO+TFA, as number in (i))

| **Fold change in average fluorescence intensity read-out** | Ratio |
|---|---|
| (4)/(1) | 6.71 |
| (4)/(2) | 6.32 |
| (4)/(3) | 6.18 |
| (4)/(5) | 8.58 |

(iii) One-way Analysis of Variance (ANOVA) and Tukey's post-hoc Honestly Significant Difference (HSD) test (FWER = 0.05) on the abundance of average fluorescence intensity in the gel (Fig. 4**G**). In Tukey HSD results, conditions are abbreviated, A: DMSO, B: DMSO:PITC, C: DMSO+TFA, D: DMSO:PITC+TFA, E: DMSO:PITC+TFA+trypsin.

**ANOVA Results (Fig. 4G)**

| | **Sum of Squares** | **df** | **F** | **p-value** |
|---|---|---|---|---|
| Group | 1.758e+07 | 4.0 | 44.2232 | 0.000003 |
| Residual | 9.936e+05 | 10.0 | — | — |

**Tukey HSD Results (Fig. 4G)**

| **Group 1** | **Group 2** | **Mean Diff** | **p-adj** | **Lower** | **Upper** | **Reject** |
|---|---|---|---|---|---|---|
| A | B | 29.0 | 1.0 | −818.02 | 876.02 | False |
| A | C | 41.0 | 0.9998 | −806.02 | 888.02 | False |
| A | D | 2695.0 | 0.0 | 1847.98 | 3542.02 | True |
| A | E | −102.67 | 0.9938 | −949.68 | 744.35 | False |
| B | C | 12.0 | 1.0 | −835.02 | 859.02 | False |
| B | D | 2666.0 | 0.0 | 1818.98 | 3513.02 | True |
| B | E | −131.67 | 0.9842 | −978.68 | 715.35 | False |
| C | D | 2654.0 | 0.0 | 1806.98 | 3501.02 | True |
| C | E | −143.67 | 0.9783 | −990.68 | 703.35 | False |
| D | E | −2797.67 | 0.0 | −3644.68 | −1950.65 | True |

**d) tvClpS2 Q31H assay with anti 6xHis antibody 488 comparing different N-terminal amino acids (phenylalanine, alanine, tryptophan, tyrosine, leucine, and no peptide gel)**

(i) Raw average fluorescence intensity values for each gel, average, and standard deviation, over the replicates for Fig. 4**H**.

| Condition name | Average intensity ExMre gel 1 | Average intensity ExMre gel 2 | Average intensity ExMre gel 3 | Average intensity over replicates | Standard deviation over replicates |
|---|---|---|---|---|---|
| Phenylalanine (F) | 5023.59508 | 3531.01217 | 2955.22601 | **3836.61109** | 1,067.51144 |
| Alanine (A) | 830.47643 | 953.50251 | 1238.16950 | **1007.38281** | 209.118923 |
| Tryptophan (W) | 804.34104 | 926.70173 | 1036.80741 | **922.61673** | 116.28701 |
| Tyrosine (Y) | 1220.48740 | 728.89890 | 1013.89616 | **987.76082** | 246.83416 |
| Leucine (L) | 6004.60776 | 3026.41741 | 3096.49085 | **4042.50534** | 1,699.59171 |
| No peptide | 966.76140 | 804.61619 | 849.28700 | **873.55486** | 83.75240 |

(ii) One-way Analysis of Variance (ANOVA) and Tukey's post-hoc Honestly Significant Difference (HSD) test (FWER = 0.05) on the abundance of average fluorescence intensity in the gel (**Fig. 4H**). In Tukey HSD results, conditions are abbreviated, A: phenylalanine, B: alanine, C: tryptophan, D: tyrosine, E: leucine, F: no peptide.

**ANOVA Results (Fig. 4H)**

| | Sum of Squares | df | F | p-value |
|---|---|---|---|---|
| Group | 3.590e+07 | 5.0 | 10.3720 | 0.000496 |
| Residual | 8.307e+06 | 12.0 | — | — |

**Tukey HSD Results (Fig. 4H)**

| Group 1 | Group 2 | Mean Diff | p-adj | Lower | Upper | Reject |
|---|---|---|---|---|---|---|
| A | B | −2829.23 | 0.0128 | −5111.04 | −547.42 | True |
| A | C | −2913.99 | 0.0104 | −5195.81 | −632.18 | True |
| A | D | −2848.85 | 0.0122 | −5130.66 | −567.04 | True |
| A | E | 205.89 | 0.9996 | −2075.92 | 2487.71 | False |
| A | F | −2963.06 | 0.0092 | −5244.87 | −681.24 | True |
| B | C | −84.77 | 1.0 | −2366.58 | 2197.05 | False |
| B | D | −19.62 | 1.0 | −2301.43 | 2262.19 | False |
| B | E | 3035.12 | 0.0077 | 753.31 | 5316.93 | True |
| B | F | −133.83 | 0.9999 | −2415.64 | 2147.98 | False |
| C | D | 65.14 | 1.0 | −2216.67 | 2346.96 | False |
| C | E | 3119.89 | 0.0063 | 838.08 | 5401.70 | True |
| C | F | −49.06 | 1.0 | −2330.87 | 2232.75 | False |
| D | E | 3054.74 | 0.0074 | 772.93 | 5336.56 | True |

| | | | | | | |
|---|---|---|---|---|---|---|
| D | F | −114.21 | 1.0 | −2396.02 | 2167.61 | False |
| E | F | −3168.95 | 0.0056 | −5450.76 | −887.14 | True |

**e) SuTEx-azide and DBCO-AF488 read-out on different N-terminal amino acids (phenylalanine, alanine, tryptophan, tyrosine, leucine, and no peptide gel)**

(i) Raw average fluorescence intensity values for each gel, average, and standard deviation, over the replicates for Fig. 4**Ii-iv**.

| **Condition name** | **Average intensity ExMre gel 1** | **Average intensity ExMre gel 2** | **Average intensity ExMre gel 3** | **Average intensity over replicates** | **Standard deviation over replicates** |
|---|---|---|---|---|---|
| Phenylalanine (F) | 2872.36801 | 3087.11648 | 3168.21230 | **3042.56559** | 152.87102 |
| Alanine (A) | 1929.72563 | 2056.63766 | 1964.85827 | **1983.74052** | 65.52916 |
| Tryptophan (W) | 3324.51890 | 2715.20300 | 3197.14845 | **3078.95678** | 321.39293 |
| Tyrosine (Y) | 9498.68800 | 9104.19872 | 9762.68304 | **9455.18992** | 331.39020 |
| No peptide | 1755.59881 | 1548.85748 | 1686.20616 | **1663.55415** | 105.21563 |

(ii) One-way Analysis of Variance (ANOVA) and Tukey's post-hoc Honestly Significant Difference (HSD) test (FWER = 0.05) on the abundance of average fluorescence intensity in the gel (Fig. 4**Ii-iv**). In Tukey HSD results, conditions are abbreviated, A: phenylalanine, B: alanine, C: tryptophan, D: tyrosine, E: no peptide.

**ANOVA Results (Fig. 4Ii-iv)**

| | **Sum of Squares** | **df** | **F** | **p-value** |
|---|---|---|---|---|
| Group | 1.228e+08 | 4.0 | 609.4172 | 6.806e-12 |
| Residual | 5.037e+05 | 10.0 | — | — |

**Tukey HSD Results (Fig. 4Ii-iv)**

| **Group 1** | **Group 2** | **Mean Diff** | **p-adj** | **Lower** | **Upper** | **Reject** |
|---|---|---|---|---|---|---|
| A | B | −1058.83 | 0.0013 | −1661.91 | −455.74 | True |
| A | C | 36.39 | 0.9996 | −566.69 | 639.47 | False |
| A | D | 6412.62 | 0.0 | 5809.54 | 7015.71 | True |
| A | E | −1379.01 | 0.0002 | −1982.09 | −775.93 | True |
| B | C | 1095.22 | 0.001 | 492.13 | 1698.30 | True |
| B | D | 7471.45 | 0.0 | 6868.37 | 8074.53 | True |
| B | E | −320.19 | 0.4505 | −923.27 | 282.90 | False |
| C | D | 6376.23 | 0.0 | 5773.15 | 6979.32 | True |

| | | | | | | |
|---|---|---|---|---|---|---|
| C | E | −1415.40 | 0.0001 | −2018.48 | −812.32 | True |
| D | E | −7791.64 | 0.0 | −8394.72 | −7188.55 | True |

**f) SuTEx-azide and DBCO-AF488 read-out on in-gel Edman degradation assay on tyrosine N-terminal amino acid peptide ($Y_1$ peptide)**

(i) Raw average fluorescence intensity values for each gel, average, and standard deviation, over the replicates for Fig. 4**Iv-viii**.

| **Condition name** | **Average intensity ExMre gel 1** | **Average intensity ExMre gel 2** | **Average intensity ExMre gel 3** | **Average intensity over replicates** | **Standard deviation over replicates** |
|---|---|---|---|---|---|
| (1) DMSO | 10755.38313 | 7406.64502 | 7933.69758 | **8698.57524** | 1800.63603 |
| (2) PITC:DMSO | 10229.71433 | 8950.04838 | 8123.95284 | **9101.23852** | 1060.99091 |
| (3) DMSO+TFA | 9624.55200 | 7020.80593 | 10008.34677 | **8884.56824** | 1625.43288 |
| (4) PITC:DMSO+TFA | 5269.54343 | 3248.57912 | 4345.90605 | **4288.00954** | 1011.72536 |
| (5) PITC:DMSO+TFA +trypsin | 4942.91732 | 3326.13486 | 2385.22161 | **3551.42460** | 1293.64540 |

(ii) One-way Analysis of Variance (ANOVA) and Tukey's post-hoc Honestly Significant Difference (HSD) test (FWER = 0.05) on the abundance of average fluorescence intensity in the gel (Fig. 4**Iv-viii**). In Tukey HSD results, conditions are abbreviated, A: DMSO, B: DMSO:PITC, C: DMSO+TFA, D: DMSO:PITC+TFA, E: DMSO:PITC+TFA+trypsin.

**ANOVA Results (Fig. 4Iv-viii)**

| | **Sum of Squares** | **df** | **F** | **p-value** |
|---|---|---|---|---|
| Group | 9.016e+07 | 4.0 | 11.6103 | 0.000893 |
| Residual | 1.941e+07 | 10.0 | — | — |

**Tukey HSD (Fig. 4Iv-viii)**

| **Group 1** | **Group 2** | **Mean Diff** | **p-adj** | **Lower** | **Upper** | **Reject** |
|---|---|---|---|---|---|---|
| A | B | 402.66 | 0.9961 | −3341.49 | 4146.82 | False |
| A | C | 185.99 | 0.9998 | −3558.16 | 3930.15 | False |
| A | D | −4410.57 | 0.0202 | −8154.72 | −666.41 | True |
| A | E | −5147.15 | 0.0076 | −8891.30 | −1403.00 | True |
| B | C | −216.67 | 0.9997 | −3960.82 | 3527.48 | False |
| B | D | −4813.23 | 0.0118 | −8557.38 | −1069.08 | True |
| B | E | −5549.81 | 0.0045 | −9293.97 | −1805.66 | True |

| | | | | | | |
|---|---|---|---|---|---|---|
| C | D | −4596.56 | 0.0157 | −8340.71 | −852.41 | True |
| C | E | −5333.14 | 0.006 | −9077.30 | −1588.99 | True |
| D | E | −736.58 | 0.9633 | −4480.74 | 3007.57 | False |

**g) Rudimentary multiplexing: SuTEx-azide, and DBCO-AF488 read-out on in-gel Edman degradation assay on $F_1Y_2L_3$ peptide; ClpS2 St-V1 with anti-HA antibody 488 read-out on $F_1Y_2L_3$ peptide combined with in-gel Edman degradation**

(i) Raw average fluorescence intensity values after SuTEx-azide and DBCO-AF488 staining of $F_1Y_2L_3$ peptide throughout in-gel Edman degradation conditions for each gel, average, and standard deviation, over the replicates for Fig. 4**Ji-iv**.

| Condition name | Average intensity ExMre gel 1 | Average intensity ExMre gel 2 | Average intensity ExMre gel 3 | Average intensity over replicates | Standard deviation over replicates |
|---|---|---|---|---|---|
| (1) DMSO | 17829.16788 | 15075.16257 | 16702.70451 | **16535.67832** | 1384.57922 |
| (2) PITC:DMSO | 15396.19454 | 14681.92096 | 15012.12870 | **15030.08140** | 357.47505 |
| (3) DMSO+TFA | 11798.65104 | 14907.26169 | 16702.26466 | **14469.39247** | 2480.95823 |
| (4) PITC:DMSO+TFA | 15971.12952 | 14460.00021 | 19864.10801 | **16765.07925** | 2788.16470 |
| (5) PITC:DMSO+TFA +trypsin | 3771.42260 | 3661.90766 | 4803.57050 | **4078.96692** | 629.90963 |

(ii) One-way Analysis of Variance (ANOVA) and Tukey's post-hoc Honestly Significant Difference (HSD) test (FWER = 0.05) on the abundance of average fluorescence intensity in the gel (Fig. 4**Ji-iv**). In Tukey HSD results, conditions are abbreviated, A: DMSO, B: DMSO:PITC, C: DMSO+TFA, D: DMSO:PITC+TFA, E: DMSO:PITC+TFA+trypsin.

**ANOVA Results (Fig. 4Ji-iv)**

| | Sum of Squares | df | F | p-value |
|---|---|---|---|---|
| Group | 3.355e+08 | 4.0 | 25.6180 | 0.000031 |
| Residual | 3.274e+07 | 10.0 | — | — |

**Tukey HSD Results (Fig. 4Ji-iv)**

| Group 1 | Group 2 | Mean Diff | p-adj | Lower | Upper | Reject |
|---|---|---|---|---|---|---|
| A | B | −1505.60 | 0.8411 | −6367.89 | 3356.70 | False |
| A | C | −2066.29 | 0.6419 | −6928.58 | 2796.01 | False |
| A | D | 229.40 | 0.9998 | −4632.89 | 5091.69 | False |
| A | E | −12456.71 | 0.0001 | −17319.00 | −7594.42 | True |
| B | C | −560.69 | 0.9949 | −5422.98 | 4301.60 | False |
| B | D | 1734.998 | 0.7652 | −3127.29 | 6597.29 | False |

| | | | | | | |
|---|---|---|---|---|---|---|
| B | E | −10951.11 | 0.0002 | −15813.41 | −6088.82 | True |
| C | D | 2295.69 | 0.5544 | −2566.61 | 7157.98 | False |
| C | E | −10390.43 | 0.0003 | −15252.72 | −5528.13 | True |
| D | E | −12686.11 | 0.0 | −17548.40 | −7823.82 | True |

(iii) Raw **maximum** fluorescence intensity values after ClpS2 St-V1 with anti-HA staining of $F_1Y_2L_3$ peptide throughout in-gel Edman degradation conditions for each gel, average, and standard deviation, over the replicates for Fig. 4**Jv-viii**.

| Condition name | Average intensity ExMre gel 1 | Average intensity ExMre gel 2 | Average intensity ExMre gel 3 | Average intensity over replicates | Standard deviation over replicates |
|---|---|---|---|---|---|
| (1) DMSO | 25581.17705 | 13869.68565 | 17844.45916 | **19098.44062** | 5955.59483 |
| (2) PITC:DMSO | 2254.36402 | 2293.97367 | 7463.28226 | **4003.87332** | 2996.00148 |
| (3) DMSO+TFA | 21829.07342 | 31254.96273 | 20085.30109 | **24389.77908** | 6009.01368 |
| (4) PITC:DMSO+TFA | 7849.21399 | 9553.45785 | 8717.72877 | **8706.80020** | 852.17449 |
| (5) PITC:DMSO+TFA +trypsin | 2593.58343 | 4042.03941 | 1369.70073 | **2668.44119** | 1337.74110 |

(iv) One-way Analysis of Variance (ANOVA) and Tukey's post-hoc Honestly Significant Difference (HSD) test (FWER = 0.05) on the abundance of **maximum** fluorescence intensity in the gel (Fig. 4**Jv-viii**). In Tukey HSD results, conditions are abbreviated, A: DMSO, B: DMSO:PITC, C: DMSO+TFA, D: DMSO:PITC+TFA, E: DMSO:PITC+TFA+trypsin.

**ANOVA Results (**Fig. 4**Jv-viii)** maximum fluorescence

| | Sum of Squares | df | F | p-value |
|---|---|---|---|---|
| Group | 1.096e+09 | 4.0 | 16.4998 | 0.000211 |
| Residual | 1.661e+08 | 10.0 | — | — |

**Tukey HSD Results (**Fig. 4**Jv-viii)** maximum fluorescence

| Group 1 | Group 2 | Mean Diff | p-adj | Lower | Upper | Reject |
|---|---|---|---|---|---|---|
| A | B | −15094.57 | 0.0075 | −26047.44 | −4141.70 | True |
| A | C | 5291.34 | 0.5344 | −5661.53 | 16244.21 | False |
| A | D | −10391.64 | 0.0649 | −21344.51 | 561.23 | False |
| A | E | −16430.00 | 0.0042 | −27382.87 | −5477.13 | True |
| B | C | 20385.91 | 0.0008 | 9433.04 | 31338.78 | True |
| B | D | 4702.93 | 0.6337 | −6249.94 | 15655.80 | False |
| B | E | −1335.43 | 0.9936 | −12288.30 | 9617.44 | False |

| | | | | | | |
|---|---|---|---|---|---|---|
| C | D | −15682.98 | 0.0058 | −26635.85 | −4730.11 | True |
| C | E | −21721.34 | 0.0005 | −32674.21 | −10768.47 | True |
| D | E | −6038.36 | 0.4168 | −16991.23 | 4914.51 | False |

(v) Raw **average** fluorescence intensity values after ClpS2 St-V1 with anti-HA staining of $F_1Y_2L_3$ peptide throughout in-gel Edman degradation conditions for each gel, average, and standard deviation, over the replicates for Fig. 4**Jv-viii**.

| **Condition name** | **Average intensity ExMre gel 1** | **Average intensity ExMre gel 2** | **Average intensity ExMre gel 3** | **Average intensity over replicates** | **Standard deviation over replicates** |
|---|---|---|---|---|---|
| (1) DMSO | 10031.64473 | 6872.36635 | 7141.00606 | **8015.00571** | 1751.61825 |
| (2) PITC:DMSO | 1817.34818 | 1679.35423 | 2016.83522 | **1837.84587** | 169.67166 |
| (3) DMSO+TFA | 8671.97730 | 10578.90190 | 9101.80781 | **9450.89567** | 1000.24353 |
| (4) PITC:DMSO+TFA | 5403.85749 | 6232.71530 | 5524.65901 | **5720.410599** | 447.76147 |
| (5) PITC:DMSO+TFA +trypsin | 1418.05203 | 1450.39934 | 1184.51938 | **1350.99025** | 145.07240 |

(vi) One-way Analysis of Variance (ANOVA) and Tukey's post-hoc Honestly Significant Difference (HSD) test (FWER = 0.05) on the abundance of **average** fluorescence intensity in the gel (Fig. 4**Jv-viii**). In Tukey HSD results, conditions are abbreviated, A: DMSO, B: DMSO:PITC, C: DMSO+TFA, D: DMSO:PITC+TFA, E: DMSO:PITC+TFA+trypsin.

**ANOVA Results** (Fig. 4**Jv-viii**) average fluorescence

| | **Sum of Squares** | **df** | **F** | **p-value** |
|---|---|---|---|---|
| Group | 1.571e+08 | 4.0 | 45.4587 | 0.000002 |
| Residual | 8.638e+06 | 10.0 | — | — |

**Tukey HSD Results** (Fig. 4**Jv-viii**) average fluorescence

| **Group 1** | **Group 2** | **Mean Diff** | **p-adj** | **Lower** | **Upper** | **Reject** |
|---|---|---|---|---|---|---|
| A | B | −6177.16 | 0.0001 | −8674.62 | −3679.70 | True |
| A | C | 1435.89 | 0.3796 | −1061.57 | 3933.35 | False |
| A | D | −2294.59 | 0.0756 | −4792.05 | 202.87 | False |
| A | E | −6664.02 | 0.0 | −9161.48 | −4166.56 | True |
| B | C | 7613.05 | 0.0 | 5115.59 | 10110.51 | True |
| B | D | 3882.57 | 0.0032 | 1385.11 | 6380.03 | True |
| B | E | −486.86 | 0.9644 | −2984.32 | 2010.60 | False |
| C | D | −3730.48 | 0.0043 | −6227.94 | −1233.02 | True |

| | | | | | | |
|---|---|---|---|---|---|---|
| C | E | −8099.91 | 0.0 | −10597.37 | −5602.45 | True |
| D | E | −4369.42 | 0.0013 | −6866.88 | −1871.96 | True |

#### Supplementary Table 10

a)

(i) Raw fluorescence intensity values for the binding assay with biotin-ClickP-aa in ExMre gels with streptavidin using Glyphic-V (Fig. 6**Ei-iii**), including the average intensity for each gel, the average and standard deviation over the replicates.

| Condition name | Average intensity ExMre gel 1 | Average intensity ExMre gel 2 | Average intensity ExMre gel 3 | Average intensity over replicates | Standard deviation over replicates |
|---|---|---|---|---|---|
| Phenylalanine | 166 | 171 | 141 | 159 | 16 |
| Glycine | 169 | 170 | 148 | 162 | 12 |
| Valine | 1,615 | 1,425 | 1,275 | 1,438 | 170 |

| Fold change | |
|---|---|
| Val/Phe | 9.03 |
| Val/Gly | 8.86 |

(ii) One-way Analysis of Variance (ANOVA) and Tukey's post-hoc Honestly Significant Difference (HSD) test (FWER = 0.05) on the abundance of average fluorescence intensity in the gel (Fig. 6**Ei-iii**). In Tukey HSD results, conditions are abbreviated, A: phenylalanine, B: glycine, C: valine.

**ANOVA results**

| | Sum of Squares | df | F | p-value |
|---|---|---|---|---|
| Group | 3264026.0 | 2.0 | 166.2718 | 0.000006 |
| Residual | 58892.0 | 6.0 | — | — |

**Tukey HSD**

| Group 1 | Group 2 | Mean Diff | p-adj | Lower | Upper | Reject |
|---|---|---|---|---|---|---|
| A | B | 3.0 | 0.9992 | −245.20 | 251.20 | False |
| A | C | 1279.0 | 0.0 | 1030.80 | 1527.20 | True |
| B | C | 1276.0 | 0.0 | 1027.80 | 1524.20 | True |

b) As in (a), but for Fig. 6**Eiv-vi**

(i)

| Condition name | Average intensity ExMre gel 1 | Average intensity ExMre gel 2 | Average intensity ExMre gel 3 | Average intensity over replicates | Standard deviation over replicates |
|---|---|---|---|---|---|
| Phenylalanine | 163 | 226 | 266 | 219 | 52 |
| Acetyl-lysine | 171 | 137 | 146 | 151 | 18 |

| | | | | | |
|---|---|---|---|---|---|
| Leucine | 547 | 3,439 | 1,986 | 1,991 | 1,446 |
| Isoleucine | 3,161 | 4,096 | 2,677 | 3,311 | 721 |
| Valine | 3,057 | 3,937 | 4,026 | 3,673 | 535 |
| Streptavidin only | 143 | 133 | 138 | 138 | 5 |

(ii) One-way Analysis of Variance (ANOVA) and Tukey's post-hoc Honestly Significant Difference (HSD) test (FWER = 0.05) on the abundance of average fluorescence intensity in the gel (Fig. 6**Eiv-vi**). In Tukey HSD results, conditions are abbreviated, A: phenylalanine, B: acetyl-lysine, C: leucine, D: isoleucine, E: valine, F: streptavidin only.

**ANOVA results (Fig. 6Eiv-vi)**

| | Sum of Squares | df | F | p-value |
|---|---|---|---|---|
| Group | 4.057e+07 | 5.0 | 16.7801 | 0.000047 |
| Residual | 5.802e+06 | 12.0 | — | — |

**Tukey HSD (Fig. 6Eiv-vi)**

| Group 1 | Group 2 | Mean Diff | p-adj | Lower | Upper | Reject |
|---|---|---|---|---|---|---|
| A | B | −67.0 | 1.0 | −1974.07 | 1840.07 | False |
| A | C | 1772.33 | 0.0742 | −134.73 | 3679.40 | False |
| A | D | 3093.0 | 0.0016 | 1185.93 | 5000.07 | True |
| A | E | 3455.0 | 0.0006 | 1547.93 | 5362.07 | True |
| A | F | −80.33 | 1.0 | −1987.40 | 1826.73 | False |
| B | C | 1839.33 | 0.061 | −67.73 | 3746.40 | False |
| B | D | 3160.0 | 0.0013 | 1252.93 | 5067.07 | True |
| B | E | 3522.0 | 0.0005 | 1614.93 | 5429.07 | True |
| B | F | −13.33 | 1.0 | −1920.40 | 1893.73 | False |
| C | D | 1320.67 | 0.2558 | −586.40 | 3227.73 | False |
| C | E | 1682.67 | 0.0961 | −224.40 | 3589.73 | False |
| C | F | −1852.67 | 0.0587 | −3759.73 | 54.40 | False |
| D | E | 362.0 | 0.9856 | −1545.07 | 2269.07 | False |
| D | F | −3173.33 | 0.0013 | −5080.40 | −1266.27 | True |
| E | F | −3535.33 | 0.0005 | −5442.40 | −1628.27 | True |

c) As in (a), but for Fig. 6**Fi-iii**

(i)

| **Condition name** | **Average intensity ExMre gel 1** | **Average intensity ExMre gel 2** | **Average intensity ExMre gel 3** | **Average intensity over replicates** | **Standard deviation over replicates** |
|---|---|---|---|---|---|

| | | | | | |
|---|---|---|---|---|---|
| Phenylalanine | 371 | 299 | 413 | 361 | 58 |
| Glycine | 114 | 117 | 119 | 117 | 3 |
| Valine | 138 | 145 | 142 | 142 | 4 |

| **Fold change** | |
|---|---|
| Phe/Gly | 3.09 |
| Phe/Val | 2.55 |

(ii) One-way Analysis of Variance (ANOVA) and Tukey's post-hoc Honestly Significant Difference (HSD) test (FWER = 0.05) on the abundance of average fluorescence intensity in the gel (**Fig. 6Fi-iii**). In Tukey HSD results, conditions are abbreviated, A: phenylalanine, B: glycine, C: valine.

**ANOVA results (Fig. 6Fi-iii)**

| | Sum of Squares | df | F | p-value |
|---|---|---|---|---|
| Group | 108430.889 | 2.0 | 48.6577 | 0.000196 |
| Residual | 6685.333 | 6.0 | — | — |

**Tukey HSD (Fig. 6Fi-iii)**

| Group 1 | Group 2 | Mean Diff | p-adj | Lower | Upper | Reject |
|---|---|---|---|---|---|---|
| A | B | −244.33 | 0.0003 | −327.96 | −160.71 | True |
| A | C | −219.33 | 0.0005 | −302.96 | −135.71 | True |
| B | C | 25.0 | 0.6502 | −58.62 | 108.62 | False |

d) As in (c), but for **Fig. 6Fiv-vi**

(i)

| Condition name | Average intensity ExMre gel 1 | Average intensity ExMre gel 2 | Average intensity ExMre gel 3 | Average intensity over replicates | Standard deviation over replicates |
|---|---|---|---|---|---|
| Phenylalanine | 4071 | 3882 | 2235 | 3396 | 1010 |
| Acetyl-lysine | 141 | 126 | 132 | 133 | 8 |
| Leucine | 392 | 491 | 441 | 441 | 50 |
| Isoleucine | 287 | 205 | 470 | 321 | 135 |
| Valine | 213 | 228 | 226 | 222 | 8 |
| Streptavidin only | 152 | 126 | 129 | 136 | 14 |

(ii) One-way Analysis of Variance (ANOVA) and Tukey's post-hoc Honestly Significant Difference (HSD) test (FWER = 0.05) on the abundance of average fluorescence intensity in the gel (**Fig. 6Fiv-vi**). In Tukey HSD results, conditions are abbreviated, A: phenylalanine, B: acetyl-lysine, C: leucine, D: isoleucine, E: valine, F: streptavidin only.

**ANOVA results (Fig. 6Fiv-vi)**

| | Sum of Squares | df | F | p-value |
|---|---|---|---|---|
| Group | 2.494e+07 | 5.0 | 28.7492 | 0.000003 |
| Residual | 2.082e+06 | 12.0 | — | — |

**Tukey HSD (Fig. 6Fiv-vi)**

| Group 1 | Group 2 | Mean Diff | p-adj | Lower | Upper | Reject |
|---|---|---|---|---|---|---|
| A | B | −3263.0 | 0.0 | −4405.39 | −2120.61 | True |
| A | C | −2954.67 | 0.0 | −4097.06 | −1812.28 | True |
| A | D | −3075.33 | 0.0 | −4217.72 | −1932.94 | True |
| A | E | −3173.67 | 0.0 | −4316.06 | −2031.28 | True |
| A | F | −3260.33 | 0.0 | −4402.72 | −2117.94 | True |
| B | C | 308.33 | 0.9375 | −834.06 | 1450.72 | False |
| B | D | 187.67 | 0.9925 | −954.72 | 1330.06 | False |
| B | E | 89.33 | 0.9998 | −1053.06 | 1231.72 | False |
| B | F | 2.67 | 1.0 | −1139.72 | 1145.06 | False |
| C | D | −120.67 | 0.9991 | −1263.06 | 1021.72 | False |
| C | E | −219.0 | 0.985 | −1361.39 | 923.39 | False |
| C | F | −305.67 | 0.9395 | −1448.06 | 836.72 | False |
| D | E | −98.33 | 0.9996 | −1240.72 | 1044.06 | False |
| D | F | −185.0 | 0.9929 | −1327.39 | 957.39 | False |
| E | F | −86.67 | 0.9998 | −1229.06 | 1055.72 | False |

e) As in (c), but for Fig. 6G

| Condition name | Average intensity ExMre gel 1 | Average intensity ExMre gel 2 | Average intensity ExMre gel 3 | Average intensity over replicates | Standard deviation over replicates |
|---|---|---|---|---|---|
| Phenylalanine | 290 | 105 | 303 | 233 | 111 |
| Acetyl-lysine | 141 | 124 | 123 | 129 | 10 |
| Leucine | 176 | 125 | 131 | 144 | 28 |
| Isoleucine | 4,300 | 6,314 | 5,352 | 5,322 | 1,008 |
| Valine | 208 | 243 | 194 | 215 | 25 |
| Streptavidin only | 123 | 122 | N/A | 123 | 1 |

**ANOVA Results (Fig. 6G)**

| | Sum of Squares | df | F | p-value |
|---|---|---|---|---|
| Group | 6.556e+07 | 5.0 | 70.1123 | 5.758e-08 |
| Residual | 2.057e+06 | 11.0 | — | — |

**Tukey HSD (Fig. 6G)**

| Group 1 | Group 2 | Mean Diff | p-adj | Lower | Upper | Reject |
|---|---|---|---|---|---|---|
| A | B | −103.33 | 0.9996 | −1307.47 | 1100.81 | False |
| A | C | −88.67 | 0.9998 | −1292.81 | 1115.47 | False |
| A | D | 5089.33 | 0.0 | 3885.19 | 6293.47 | True |
| A | E | −17.67 | 1.0 | −1221.81 | 1186.47 | False |
| A | F | −110.17 | 0.9997 | −1456.43 | 1236.10 | False |
| B | C | 14.67 | 1.0 | −1189.47 | 1218.81 | False |
| B | D | 5192.67 | 0.0 | 3988.53 | 6396.81 | True |
| B | E | 85.67 | 0.9998 | −1118.47 | 1289.81 | False |
| B | F | −6.83 | 1.0 | −1353.10 | 1339.43 | False |
| C | D | 5178.0 | 0.0 | 3973.86 | 6382.14 | True |
| C | E | 71.0 | 0.9999 | −1133.14 | 1275.14 | False |
| C | F | −21.5 | 1.0 | −1367.77 | 1324.77 | False |
| D | E | −5107.0 | 0.0 | −6311.14 | −3902.86 | True |
| D | F | −5199.5 | 0.0 | −6545.77 | −3853.23 | True |
| E | F | −92.5 | 0.9999 | −1438.77 | 1253.77 | False |

f) As in (c), but for Fig. 6H

| Condition name | Average intensity ExMre gel 1 | Average intensity ExMre gel 2 | Average intensity ExMre gel 3 | Average intensity over replicates | Standard deviation over replicates |
|---|---|---|---|---|---|
| Phenylalanine | 223 | 401 | 756 | 460 | 271 |
| Acetyl-lysine | 146 | 126 | 191 | 155 | 33 |
| Leucine | 2310 | 3766 | 3435 | 3,170 | 763 |
| Isoleucine | 2523 | 4157 | 2196 | 2,959 | 1050 |
| Valine | 2239 | 2189 | 2076 | 2,168 | 83 |
| Streptavidin only | 166 | 122 | 125 | 137 | 25 |

**ANOVA Results (Fig. 6H)**

| | Sum of Squares | df | F | p-value |
|---|---|---|---|---|

| | | | | |
|---|---|---|---|---|
| **Group** | 3.034e+07 | 5.0 | 20.5832 | 0.000017 |
| **Residual** | 3.537e+06 | 12.0 | — | — |

**Tukey HSD (Fig. 6H)**

| Group 1 | Group 2 | Mean Diff | p-adj | Lower | Upper | Reject |
|---|---|---|---|---|---|---|
| A | B | −305.67 | 0.9797 | −1794.65 | 1183.32 | False |
| A | C | 2710.33 | 0.0006 | 1221.35 | 4199.32 | True |
| A | D | 2498.67 | 0.0012 | 1009.68 | 3987.65 | True |
| A | E | 1708.0 | 0.0217 | 219.02 | 3196.98 | True |
| A | F | −322.33 | 0.9746 | −1811.32 | 1166.65 | False |
| B | C | 3016.0 | 0.0002 | 1527.02 | 4504.98 | True |
| B | D | 2804.33 | 0.0004 | 1315.35 | 4293.32 | True |
| B | E | 2013.67 | 0.0068 | 524.68 | 3502.65 | True |
| B | F | −16.67 | 1.0 | −1505.65 | 1472.32 | False |
| C | D | −211.67 | 0.9961 | −1700.65 | 1277.32 | False |
| C | E | −1002.33 | 0.2802 | −2491.32 | 486.65 | False |
| C | F | −3032.67 | 0.0002 | −4521.65 | −1543.68 | True |
| D | E | −790.67 | 0.5092 | −2279.65 | 698.32 | False |
| D | F | −2821.0 | 0.0004 | −4309.98 | −1332.02 | True |
| E | F | −2030.33 | 0.0064 | −3519.32 | −541.35 | True |

g) As in (c), but for Fig. 6I

| Condition name | Average intensity ExMre gel 1 | Average intensity ExMre gel 2 | Average intensity ExMre gel 3 | Average intensity over replicates | Standard deviation over replicates |
|---|---|---|---|---|---|
| Phenylalanine | 387 | 303 | 239 | 310 | 74 |
| Acetyl-lysine | 2557 | 2454 | 2258 | 2423 | 152 |
| Leucine | 269 | 194 | 275 | 246 | 45 |
| Isoleucine | 397 | 218 | 454 | 356 | 123 |
| Valine | 363 | 205 | 262 | 277 | 80 |
| Streptavidin only | 454 | 286 | 283 | 341 | 98 |

**ANOVA Results (Fig. 6I)**

| | Sum of Squares | df | F | p-value |
|---|---|---|---|---|
| Group | 1.123e+07 | 5.0 | 218.1816 | 2.399e-11 |

| | | | | |
|---|---|---|---|---|
| Residual | 1.235e+05 | 12.0 | — | — |

**Tukey HSD (Fig. 6I)**

| Group 1 | Group 2 | Mean Diff | p-adj | Lower | Upper | Reject |
|---|---|---|---|---|---|---|
| A | B | 2113.33 | 0.0 | 1835.08 | 2391.59 | True |
| A | C | −63.67 | 0.9679 | −341.92 | 214.59 | False |
| A | D | 46.67 | 0.9917 | −231.59 | 324.92 | False |
| A | E | −33.0 | 0.9984 | −311.25 | 245.25 | False |
| A | F | 31.33 | 0.9987 | −246.92 | 309.59 | False |
| B | C | −2177.0 | 0.0 | −2455.25 | −1898.75 | True |
| B | D | −2066.67 | 0.0 | −2344.92 | −1788.41 | True |
| B | E | −2146.33 | 0.0 | −2424.59 | −1868.08 | True |
| B | F | −2082.0 | 0.0 | −2360.25 | −1803.75 | True |
| C | D | 110.33 | 0.7633 | −167.92 | 388.59 | False |
| C | E | 30.67 | 0.9988 | −247.59 | 308.92 | False |
| C | F | 95.0 | 0.8527 | −183.25 | 373.25 | False |
| D | E | −79.67 | 0.9216 | −357.92 | 198.59 | False |
| D | F | −15.33 | 1.0 | −293.59 | 262.92 | False |
| E | F | 64.33 | 0.9665 | −213.92 | 342.59 | False |

#### Supplementary Table 11

Table recording the highest ratio of known oxidation peptide species to original peptide in the ExM gels (descriptive values of **Supplementary Figure 8**).

| | **Highest ratio of oxidized species to original peptide (oxidized/non-oxidized)** | |
|---|---|---|
| **Methionine** | 13817063/3628434.7 | 3.80800 |
| **Cysteine** | 299666.7/54427.54 | 5.50579 |
| **Tyrosine** | 30242.02/2509423.3 | 0.01205 |
| **Phenylalanine** | 1166.65/18738042 | 0.00006 |
| **Tryptophan** | 108509.7/2119325.3 | 0.05120 |
| **Histidine** | 30999.35/441471.91 | 0.07022 |
| **Proline** | 294169.1/5514152.5 | 0.05335 |
| **Arginine** | 6285.23/4906487.6 | 0.00128 |

#### Supplementary Table 12

This table relates to **Supplementary Figure 17** and the kinetic parameters of the binders used in the simulation of *in situ* protein sequencing. The following table shows the binding probabilities for different specificity regimes at a 1 μM binder concentration reaching equilibrium and a subsequent wash time of 30 min. The ratio α represents the ratio between $k_{off}$ of the on-target versus the off-target.

The bound probability for the on-target (correct) is $p_{correct} = \left(\frac{[C]}{[C] + K_d^{on-target}}\right) \cdot e^{-k_{off}^{on-target} t_{wash}}$, the unbound probability for the on-target is $p_{correct-unbound} = 1 - p_{correct}$, whereas the bound probability for off-target (incorrect) is $p_{incorrect} = \left(\frac{[C]}{[C] + K_d^{off-target}}\right) \cdot e^{-k_{off}^{off-target} t_{wash}}$, and the unbound probability for the off-target is $p_{incorrect-unbound} = 1 - p_{incorrect}$. The $k_{off}$ ($s^{-1}$) is the inverse of the dwell time, and $t_{wash}$ (min) is the wash time. The $k_{on}$ value is assumed constant across all binders with a value of $10^6$.

| Specificity | Ratio (α) | Bound probability | Unbound probability | Concentration of binder (μM) | Wash Time (min) | k_on ($M^{-1}$ $s^{-1}$) | k_off ($s^{-1}$) | k_D (M) |
|---|---|---|---|---|---|---|---|---|
| Perfect | ∞ | 1 | 0 | - | - | - | - | - |
| Perfect (off-target) | ∞ | 0 | 1 | - | - | - | - | - |
| Very High | 50,000 | 0.9139 | 0.0861 | 1 | 30 | $10^6$ | $5*10^{-5}$ | $5*10^{-11}$ |
| Very High (off-target) | 50,000 | ~0 | ~1 | 1 | 30 | $10^6$ | 2.5 | $2.5*10^{-6}$ |
| High | 5000 | 0.9139 | 0.0861 | 1 | 30 | $10^6$ | $5*10^{-5}$ | $5*10^{-11}$ |
| High (off-target) | 5000 | ~0 | ~1 | 1 | 30 | $10^6$ | 0.25 | $2.5*10^{-7}$ |
| Medium | 500 | 0.9139 | 0.0861 | 1 | 30 | $10^6$ | $5*10^{-5}$ | $5*10^{-11}$ |
| Medium (off-target) | 500 | ~0 | ~1 | 1 | 30 | $10^6$ | $2.5*10^{-2}$ | $2.5*10^{-8}$ |
| Low | 50 | 0.9139 | 0.0861 | 1 | 30 | $10^6$ | $5*10^{-5}$ | $5*10^{-11}$ |
| Low (off-target) | 50 | 0.0111 | 0.9890 | 1 | 30 | $10^6$ | $2.5*10^{-3}$ | $2.5*10^{-9}$ |
| Very Low | 5 | 0.9139 | 0.0861 | 1 | 30 | $10^6$ | $5*10^{-5}$ | $5*10^{-11}$ |
| Very Low (off-target) | 5 | 0.6375 | 0.3625 | 1 | 30 | $10^6$ | $2.5*10^{-4}$ | $2.5*10^{-10}$ |

#### Supplementary Table 13

| Expansion factor | Theoretical protein size after expansion* | Microscope | Theoretical resolution (including dividing by expansion factor) | Imaging depth |
|---|---|---|---|---|
| ~ 300X | 3 µm | Confocal microscopy (e.g., Nikon CSU-W1 with SoRa) | ~ 0.8 nm | < ~ 200 µm (depends on the lens) |
| ~ 100X | 1 µm | Nikon CSU-W1 SoRa with TIRF | ~ 0.5 nm (with dSTORM) | HILO:<br>< 10 um |
| ~ 20X | 0.2 µm | iPALM | ~ 1 nm | < 0.75 µm |

* Assuming ~ 10 nm sized protein

#### REFERENCES

1. Hoshi, T. & Heinemann, S. H. Regulation of cell function by methionine oxidation and reduction. *J. Physiol.* **531**, 1–11 (2001).
2. Jortzik, E., Wang, L. & Becker, K. Thiol-Based Posttranslational Modifications in Parasites. *Antioxid. Redox Signal.* **17**, 657–673 (2011).
3. Giulivi, C., Traaseth, N. J. & Davies, K. J. A. Tyrosine oxidation products: analysis and biological relevance. *Amino Acids* **25**, 227–232 (2003).
4. Galano, A. & Cruz-Torres, A. OH radical reactions with phenylalanine in free and peptide forms. *Org. Biomol. Chem.* **6**, 732–738 (2008).
5. Todorovski, T., Fedorova, M., Hennig, L. & Hoffmann, R. Synthesis of peptides containing 5-hydroxytryptophan, oxindolylalanine, N-formylkynurenine and kynurenine. *J. Pept. Sci.* **17**, 256–262 (2011).
6. Uchida, K. Histidine and lysine as targets of oxidative modification. *Amino Acids* **25**, 249–257 (2003).
7. Xu, G. & Chance, M. R. Hydroxyl Radical-Mediated Modification of Proteins as Probes for Structural Proteomics. *Chem. Rev.* **107**, 3514–3543 (2007).
8. Phang, J. M., Liu, W. & Zabirnyk, O. Proline Metabolism and Microenvironmental Stress. *Annu. Rev. Nutr.* **30**, 441–463 (2010).
9. Petrov, D., Daura, X. & Zagrovic, B. Effect of Oxidative Damage on the Stability and Dimerization of Superoxide Dismutase 1. *Biophys. J.* **110**, 1499–1509 (2016).
10. Tayeh, M. A. & Marletta, M. A. Macrophage Oxidation of L-Arginine to Nitric Oxide, Nitrite, and Nitrate: Tetrahydrobiopterin is required as a cofactor*. *J. Biol. Chem.* **264**, 19654–19658 (1989).
11. Wang, S. *et al.* Single-shot 20-fold expansion microscopy. *Nat. Methods* 1–7 (2024).
12. Edman, P., Högfeldt, E., Sillén, L. G. & Kinell, P. O. Method for Determination of the Amino Acid Sequence in Peptides. *Acta Chemica Scandinavica* **4**, 283–293 (1950).

13. Edman, P. & Begg, G. A Protein Sequenator. *Eur. J. Biochem.* **1**, 80–91 (1967).
14. Kostić, M. D., Mihajlović, K. & Divac, V. M. Kinetic Study of the Pyridine-Catalyzed Selenolactonization of 4-Pentenoic Acid. *Catal. Letters* **150**, 2076–2081 (2020).
15. Laursen, R. A. [27] Automatic solid-phase Edman degradation. in *Methods in Enzymology* vol. 25 344–359 (Academic Press, 1972).
16. Laursen, R. A. Solid-phase Edman degradation. An automatic peptide sequencer. *Eur. J. Biochem.* **20**, 89–102 (1971).
17. Swaminathan, J. *et al.* Highly parallel single-molecule identification of proteins in zeptomole-scale mixtures. *Nat. Biotechnol.* 10.1038/nbt.4278 (2018).
18. Boehnert, M. & Schlesinger, D. H. Improved manual sequential analysis of peptides. *Anal. Biochem.* **96**, 469–473 (1979).
19. Eriksson, S. & Sjöquist, J. Quantitative determination of N-terminal amino acids in some serum proteins. *Biochim. Biophys. Acta* **45**, 290–296 (1960).
20. Sullivan, S. & Wong, T. W. A manual sequencing method for identification of phosphorylated amino acids in phosphopeptides. *Anal. Biochem.* **197**, 65–68 (1991).
21. Tarr, G. E. A general procedure for the manual sequencing of small quantities of peptides. *Anal. Biochem.* **63**, 361–370 (1975).
22. Zimmerman, C. L., Appella, E. & Pisano, J. J. Rapid analysis of amino acid phenylthiohydantoins by high-performance liquid chromatography. *Anal. Biochem.* **77**, 569–573 (1977).
23. Pramanik, B. C., Hinton, S. M., Millington, D. S., Dourdeville, T. A. & Slaughter, C. A. Analysis of phenylthiohydantoin amino acid mixtures for sequencing by thermospray liquid chromatography/mass spectrometry. *Anal. Biochem.* **175**, 305–318 (1988).
24. Matsudaira, P. Sequence from picomole quantities of proteins electroblotted onto polyvinylidene difluoride membranes. *J. Biol. Chem.* **262**, 10035–10038 (1987).
25. Iida, T., Santa, T., Toriba, A. & Imai, K. Semi-automatic amino acid sequencing and D/L-configuration determination of peptides with detection of liberated N-terminal

phenylthiocarbamoylamino acids. *Analyst* **123**, 2829–2834 (1998).

26. Matsunaga, H. *et al.* Proton: a major factor for the racemization and the dehydration at the cyclization/cleavage stage in the Edman sequencing method. *Anal. Chem.* **68**, 2850–2856 (1996).
27. Zheng, L., Sun, Y., Eisenstein, M. & Soh, H. T. Peptide sequencing via reverse translation of peptides into DNA. *bioRxiv* (2024) doi:10.1101/2024.05.31.596913.
28. Mustățea, G., Ungureanu, E. & Iorga, E. Protein acidic hydrolysis for amino acids analysis in food-progress over time : A short review. (2019).
29. Giera, M., Aisporna, A., Uritboonthai, W. & Siuzdak, G. The hidden impact of in-source fragmentation in metabolic and chemical mass spectrometry data interpretation. *Nat. Metab.* **6**, 1647–1648 (2024).
30. van der Rest, G., He, F., Emmett, M. R., Marshall, A. G. & Gaskell, S. J. Gas-phase cleavage of PTC-derivatized electrosprayed tryptic peptides in an FT-ICR trapped-ion cell: mass-based protein identification without liquid chromatographic separation. *J. Am. Soc. Mass Spectrom.* **12**, 288–295 (2001).
31. Reed, B. D. *et al.* Real-time dynamic single-molecule protein sequencing on an integrated semiconductor device. *Science* (2022) doi:10.1126/science.abo7651.
32. Tullman, J., Callahan, N., Ellington, B., Kelman, Z. & Marino, J. P. Engineering ClpS for selective and enhanced N-terminal amino acid binding. *Appl. Microbiol. Biotechnol.* **103**, 2621–2633 (2019).
33. Tullman, J., Christensen, M., Kelman, Z. & Marino, J. P. A ClpS-based N-terminal amino acid binding reagent with improved thermostability and selectivity. *Biochem. Eng. J.* **154**, 107438 (2020).
34. Alon, S. *et al.* Expansion sequencing: Spatially precise in situ transcriptomics in intact biological systems. *Science* **371**, eaax2656 (2021).
35. Chen, F. *et al.* Nanoscale Imaging of RNA with Expansion Microscopy. *Nat. Methods* **13**, 679–684 (2016-8).

36. Stein, B. J., Grant, R. A., Sauer, R. T. & Baker, T. A. Structural Basis of an N-Degron Adaptor with More Stringent Specificity. *Structure* **24**, 232–242 (2016).
37. Callahan, N., Siegall, W. B., Bergonzo, C., Marino, J. P. & Kelman, Z. Contributions from ClpS surface residues in modulating N-terminal peptide binding and their implications for NAAB development. *Protein Eng. Des. Sel.* **36**, gzad007 (2023).
38. Sass, H.-J., Musco, G., Stahl, S. J., Wingfield, P. T. & Grzesiek, S. Solution NMR of proteins within polyacrylamide gels: Diffusional properties and residual alignment by mechanical stress or embedding of oriented purple membranes. *J. Biomol. NMR* **18**, 303–309 (2000).
39. Albe, K. R., Butler, M. H. & Wright, B. E. Cellular concentrations of enzymes and their substrates. *J. Theor. Biol.* **143**, 163–195 (1990).
40. Murray, E. *et al.* Simple, Scalable Proteomic Imaging for High-Dimensional Profiling of Intact Systems. *Cell* **163**, 1500–1514 (2015).
41. Crank, J. *The Mathematics of Diffusion. United Kingdom.* (Clarendon Press, 1964).
42. Fischer, H., Polikarpov, I. & Craievich, A. F. Average protein density is a molecular-weight-dependent function. *Protein Sci.* **13**, 2825–2828 (2004).
43. Trnková, L., Dršata, J. & Boušová, I. Oxidation as an important factor of protein damage: Implications for Maillard reaction. *J. Biosci.* **40**, 419–439 (2015).
44. Heinonen, M., Gürbüz, G. & Ertbjerg, P. Chapter 3 - Oxidation of proteins. in *Chemical Changes During Processing and Storage of Foods* (eds. Rodriguez-Amaya, D. B. & Amaya-Farfan, J.) 85–123 (Academic Press, 2021).
45. Williams, B. J., Barlow, C. K., Kmiec, K. L., Russell, W. K. & Russell, D. H. Negative ion fragmentation of cysteic acid containing peptides: cysteic acid as a fixed negative charge. *J. Am. Soc. Mass Spectrom.* **22**, 1622–1630 (2011).
46. Gao, R. *et al.* A highly homogeneous polymer composed of tetrahedron-like monomers for high-isotropy expansion microscopy. *Nat. Nanotechnol.* **16**, 698–707 (2021).
47. Lee, H., Yu, C.-C., Boyden, E. S., Zhuang, X. & Kosuri, P. Tetra-gel enables superior

accuracy in combined super-resolution imaging and expansion microscopy. *Sci. Rep.* **11**, 16944 (2021).

48. Yao, Y., Docter, M., van Ginkel, J., Ridder, D. de & Joo, C. Single-molecule protein sequencing through fingerprinting: computational assessment. *Phys. Biol.* **12**, 055003 (2015).
49. Swaminathan, J., Boulgakov, A. A. & Marcotte, E. M. A Theoretical Justification for Single Molecule Peptide Sequencing. *PLoS Comput. Biol.* **11**, e1004080 (2015).
50. Ohayon, S., Girsault, A., Nasser, M., Shen-Orr, S. & Meller, A. Simulation of single-protein nanopore sensing shows feasibility for whole-proteome identification. *PLoS Comput. Biol.* **15**, e1007067 (2019).
51. Smith, M. B., Simpson, Z. B. & Marcotte, E. M. Amino acid sequence assignment from single molecule peptide sequencing data using a two-stage classifier. *PLoS Comput. Biol.* **19**, e1011157 (2023).
52. Mapes, J. H. *et al.* Robust and scalable single-molecule protein sequencing with fluorosequencing. *bioRxivorg* (2023) doi:10.1101/2023.09.15.558007.
53. Smith, M. B. *et al.* Estimating error rates for single molecule protein sequencing experiments. *PLoS Comput. Biol.* **20**, e1012258 (2024).
54. Pham, T. L. H. *et al.* Peptide sequencing via protein language models. *arXiv [q-bio.BM]* (2024) doi:10.48550/arXiv.2408.00892.
55. Shaib, A. H. *et al.* One-step nanoscale expansion microscopy reveals individual protein shapes. *Nat. Biotechnol.* 1–9 (2024).
56. Thavarajah, R., Mudimbaimannar, V. K., Elizabeth, J., Rao, U. K. & Ranganathan, K. Chemical and physical basics of routine formaldehyde fixation. *J. Oral Maxillofac. Pathol.* **16**, 400–405 (2012).
57. Asano, S. M. *et al.* Expansion Microscopy: Protocols for Imaging Proteins and RNA in Cells and Tissues. *Curr. Protoc. Cell Biol.* **80**, e56 (2018-9).
58. Gambarotto, D. *et al.* Imaging cellular ultrastructures using expansion microscopy

(U-ExM). *Nat. Methods* **16**, 71–74 (2019).

59. Tillberg, P. W. *et al.* Protein-retention expansion microscopy of cells and tissues labeled using standard fluorescent proteins and antibodies. *Nat. Biotechnol.* **34**, 987–992 (2016).
60. Truckenbrodt, S., Sommer, C., Rizzoli, S. O. & Danzl, J. G. A practical guide to optimization in X10 expansion microscopy. *Nat. Protoc.* **14**, 832–863 (2019).
61. Metz, B. *et al.* Identification of formaldehyde-induced modifications in proteins: reactions with model peptides. *J. Biol. Chem.* **279**, 6235–6243 (2004).
62. Bucur, O. *et al.* Nanoscale imaging of clinical specimens using conventional and rapid-expansion pathology. *Nat. Protoc.* **15**, 1649–1672 (2020).
63. Schnell, U., Dijk, F., Sjollema, K. A. & Giepmans, B. N. G. Immunolabeling artifacts and the need for live-cell imaging. *Nat. Methods* **9**, 152–158 (2012).
64. Chung, K. *et al.* Structural and molecular interrogation of intact biological systems. *Nature* **497**, 332–337 (2013).
65. Klimas, A. *et al.* Magnify is a universal molecular anchoring strategy for expansion microscopy. *Nat. Biotechnol.* **41**, 858–869 (2023).
66. Ku, T. *et al.* Multiplexed and scalable super-resolution imaging of three-dimensional protein localization in size-adjustable tissues. *Nat. Biotechnol.* **34**, 973–981 (2016-9).
67. Cui, Y. *et al.* Expansion microscopy using a single anchor molecule for high-yield multiplexed imaging of proteins and RNAs. *PLoS One* **18**, e0291506 (2023).
68. Keil, B. *Specificity of Proteolysis*. (Springer, Berlin, Germany, 2012).
69. Morihara, K. & Tsuzuki, H. Specificity of proteinase K from Tritirachium album limber for synthetic peptides. *Agricultural and biological chemistry* **39**, 1489–1492 (1975).
70. Saenger, W. Proteinase K. in *Handbook of Proteolytic Enzymes* 3240–3242 (Elsevier, 2013).
71. Yu, C.-C. J. *et al.* Expansion microscopy of C. elegans. *Elife* **9**, (2020).
72. Wang, U.-T. T. *et al.* Protein and lipid expansion microscopy with trypsin and tyramide

signal amplification for 3D imaging. *Sci. Rep.* **13**, 21922 (2023).

73. Kang, J. *et al.* Multiplexed expansion revealing for imaging multiprotein nanostructures in healthy and diseased brain. *Nat. Commun.* **15**, 9722 (2024).
74. Valdes, P. A. *et al.* Improved immunostaining of nanostructures and cells in human brain specimens through expansion-mediated protein decrowding. *Sci. Transl. Med.* **16**, eabo0049 (2024).
75. Olsen, J. V., Ong, S.-E. & Mann, M. Trypsin cleaves exclusively C-terminal to arginine and lysine residues. *Mol. Cell. Proteomics* **3**, 608–614 (2004).
76. Raijmakers, R., Neerincx, P., Mohammed, S. & Heck, A. J. R. Cleavage specificities of the brother and sister proteases Lys-C and Lys-N. *Chem. Commun. (Camb.)* **46**, 8827–8829 (2010).
77. Ebeling, W. *et al.* Proteinase K from Tritirachium album limber. *Eur. J. Biochem.* **47**, 91–97 (1974).
78. Bonissone, S., Gupta, N., Romine, M., Bradshaw, R. A. & Pevzner, P. A. N-terminal Protein Processing: A Comparative Proteogenomic Analysis. *Mol. Cell. Proteomics* **12**, 14–28 (2013-1).
79. Deng, S. & Marmorstein, R. Protein N-terminal acetylation: Structural basis, mechanism, versatility, and regulation. *Trends Biochem. Sci.* **46**, 15–27 (2021).
80. Saveliev, S. *et al.* Trypsin/Lys-C protease mix for enhanced protein mass spectrometry analysis. *Nat. Methods* **10**, i–ii (2013).
81. Kashina, A. S. & Yates, J. R., Iii. Analysis of arginylated peptides by subtractive Edman degradation. *Methods Mol. Biol.* **2620**, 153–155 (2023).
82. Khoury, G. A., Baliban, R. C. & Floudas, C. A. Proteome-wide post-translational modification statistics: frequency analysis and curation of the swiss-prot database. *Sci. Rep.* **1**, 90 (2011).
83. Landry, J. P., Ke, Y., Yu, G.-L. & Zhu, X. D. Measuring affinity constants of 1450 monoclonal antibodies to peptide targets with a microarray-based label-free assay

platform. *J. Immunol. Methods* **417**, 86–96 (2015).
